# Integration of differential expression under drought with gene family expansion unique to drought tolerant species predicts candidate genes for drought adaptation in Brassicaceae species

**DOI:** 10.1101/2025.01.07.631653

**Authors:** Carolin Uebermuth-Feldhaus, Heiko Schoof

**Affiliations:** INRES Crop Bioinformatics, University of Bonn, Katzenbugweg 2, 53115 Bonn, Germany

**Keywords:** Adaptation, drought, comparative genomics, gene family expansion, differential expression, evolution, Brassicaceae, diversifying selection, abiotic stress resistance, bioinformatics

## Abstract

**Background:** In order to predict candidate genes for drought adaptation, we analyze genomic data of the more drought resistant Brassicaceae species *Eutrema salsugineum* and *Arabidopsis lyrata* compared to more drought sensitive species *Arabidopsis thaliana* and *Brassica napus.* We combine gene family expansion, which is an important driver of evolution in plants, unique to the drought resistant species with differential expression under drought (DE).

**Results:** We show that combining trait-specific gene family expansion with differential expression predicts a concise set of candidate genes. To demonstrate that these are relevant for drought adaptation in tolerant species we show enrichment of DE conserved between both tolerant species, DE unique to the tolerant species, and up regulation. We show that specific functions are enriched, and that the set contains genes with functions such as root development in line with drought adaptation based on evidence from other species, while the background of all differentially expressed genes (DEGs) contains many general stress reaction genes. Whereas DEGs in general are rarely under diversifying selection, signatures of diversifying selection are slightly enriched in the candidate gene families, highly significantly enriched in DEGs in tolerant species-specific expanded gene families, and in contrast not enriched in DEGs in sensitive species-specific expanded gene families.

**Conclusions:** Our approach identifies a concise and functionally relevant set of candidate genes for drought adaptation with promising targets for functional studies and crop improvement for drought tolerance. We propose that our method can also be used to predict candidate genes for adaptation to other environmental factors.

## Background

Drought stress experiments often identify thousands of differentially expressed genes and many of them are general stress genes ([1], [2], [3]). Moreover, drought reactions of plants include effects which are unfavorable for crop production, such as the reduction of photosynthesis ([4], [5], [6], [7]), because they reduce biomass accumulation. The goal of this study is to prioritize genes which function in the adaptation to drought from all genes which react to drought. For this purpose, we combine gene family expansion unique to drought resistant plant species with differential expression under drought (DE).

Gene family expansion through whole genome duplication (WGD) is common in plants ([8], [9]). Gene duplicates can acquire new functions (neofunctionalization, including new regulation patterns) or split their functions (subfunctionalization) (reviewed in e.g. [10]). Examples of experimentally confirmed duplicates which confer abiotic stress tolerance in *Arabidopsis thaliana* are 1) a duplication in a E3 ubiquitin ligase, which is responsible for drought tolerance [11], and 2) a duplication of a mitochondrial GrpE (Mge) protein which is responsible for heat stress tolerance [12]. In *Eutrema salsugineum,* there are gene family expansions which were experimentally shown to be relevant for its salt tolerance: 1) a duplication in a calcium sensor (*ATCBL10*, AT4G33000) in *Eutrema salsugineum* (EUTSA_v10026019mg and EUTSA_v10028908mg) is important for calcium-signaling and increased salt tolerance in *Arabidopsis thaliana* mutants [13], and 2) HKT1 (AT4G10310) is duplicated and relevant for salt tolerance in *Eutrema salsugineum* [14].

Differential expression upon drought can link a gene duplication to a function under drought stress. The authors of different studies have pointed out expanded gene families among differentially expressed genes, e.g. in a salt tolerant *Populus* species under salt treatment [15], in a desiccation tolerant *Lindernia* species under drought and recovery [16] or in roots of soy beans during nodulation [17]. The relationship between differential expression under abiotic stress and gene family expansion has been systematically described: 30 % of duplicated genes in *Eutrema salsugineum* and 26-27 % of duplicated genes in *Arabidopsis thaliana* respond to abiotic stresses [18], and tandem duplicates in the cold tolerant grass *Lolium perenne* responded significantly more (17.48 %) often to cold treatment than all genes (9.91 %) [19].

A duplication of the NICOTIANAMINE SYNTHASE3 (AT1G09240) in *Arabidopsis lyrata* is propose as candidate gene for stress resistance, as it has likely neofunctionalized, i.e. acquired expression in a different tissue and under drought and salt stress [20]. Laha and colleagues [21] review the research on stress genomics focused on gene family expansion in *Brassicaceae* species and discuss approaches to identify stress-responsive duplicates. They propose a strategy to identify stress-responsive allelic variants for the engineering of stress tolerant plants by characterizing stress-responsive genes retained as multiple copies in tolerant species, while present only as a single copy in the susceptible *Arabidopsis thaliana*. Oh and Dassanayake [22] state that it is important to perform comparative studies on species with natural tolerance to abiotic stress because it is not possible to identify stress tolerance relevant expression and physiological reactions in species which are sensitive to this stress.

In this study, we apply a generalizable approach to identify trait-specific evolutionary adaptations [23] to drought tolerance in Brassicaceae. We identify gene family expansion in the drought tolerant Brassicaceae species *Arabidopsis lyrata* (*Aly*) and *Eutrema salsugineum* (*Esa*) compared to the more drought sensitive *Arabidopsis thaliana* (*Ath*) and *Brassica napus* (*Bna*) (Figure 1) and combine this with genes which are differentially expressed under drought. *Arabidopsis lyrata* is a biennial or perennial which naturally grows on dry soils. Bouzid and colleagues [24] observed drought avoidance and tolerance mechanisms in *Aly:* The leaf water content decreased slower before wilting and the photosynthetic capacity was kept high at wilting compared to *Ath*. Additionally, *Aly* plants had a higher survival rate and showed less damage from wilting at rewatering ([25], [24]). *Aly* reacts earlier to drought than *Esa* and *Ath* with transcriptional change and growth reduction [25] and wilting compared to *Ath* [24]. Marín-de la Rosa and colleagues [25] showed that the stomata in *Aly* are less open already under normal conditions compared to *Ath* and they close less upon ABA treatment. Congruent with this, several ABA receptors are constantly higher expressed. They observed this “stress-aware” state in *Aly* and *Esa*. *Eutrema salsugineum* (syn: *Thellungiella salsuginea*) is a halophytic model plant which also shows tolerance towards several other stresses, including drought tolerance (reviewed in [26]). Traits related to drought tolerance in *Esa* include morphological plasticity depending on growth conditions and an increased accumulation of solutes ([27]). *Esa* shows an increased prolin content and a constantly high expression of the prolin biosynthesis gene P5CS1, even under well watered conditions [25]. This together with the limited opening of stomata, described above for *Aly,* suggests that *Esa* has several “stress-awareness” mechanisms. *Esa* can maintain the plant water content even under drought conditions [26]. Further, *Esa* responds later with growth reduction, i.e. at a lower soil water content, than *Aly* and *Ath* [25]. Its survival rate after rewatering is higher than that of *Ath* and equal to that of *Aly* [25], probably due to several tolerance mechanisms, some of which are described above. *Arabidopsis thaliana* is an annual plant which can escape drought by quickly terminating its life cycle. *Ath* also shows some drought avoidance mechanisms, e.g. *Ath* was able to maintain thicker leaves and started wilting later than *Aly* [24]. But the survival rate of *Ath* after wilting is much lower than that of *Aly* and *Esa* ([24], [25]), which is eminently important for crop production, i.e. yield stability. *Brassica napus* is an annual or biennial oil seed crop. It is an allopolyploid species which contains the A and C *Brassica* genome with 2n = 4x = 38 [28]. Its yield is highly reduced by drought [29]

**Figure 1:**
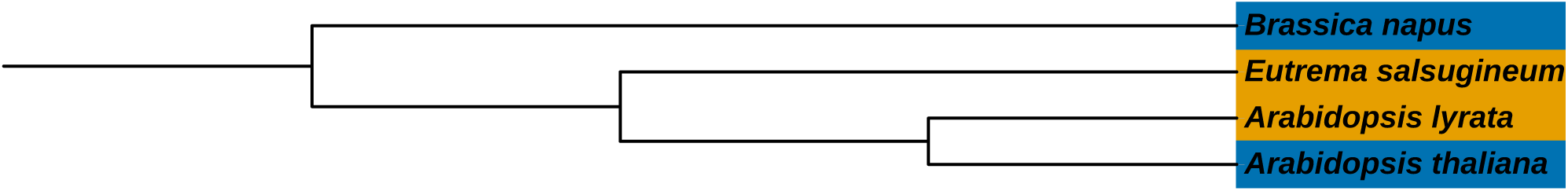
Phylogenetic topology of the Brassicaceae species in this study as calculated by OrthoFinder [30]. Orange marks the drought tolerant and blue marks the drought sensitive species.

Knowledge about the genetic basis of drought adaptation is relevant for breeding of drought resistant crops. While there are many indicators that neo- or subfunctionalized gene duplicates play an important role in adaptation to abiotic stresses, the knowledge about gene family expansion which confers drought resistance to adapted Brassicaceae species is low. We took advantage of these indicators to predict candidate genes for drought resistance in plants from the general drought-responsive genes using our recently published automated workflow [23]. For this we integrate publicly available transcriptomic and genomic data from the tolerant and the sensitive Brassicaceae species. We systematically describe the relationship between gene family expansion and differential expression under drought with focus on the drought tolerant species. We evaluate the prediction of candidate genes by identifying enrichment with signatures of diversifying selection.

## Results

### Protein coding gene families in four Brassicaceae species

We assigned the genes of proteins with a minimum length of 100 amino acids (aa) from the four Brassicaceae species *Eutrema salsugineum (Esa)*, *Arabidopsis lyrata (Aly)*, *Arabidopsis thaliana (Ath)* and *Brassica napus* (*Bna*) to gene families (orthologous groups, HOGs) using OrthoFinder ([30], version 2.5.4) in order to perform comparative genome analyses. There are 17,488 gene families which are conserved between the four species (Supplementary Figure 2, Additional File 1). These conserved gene families contain 19,990 genes from *Esa*, 21,173 genes from *Aly*, 19,919 genes from *Ath* and 38,861 genes from *Bna.* There are considerably less HOGs in all other combinations, e.g. species-specific HOGs or HOGs with representatives of only some species, less than 1000 with the exception of *Bna* specific HOGs (7,573). All analyses in this study were performed on the set of HOGs which are conserved between all four species (Conserved Set, 17,488 HOGs, *Conserved_dataset_matrix.tsv*, Additional File 4), all other HOGs were not analyzed further.

### Upon drought, differential expression as well as regulation are conserved between the four Brassicaceae species

We inferred differentially expressed genes (DEGs) between drought and control from publicly available leaf transcriptomes from all four species in this study (p-adj. <= 0.1, see Methods). In the Conserved Set, 13,531 HOGs (77.4 %) contain at least one DEG. There are 7,790 DEGs from *Esa* in 7,434 HOGs (42.5 %). In *Aly*, there are 9,478 DEGs in 8,866 HOGs (50.7 %). In *Ath*, there are 3,141 DEGs in 2,865 HOGs (16.4 %). In *Bna*, there are 10,065 DEGs in 6,956 HOGs (39.8 %) (background pie charts in Supplementary Figure 4, Additional File 1).

In order to find out if the differential expression (DE) is conserved between the four species or rather species specific, we compared the frequency of HOGs with DEG from all four species and HOGs with DEG from none of the species to the frequencies expected from the distribution of HOGs with DEG per species. Using a hypergeometric test for over representation at significance level 0.05, we found significantly more HOGs than expected which contain DEG from all four species (observed: 846/17488 = 4.8 %, expected = 245 based on *p_s4_*= 1.4 %, p < 2.2e-16, see Methods) as well as significantly more HOGs with no DEG from any of the four species (observed: 3957/17488 = 22.6 %, expected: 2500 based on *p_f4_* = 14.3 %, p < 2.2e-16). We conclude that differential expression upon drought is mostly conserved between the four species.

We then wanted to know how genes are predominantly regulated, i.e. if the reaction to drought is manifested rather by up or down regulation and if the regulation is rather conserved or if it differs between the species. For this purpose, we categorized the DE gene families with at least 2 DEGs regarding their pattern of regulation, where “up” means that all DEGs are up regulated, “down” means that all DEGs are down regulated and “up_and_down” means that there are up regulated as well as down regulated genes in the HOG. The regulation within the HOGs with at least two DEGs in the Conserved Set is equally distributed among the regulation categories (2768 only up, 2935 only down and 3097 gene families with up- and down regulated genes). Hence, in 2/3 of the HOGs which have more than one DEG, these genes are equally regulated. In only 1/3 of the tested HOGs, the genes were regulated in opposing directions (light blue columns in Supplementary Figure 5, Additional File 1). We found that HOGs which show contrasting regulation between the species mostly involve DE in *Bna* (Supplementary Figure 5,, Additional File 1, highlighted with asterisks). There is also more within-species divergent regulation in *Bna* compared to the other species (down_and_up in *Bna*: 230 HOGs, *Aly* 76 HOGs, *Esa* 49 HOGs and *Ath* 6 HOGs, Supplementary Figure 5, Additional File 1).

### Gene family expansion

From the gene families which are conserved between the four species we identified 727 gene families which are expanded in the drought tolerant species *Esa* compared to the drought sensitive species *Ath* and *Bna*, and 1075 gene families which are expanded in *Aly* compared to both sensitive species. 144 of these gene family expansions in *Esa* or *Aly* are common to both tolerant species. As for the drought sensitive species, 375 gene families are expanded in *Ath* compared to the drought tolerant species *Esa* and *Aly*, and 1825 gene families are expanded in *Bna* compared to both tolerant species. 58 of these gene family expansions in *Ath* or *Bna* are common to both sensitive species.

When considering the lineage-specific duplications, we identified 449 gene families which are commonly expanded in *Esa* and both *Arabidopsis* species compared to *Bna* and 1762 gene families which are expanded in *Bna* compared to *Esa* and both *Arabidopsis* species. 441 gene families are expanded in both *Arabidopsis* species compared to *Bna* and *Esa* and 103 gene families are commonly expanded in *Bna* and *Esa* compared to both *Arabidopsis* species.

### Differential expression under drought in expanded gene families in tolerant and sensitive species

Next we analyzed the differential expression (DE) of genes in expanded gene families. We observe that DE in drought tolerant species correlates with gene family expansion unique to drought tolerant species. The gene families which are expanded in *Esa* but not in any of the sensitive species are significantly more often differentially expressed in *Esa* than expected from the distribution of DEGs in the Conserved Set (363/727 vs. 7434/17488, hypergeometric test for over representation p = 3.2e-05, Supplementary Figure 4 c, Additional File 1). The same applies for DE and expansion in *Aly* (613/1075 vs. 8866/17488, p = 1.0e-05, Supplementary Figure 4 d, Additional File 1), but also for DE in a drought sensitive species in gene families which are expanded in a drought sensitive species (*Ath:* 87/375 vs. 2865/17488, p = 0.00035, *Bna:* 969/1825 vs. 6956/17488, p = 3.8e-34, Supplementary Figure 4 g and h, Additional File 1). Gene families which are expanded in both species with the same tolerance compared to the species with contrasting drought tolerance are slightly, but not significantly, enriched for DE of both expanded species (tolerant species 47/144 vs. 4638/17488, p = 0.06, sensitive species 6/58 vs. 1436/17488, p = 0.34, Supplementary Figure 4 b and f, Additional File 1). We conclude that duplicate genes in general are more often DE.

Thus, we asked if there is a significant enrichment of conserved DE, that is, genes from both species are differentially expressed, in the expanded gene families. Of the 93 HOGs expanded in both tolerant with DE in at least one of the tolerant species, 47 HOGs (50 %) show DE in both tolerant species (Supplementary Figure 4 a and b, Additional File 1). This is significantly more than expected from the background, where 4,638 (40 %) from 11,662 HOGs which are differentially expressed in any of the two tolerant species are differentially expressed in both tolerant species (p = 0.022). In contrast, of the 39 HOGs expanded in both sensitive species with DE in at least one of the sensitive species, only 6 HOGs (15 %) show DE in both sensitive species (Supplementary Figure 4 e and f, Additional File 1). This is slightly less than in the background, where from 8,385 HOGs which are differentially expressed in any sensitive species, 1,436 HOGs (17 %) are differentially expressed in both sensitive species. Thus, conserved DE is significantly enriched only in gene families expanded in both drought tolerant species, but not in gene families expanded in both sensitive species.

Then, we analyzed the direction of regulation in the expanded gene families, i.e. whether up or down regulation under drought is more frequent (see above) and if this varies between the expanded species. We found that in the gene families which are expanded and differentially expressed in the tolerant species, a significantly larger proportion contain only up regulated genes (51 %, p = 3.9e-3 in both, 38 %, p = 5e-4 in *Esa* and 39 %, p = 4e-4 in *Aly,* Figure 2 a-c) when compared to all gene families which show DE in the respective species, where all three regulation categories are equally frequent (Figure 2 a-c background pie charts). In contrast to this, the HOGs expanded in both sensitive species more often show divergent, both up and down, regulation (67 % HOGs). While this is not significantly more than in the background of all HOGs with DEGs from both sensitive species (Figure 2 d, p = 0.195), the sample is only four out of six gene families expanded in both sensitive species. Looking at the sensitive species individually, the HOGs expanded in *Ath* show significant enrichment with divergent regulation compared to all HOGs with DEG from *Ath* (39 %, p = 0.015, Figure 2 e), while they are not significantly enriched for only up regulation (32 %, p = 0.195, Figure 2 e). The regulation categories of HOGs expanded in *Bna* are distributed similarly to those in all HOGs with DEG from *Bna*: nearly half (47 %, Figure 2 f) of HOGs show up and down regulation.

**Figure 2:**
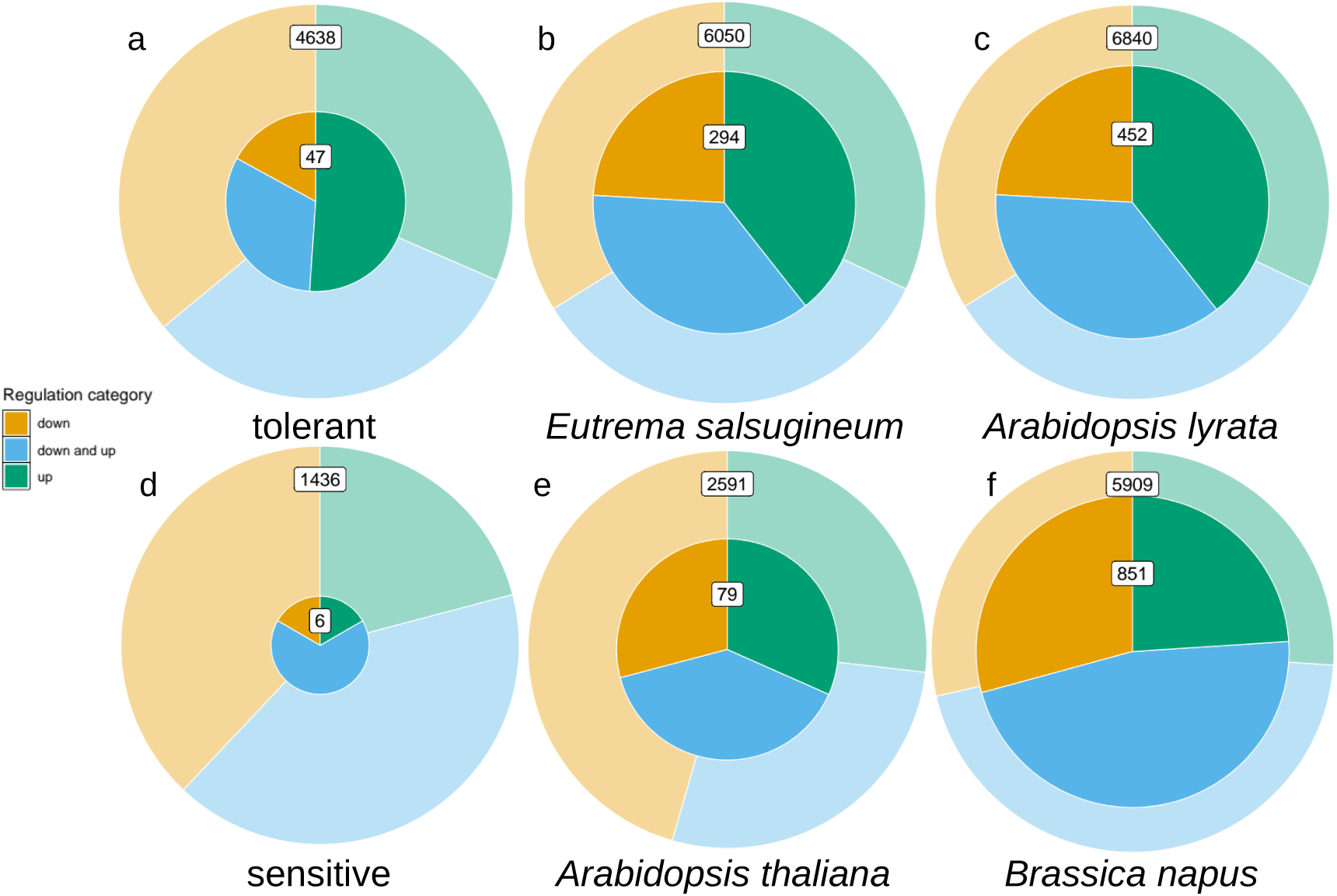
Pie charts of the proportions of gene families per regulation category. Green = “up”, Orange = “down”, Blue = “down_and_up”. Each pie chart represents all gene families with DEGs from the depicted species in the Conserved Set (background) and in its subset (foreground) of gene families expanded in the depicted species. Only gene families with at least two DEGs regardless of the species are considered.

To summarize, in tolerant species, but not in sensitive species we observe enrichment of gene families with a) conserved differential expression and b) specifically only up regulation in expanded families vs. the background.

Interestingly and in contrast to the observation that differential expression is likely to be conserved across species within a gene family, in the HOGs expanded only in tolerant species, we find that most genes are differentially expressed only in the tolerant species but not the sensitive species: We found that 28 (59 %) out of 47 HOGs (shown in blue in Figure 3) which are expanded and show DE in both tolerant species do not show DE in any of the sensitive species (Figure 3). This is significantly more (p = 4.6e-4) than in the background, where only 1,621 (35 %) of 4,638 HOGs which show DE in both tolerant species are not differentially expressed in the sensitive species (Supplementary Figure 3, Additional File 1).

**Figure 3:**
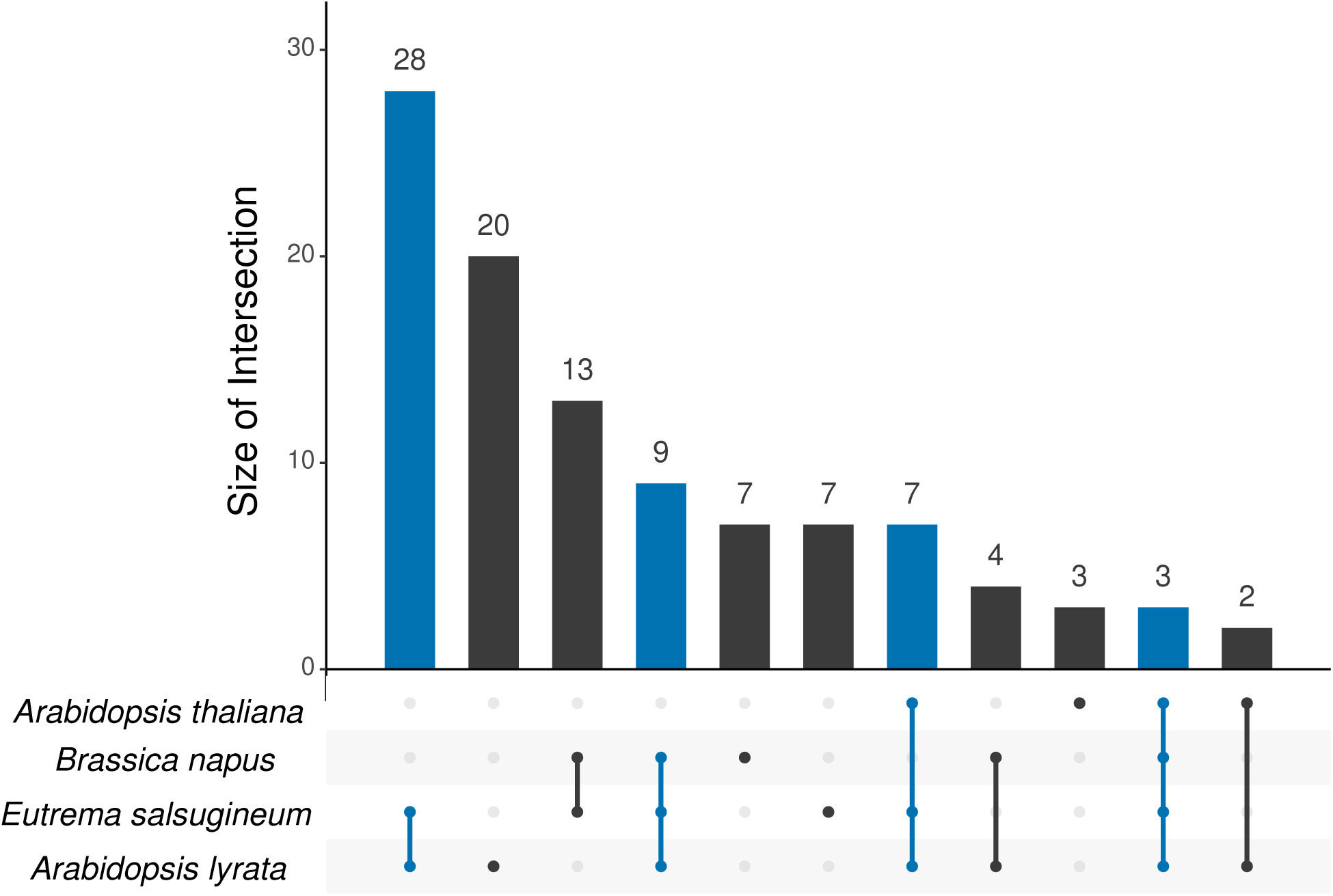
UpSet plot showing all gene families expanded in both tolerant species that contain a DEG. The intersections of gene families with DEG from the given species are shown, where blue marks intersections with both tolerant species. The species are ordered by the number of gene families, with the largest set on the bottom.

Given their strong correlations to the drought tolerance phenotype we define the gene families which are commonly expanded in the tolerant species and differentially expressed under drought in at least one of the tolerant species as candidate genes for drought adaptation. We put special focus on two subsets, a) genes which show conserved differential expression in both tolerant species and b) out of these those showing only up regulation and no DE in sensitive species. Information on these candidate gene families, including regulation under drought of the genes from all four species, are provided in *Candidate_gene_families_and_tolerant_specific_expansions.xlsx*, Additional File 2, where the corresponding subsets are indicated.

### Functions of candidate gene families for drought adaptation

We explored the functions of the predicted candidate gene families for drought adaptation based on function prediction generated by AHRD, compared them to the functions of all gene families in the Conserved Set as well as all DE gene families and tested for functional enrichment. In addition, we explored the functions of gene families which are expanded and differentially expressed only in one tolerant species, i.e. in *Esa* only or in *Aly* only (see Supplement). The candidate gene families as well as the tolerant species-specific expansions are listed in *Candidate_gene_families_and_tolerant_specific_expansions.xlsx*, Additional File 2, see above, including their predicted functions (human readable descriptions, HRD).

The set of candidate gene families has diverse functions. Only the biological processes “modification-dependent protein catabolic process” and “mRNA splicing, via spliceosome” were significantly enriched in the set of 93 candidate gene families (Supplementary Table 3, Additional File 1). In its subset of 47 candidate gene families which are DE in both tolerant species, the biological processes “modification-dependent protein catabolic process” and “translation” were significantly over represented (Supplementary Table 4, Additional File 1). Interestingly, neither “modification-dependent protein catabolic process” nor any related post translational regulatory process was enriched in the background of all HOGs which show DE in any or in both tolerant species (Supplementary Figure 7 and 8, Additional File 1). Candidate gene families which function in the enriched processes are described in detail in the Supplementary Results, Additional File 1. In the following, we focus on the candidate gene families which are uniquely up regulated in both tolerant species. Because of the small sample size of 17 gene families, we have not tested for enriched functions but display all functions in Supplementary Figure 12, Additional File 1. We inspected all 17 gene families manually and included these notes in *Candidate_gene_families_and_tolerant_specific_expansions.xlsx*, Additional File 2.

The following examples show an interesting relation to drought tolerance. N0.HOG0007350 (Lysine-specific demethylase REF6, Supplementary Figure 13, Additional File 1), which functions in “positive regulation of lateral root development” (GO:1901333) and “ABA catabolism” (GO:0046345), promotes lateral root formation in *Ath* ([31]) and is involved in the regulation of flowering ([32]). N0.HOG0008546 (AT3G16630) encodes a Kinesin-like protein KIN-13A (Supplementary Figure 14, Additional File 1) which functions in trichome morphogenesis ([33]) and in the formation of secondary cell wall pits ([34]).

There are several gene families which function in regulatory processes: N0.HOG0007005 encodes transcription initiation factor TFIID subunit 9 (AT1G54140, *TAF9*), which functions in “DNA-templated transcription initiation” (GO:0006352). It is duplicated in both tolerant species and highly up regulated in *Aly* (Supplementary Figure 15, Additional File 1). N0.HOG0005205 (AT3G49000), DNA-directed RNA polymerase III subunit RPC3, functions in “DNA-templated transcription” (GO:0006351). Moreover, two gene families function in epigenetic transcription regulation: 1) the highly expanded N0.HOG0005443, which is most likely a family of Histone-lysine N-methyltransferases (AT5G47150). One duplicate from *Aly* is up regulated and shows signatures of diversifying selection. Both duplicates from *Esa* in this gene family are also up regulated. 2) The above mentioned Lysine-specific demethylase REF6, which functions in “positive regulation of lateral root development”. Gene family N0.HOG0008832 is a family of E3 UFM1-protein ligase 1 homologs and regulates protein homeostasis (Supplementary Figure 9, Additional File 1). One of the up regulated duplicates from *Esa* is under positive selection. It functions in “response to endoplasmic reticulum stress”, “protein K69-linked ufmylation” and “regulation of proteasomal ubiquitin-dependent protein catabolic process”.

### Diversifying selection is enriched in candidate gene families for drought adaptation

We tested the genes within the conserved gene families for signatures of diversifying selection in their coding sequences by performing an exploratory analysis using the adaptive branch site model absrel [35]. 72 % of the conserved genes from *Esa* (14,412 / 19,990), 71 % from *Aly* (14,974 / 21,173), 72 % from *Ath* (14,408 / 19,919) and 72 % from *Bna* (27,962 / 38,861) passed the quality filter criteria (see Methods) and were tested (*OGs_pos_selection_pvalues.tsv*, Additional File 4). From these, 1,371 (9.5 %) genes from *Esa*, 1,423 (9.5 %) genes from *Aly,* 1,165 (8.1 %) genes from *Ath,* and 2,732 (9.8 %) genes from *Bna* show signatures of diversifying selection on significance level 0.05. We used the signatures of diversifying selection as a marker for an increased probability of a gene to be relevant for adaptation. To use this marker to evaluate the prediction of candidate genes for drought adaptation, we tested if the sets of 1) DEGs (Table 1), 2) expanded gene families (Table 2) and 3) differentially expressed and expanded gene families (Table 3), including the candidate gene families for drought adaptation, were enriched or depleted for genes with signatures of diversifying selection using a one-sided hypergeometric test at significance level 0.05. We found that DEGs from all four species are significantly less often under diversifying selection than all genes from the respective species in the Conserved Set (Table 1). We expected that gene duplicates are frequently under diversifying selection because neofunctionalization including positive selection is an important model for the fixation of a gene duplicate in a population. We confirm this hypothesis: duplicated genes of each of the four species, which are not duplicated in the species with contrasting drought tolerance, are significantly more often under diversifying selection compared to all genes of the respective species in the Conserved Set (Table 2). The enrichment is smaller in duplicates from *Bna* compared to duplicates from the other species.

**Table 1:** Number of genes for which diversifying selection was predicted by absrel [35].

| Species | DEGs | Background | $p_{lower}$ |
| --- | --- | --- | --- |
| *Aly* | 544 (8 %) | 1423 (10 %) | 1.6e-15 |
| *Ath* | 133 (6 %) | 1165 (8 %) | 1.1e-05 |
| *Esa* | 489 (8 %) | 1371 (10 %) | 2.6e-06 |
| *Bna* | 596 (8 %) | 2732 (10 %) | 3.5e-14 |

$P_{lower}$ = p-value from hypergeometric test for under representation in the set of differentially expressed genes, compared to the background of all genes from the given species in the Conserved Set.

**Table 2:** Number of genes for which diversifying selection was predicted by absrel [35] in expanded gene families.

| Species | Gene family expansion in | Genes under diversifying selection | Genes under diversifying selection in background | $p_{upper}$ |
| --- | --- | --- | --- | --- |
| *Aly* | *Aly* | 326 (25 %) | 1423 (10 %) | 5.9e-65 |
| ***Aly*** | ***Aly* and *Esa*** | **41 (24 %)** | **1423 (10 %)** | **3.2e-08** |
| *Esa* | *Esa* | 209 (21 %) | 1371 (10 %) | 1.4e-30 |
| ***Esa*** | ***Esa* and *Aly*** | **33 (20 %)** | **1371 (10 %)** | **3.8e-05** |
| *Ath* | *Ath* | 97 (20 %) | 1165 (8 %) | 3.7e-17 |
| ***Ath*** | ***Ath* and *Bna*** | **17 (25 %)** | **1165 (8 %)** | **2.1e-05** |
| *Bna* | *Bna* | 565 (10.4 %) | 2732 (9.8 %) | 0.04 |
| ***Bna*** | ***Bna* and *Ath*** | **21 (15 %)** | **2732 (10 %)** | **0.04** |

For a given species, the number of genes showing diversifying selection is given in families expanded in the given species (upper row, normal letters) or in families expanded in both species with the same tolerance phenotype (lower row, bold letters), but not expanded in species of the opposite tolerance phenotype. $P_{upper}$ = p-value from hypergeometric test for over representation in the set of expanded gene families, compared to the background of all genes from the given species in the Conserved Set.

**Table 3:** Number of differentially expressed genes for which diversifying selection was predicted by absrel [35] in gene families which are expanded in the given species (but not in any species with opposite tolerance phenotype).

| Species | Gene family | DEGs under | DEGs under | $p_{upper}$ |
| --- | --- | --- | --- | --- |

|  | expansion in | diversifying selection | diversifying selection in background |  |
| --- | --- | --- | --- | --- |
| *Aly* | *Aly* | 70 (17 %) | 544 (8%) | 1.3e-10 |
| *Ath* | *Ath* | 6 (10 %) | 133 (6 %) | 0.16 |
| *Esa* | *Esa* | 41 (14 %) | 489 (8 %) | 5.0e-4 |
| *Bna* | *Bna* | 109 (8 %) | 596 (8 %) | 0.29 |
| *Aly* or *Esa* | *Aly* and *Esa* | 12 (13 %) | 1033 (8 %) | 0.06 |
| *Ath* or *Bna* | *Ath* and *Bna* | 4 (8 %) | 729 (7 %) | 0.48 |

$P_{upper}$ = p-value from hypergeometric test for over representation compared to the background of all DEGs from the given species under diversifying selection in the Conserved Set.

Genes duplicated in both species with the same tolerance phenotype, which are not duplicated in the species with opposite drought tolerance, are also enriched for signatures of diversifying selection, as are expansions common to phylogenetically closely related species (Supplementary Table 7, Additional File 1). However, expansions in *Bna* or expansions common to *Bna* and *Esa* are not enriched for signatures of diversifying selection.

In order to evaluate the quality of the predicted candidate genes for drought adaptation we used the enrichment of genes with signatures of diversifying selection as a marker for the relevance of the gene for adaptation (Table 3). In the background of all DEGs from any of the tolerant species, 8 % are under diversifying selection, whereas 12 of 92 testedcandidate gene families (13 %) have a DEG from any of the tolerant species under diversifying selection. However, this enrichment has a low level of significance (p = 0.06). We do not see enrichment in the contrasting set, which are DEGs from any of the sensitive species in gene families expanded in both sensitive species, but not in the tolerant species (8 % vs. 7 %, p = 0.48). As the number of genes under diversifying selection is very low in the candidate and contrasting gene sets, we also analyzed the enrichment in gene families expanded in only one species, but not expanded in the species with opposite drought tolerance phenotype. When looking at DEGs in species-specific gene family expansions, we found that DEGs in families which are expanded in either of the tolerant species, but not in any sensitive species, are highly significantly enriched for DEGs with signatures of diversifying selection (Table 3). This is most prominent for gene family expansions in *Aly*, where 17 % of the DEGs are under diversifying selection, whereas only 8 % of all DEGs from *Aly* are under diversifying selection (p = 1.26e-10). We did not find significant enrichment in DEGs expanded in the sensitive species.

### Candidate gene families with differentially expressed genes under diversifying selection

There are 12 candidate gene families which have a DEG from a tolerant species under diversifying selection (Table 3): 7 of these DEGs are from *Esa* and 5 from *Aly,* while there are no gene families which have a DEG from both tolerant species under diversifying selection. Within the focus set of gene families expanded and specifically up regulated in both tolerant species, we find N0.HOG0008832 (E3 UFM1-protein ligase 1, Supplementary Figure 9, Additional File 1), N0.HOG0007350 (Lysine-specific demethylase REF6, Supplementary Figure 13, Additional File 1) and N0.HOG0005443, which is most likely a family of Histone-lysine N-methyltransferases. More example gene families have been described above or in Supplementary Results, Additional File 1. Detailed information on the candidate gene families with a DEG under positive selection can be found in *Candidate_gene_families_and_tolerant_specific_expansions.xlsx,* Additional File 2. We also describe the functions and examples of DEGs which show signatures of diversifying selection in gene families expanded in either *Esa* or *Aly* in the Supplementary Results, Additional File 1.

## Discussion

We hypothesized that gene family expansion unique to drought tolerant species is a criterion to select from differentially expressed genes (DEGs) those which underlie drought adaptation of the tested species. Gene family expansion is an important driver for the evolution of new functions, and retention of duplicates indicates a selective advantage. Correlation with a tolerance trait can be indicative of successful adaptation.

Using our A2TEA workflow and webapp [23] to combine trait-specific gene family expansion with differential expression (DE) we predict a more concise set of candidate genes. In order to demonstrate that this is a meaningful set we analyze features that are relevant for drought adaptation and show that the candidate gene set is enriched for 1) DE under drought conserved in both tolerant species, 2) DE unique to the tolerant species, and 3) up regulation. Specific functions are enriched, and the set contains genes with functions in line with drought adaptation based on evidence from other species. We show that signatures of diversifying selection are slightly enriched in the candidate gene families (expanded in both tolerant and DE in any tolerant species) and highly significantly enriched in DEGs in tolerant species-specific expanded gene families. Strikingly, they are not enriched in differentially expressed gene families which are expanded in sensitive species.

The prediction of candidate genes in this study is based on the classification of drought resistance of the four analyzed *Brassicaceae* species. Drought resistance involves many processes and phenotypic effects, which complicates its definition (reviewed by [36]). For plant production in areas with periods of drought, a high survivability with a minimum of growth reduction is desirable ([37], [38], [24], [39]). Based on the literature, we classified the extremophytes *Eutrema salsugineum* (*Esa*) and *Arabidopsis lyrata (Aly*) as more drought resistant than their relatives *Arabidopsis thaliana* (*Ath*) and *Brassica napus* (*Bna*) (see Background). A strength of this set is that gene family expansion, one important pillar on which we base our prediction of candidate genes for drought adaptation, is detached from the phylogenetic relationship of the analyzed species, as we have used one tolerant and one sensitive species from the *Brassicaceae* lineages I and II, each ([40]). Thereby, our study most likely identifies old gene family expansions which are derived from whole genome duplication (WGD) in the common ancestor of the Brassicaceae ([41], [42], [43]) but were kept only in the drought tolerant species. This indicates their selective advantage under drought rather than phylogenetic conservation of lineage-specific expansions not related to the trait, such as the 449 lineage-specific expansions in *Esa*, *Aly* and *Ath*. Similarly, the 441 lineage-specific expansions in both *Arabidopsis* species and the 103 lineage-specific expansions in *Esa* and *Bna* are phylogenetically conserved but we cannot relate their conservation to drought adaptation.

We argue that retention of duplicates in both drought resistant species but not sensitive species, even if phylogenetically closer, is indicative of adaptation. Alternatively, two independent duplications in both tolerant species could have led to adaptation through convergent evolution.

### Gene family expansions in DE gene families

Differential expression under drought allows identifying genes which are functionally linked to drought response but not necessarily the result of adaptation. Based on the prediction of orthologous groups (HOGs), we prioritized from the thousands of gene families which are DE under drought hundreds of gene families which are expanded in any of the drought resistant Brassicaceae species. From these we selected 93 candidate gene families which are expanded in both tolerant species and DE in at least one tolerant species.

This prediction of candidate genes for drought adaptation relies on the prediction of HOGs by Orthofinder [30]. Additionally, we used an automated tool, the A2TEA.WebApp [23], to manually evaluate the predicted candidate gene families. This quality control included the visualization of the MSA with the closest related HOGs and the base mean expression per gene in the HOG. We observed several cases where the MSA of gene duplicates was fragmented. This could be due to pseudogenization, or sequencing, assembly, or annotation errors. Nevertheless, we also expect such cases in the background and did not exclude them from enrichment analysis. Furthermore, when manually validating the candidate gene families, we found several HOGs where Orthofinder predicted an expansion only in the tolerant species, but likely failed to include additional gene copies in *Bna*: when adding to the phylogenetic tree the most closely related HOGs, we observed that genes from a HOG composed of exclusively *Bna* genes are probably members of the HOG in question. This means that the predicted expansion in the tolerant species is not exclusive, but also found in a sensitive species. In the complete dataset, there is an unexpectedly large number of HOGs specific for *Bna*, so this may be a frequent prediction error.

We also found discrepancies between the HOGs predicted in our study to the gene families predicted in other studies. For example, Ali and colleagues [14] found gene family expansion in *HKT1* in *Esa* but not in *Ath*. Laha and colleagues [21] found even three homologs to the single Arabidopsis *HKT1* gene At4G10310 in *Esa*. We did not find this gene duplication in the candidate gene families, but instead, we identified *HKT1* from *Ath* in N0.HOG0002756 together with a single homolog from *Aly*, which is up regulated under drought. The three homologs from *Esa* were found separately, each in a HOG with two *Bna* homeologs, (N0.HOG0002757: EUTSA_v10028767mg, N0.HOG0002758: EUTSA_v10028594mg and N0.HOG0002759: EUTSA_v10028595mg, refer to *A2TEA_finished.Rdata,* Additional File 4). So *HKT1* is expanded both in *Esa* and in *Bna*. Thus, as the triplication is also found in the drought sensitive species *Bna*, in our study these are not candidate genes for drought adaptation. We speculate that the neofunctionalization of HKT1 under increased salt concentrations is a special adaptation of *Esa* to salt stress.

In contrast to Das and colleagues [20], who found a duplication in *Aly* in *NICOTINAMINE SYNTHASE3* genes which was not present in *Ath*, we found that the duplication in *NICOTINAMINE SYNTHASE3* genes is present in all four analyzed Brassicaceae species and that the duplicates show divergent regulation under drought in the species in our study (N0.HOG0003383 and N0.HOG0003384, refer to *A2TEA_finished.Rdata,* Additional File 4). Hence, we did not identify this gene family as a candidate gene family for drought adaptation according to our criteria. Nevertheless, the duplicates might be neofunctionalized in all these species.

The definition of orthologous groups is crucial for the reliable prediction of candidate gene families with our criteria. To achieve the best possible reliability, we have compared tolerant and sensitive species of the same family and have used the most recent version of Orthofinder. Our manual inspection allowed us to identify some prediction errors, and the *A2TEA.WebApp* offers a useful visualization for this. However, even though in some cases we come to different conclusions than other studies, we see no indication that the reliability of the candidate prediction is insufficient for statistical evaluation.

We characterized the predicted candidate genes for drought adaptation regarding their patterns of regulation and their biological functions.

Differential expression is more conserved in the candidate gene set. Among the set of 93 candidate gene families we found differential expression under drought in both species more often than expected: 47 (50 %) show differential expression in both species, whereas only 6 (15 %) of the contrasting set (expanded in both sensitive species with differential expression in at least one sensitive species) show differential expression in both. Also with respect to the background set, where 4,638 (40 %) are differentially expressed in both tolerant species, in tolerant species conserved differential expression is significantly enriched in the candidate set: 47 (50 %) show DE in both tolerant species (p = 0.02). We do not find enrichment in the contrasting set. We interpret this conservation to indicate common drought adaptation, which resulted in conserved gene family expansion and regulation under drought in tolerant species, but not sensitive species. This underpins the prediction as candidate gene families for drought adaptation. Interestingly, in the candidate gene families where both species show differentially expressed genes, the direction of differential expression is more often only up regulation and less often only down regulation (Figure 2 a). This is different from the background of all gene families with conserved differential expression in both tolerant species, where up regulation is equally frequent as down regulation. We infer from this that up regulated genes are more likely to have been targets for adaptation to drought. As a result, these duplicates are retained specifically in the drought tolerant species, along with their expression pattern.

A different perspective regards differential expression unique to tolerant species, where genes from sensitive species are not differentially expressed. 35 % of gene families from the Conserved Set which are regulated in both tolerant species are not regulated in the sensitive species, i.e. a majority of gene families shows differential expression also in a sensitive species. This is in line with the observation that differential expression is mostly conserved between species. In contrast, in the candidate gene families the proportion showing unique differential expression, i.e. only in the tolerant species but not in any sensitive species, is significantly increased to 59 % (p = 4.6e-4, Figure 3). This could point to emergence of differential expression under drought as a result of adaptation, which would be cases of neofunctionalization [10], or to the loss of differential expression in the sensitive species.

To summarize, the candidate genes for drought adaptation predicted by our approach, which show DE in both tolerant species, are more often up than down regulated, and mostly uniquely differentially expressed in the tolerant species but not the sensitive species. These observations are consistent with the hypothesis that our approach selects genes that are adaptive for drought.

With respect to gene function, we observe that the selected candidate genes do not show enrichment of functions directly relevant to drought, e.g. general stress response genes or known drought response functions. However, focusing on genes uniquely up regulated in both tolerant species enriches for genes which function in processes such as “plant-type secondary cell wall biogenesis”, “trichome morphogenesis”, “sugar mediated signaling” and “positive regulation of lateral root development”, functions which could reasonably play a role in drought adaptation. It needs to be noted that this excludes genes differentially expressed upon drought in *Ath*. As many functional annotations are deduced from experiments conducted in *Ath,* the genes annotated with drought response functions are mostly differentially expressed upon drought in *Ath*. It is not surprising that the 17 gene families of interest are not annotated with general drought response functions as they are not regulated upon drought in *Ath*. Instead, we conclude that their regulation upon drought is acquired through adaptation in the drought resistant species. This hypothesis is supported by genes from this set that function in processes known to impact drought tolerance such as root development [37] or cell wall organization ([44], [45]) and that are differentially regulated upon drought only in tolerant species.

In contrast to candidate genes uniquely up regulated in both tolerant species, the background of all DEGs from the tolerant species is enriched for many stress response processes such as “response to water deprivation”, “response to salt stress”, “response to cold”. These results confirm that drought response genes are often general stress response genes (see Background). Moreover, the high enrichment of genes which function in “photoinhibition”, “photosynthesis” and “photosynthesis, light harvesting in photosystem I” underpins that drought strongly affects photosynthesis, which can have negative effects on crop yield ([5], [6], [7]). Apparently, by selecting from the DEGs candidate gene families which are expanded and up regulated only in the tolerant species, we differentiate processes which are relevant for drought adaptation from the many general stress responses in all DEGs. We conclude that our approach identifies a concise and functionally relevant set of candidate genes for drought adaptation and allows identifying novel functions not linked to drought response in *Ath*.

We evaluated the prediction of candidate genes for drought adaptation by identifying enrichment with signatures of diversifying selection to underpin a gene’s relevance for adaptation. While the enrichment of diversifying selection in the candidate gene families was not significant (p = 0.06), we could still show that selecting gene family expansion unique to drought tolerant species from DEGs under drought links the selected DEGs to drought adaptation because of the highly significant enrichment of diversifying selection in the DEGs in tolerant species-specific expanded gene families, while there was no enrichment in the DEGs in in sensitive species expanded gene families. The proportion of genes which were tested for diversifying selection was 71 – 72 % for each of the four species, which is why we do not expect biases between the species in the sets of tested genes.

We show that differential expression (DE) under drought is overall not linked to diversifying selection in any of the species. This is in accordance to our observation that DE under drought is conserved between the four species and as such we expect that DEGs are rather under purifying selection. Gene families specifically expanded in both drought tolerant or drought sensitive species are enriched for diversifying selection, thus these genes, while they are probably a result of adaptation, are not specifically adapted to drought. By considering DE under drought, we can link them to an adaptive function under drought. Even though we find a strong correlation of DE and gene family expansion in all four species, diversifying selection is only enriched in gene families which combine these criteria in the tolerant, but not in the sensitive species. This indicates that expansions specific to tolerant species are adaptive for the tolerance trait, whereas expansions specific to sensitive species are adaptive for other traits. So while diversifying selection is not generally linked to DE, it is linked to genes DE under drought that are specifically expanded in drought tolerant plants. We propose that in these genes, the signatures of diversifying selection specifically indicate adaptation to drought. This supports our hypothesis that combining expansion in tolerant species with differential expression under drought selects candidate genes related to adaptation to drought.

There are 14 % and 17 % of DE gene families expanded in *Esa* or in *Aly*, respectively, in which a DEG from the respective drought tolerant species shows signatures of diversifying selection. This also means that there are many expanded gene families in which the DE duplicates have not experienced diversifying selection. These gene duplications were most likely subfunctionalized and fixed by genetic drift according to the DDC model ([46], reviewed in [47] and [10]) and are as relevant for adaptation as the ones which have experienced diversifying selection.

### Discussion of example candidate genes

Among all 93 candidate genes, we identified several processes which are implicated in drought tolerance. We propose that the candidate genes which function in processes whose relevance has not yet been implicated in drought tolerance are likely to be relevant, too. We describe some interesting cases here. We identified the gene family of *CER9* (N0.HOG0006674) among our candidate genes, where one of two *CER9* homologs is up regulated in *Aly*. *CER9* encodes a E3 ubiquitin-ligase with a negative regulatory role in the early cuticle wax and lipid biosynthesis [48]) Further, a mutant which is deficient in *cer9* in *Ath* wilts later [48]. Hence, the up regulation of the gene duplicate in *Aly* might be related to the earlier wilting which Bouzid and colleagues [24] observed in *Aly*. Based on these two findings we speculate that an over expression in *Ath* might lead to earlier wilting. The gene is also duplicated but not significantly regulated in *Esa.* This could be due to the generally later reaction to drought of *Esa* compared to fast-responding *Aly* [25]. *Aly* has mechanisms to tolerate earlier witling, for example keep the photosynthetic rate high under wilting, and can better survive wilting than *Ath* ([25], [24]). Possibly, even if the earlier wilting is a negative trait, it was retained because the up regulated duplicate regulates new targets and influences a positive mechanism for drought tolerance. Actually, the deficiency in *cer9* results in an up regulation of 21 genes which function in protein ubiquitination and degradation [48]. These are potential targets for a diverged regulation by the drought specific up regulated gene duplicate in *Aly*. The *cer9* mutants also showed an increase in very long-chain fatty acids (VLCFAs), and the effects of *cer9* were mostly additive to mutations in Long chain acyl-CoA synthetase 1 (LACS1, CER8) [48], which is a second important component of the early wax and cutin biosynthesis pathway. Mutations in LACS1 more than doubled the amount of VLCFAs in *Ath* [49]. We found LACS1 (N0.HOG0008907) among our candidate genes, where one of the homologs from *Esa* and from *Bna,* each, are down regulated. Even though the increase in VLCFAs has not been shown to be causal for an increased barrier for cuticular water loss, structural differences in cuticular wax composition influence the transpiration ([48], [50]). It is yet unclear, how the duplication of the here described wax biosynthesis genes underlie drought adaptation, but the conservation of the duplication in the tolerant species and their regulation under drought indicate their relevance for the drought tolerance.

Among our candidate gene families which are up regulated uniquely in the tolerant species is a family coding for a Kinesin-like protein (KIN-13A, N0.HOG0008546, AT3G16630) which functions in the morphogenesis of trichomes [33] and in the formation of secondary cell wall pits [34]. In a hot and dry adapted *Shepherdia* hybrid, trichome density was increased under drought and positively influenced light reflectance and leaf temperature regulation under heat [51]. While *Aly* populations with trichomes were more resistant to herbivory compared to populations without trichomes [52], trichome production did not increase tolerance to drought in *Aly* [53]. But the function of KIN-13A in the development of metaxylem could explain the adaptive role of its duplication in both tolerant species: the surface of secondary cell wall pits, areas where no secondary cell wall is deposited, increased when over expressing KIN-13A in *Ath* [34], which could influence the xylem sap transport. We suggest that the duplication and up regulation of KIN-13A in *Aly* and *Esa* is adaptive for drought by altering the secondary cell wall patterns in developing xylem tissues in response to drought, which could potentially increase the xylem sap transport. The formation of secondary cell wall pits depends on the depolymerization of the cortical microtubules by the interaction of KIN-13A with ROP11 and MIDD1 [34]. Interestingly, both KIN-13A, one of two ROP11 (N0.HOG0005842) homologs, and the single MIDD1 (N0.HOG0021372) are up regulated in *Esa* under drought (refer to *A2TEA_finished.Rdata,* Additional File 4). While KIN-13A is duplicated uniquely in the two tolerant species, ROP11 is duplicated in all four species and MIDD1 is not duplicated. It is less clear, how the pathway is regulated in *Aly*, where both KIN-13A homologs are, similarly to *Esa*, up regulated, while both ROP11 (N0.HOG0005842) homologs are down regulated under drought. In our study (genome assembly GCA_000004255.1), we did not find MIDD1 in *Aly*, but an OrthoDB search and an NCBI blast search (GCF_000004255.2) for orthologs to the MIDD1 homolog from *Ath* (AT3G53350) identified a single homolog (110228692) in *Aly* of which we do not have information regarding its expression under drought. These findings suggest that the up regulation of the *KIN-13A-ROP11-MIDD1* pathway is adaptive for drought at least in *Esa*.

Another candidate gene family regulates flowering in *Ath*: The Lysin-specific demethylase REF6 (AT3G48430, N0.HOG0007350) is expanded and up regulated under drought in both tolerant species but in none of in the sensitive species. Both duplicates from *Esa* show signatures of diversifying selection. Over expression of *REF6* in *Ath* lead to early flowering and altered leaf development [54]. It also promotes lateral root formation in *Ath* by demethylation of the repressive H3K27me3 mark on the Chromatin of *PIN1/3/7* [31]. We propose that the over expression of REF6, as we observe it in the up regulation of two homologs in each of the tolerant species, can improve drought tolerance, also in the sensitive species, likely by increasing the root biomass or altering the regulation of flowering.

Among the candidate gene families, transcription factor II D subunit 9 (TAF9, N0.HOG0007005) is uniquely up regulated in both tolerant species. We speculate that the up regulation of both duplicates in the tolerant *Aly* underlies the generally fast [25] and strong reaction of *Aly* to drought, whereas in contrast in *Esa* only one of the gene duplicates is slightly up regulated while the other is not regulated under drought. We found that an additional TFIID subunit (TAF14b, N0.HOG0011415) is expanded and up regulated in *Aly,* while the second homolog is not expressed in this experiment and lacks large parts of N- and C-terminal domains. As loss and gain of domains is common in the evolution of diverged TAFs [55], the diverged gene copies in our study likely have specialized functions. TAF14 is a regulator of FLC [56], and we speculate that its duplication can affect the regulation of flowering under drought.

The described cases require further research to prove their relevance in drought adaptation. We report them here to highlight that further investigations about the unknown candidates will likely be rewarding.

## Conclusions

We predicted a concise set of candidate genes for drought adaptation in two drought tolerant Brassicaceae species using comparative genomics and transcriptomics. DEGs which are expanded in drought tolerant, but not in sensitive Brassicaceae species can be related to drought adaptation more than the background of all genes which react to drought when regarding conserved DE, DE specific to tolerant species, or up-regulation. Based on functional annotation we found candidate genes involved in processes which are relevant for drought adaptation, while the background set of DEGs is enriched for general stress reactions. As an independent evaluation of the relevance of the predicted candidate genes we found that only DEGs which are duplicated uniquely in a tolerant, but not DEGs duplicated uniquely in the sensitive species, are enriched for signatures of diversifying selection. Our method allows identifying both conserved adaptations as well as species-specific adaptations in only one tolerant species, see Supplement.

As our approach is exploratory and not based on predefined groups of genes, the prediction of candidate genes which have not yet been implicated in drought tolerance presents novel but probable hypotheses for further analyses. *In vivo* experiments using mutations or genetic engineering of candidate genes, e.g. following a strategy outlined in [21], will be necessary to demonstrate a role in drought tolerance and give more insights into which molecular mechanisms underlie the drought tolerance traits of *Esa* and *Aly*.

## Methods

### Data

Genomic data from *Eutrema salsugineum* [57], *Arabidopsis lyrata* [58], *Arabidopsis thaliana* [59] and *Brassica napus* [28] were downloaded from ensemblgenomes [60] as listed in Supplementary Table 1, Additional File 1, on June 07 2022. RNA-Seq Data from drought stress studies in *Eutrema salsugineum*, *Arabidopsis lyrata* and *Arabidopsis thaliana* [25] and *Brassica napus* [61] were downloaded from the Sequence Read Archive (SRA, [62]) as listed in Supplementary Table 2, Additional File 1, on March 16 2022. The proteome files were filtered for a minimum sequence length of 100 amino acids using the scripts *filter_min100aa.sh,* Additional File 3 followed by *normalize.sh,* Additional File 3.

### Candidate gene prediction with the *A2TEA.Workflow*

We calculated the differential expression, the orthologous group prediction and the gene family expansion using our recently published automated snakemake workflow *A2TEA.Workflow* [23] commit e6d4aad58e0a685e4e8eec6fc0b7a99521730ea4. Briefly, *A2TEA.Workflow* uses OrthoFinder ([30], version 2.5.4) to predict orthologous groups from proteome files of the species of interest, and selects expanded gene families from the predicted orthologous groups, where the expansion criteria are defined by the user. Additionally, it calculates differential gene expression using DESeq2 [63] from RNA-Seq reads of the same species and combines the information of differential gene expression with the gene family expansion (Supplementary Figure 1 a, Additional File 1).

Raw genomic files (cdna and annotation .gff3) as listed in Supplementary Table 1, Additional File 1, and the filtered proteome files were used as input to the *A2TEA.Workflow*. Functional annotation files from a previous *A2TEA.Workflow* run, which was based on the same input data, were used and depicted in the column “function” in *config/species.tsv*. This functional annotation file had the first two lines (# and empty lines) deleted, as it is stated in the *A2TEA.Workflow README.md* file. Parameters in the *config.yaml* were adjusted to use only the longest isoform when there are several isoforms of one protein (*auto-isoform-filtering=YES*). For the expression quantification, featurecounts was adjusted to quantify on gene level (*transcript_level_quantification=NO).* For the differential expression, a false discovery rate of 0.1 was defined to identify differentially expressed genes between drought and control conditions (*DEG_FDR=0.1).* During the workflow, phylogenetic trees of expanded gene families were calculated with additionally up to five closest related HOGs (*add_OGs=5*) for subsequent visual inspection using the *A2TEA.WebApp* v1.1.5. We used the *A2TEA.Workflow* to identify gene families which were expanded in both of the tolerant species but not in any of the sensitive species (Hypothesis 1), or in each of the tolerant species but not in any of the sensitive species (Hypotheses 2 and 3), and *vice versa* (Hypotheses 4-6 in the *config/hypotheses.tsv*). As *Bna* is a young (∼7500 y.a.) allopolyploid with the A and C *Brassica* genome (2n = 4x = 38, [28]), we account for the doubled number of genes in *Bna* when calculating gene family expansions by setting the column “ploidy” in the *config/species.tsv to “4”*.

The results of the *A2TEA.Workflow*, including differential expression, information about gene family expansion on all gene families, as well as MSA and phylogenetic trees of gene families of each of the tested hypotheses, are in *A2TEA_finished.Rdata*, Additional File 3.

### Analyses

We characterized the predicted candidate genes for drought adaptation regarding their functions, differential expression upon drought, calculated their phylogeny and predicted signatures of diversifying selection (Supplementary Figure 1 b, Additional File 1). The analyses on the differential expression of the gene families and plotting were performed using the R programming language (vers. 4.4, [64]) with the Analysis scripts *Analysis_A2TEA_output.R,*Additional File 3 and *Analysis_expression_pattern_per_OG_only_at_least_2_sig.R*, Additional File 3).

#### Conservation of DE

To test if the DE was more conserved than expected from the distribution of DEGs from each of the species in the Conserved Set, we calculated the probability of success (*p_s_ = s/n)* and failure (*p_f_*= 1 - *p_s_*) of a HOG to contain a DEG per species from the number of HOGs with at least one DEG from species (*s*) in the background of all conserved HOGs (*n* = 17488). We then calculated the probability of a HOG to have a DEG from all 4 species as *p_s4_ = p_sAt_ * p_sAl_ * p_sEs_ * p_sBn_*, and the probability of a HOG not to contain a DEG from any of the four species as *p_f4_ = p_fAt_ * p_fAl_ * p_fEs_ * p_fBn_.* We compared the probabilities to the observed number of HOGs with a DEG from all four species or from none of the four species. We computed the chi-square test using the function *chisq.test()* for each of the two hypotheses on a level of significance alpha = 0.05 (*Analysis_A2TEA_output.R*, Additional File 3).

#### GO term over representation analyses

The GO term enrichment analyses were calculated using the integrated TopGO analysis in the A2TEA.WebApp, always using the Conserved Set as a background distribution. GO terms were only considered over represented, when the one sided fisher’s test p-value was below 0.05 and the GO term was annotated at least twice in the set of interest.

#### Revigo Maps of gene functions

Gene functions were summarized and visualized in maps using the Revigo webtool [65] with default parameters.

#### Phylogenetic trees of candidate gene families

The candidate gene families were evaluated by plotting the phylogeny including up to 5 closest HOGs and the underlying multiple sequence alignment (MSA) of the peptide sequences with the *A2TEA.Webapp* [23].

#### Diversifying selection

Coding sequences of the four species were downloaded from ensemblgenomes [60] as listed in Supplementary Table 1, Additional File 1. We applied a snakemake ([66], version 7.0.0) workflow to test the gene families predicted by OrthoFinder (HOGs) for signatures of diversifying selection using the exploratory mode in *aBSREL* [35], an adaptive branch-site model. The workflow is publicly available in a github repository [67]. It includes the following steps: 1) We supplied the coding sequences of the proteins from all four species and a matrix which contained the gene families with the corresponding protein names. 2) The data were pre-processed to obtain a codon aware multiple sequence alignment per HOG for the aBSREL test for diversifying selection. First, all proteins whose coding sequence contained “N” were removed from the workflow. Then, the amino acid sequences were translated from the coding sequences using the hyphy [68] *pre-msa.bf* script (https://github.com/veg/hyphy-analyses/tree/master/codon-msa) using the --E 0.05 option to allow alignment of sequences with low homology. 3) The amino acid sequences of each HOG were then aligned to each other with muscle ([69] version 5.1) and the nucleotide multiple sequence alignment (MSA) was obtained from the peptide alignment with the hyphy *post-msa.bf* script. 4) The phylogenetic trees were calculated with fasttree ([70], version 2.1) only for the HOGs which did not contain “?” in the nucleotide MSA. 5) The test for diversifying selection was then applied to all remaining HOGs. We used a false discovery rate of 0.05 to define signatures of diversifying selection in a gene. Hence, genes with a corrected p-value <= 0.05 have most likely experienced diversifying selection. The results of the test for diversifying selection were analyzed with the script *Analysis_A2TEA_output_with_Selection_v04_with_corrected_pvalues.R,* Additional File 3. The over representation analyses of the signatures of diversifying selection within the subsets of candidate gene families were calculated with the script *Selection_overrepresentation_v04_with_corrected_pvalues.Rmd,* Additional File 3.

## Abbreviations

AA: Amino acid
ABA: Abscisic acid
*Aly*: *Arabidopsis lyrata*
*Ath*: *Arabidopsis thaliana*
Bna: Brassica napus
BP: “Biological Process”
DE: Differential expression
DEG: Differentially expressed gene
*Esa*: *Eutrema salsugineum*
GO: Gene ontology
HOG: Hierarchical Orthologous Group
HRD: Human readable description
MSA: Multiple sequence alignment
VLCFA: Very long-chain fatty acid
WGD: Whole genome duplication

## Declarations

### Ethics approval and consent to participate

’Not applicable’

### Consent for publication

’Not applicable’

### Availability of data

- No datasets were generated in this study. The analyses were performed on publicly available genomic ([57], [58], [59], [28]) and transcriptomic ([25], [61]) data (details see Methods).
- The raw results from the A2TEA.Workflow (*A2TEA_finished.Rdata)*, a table of all conserved gene families and the respective genes of the four species (*Conserved_dataset_matrix.tsv)* and a table with the p-values of the test for diversifying selection for all tested genes (*OGs_pos_selection_pvalues.tsv)* are available in a bonndata repository [71] under a CC-BY-4.0 license.
- All analysis scripts are in **Additional File 3**: *Scripts.zip*.
- The code for the workflow for predicting signatures of diversifying selection is available in a github repository [67] under a Apache-2.0 license.

### Competing interests

The authors declare that they have no competing interests.

### Funding

Carolin Uebermuth-Feldhaus was funded by a PhD scholarship from Evangelisches Studienwerk e. V. Villigst

### Authors’ contributions

CUF and HS designed the study. CUF performed all analyses. CUF and HS interpreted the results of the analyses. CUF and HS wrote the manuscript.

## Acknowledgements

We thank Tyll Stöcker for advice and help with the *A2TEA.Workflow*.

## Supplementary Information

- **Additional File 1**: *Supplementary_Text_Figures_and_Tables.pdf,* Text file which contains Supplementary Figures 1-26 and Supplementary Tables 1-7, Supplementary Methods, Results and Discussion.
- **Additional File 2:** *Candidate_gene_families_and_tolerant_specific_expansions.xlsx*, Table which includes all relevant info about the gene families which are expanded in *Eutrema salsugineum* and/or *Arabidopsis lyrata* and DE in *Eutrema salsugineum* and/or *Arabidopsis lyrata*, including manual notes. Contains two sheets: Sheet 1 contains the actual information, the column headers are explained in Sheet 2.
- **Additional File 3**: *Scripts.zip*: *filter_min100aa.sh, normalize.sh* (bash scripts for pre-processing of raw genomic data), *Analysis_A2TEA_output.R*, *Analysis_expression_pattern_per_OG_only_at_least_2_sig.R*, *Analysis_A2TEA_output_with_Selection_v04_with_corrected_pvalues.R*, *Selection_overrepresentation_v04_with_corrected_pvalues.Rmd*, *Selection_overrepresentation_v04_with_corrected_pvalues.pdf* and *merge_all_info_of_candidate_gene_families.R* (Rscripts for the Analyses performed on the raw *A2TEA.Workflow* results and on the results including the prediction of diversifying selection)
- **Additional File 4**: [71] contains *A2TEA_finished.Rdata* (the raw results from the *A2TEA.Workflow*), *Conserved_dataset_matrix.tsv* (a table of all conserved gene families and the respective genes of the four species) and *OGs_pos_selection_pvalues.tsv* (a table with the p-values of the test for diversifying selection for all tested genes)
- **Additional File 5**: *GO_term_enrichment_analyses.xlsx*: Tables with the Results of GO term enrichment analyses. Contains nine sheets: Sheet 1: *Sig_enriched_GO_terms_DEG_Aly_and_Esa*, Sheet 2: *Sig_enriched_GO_terms_DEG_Aly_or_Esa*, Sheet 3: *Sig_enriched_GO_terms_DEG_Aly*, Sheet 4: *Sig_enriched_GO_terms_DEG_Esa*, Sheet 5: *Sig_enriched_GO_terms_exp_Aly_and_Esa*, Sheet 6: *Sig_enriched_GO_terms_exp_Aly*, Sheet 7: *Sig_enriched_GO_terms_exp_Esa*, Sheet 8: *Sig_enriched_GO_terms_exp_and_DEG_Aly*, Sheet 9: *Sig_enriched_GO_terms_exp_and_DEG_Esa*

## Notes

### Competing Interest Statement

The authors have declared no competing interest.

