## Additional File 1 for "Integration of differential expression under drought with gene family expansion unique to drought tolerant species predicts candidate genes for drought adaptation in Brassicaceae species"

### Supplementary Text, Supplementary Figures and Tables

#### Supplementary Figures

#### Supplementary Tables

#### Supplementary Methods

*Supplementary Table 1: Genomic Data used in this study.*

| Species | Genome assembly | pep.fasta | cDNA.fasta | CDS.fasta | Annotation.gff3 | Publication |
|---|---|---|---|---|---|---|
| *Eutrema salsugineum* | http://ftp.ensemblgenomes.org/pub/plants/release-53/fasta/eutrema_salsugineum/dna/Eutrema_salsugineum.Eutsalg1_0.dna.toplevel.fa.gz | http://ftp.ensemblgenomes.org/pub/plants/release-53/fasta/eutrema_salsugineum/pep/Eutrema_salsugineum.Eutsalg1_0.pep.all.fa.gz | http://ftp.ensemblgenomes.org/pub/plants/release-53/fasta/eutrema_salsugineum/cdna/Eutrema_salsugineum.Eutsalg1_0.cdna.all.fa.gz | http://ftp.ensemblgenomes.org/pub/plants/release-53/fasta/eutrema_salsugineum/cds/Eutrema_salsugineum.Eutsalg1_0.cds.all.fa.gz | http://ftp.ensemblgenomes.org/pub/plants/release-53/gff3/eutrema_salsugineum/Eutrema_salsugineum.Eutsalg1_0.53.gff3.gz | [1] |
| *Arabidopsis lyrata* | http://ftp.ensemblgenomes.org/pub/plants/release-53/fasta/arabidopsis_lyrata/dna/ | http://ftp.ensemblgenomes.org/pub/plants/release-53/fasta/arabidopsis_lyrata/pep/ | http://ftp.ensemblgenomes.org/pub/plants/release-53/fasta/arabidopsis_lyrata/cdna/ | http://ftp.ensemblgenomes.org/pub/plants/release-53/fasta/arabidopsis_lyrata/cds/ | http://ftp.ensemblgenomes.org/pub/plants/release-53/gff3/arabidopsis_lyrata/Arabidopsis_lyrat | [2] |

| | | | | | | |
|---|---|---|---|---|---|---|
| | Arabidopsis_lyrata.v.1.0.dna.toplevel.fa.gz | Arabidopsis_lyrata.v.1.0.pep.all.fa.gz | Arabidopsis_lyrata.v.1.0.cdna.all.fa.gz | Arabidopsis_lyrata.v.1.0.cds.all.fa.gz | a.v.1.0.53.gff3.gz | |
| *Arabidopsis thaliana* | http://ftp.ensemblgenomes.org/pub/plants/release-53/fasta/arabidopsis_thaliana/dna/Arabidopsis_thaliana.TAIR10.dna.toplevel.fa.gz | http://ftp.ensemblgenomes.org/pub/plants/release-53/fasta/arabidopsis_thaliana/pep/Arabidopsis_thaliana.TAIR10.pep.all.fa.gz | http://ftp.ensemblgenomes.org/pub/plants/release-53/fasta/arabidopsis_thaliana/cdna/Arabidopsis_thaliana.TAIR10.cdna.all.fa.gz | http://ftp.ensemblgenomes.org/pub/plants/release-53/fasta/arabidopsis_thaliana/cds/Arabidopsis_thaliana.TAIR10.cds.all.fa.gz | http://ftp.ensemblgenomes.org/pub/plants/release-53/gff3/arabidopsis_thaliana/Arabidopsis_thaliana.TAIR10.53.gff3.gz | [3] |
| *Brassica napus* | http://ftp.ensemblgenomes.org/pub/plants/release-53/fasta/brassica_napus/dna/Brassica_napus.AST_PRJEB5043_v1.dna.toplevel.fa.gz | http://ftp.ensemblgenomes.org/pub/plants/release-53/fasta/brassica_napus/pep/Brassica_napus.AST_PRJEB5043_v1.pep.all.fa.gz | http://ftp.ensemblgenomes.org/pub/plants/release-53/fasta/brassica_napus/cdna/Brassica_napus.AST_PRJEB5043_v1.cdna.all.fa.gz | http://ftp.ensemblgenomes.org/pub/plants/release-53/fasta/brassica_napus/cds/Brassica_napus.AST_PRJEB5043_v1.cds.all.fa.gz | http://ftp.ensemblgenomes.org/pub/plants/release-53/gff3/brassica_napus/Brassica_napus.AST_PRJEB5043_v1.53.gff3.gz | [4] |

*Supplementary Table 2: RNA-Seq reads used in this study.*

| Species (genotype) | Publication | Project ID | Libraries Drought | Libraries Control | Organ | Treatment |
|---|---|---|---|---|---|---|
| *Eutrema salsugineum* (accession Shandong) | [5] | SRP155798 | SRR7624684, SRR7624685, SRR7624692 | SRR7624687, SRR7624721, SRR7624722 | rosette leaf | Watering was stopped when leaf 6 was initiated on the apex and RNA samples were taken on day 11 of watering stop (T11). |
| *Arabidopsis lyrata* (strain MN47, same as Genome publication) | [5] | SRP155798 | SRR7624680, SRR7624702, SRR7624703 | SRR7624732, SRR7624733, SRR7624742 | rosette leaf | Watering was stopped when leaf 6 was initiated on the apex and RNA samples were taken on day 11 of watering stop (T11). |
| *Arabidopsis thaliana* (Col-0) | [5] | SRP155798 | SRR7624694, SRR7624696, SRR7624697 | SRR7624710, SRR7624714, SRR7624723 | rosette leaf | Watering was stopped when leaf 6 was initiated on the apex and RNA samples were taken on day 11 of watering stop (T11). |
| *Brassica napus* (Cultivar 29005) | [6] | SRP277041, GSE156029 | SRR12429701, SRR12429702, SRR12429703 | SRR12429698, SRR12429699, | leaf | Before bolting stage, watering was stopped and RNA |

| | |
|---|---|
| SRR12429700 | samples were taken on day 3 of watering stop. |

5

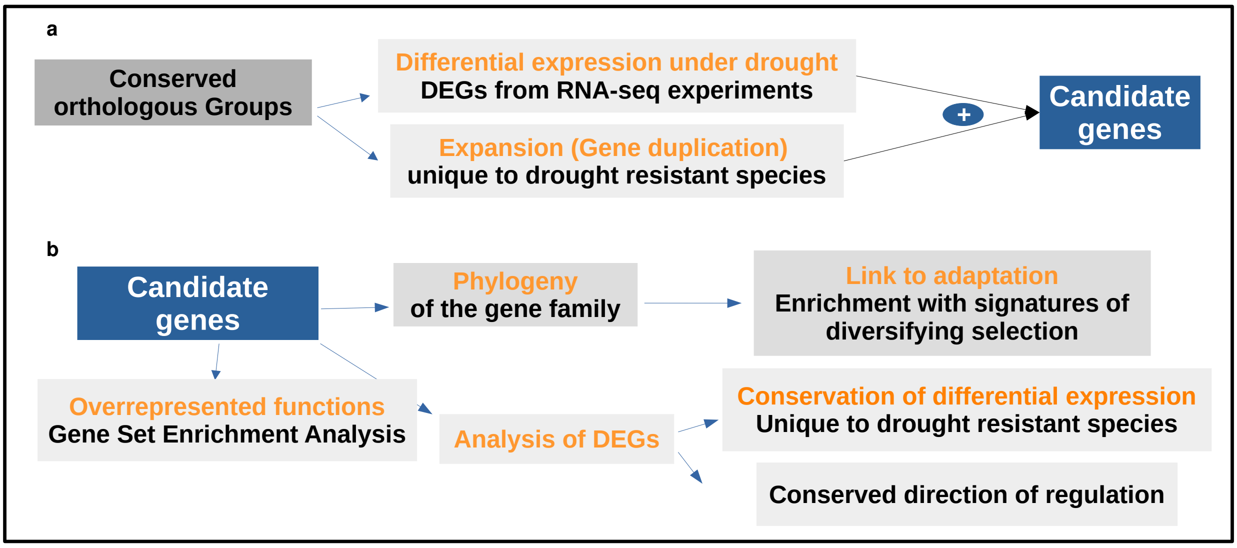

*Supplementary Figure 1: Workflow of the prediction of candidate gene families for drought adaptation and their characterization and evaluation.*

*A) Integration of differential gene expression with gene family expansion predicts candidate genes for drought adaptation. B) The candidate genes are characterized: their functions, the conservation of differential expression and the direction of regulation are analyzed. Moreover, the evolution of the candidate gene families is described by their phylogeny. Signatures of diversifying selection as predicted by absrel ([7]) are used as evaluation criterion for a gene family's relevance for adaptation.*

6

### Supplementary Results and Discussion

#### Gene families

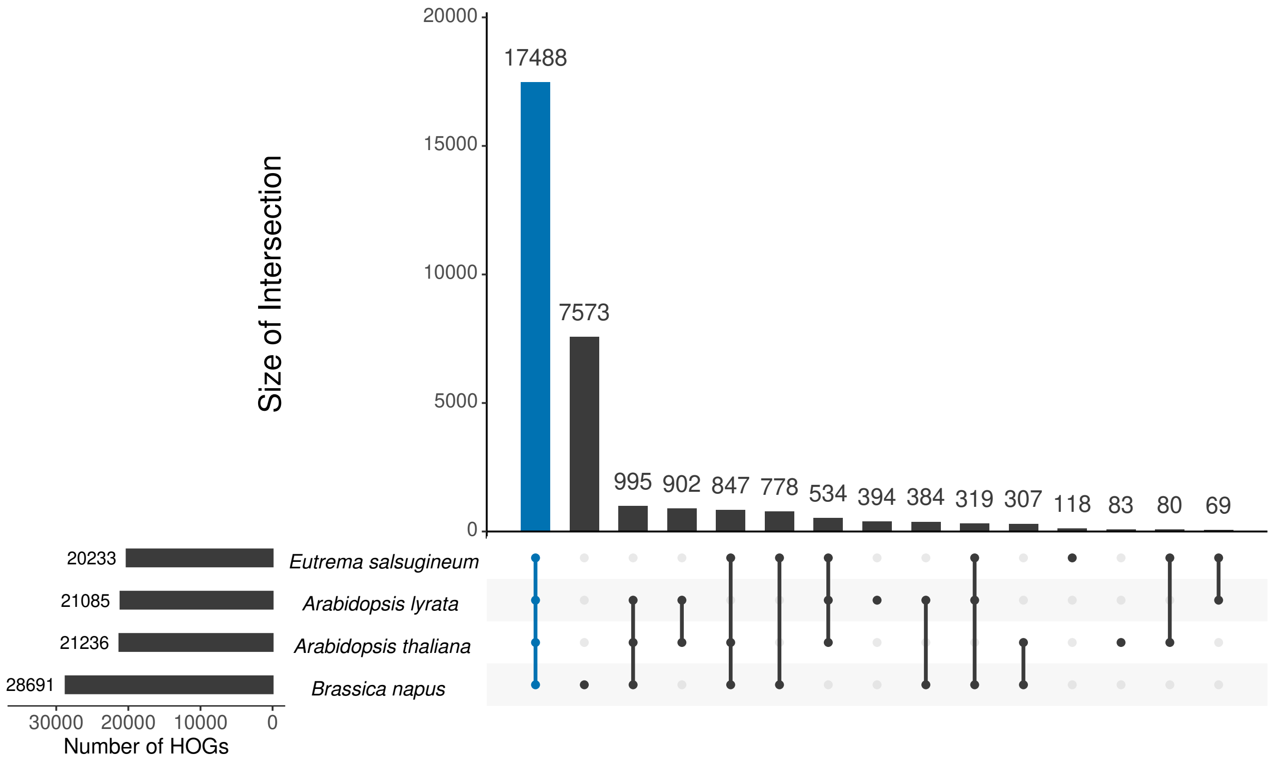

*Supplementary Figure 2: UpSet plot showing all gene families (HOGs, Hierarchical Orthologous Groups) predicted by orthofinder.*

*The intersections of gene families are shown, where blue marks the Conserved Set, i.e. intersections with all four species in the study. Black bars on the left show the total number of gene families per species.*

#### 10 Gene regulation under drought: Differential expression between drought and control growth conditions

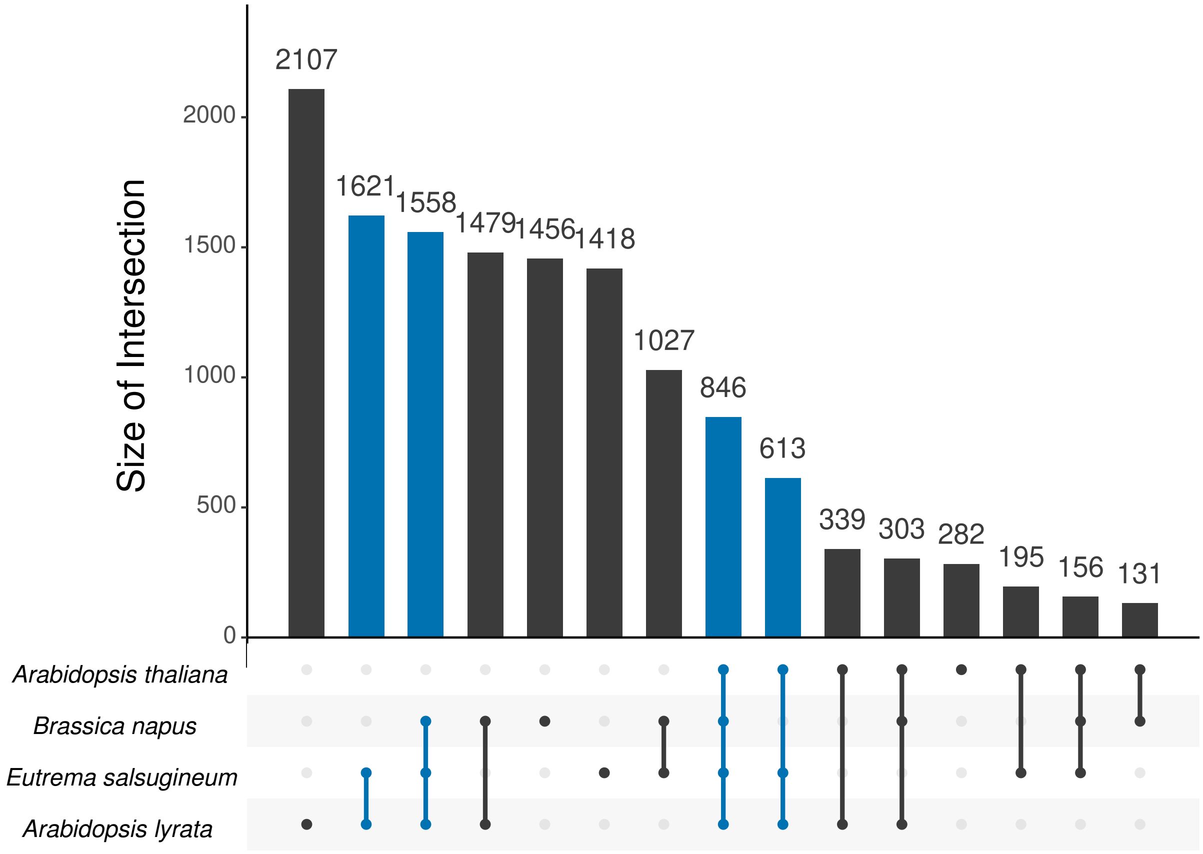

*Supplementary Figure 3: UpSet plot showing all gene families in the Conserved Set.*

*The intersections of gene families with DEG from the given species are shown, where blue marks intersections with both tolerant species. The species are ordered by the number of gene families, with the largest set on the bottom.*

**Differential expression under drought of expanded gene families in tolerant and sensitive species**

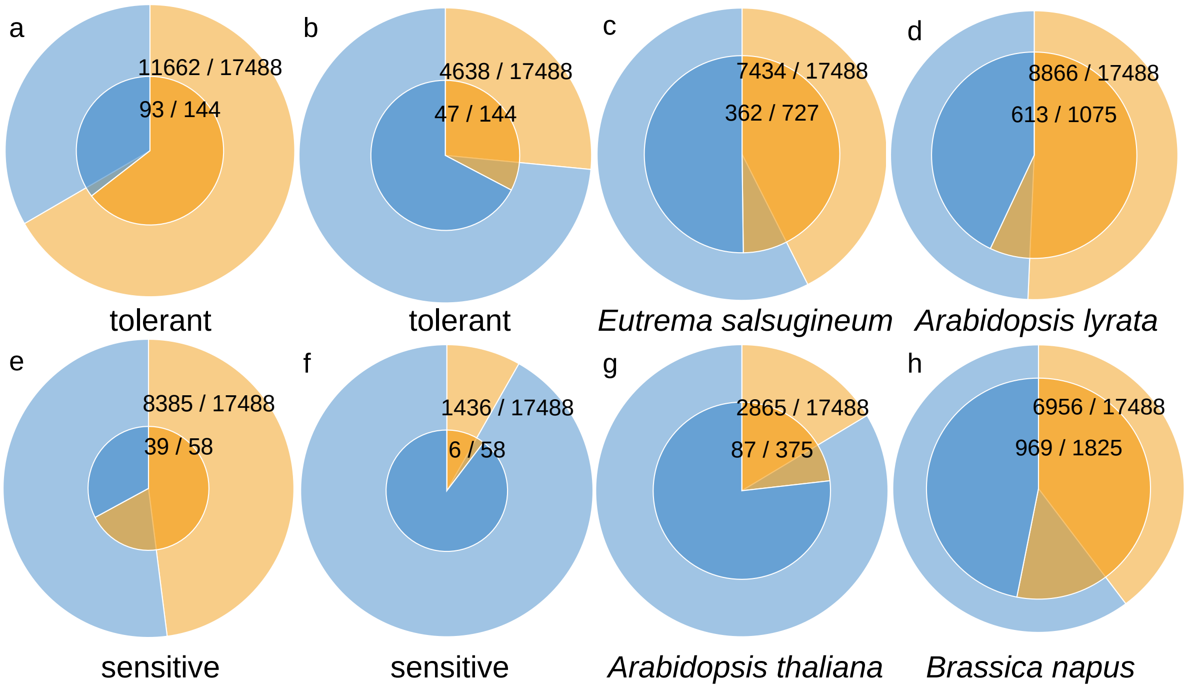

*Supplementary Figure 4: Pie charts of the number of gene families with DEG (orange) or without DEG (blue) within the Conserved Set (outer circle) or within the subset of expanded HOGs (inner circle, log10-scale) as defined by the names below the circles.*

*The DEGs are from a) any of the tolerant species, b) both tolerant species, c) Esa, d) Aly, e) any of the sensitive species, f) both sensitive species, g) Ath or h) Bna.*

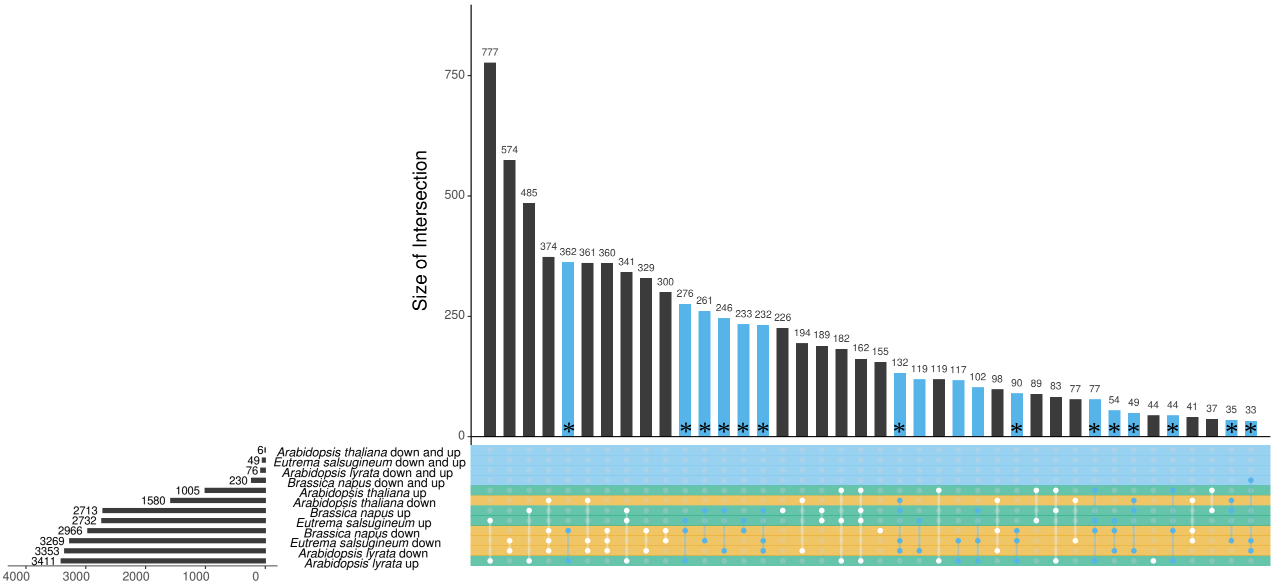

*Supplementary Figure 5: UpSet plot of the intersections of regulation categories over all DEGs from each of the species in the Conserved Set.*

*Colors of the rows label the category of regulation of each set of gene families per each species. Black bars on the left show the total number of gene families in each regulation category. Blue colored intersects are those gene families which show up and down regulation and within these, asterisks mark intersections which include Brassica napus. Only gene families with a minimum of two DEGs are considered and only the 40 largest intersections are displayed.*

The enrichment with differential expression unique to both tolerant species can also slightly be observed in gene families which are only up regulated: In the candidate gene families, from all gene families which show only up regulation in at least both tolerant species (24 HOGs, green bars in Supplementary Figure 6), 17 (71 %) are not regulated in any sensitive species. In contrast: In the Conserved Set, from 1472 HOGs which show only up regulation in at least both tolerant species, 777 (53 %) are not regulated in any sensitive species (Supplementary Figure 5). However, this enrichment has a low level of significance (p = 0.06).

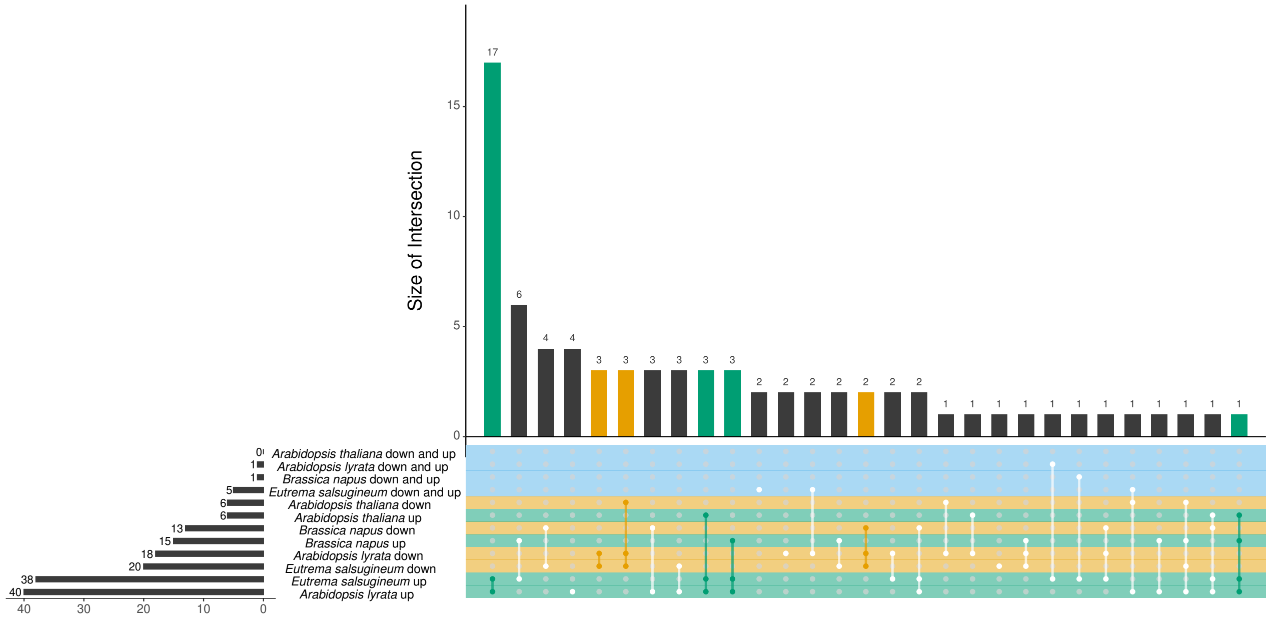

*Supplementary Figure 6: UpSet plot of the intersections of regulation categories over all DEGs from each of the species in the gene families expanded in both tolerant species.*

*Black bars on the left show the total number of gene families in each regulation category. Colors of the rows label the category of regulation of each set of gene families per each species. Colored intersects are those gene families which are either only up (green) or only down (orange) regulated and in which both tolerant species have the same regulation category. Only gene families with a minimum of two DEGs are considered and only the 40 largest intersections are displayed.*

### Candidate gene families

#### Enriched functions of candidate gene families

*Supplementary Table 3: Significantly enriched GO terms of candidate gene families.*

| GO.ID | Term | Annotated | Significant | Expected | p-value |
|---|---|---|---|---|---|
| GO:0000398 | mRNA splicing, via spliceosome | 167 | 6 | 0.92 | 0.00544 |
| GO:0019941 | modification-dependent protein catabolic process | 419 | 5 | 2.31 | 0.04743 |

*The enrichment is calculated compared to the Conserved Set.*

*Supplementary Table 4: Significantly enriched GO terms of candidate gene families which are conserved DE in the tolerant species.*

| GO.ID | Term | Annotated | Significant | Expected | p-value |
|---|---|---|---|---|---|
| GO:0019941 | modification-dependent protein catabolic process | 419 | 3 | 1.15 | 0.02397 |
| GO:0006412 | translation | 634 | 7 | 1.74 | 0.0395 |

*The enrichment is calculated compared to the Conserved Set.*

#### Enriched functions of differentially expressed genes in any or in both tolerant species

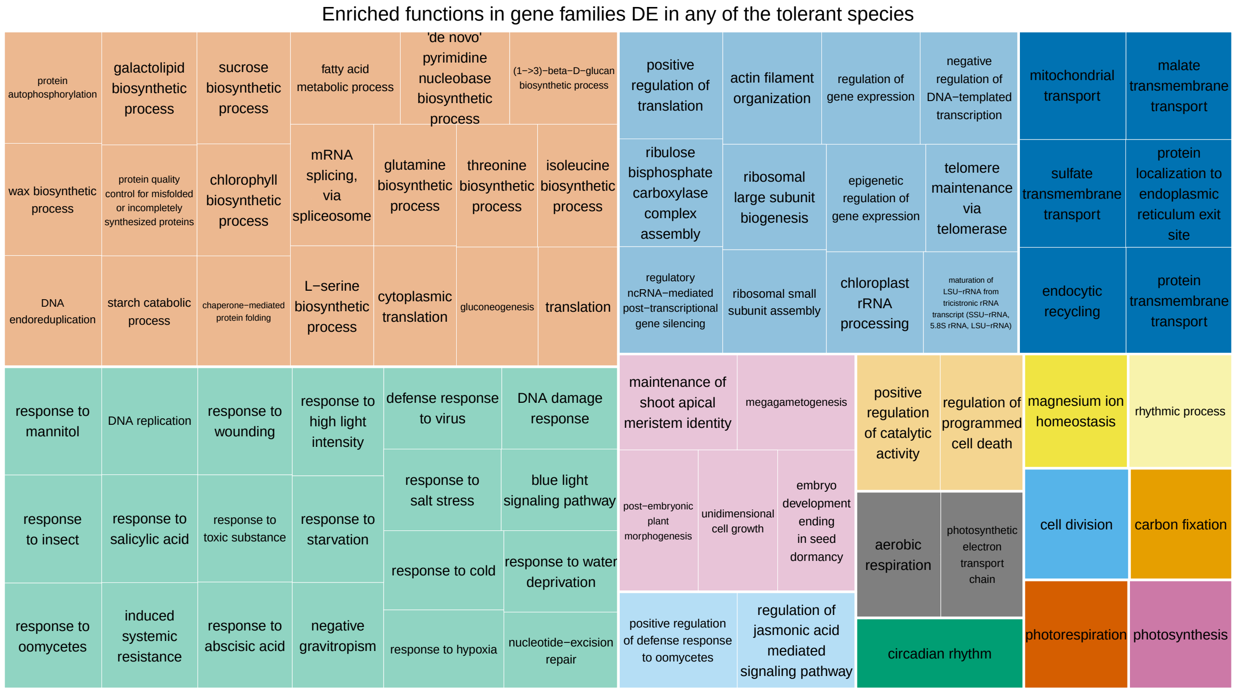

*Supplementary Figure 7: Revigo TreeMap of significantly (p <= 0.05) over represented functions of gene families which show DE in any tolerant species.*

*Loosely related terms are clustered together. Square size is the log10(p-value) of the one-sided fisher test for over representation. Refer to GO_term_enrichment_analyses.xlsx, Additional File 5 for tables with values.*

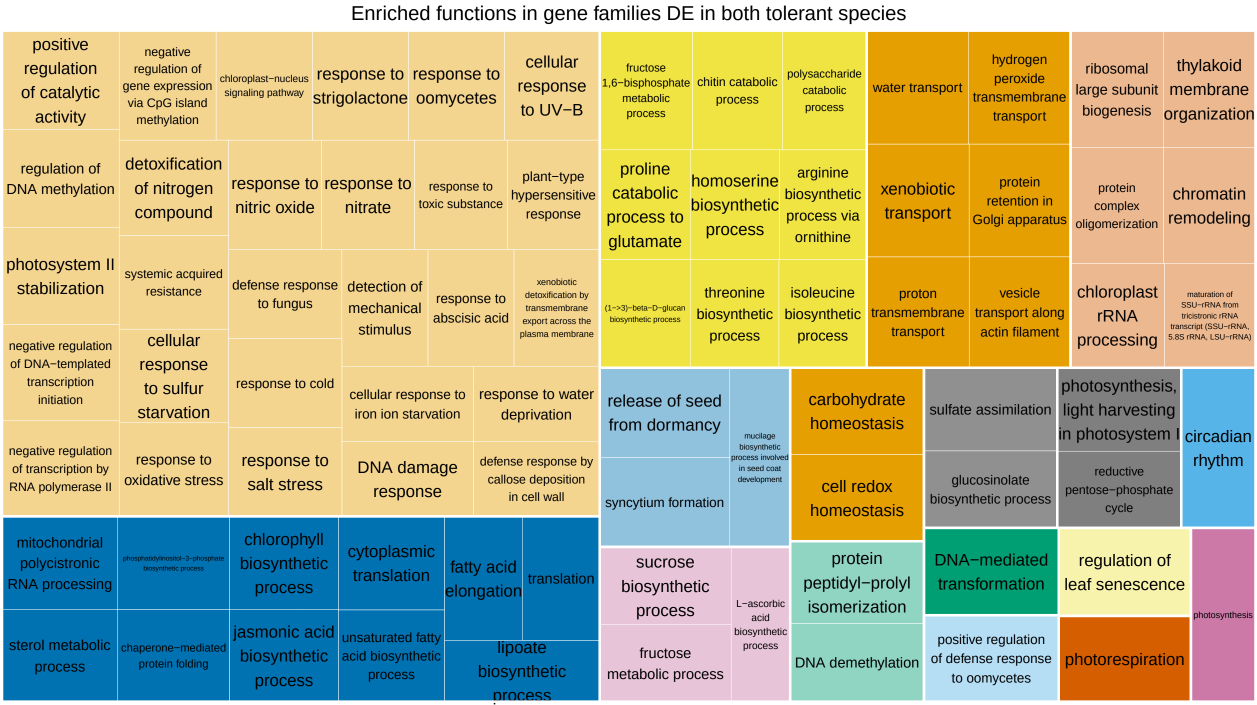

*Supplementary Figure 8: Revigo TreeMap of significantly (p <= 0.05) over represented functions of gene families which show DE in both tolerant species.*

*Loosely related terms are clustered together. Square size is the log10(p-value) of the one-sided fisher test for over representation. Refer to GO_term_enrichment_analyses.xlsx, Additional File 5 for tables with values.*

#### Candidate Gene families with enriched functions

There are several candidate gene families which function in “modification-dependent protein catabolic process”:

1) N0.HOG0008832 is a E3 UFM1-protein ligase 1 homolog (AT3G46220) that mediates ufmylation (TAIR Curated Description). Both duplicates from *Esa* are up regulated under drought (Supplementary Figure 9) and one of these, EUTSA_v10002414mg, shows signatures of diversifying selection. One of two homologs is also up regulated in *Aly and* the other homolog (fgenesh1_pm_C_scaffold_1003137), which is not regulated under drought, is under diversifying selection. The single homologs from each of the sensitive species are neither regulated nor under positive selection. In *Ath* it is repressed under heat by the nat-siRNA in AT3G46230, which is up regulated under heat ([8]). 2) N0.HOG0007461 encodes a TOM1-LIKE (TOL) protein (AT5G16880), which regulates growth in *Ath* ([9]). Both *Aly* duplicates are up regulated (Supplementary Figure 10). 3)

N0.HOG0006674 *(CER9,* AT4G34100) encodes a E3 ubiquitin ligase which plays a role in cuticle biosynthesis and is known to be involved in plant drought tolerance in *Ath* ([10]). CER9 is duplicated in both tolerant species and up regulated in *Aly* (Supplementary Figure 11). Unfortunately, the function wax biosynthetic process (GO:0010025) was not annotated in our analysis.

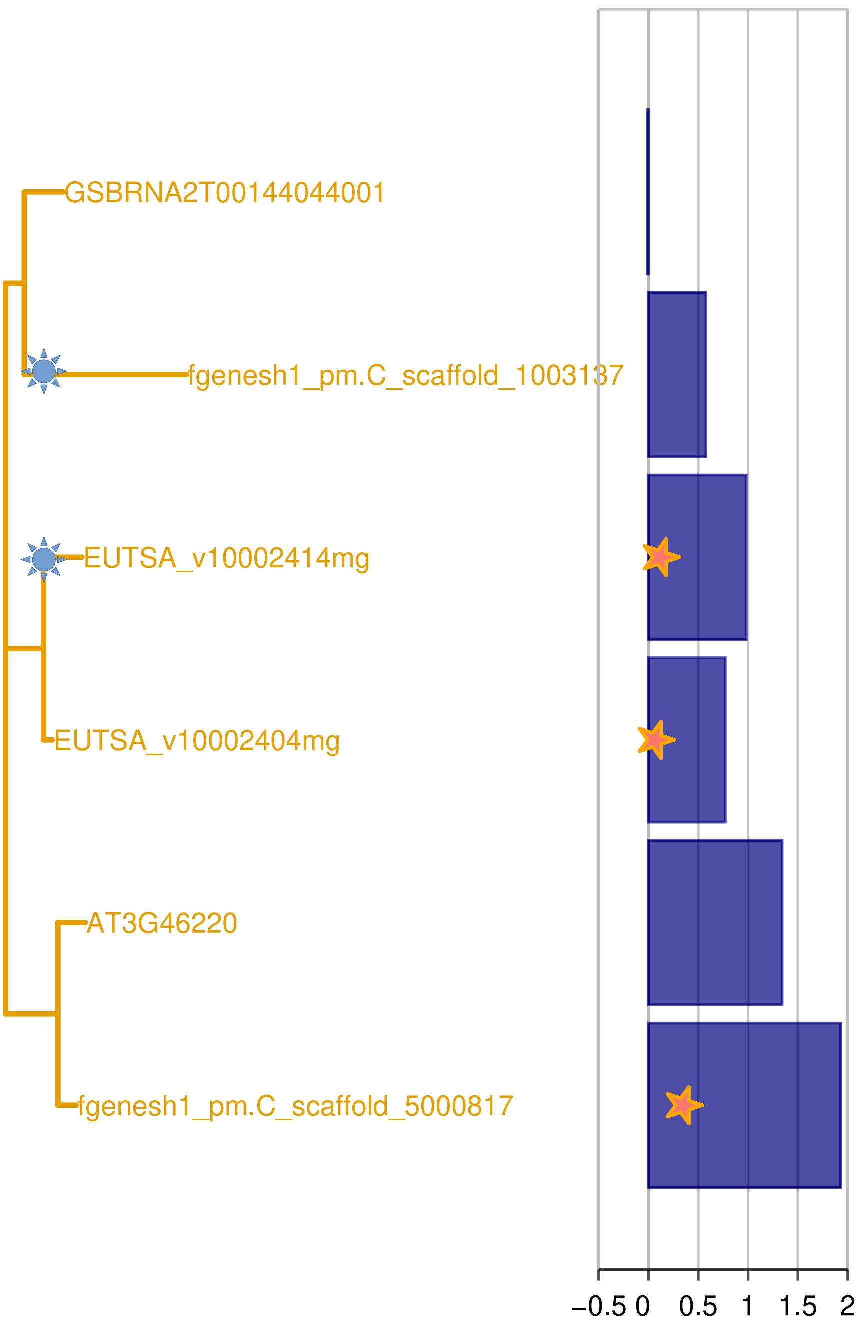

*Supplementary Figure 9: Phylogenetic tree of the genes from the four Brassicaceae species of the family N0.HOG0008832.*

*Blue bars represent differential expression between drought and control (log2FC), where stars indicate a corrected p-value <= 0.1). Genes with IDs starting with "EUTSA" are from Esa, with "fgenesh" or "scaffold" are from Aly, with "AT" are from Ath and with "GSBRN" are from Bna. Sun symbols indicate genes with signatures of diversifying selection.*

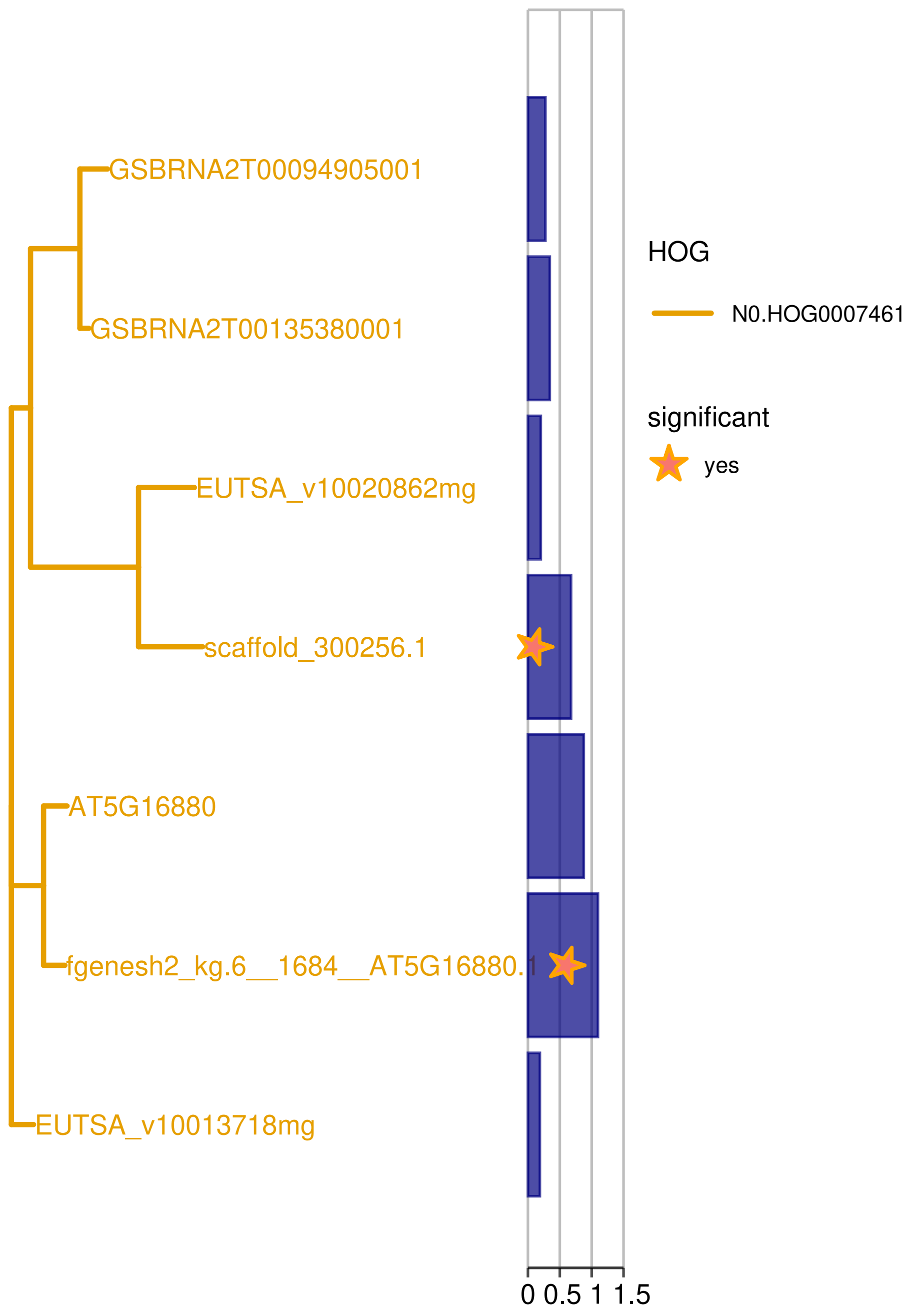

*Supplementary Figure 10: Phylogenetic tree of the genes from the four Brassicaceae species of the family N0.HOG0007461.*

*Blue bars represent differential expression between drought and control (log2FC), where stars indicate a corrected p-value <= 0.1). Genes with IDs starting with "EUTSA" are from Esa, with "fgenesh" or "scaffold" are from Aly, with "AT" are from Ath and with "GSBRN" are from Bna.*

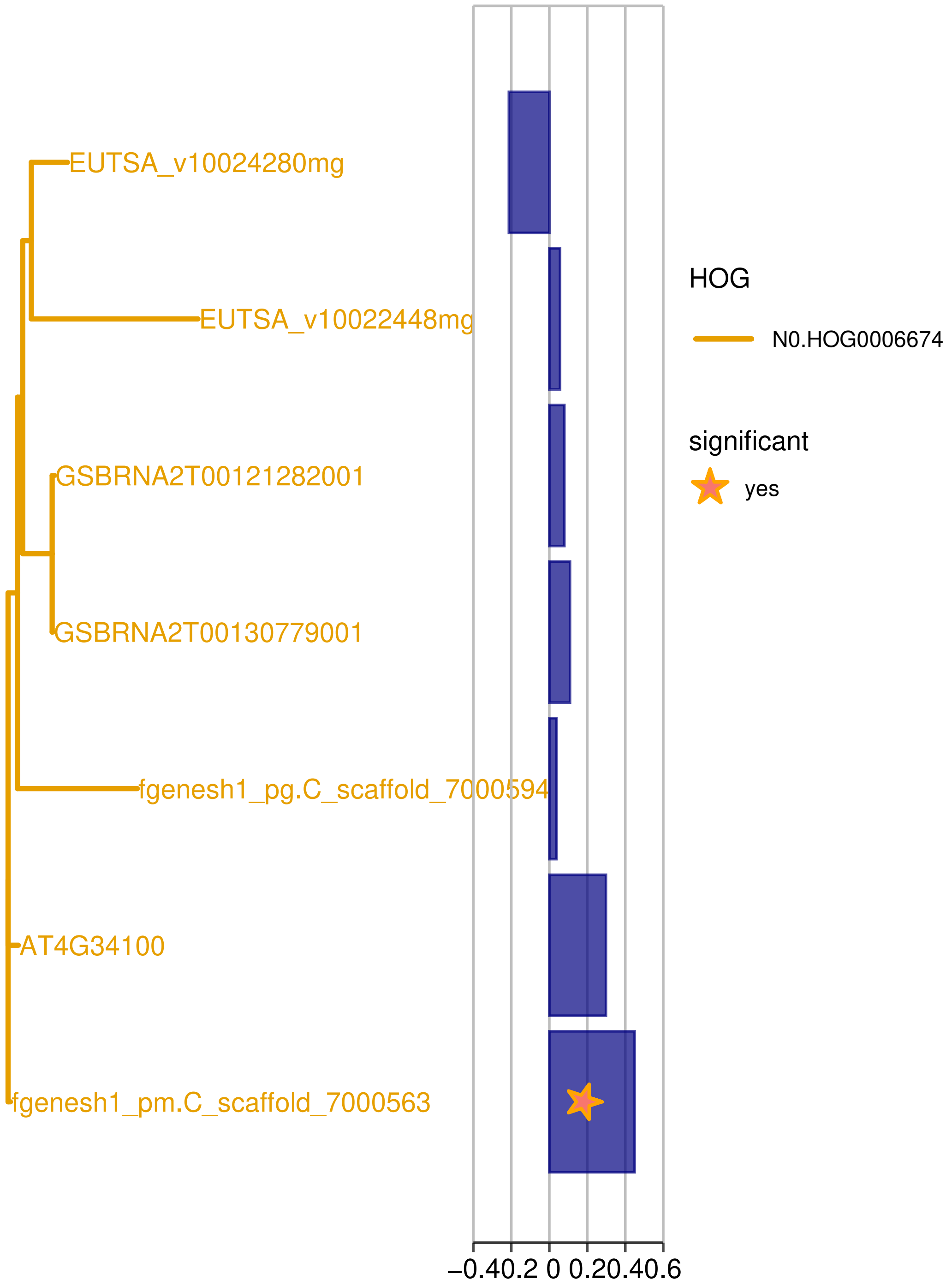

*Supplementary Figure 11: Phylogenetic tree of the genes from the four Brassicaceae species of the family N0.HOG0006674.*

*Blue bars represent differential expression between drought and control (log2FC), where stars indicate a corrected p-value <= 0.1). Genes with IDs starting with "EUTSA" are from Esa, with "fgenesh" or "scaffold" are from Aly, with "AT" are from Ath and with "GSBRN" are from Bna.*

#### 55 Functions of candidate gene families which are uniquely up regulated in the tolerant species

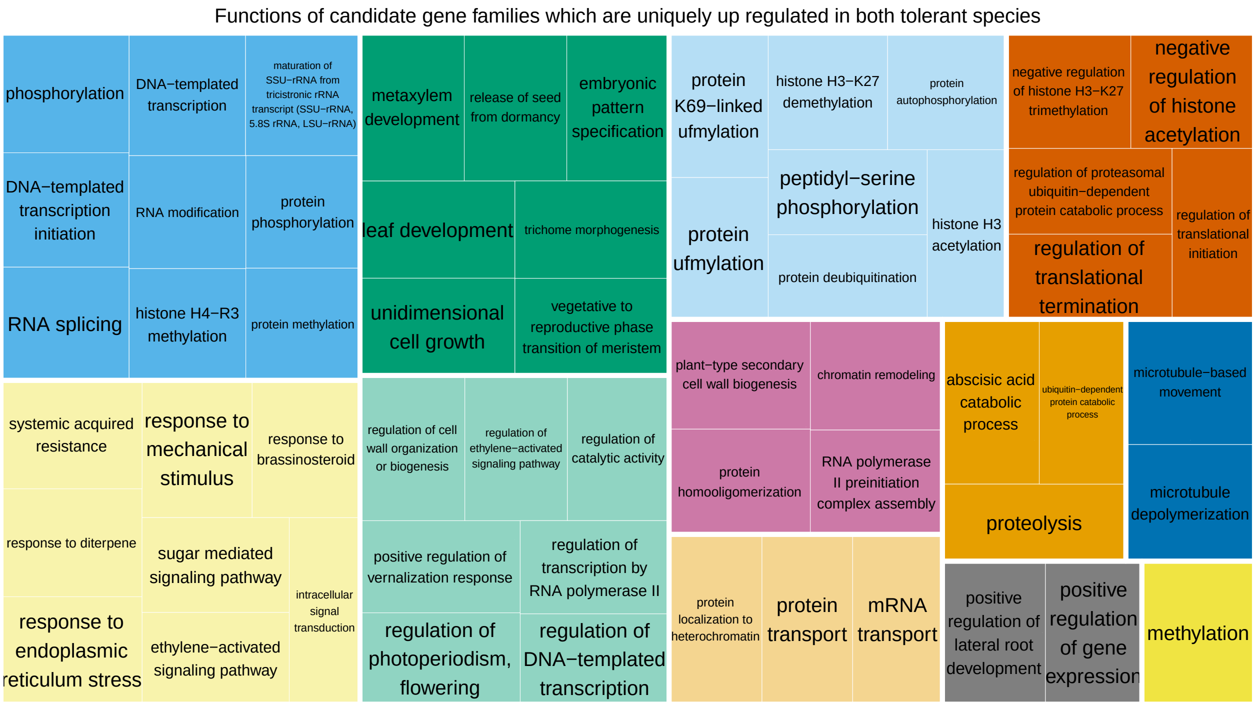

*Supplementary Figure 12: Revigo TreeMap of functions of candidate gene families which are conserved expanded and uniquely up regulated in both tolerant species.*

*Loosely related terms are clustered together. Square size is the uniqueness of the term.*

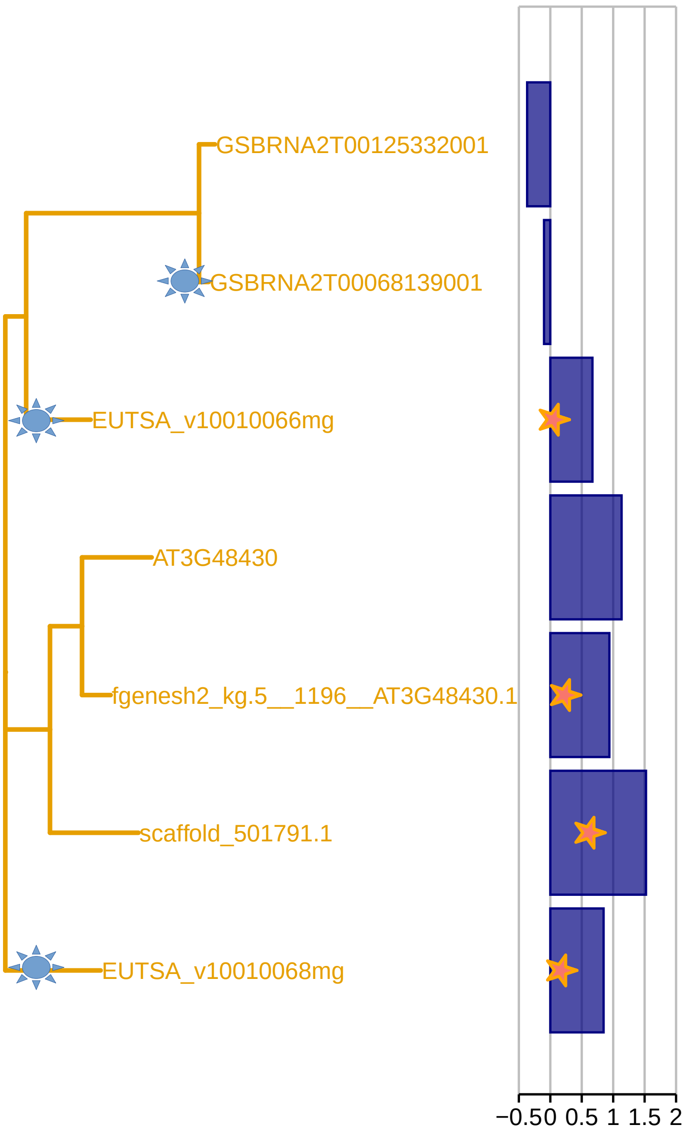

*Supplementary Figure 13: Phylogenetic tree of the genes from the four Brassicaceae species of the family N0.HOG0007350 (Lysine-specific demethylase REF6).*

*Blue bars represent differential expression between drought and control (log2FC), where stars indicate a corrected p-value <= 0.1). Genes with IDs starting with "EUTSA" are from Esa, with "fgenesh" or "scaffold" are from Aly, with "AT" are from Ath and with "GSBRN" are from Bna. Sun symbols indicate genes with signatures of diversifying selection.*

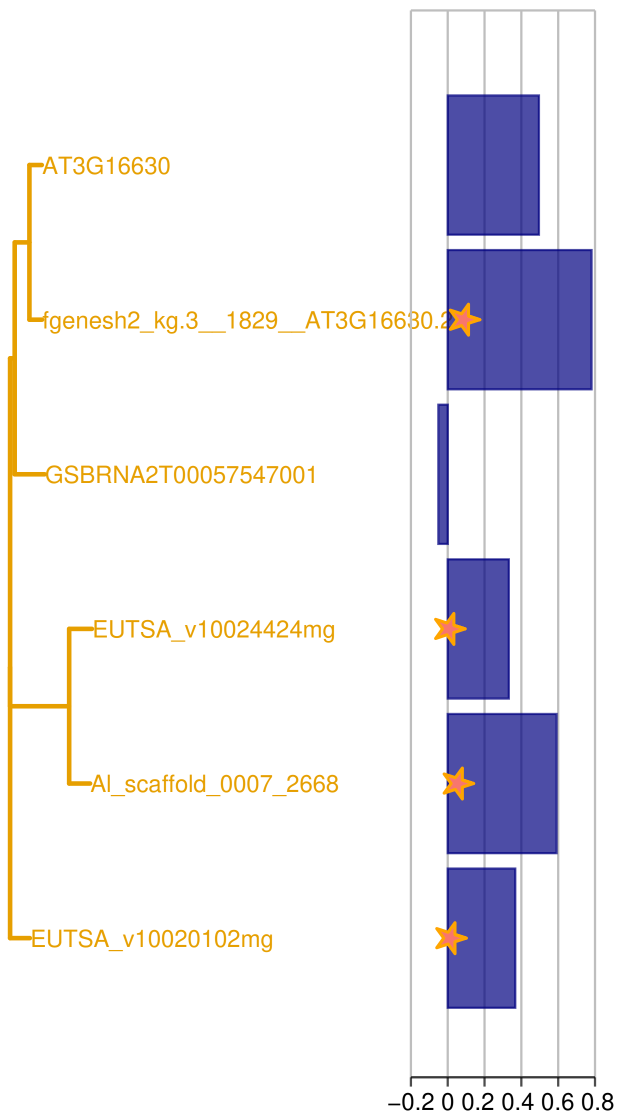

*Supplementary Figure 14: Phylogenetic tree of the genes from the four Brassicaceae species of the family N0.HOG0008546 (Kinesin-like protein KIN-13A).*

*Blue bars represent differential expression between drought and control (log2FC), where stars indicate a corrected p-value <= 0.1). Genes with IDs starting with "EUTSA" are from Esa, with "fgenesh" or "scaffold" are from Aly, with "AT" are from Ath and with "GSBRN" are from Bna.*

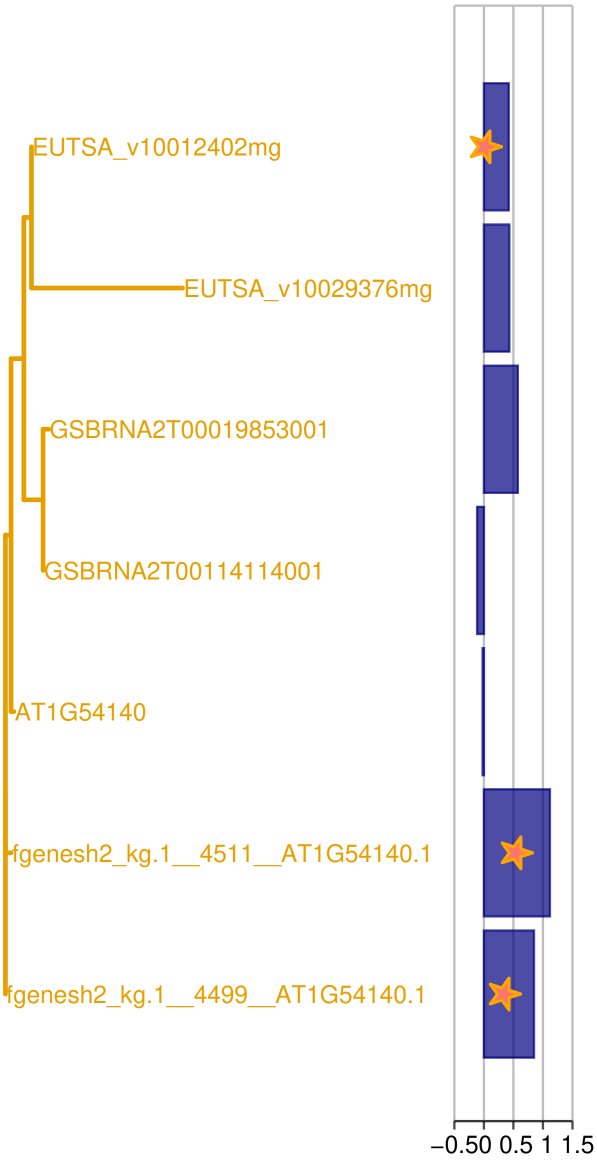

*Supplementary Figure 15: Phylogenetic tree of the genes from the four Brassicaceae species of the family N0.HOG0007005 (TFIID subunit 9).*

*Blue bars represent differential expression between drought and control (log2FC), where stars indicate a corrected p-value <= 0.1). Genes with IDs starting with "EUTSA" are from Esa, with "fgenesh" or "scaffold" are from Aly, with "AT" are from Ath and with "GSBRN" are from Bna.*

Among the candidate gene families which function in “mRNA splicing, via spliceosome” are 1) N0.HOG0006476, AT5G18810, the Serine/arginine-rich SC35-like splicing factor SCL28. The sequences of both homologs from *Aly* are quite diverged but expressed and one of the duplicates from *Esa* is up regulated together with both homeologs from *Bna*. 2) N0.HOG0007326, AT2G27100, Serrate RNA effector molecule, which is annotated as “regulator of meristem activity and adaxial leaf fate via the miRNA gene-silencing pathway” in Uniprot ([11]), which lists several publications. One of the two homologs of *Esa* is down regulated under drought. 3) N0.HOG0008634, AT1G77180, a SNW/SKI-interacting protein, is annotated as “splicing factor involved in post-transcriptional regulation of circadian clock and flowering time genes” in Uniprot ([11]). Both homologs from *Aly* are up regulated, whereas the sequence of one of two homologs from *Esa* is quite diverged and the gene is not expressed in this experiment. The duplication might have a common origin in both tolerant species but the neofunctionalization happened only in *Aly*, while the duplicate is probably pseudogenizing in *Esa*. As we know that the drought reactions differ between the two species , i.e. *Aly* responds earlier than *Esa* and *Ath* with growth reduction ([5]), this is an interesting candidate gene family for drought adaptation in *Aly*.

#### Gene families which are expanded and differentially expressed in *Esa*

There are 362 gene families expanded and DE in *Esa* (Supplementary Figure 4 c). Information on these gene families, including regulation under drought of the genes from all four species, their predicted functions and notes taken upon manual inspection, are provided in *Candidate_gene_families_and_tolerant_specific_expansions.xlsx*, Additional File 2. These gene families are over represented with some biological processes (Supplementary Figure 16, table with values in *GO_term_enrichment_analyses.xlsx*, Additional File 5), of which several are relevant for drought adaptation. These include the “regulation of ethylene-activated signaling pathway” ($p = 2.7e-03$, 3/22 HOGs, expected = 0.48), which is also significantly enriched in the whole subset of gene families expanded in *Esa* (*GO_term_enrichment_analyses.xlsx*, Additional File 5), “wax biosynthetic process” ($p = 0.03$, 3/32 HOGs, expected = 0.69), which is also significantly enriched in the whole subset of gene families expanded in *Esa*, and “stomatal complex development” ($p = 0.03$, 4/58 HOGs, expected = 1.25), which is not enriched in all gene families expanded in *Esa* not in all DEGs from *Esa* *GO_term_enrichment_analyses.xlsx*, Additional File 5).

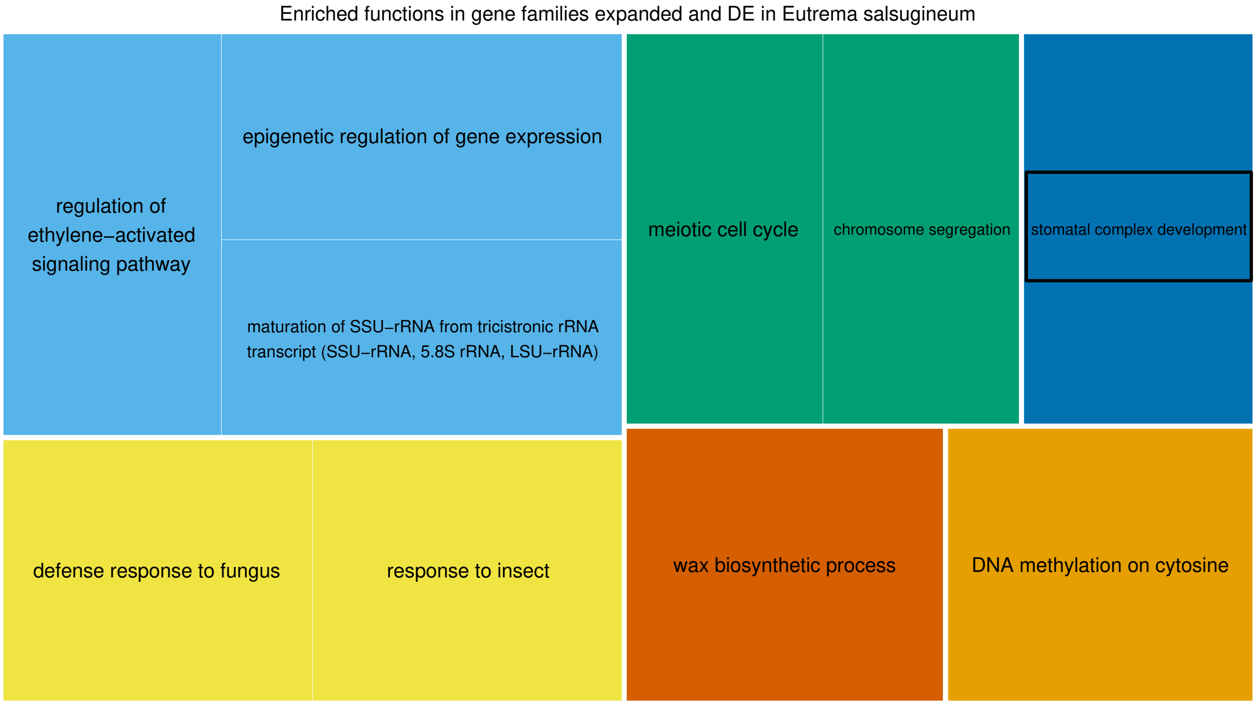

*Supplementary Figure 16: Revigo TreeMap of significantly (p <= 0.05) over represented biological processes of gene families expanded and differentially expressed in Esa.*

*Loosely related terms are clustered together. Square size is the log10(p-value) of the one-sided fisher test for over representation. Black ovals mark terms which are not enriched in all gene families either differentially expressed or expanded in Esa, but only in the combination of both. Refer to supplementary material for tables with values.*

##### *Example gene families with enriched functions*

**Wax biosynthetic process**

N0.HOG0001915 encodes a family of wax ester synthase/diacylglycerol acyltransferases 8 and 7 (AT5G16350 and AT5G12420). The gene family is expanded in all four analyzed species but has additional duplicates in *Esa*. Two homologs from *Bna* are highly down regulated under drought while one homolog from *Esa* is highly up regulated. I speculate that this contrasting regulation is related to the contrasting drought resistance of the species. Wax ester synthase / diacylglycerol acyltransferase 1 is the main enzyme in the late steps of cuticular wax synthesis in *Ath* ([12]).

N0.HOG0008907, AT2G47240, codes for Long chain acyl-CoA synthetase 1 (LACS1, CER8), which is involved in the early steps of cutin and wax biosynthesis. Together with LACS2, it is essential for normal cuticle synthesis ([13]). The gene family is expanded in *Esa* and *Aly* and one homolog from *Esa* is down regulated under drought, while the other homolog is not expressed in this experiment (Supplementary Figure 17). One *Bna* homeolog is down regulated under drought.

**Stomatal complex development**

N0.HOG0006742 encodes Cyclin-A2-4 (AT1G80370) , which is, redundantly with Cyclin-A2-1, required for the vein development ([14]). There are two gene duplications in *Esa.* Upon closer inspection including the next closest HOGs, there are also two duplications in *Bna*. One homolog from *Esa* is up regulated under drought. A study of the large cyclin family in *Brassica rapa* identified hormone-correlated responsive elements in the promoter regions of CYCA2;3, CYCA2;4 ([15]).

N0.HOG0005324, AT3G24140 encodes Transcription factor FAMA which is responsible for guard cell differentiation ([16]). It is expanded in both tolerant species and, upon closer inspection including the next closest HOGs, there is also a duplication in *Bna*. One duplicate is up regulated under drought in *Esa.* Its corresponding homolog in *Bna* is also up regulated but the other homeolog is down regulated. The duplicate from *Esa* is not expressed in this experiment. FAMA is also duplicated in the CAM-plant *Kalanchoe laxiflora* ([17]).

N0.HOG0009807 encodes the Lysine-specific demethylase JMJ25 (AT4G00990, Supplementary Figure 18). Both homologs from *Esa* and the single homolog from *Aly* are up regulated under drought. One of the homologs from *Esa* has an insertion in the middle of the peptide sequence. Upon closer inspection including the next closest HOGs, there is also a duplication in *Bna* which is not regulated under drought. Interestingly, the corresponding homolog of *Talinum triangulare*, KMT10399, was identified as a possible regulator of CAM photosynthesis ([18]), an important drought tolerance mechanism. Moreover, it positively regulates drought-stress responses in *Ath* ([19]). A loss-of-function mutant in a related histone demethylase JMJ27 increased dehydration stress tolerance in *Ath* by regulating OST1 (OPEN STOMATA 1) and other drought stress regulators ([19]).

N0.HOG0010477 encodes BASL (BREAKING OF ASYMMETRY IN THE STOMATAL LINEAGE, AT5G60880), which controls polarity in the development of stomata ([20], [21]). It is duplicated in *Kalanchoe laxiflora* which is adapted to drought by using CAM photosynthesis, while it is lost in the basic Eudicot *Nymphaea colorata* ([17]). The gene family is duplicated in *Esa.* One of the homologs is down regulated under drought, while the second homolog is not expressed in this experiment. Upon closer inspection including the next closest HOGs, there is also a duplication in *Bna*. Both homologs from *Bna* have one homeolog, each, down regulated. The observation that the second homolog from *Esa* is not expressed in this experiment leads us to speculate that it has acquired a function under different conditions or in a different tissue.

In gene families which function in stomatal complex development, duplications in *Esa* seem to be conserved with *Bna,* which suggests that they are relevant for common traits between the two species. On the other hand, their regulation under drought differs between the two species, suggesting that in *Bna,* the duplicates might be relevant for other than drought adaptation traits. We do not have an explanation for why OrthoFinder did not predict the duplications in *Bna* as orthologs to the respective gene family.

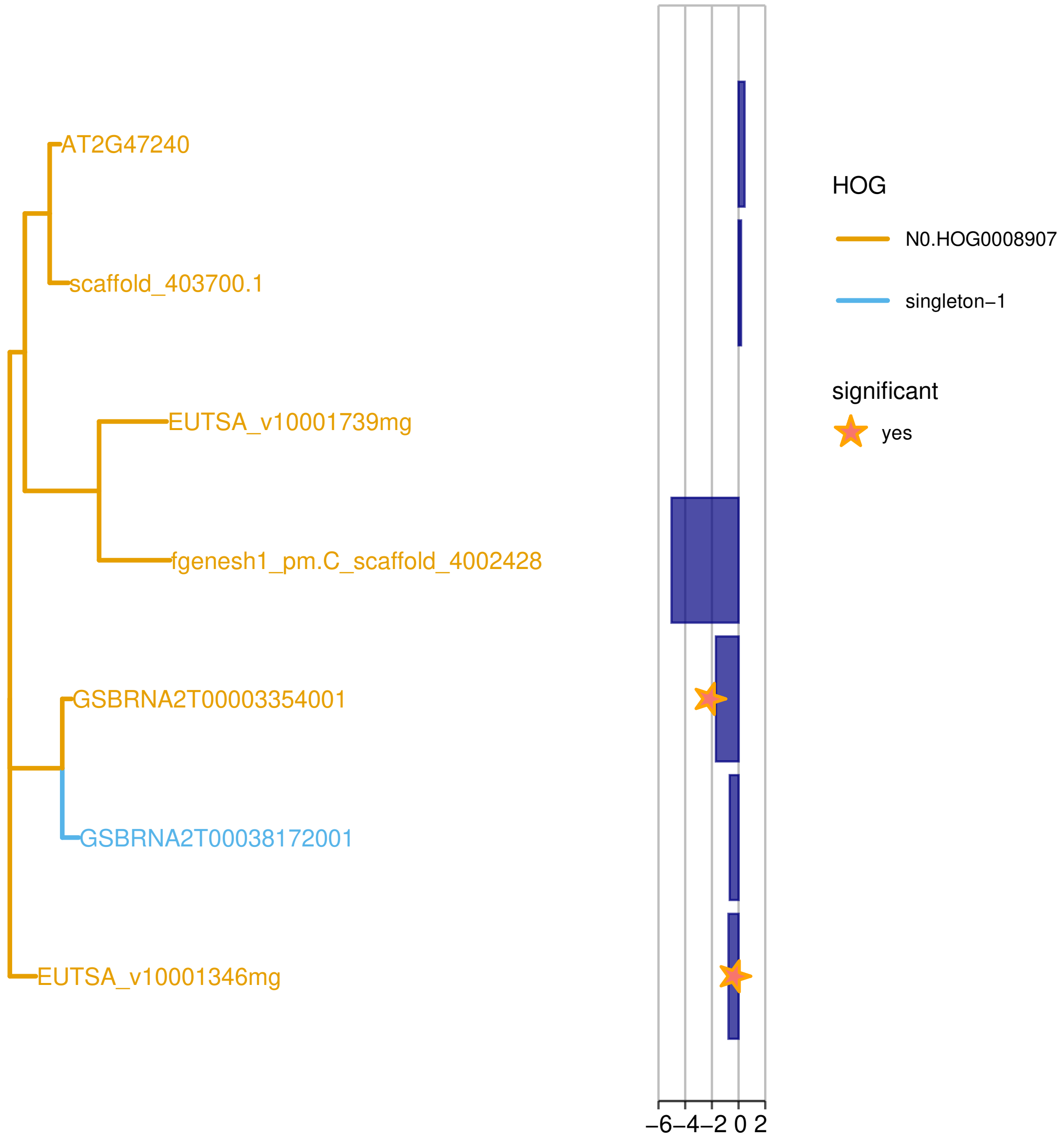

*Supplementary Figure 17: Phylogenetic tree of the genes from the four Brassicaceae species of the family N0.HOG0008907 (LACS1, CER8).*

*Blue bars represent differential expression between drought and control (log2FC), where stars indicate a corrected p-value <= 0.1). Genes with IDs starting with “EUTSA” are from Esa, with “fgenesh” or “scaffold” are from Aly, with “AT” are from Ath and with “GSBRN” are from Bna.*

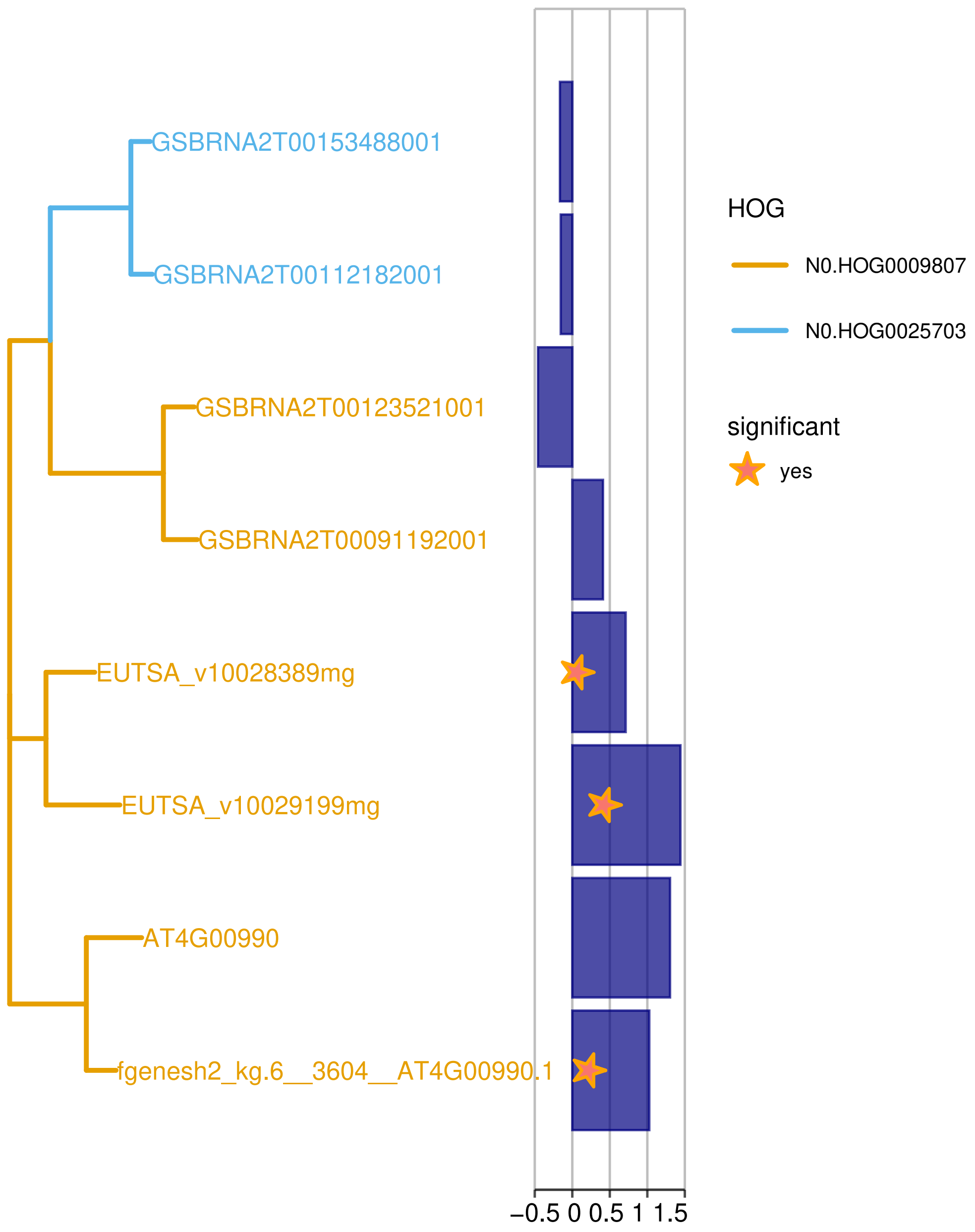

*Supplementary Figure 18: Phylogenetic tree of the genes from the four Brassicaceae species of the family N0.HOG0009807 (Lysine-specific demethylase JMJ25).*

*Blue bars represent differential expression between drought and control (log2FC), where stars indicate a corrected p-value <= 0.1). Genes with IDs starting with "EUTSA" are from Esa, with "fgenesh" or "scaffold" are from Aly, with "AT" are from Ath and with "GSBRN" are from Bna.*

##### *Subfunctionalization*

We can detect regulatory subfunctionalization when the duplicates are expressed under different conditions ([22]). We identified 12 gene families expanded in *Esa* in which one of the copies is up and at least one other copy is down regulated under drought. There are no enriched functions in these HOGs.

**Examples of up and down regulation of duplicates**

N0.HOG0002990 encodes the E3 ubiquitin-protein ligase SP1. The homologs from all species in this study are up regulated and one additional homolog from *Esa* is down regulated (Supplementary Figure 19). The two *Esa* homologs are not tandem duplicates. In *Ath*, overexpression of SP1 (AT1G63900) increased stress tolerance ([23]).

N0.HOG0015608 encodes the UDP-glycosyl- transferase 84A1. The single homolog from *Ath* and one homolog from *Esa* are up regulated (Supplementary Figure 20). Additionally, the other homolog from *Esa,* the single homolog from *Aly* and both homeologs from *Bna* are down regulated. The homologs from *Esa* are tandem duplicates. In *Ath,* UDP-glycosyl-transferase 84A1 (AT4G15480) responds to UV-B ([24]).

N0.HOG0000198, Disease resistance protein (AT5G46450, TIR-NBS-LRR class family), is duplicated in *Esa* with one homolog up and one homolog down regulated. It is also duplicated in *Aly* and one of the three *Aly* homologs is also down regulated. The other two homologs from *Aly* and the homolog from *Bna* are nearly not expressed in this experiment, while the homolog from *Ath* is constantly expressed.

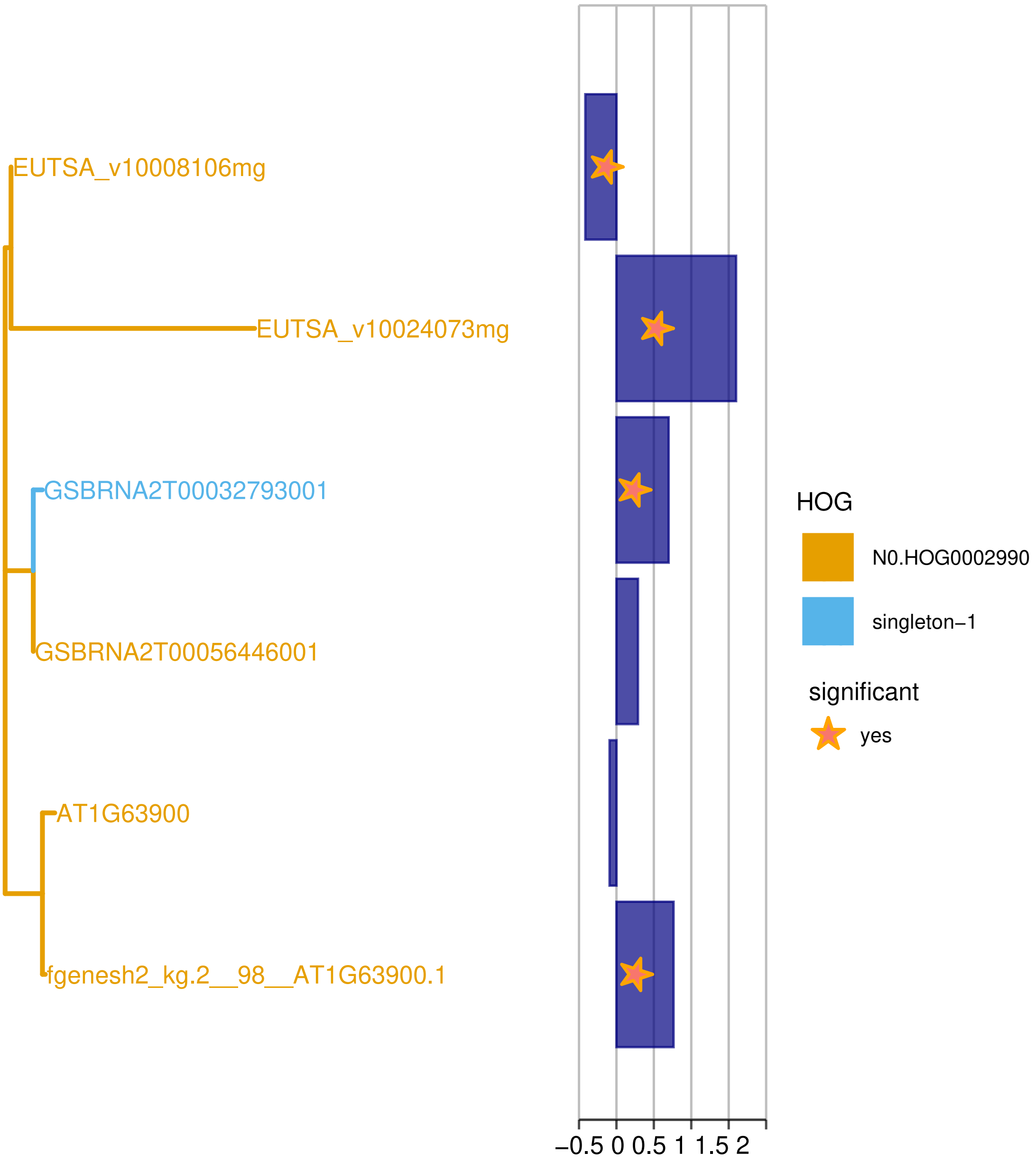

*Supplementary Figure 19: Phylogenetic tree of the genes from the four Brassicaceae species of the family N0.HOG0002990 (E3 ubiquitin-protein ligase SP1).*

*Blue bars represent differential expression between drought and control (log2FC), where stars indicate a corrected p-value <= 0.1). Genes with IDs starting with "EUTSA" are from Esa, with "fgenesh" or "scaffold" are from Aly, with "AT" are from Ath and with "GSBRN" are from Bna.*

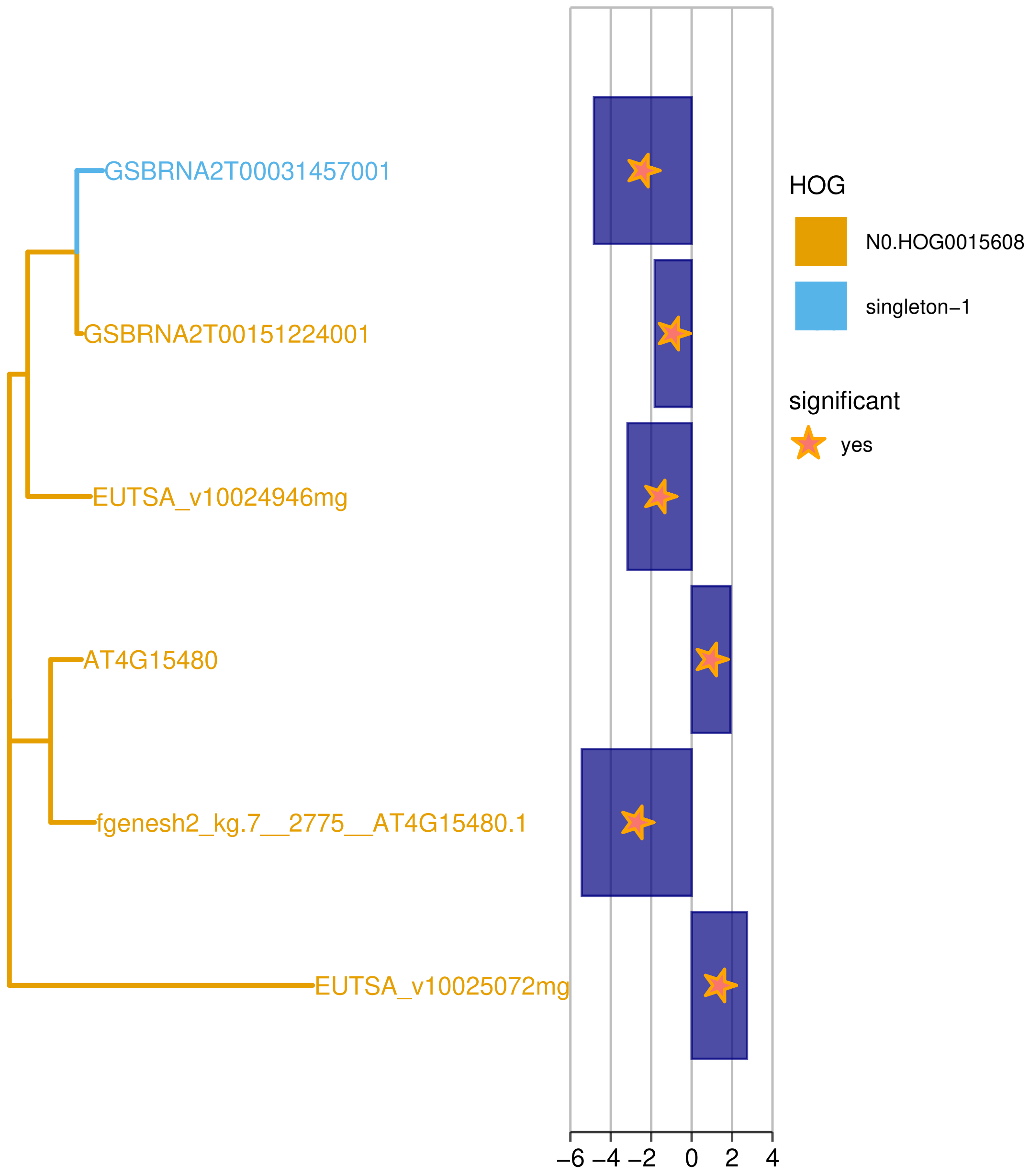

*Supplementary Figure 20: Phylogenetic tree of the genes from the four Brassicaceae species of the family N0.HOG0015608 (UDP-glycosyl-transferase 84A1) .*

*Blue bars represent differential expression between drought and control (log2FC), where stars indicate a corrected p-value <= 0.1). Genes with IDs starting with "EUTSA" are from Esa, with "fgenesh" or "scaffold" are from Aly, with "AT" are from Ath and with "GSBRN" are from Bna.*

#### Gene families which are expanded and differentially expressed in *Aly*

There are 613 gene families expanded in *Aly* which show DE in *Aly* (Supplementary Figure 4 d). Information on these gene families, including regulation under drought of the genes from all four species, their predicted functions and notes taken upon manual inspection, are provided in *Candidate_gene_families_and_tolerant_specific_expansions.xlsx*, Additional File 2. These gene families are over represented with some biological processes (Supplementary Figure 21, table with values in *GO_term_enrichment_analyses.xlsx,* Additional File 5), of which several are relevant for drought adaptation. These include the “regulation of timing of transition from vegetative to reproductive phase” (4/35, expected = 1.27, p = 0.03655), a drought escape strategy, “regulatory ncRNA-mediated post-transcriptional gene silencing” (7/55 HOGs, expected = 1.99, p = 0.01), "chromatin remodeling" (12/165 HOGs, expected = 5.97, p = 0.00397) and “response to JA” (10/185 HOGs, p = 0.03617, expected = 6.69). Interestingly, these processes are not enriched in the subset of all DEGs from *Aly* (*GO_term_enrichment_analyses.xlsx,* Additional File 5) nor in the subset of all gene families expanded in *Aly* (*GO_term_enrichment_analyses.xlsx,* Additional File 5).

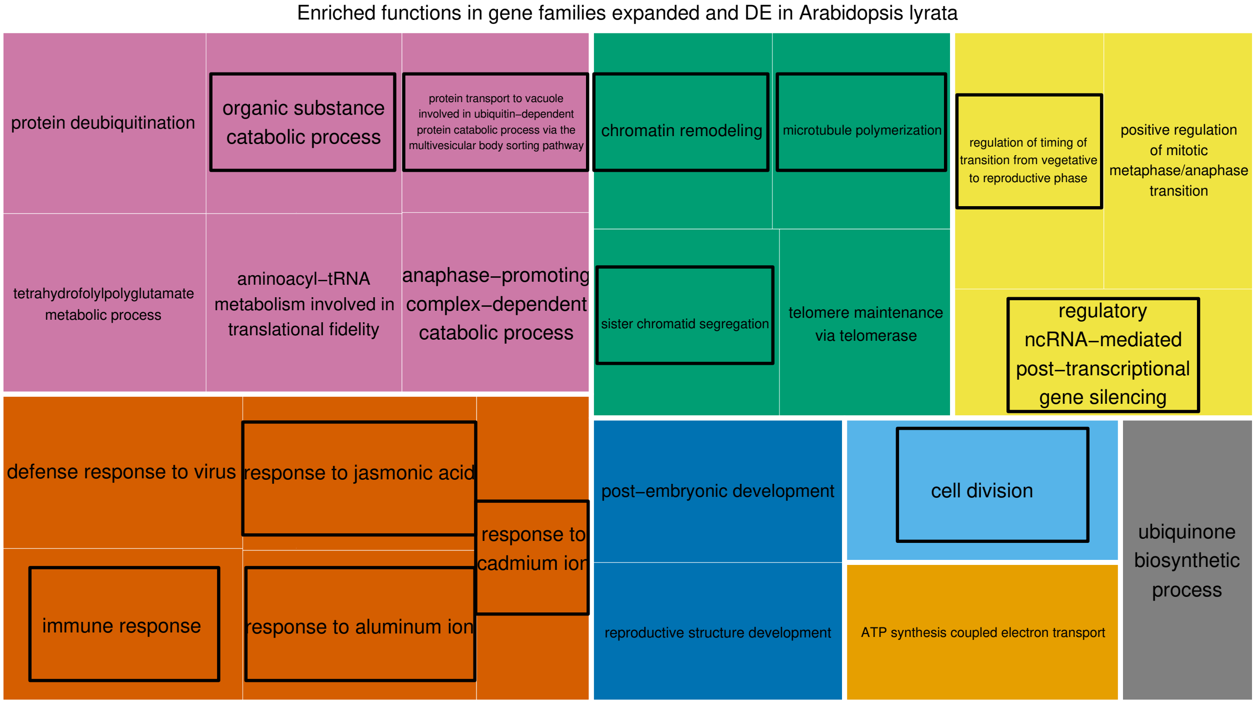

*Supplementary Figure 21: Revigo TreeMap of significantly (p <= 0.05) over represented biological processes of gene families expanded and differentially expressed in Aly.*

*Loosely related terms are clustered together. Square size is the log10(p-value) of the one-sided fisher test for over representation. Black ovals mark terms which are not enriched in all gene families either differentially expressed or expanded in Aly, but only in the combination of both. Refer to GO_term_enrichment_analyses.xlsx, Additional File 5 for tables with values.*

##### ***Example gene families with enriched functions***

One gene family (N0.HOG0008621) which is expanded and shows DE in *Aly* is NFYA4, a HAP2 (yeast HEME ACTIVATOR PROTEIN, AT2G34720) gene which is involved in the regulation of flowering in *Ath* ([25]). N0.HOG0014978, AT5G23880, which codes for subunit 2 of the Cleavage and polyadenylation specificity factor, is also involved in flowering in *Ath* ([26]). It functions in the post-transcriptional gene silencing via ncRNA and is duplicated and up regulated only in *Aly* under drought.

N0.HOG0005103 is a family of small nuclear ribonucleoprotein SmD1a which is duplicated in *Aly* and the duplicate is down regulated under drought. It is involved in post-transcriptional gene silencing in *Arabidopsis thaliana* (AT3G07590, [27]).

N0.HOG0009792 (Protein STABILIZED1, AT4G03430) is duplicated in *Aly* and one homolog is up regulated in *Aly,* one homolog is up regulated in *Bna* as well as the single homolog in *Esa*. It encodes a pre-mNA splicing factor acting in post-transcriptional gene silencing and chromatin remodeling. According to Uniprot ([11]), which lists several publications, it is involved in responses to abiotic stresses in *Ath*. We speculate that the other homolog in *Aly* could be subfunctionalized and thus function under conditions or in tissues which were not measured in this experiment.

Several other gene families expanded and differentially expressed in *Aly* are involved in chromatin remodeling: 1) the above mentioned lysin-specific demethylase REF6 (Additional Figure 13), 2) GTP-BINDING PROTEIN RELATED1 (GPR1, N0.HOG0004948, AT3G23860), which is a regulator of fertilization ([28]), 3) a Transcription factor jumonji (jmj) family protein / zinc finger (C5HC2 type) family protein (N0.HOG0008456, AT2G38950) which is not yet further characterized, 4) Transcription factor GTE12 (N0.HOG0008575, AT5G46550) which responds to ABA and sugar signaling in *Ath* ([29]), 5) Transcription initiation factor TFIID subunit 14b (N0.HOG0011415, AT5G45600, TAF14b, Supplementary Figure 22), 6) a DNA-binding bromodomain-containing protein (N0.HOG0009284, AT1G58025) and 7) a chromatin structure-remodeling complex protein BSH, “BUSHY” (N0.HOG0010933, AT3G17590). The two last mentioned gene families are both involved in the SWI/SNF complex-mediated +1 nucleosome positioning and transcription start site determination ([30]) and the resulting regulation of transcription.

One expanded and DE gene family which functions in the segregation of sister chromatids is the Sister-chromatid cohesion protein 3 gene family (N0.HOG0010279, AT2G47980), which has both homologs from *Aly* highly up regulated under drought.

Among the gene families which are involved in the reaction to the phytohormone jasmonic acid are 1) N0.HOG0003008 (AT4G32940), which encodes an Asparagine-specific endopeptidase involved in seed development in *Ath* ([31]). *Aly* has three homologs to AT4G32940, of which one is very fragmented and not expressed but both other homologs are up regulated. The homologs from *Bna* and *Ath* are also up regulated. 2) N0.HOG0008614 (AT3G11670) codes for the chloroplastic DGD1 Digalactosyldiacylglycerol synthase 1, which catalyzes the assembly of galactolipids in photosynthetic membranes, providing stability to the photosystem I (PSI) complex ([32]). Importantly, DGD1 synthase was also shown to be relevant for growth regulation by jasmonic acid: Mutants deficient in DGD synthase 1 (*dgd1*) showed reduced inflorescence stem elongation, which [33] could relate to an increased JA level and which was independent from the different chloroplast phenotype in this mutant. It mainly reduced growth in vascular tissue. N0.HOG0008614 is duplicated in *Aly* and both duplicates are up regulated. 1/5 of the AA sequence is missing at the C-terminus of one duplicate. We speculate that the up regulation can decrease the JA level and by this maintain cell cycle progression ([34]) to promote growth especially in vascular tissue. By this, the drought tolerance effect would be related to the maintenance of water and nutrient supply or inflorescence stem elongation. In *Esa*, we identified a duplication and up regulation in a gene where the mutation inhibited vein growth (see above, N0.HOG0006742). We speculate that there is a similar adaptation mechanism: up regulation would promote the growth of veins, which contain vascular tissue.

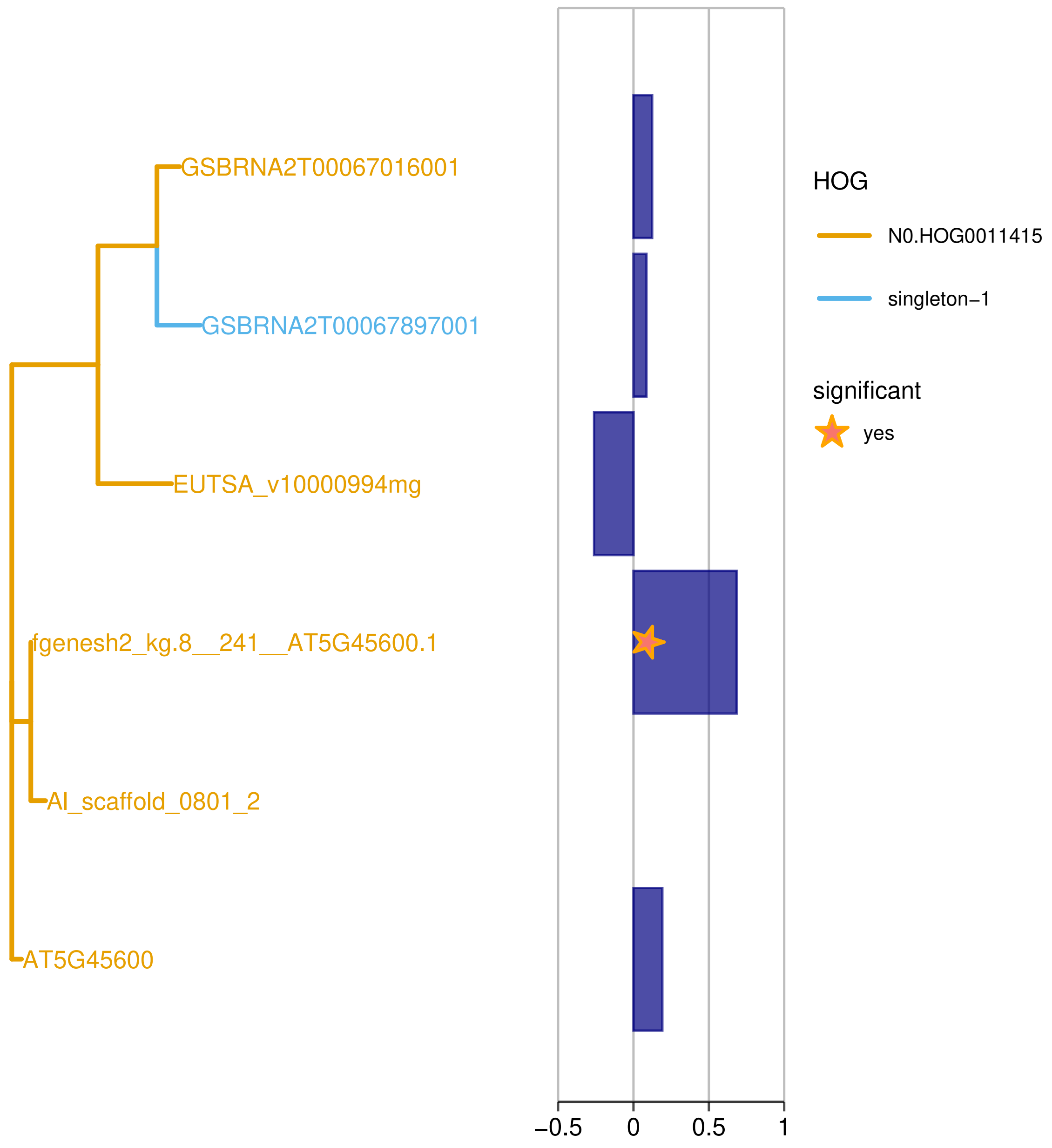

*Supplementary Figure 22: Phylogenetic tree of the genes from the four Brassicaceae species of the family N0.HOG0011415.*

*Blue bars represent differential expression between drought and control (log2FC), where stars indicate a corrected p-value <= 0.1). Genes with IDs starting with "EUTSA" are from Esa, with "fgenesh" or "scaffold" are from Aly, with "AT" are from Ath and with "GSBRN" are from Bna.*

##### *Subfunctionalization*

According to the signature of subfunctionalization described for gene families expanded in *Esa*, we identified 15 gene families expanded in *Aly* in which one of the homologs is up and at least one other homolog is down regulated under drought. There are no enriched functions in these HOGs.

**Examples of up and down regulation of duplicates**

N0.HOG0011469 encodes a family of chloroplastic Probable envelope ADP,ATP carrier protein genes (AT3G51870), which function in “photoprotection”. One homolog of *Esa*, *Bna* and *Aly*, each, is down regulated, while the second homolog from *Aly* is up regulated (Supplementary Figure 23). Interestingly, this homolog shows signatures of diversifying selection.

N0.HOG0001580 highly expanded gene family, which encodes Disease resistance protein RPS4B. The duplication is common to both *Arabidopsis* species, but the duplicates have further propagated in *Aly* (Supplement Figure 24). One of two homologs from *Ath* is up regulated and 5 of 9 homologs from *Aly* are up regulated. Interestingly, within the group of homologs which is specific for *Aly*, one homolog is down regulated and the other two homologs show very low or no expression in this experiment, which indicates that these are rather not relevant for the drought reaction. We can only speculate about what function the disease resistance gene, which is known to function in the recognition of *P. syringae* effectors ([35]), has in drought adaptation, but the expansion together with the up regulation of the duplicates under drought are strong indicators of such a function.

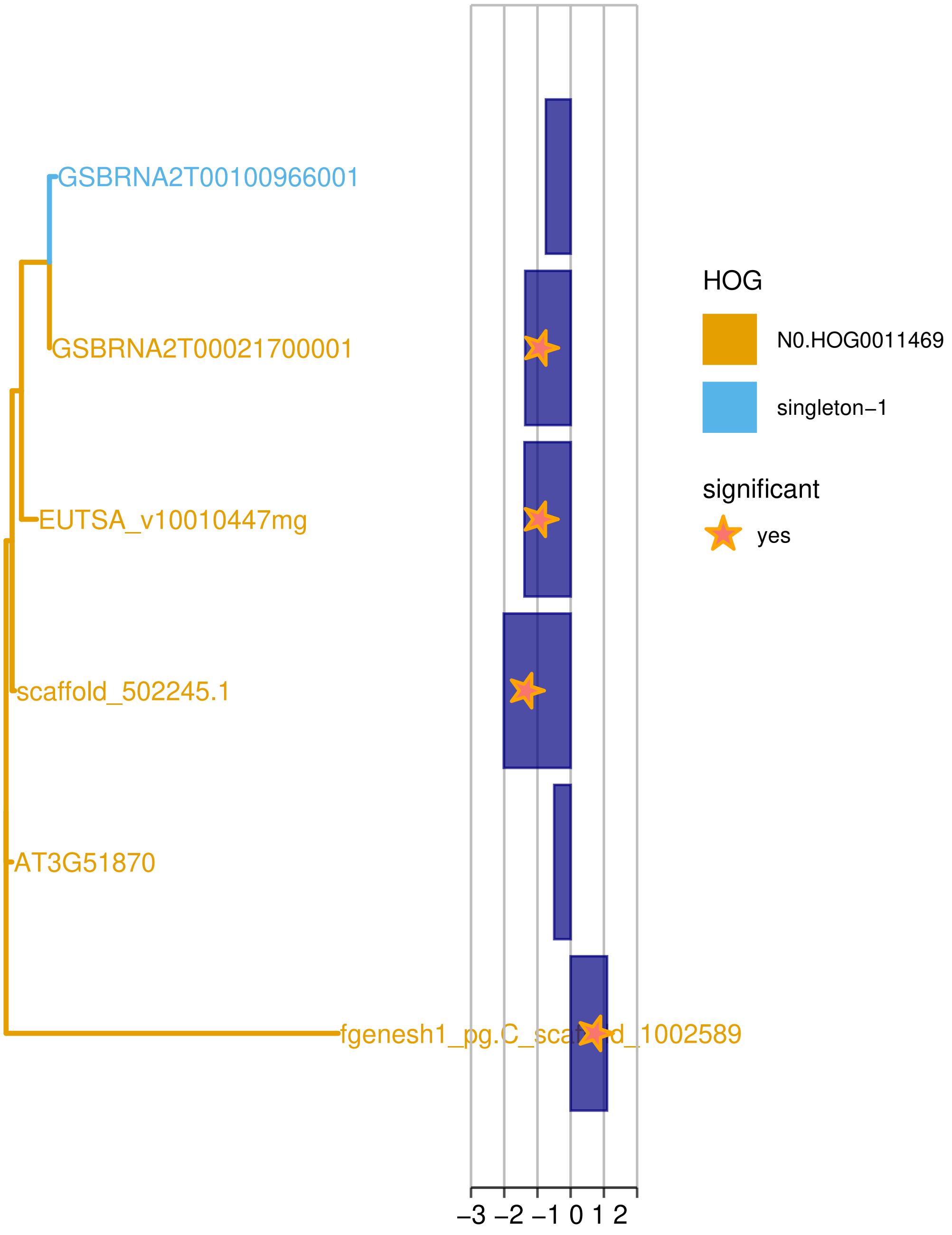

*Supplementary Figure 23: Phylogenetic tree of the genes from the four Brassicaceae species of the family N0.HOG0011469 (Probable envelope ADP,ATP carrier protein, chloroplastic).*

*Blue bars represent differential expression between drought and control (log2FC), where stars indicate a corrected p-value <= 0.1). Genes with IDs starting with "EUTSA" are from Esa, with "fgenesh" or "scaffold" are from Aly, with "AT" are from Ath and with "GSBRN" are from Bna.*

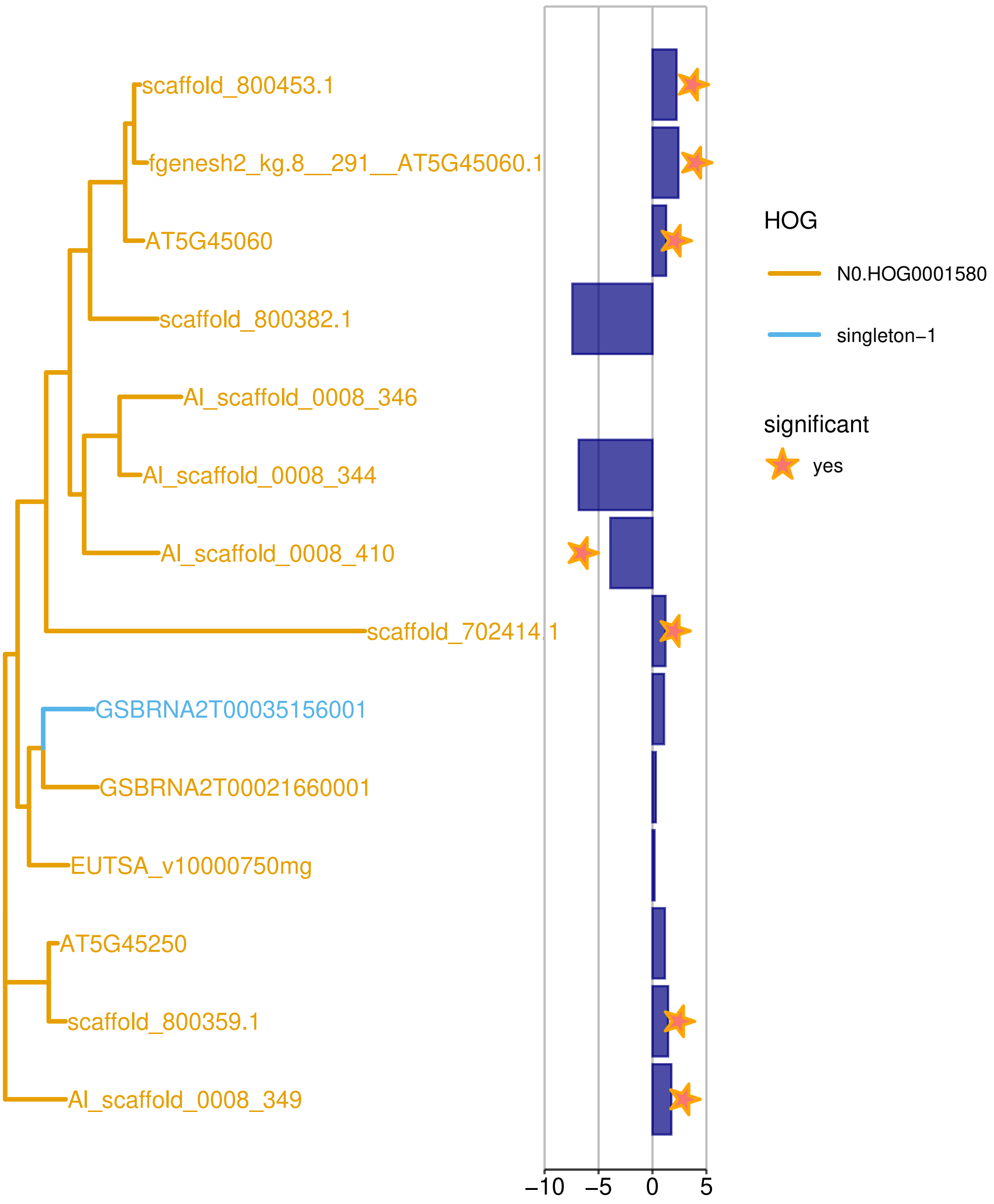

*Supplementary Figure 24: Phylogenetic tree of the genes from the four Brassicaceae species of the family N0.HOG0001580 (Disease resistance protein RPS4B).*

*Blue bars represent differential expression between drought and control (log2FC), where stars indicate a corrected p-value <= 0.1). Genes with IDs starting with “EUTSA” are from Esa, with “fgenesh” or “scaffold” are from Aly, with “AT” are from Ath and with “GSBRN” are from Bna.*

#### Gene families with exactly one up regulated duplicate in *Esa* or in *Aly*

Another form of regulatory subfunctionalization or neofunctionalization is manifested in such a way that one of the duplicates functions constantly, while another duplicate has acquired a drought stress specific function. We can observe this in the expression of the duplicates, where one duplicate is regulated under drought stress, while the other(s) is/are constantly expressed or not expressed.

In *Esa* we found 114 gene families which have exactly one of the duplicates up regulated and 91 gene families which have exactly one of the duplicates down regulated. In *Aly,* we found 225 gene families which have exactly one of the duplicates up regulated and 149 gene families which have exactly one of the duplicates down regulated.

We identified 6 (Supplementary Table 5) and 10 (Supplementary Table 6) over represented biological processes in the gene families where exactly one duplicate from *Esa* or from *Aly*, respectively, is up regulated. By comparing the over represented terms between the two groups of gene families, we found that gene families which function in cell cycle (GO:0007049) are over represented in both groups. Interestingly, the 7 gene families with this function in *Esa* do not overlap with the 17 gene families with this function in *Aly.*

*Supplementary Table 5: Significantly enriched GO terms in the subset of HOGs which are expanded in Esa and where exactly one of the duplicates is up regulated.*

| GO.ID | Term | Annotated | Significant | Expected | p-value |
|---|---|---|---|---|---|
| GO:0009303 | rRNA transcription | 11 | 2 | 0.07 | 0.0024 |
| GO:0007049 | cell cycle | 619 | 7 | 4.15 | 0.0039 |
| GO:0007059 | chromosome segregation | 133 | 3 | 0.89 | 0.0057 |
| GO:0009809 | lignin biosynthetic process | 55 | 3 | 0.37 | 0.006 |
| GO:0040031 | snRNA modification | 3 | 2 | 0.02 | 0.0066 |
| GO:0043966 | histone H3 acetylation | 20 | 2 | 0.13 | 0.0078 |

*The enrichment is calculated compared to the Conserved Set.*

*Supplementary Table 6: Significantly enriched GO terms in the subset of HOGs which are expanded in Aly and where exactly one of the duplicates is up regulated.*

| GO.ID | Term | Annotated | Significant | Expected | p-value |
|---|---|---|---|---|---|
| GO:0045842 | positive regulation of mitotic metaphase/anaphase transition | 8 | 3 | 0.11 | 0.00013 |
| GO:0035551 | protein initiator methionine removal involved in protein maturation | 4 | 2 | 0.05 | 0.00107 |
| GO:0031145 | anaphase-promoting complex-dependent catabolism | 17 | 3 | 0.23 | 0.00143 |
| GO:0030259 | lipid glycosylation | 6 | 2 | 0.08 | 0.00262 |
| GO:0006777 | Mo-molybdopterin cofactor biosynthetic process | 7 | 2 | 0.09 | 0.00364 |
| GO:0006338 | chromatin remodeling | 121 | 7 | 1.63 | 0.00438 |
| GO:0006904 | vesicle docking involved in exocytosis | 9 | 2 | 0.12 | 0.00613 |
| GO:0032012 | regulation of ARF protein signal transduction | 10 | 2 | 0.13 | 0.00759 |
| GO:0007049 | cell cycle | 619 | 17 | 8.35 | 0.00832 |
| GO:0009791 | post-embryonic development | 1426 | 23 | 19.24 | 0.00883 |

*The enrichment is calculated compared to the Conserved Set.*

##### *Examples of expanded gene families with exactly one up regulated duplicate which function in Cell Cycle*

N0.HOG0014350 encodes a family of Cyclin-dependent kinase A-1, which is expanded in *Esa* and has exactly one duplicate up regulated. It is involved in stomatal development in *Ath* (AT3G48750, [36]), which is a process relevant for drought adaptation. Furthermore, there are two kinetochore or kinetochore-associated proteins which are expanded and exactly one duplicate is up regulated in *Esa:* 1) N0.HOG0008466 (AT3G48210, Kinetochore protein SPC25 homolog) is duplicated only in *Esa*. Exactly one duplicate is up regulated and it shows signatures of diversifying selection. The orthologs from the other species are not differentially but constantly, while in *Bna* very lowly, expressed. 2) N0.HOG0009827 (AT5G06590, Spindle and kinetochore-associated protein 3) is also duplicated only in *Esa* and only the duplicate is up regulated. The orthologs from the other species are not differentially expressed, but in *Ath* and *Aly* constantly expressed, and in *Bna* not expressed.

In *Aly*, there are 3 cell division cycle proteins (Cyclin or Cyclin-dependent kinase) which are duplicated and exactly one is up regulated: N0.HOG0006690 (AT5G48630, Cyclin-C1-2) are tandem duplicates, of which one is up regulated while the other is constantly expressed. N0.HOG0008521 (AT4G05440, Cell division cycle protein 123 homolog) has one of the homologs from *Aly* up regulated. The single *Esa* ortholog is also up regulated. 3) N0.HOG0008711 (AT3G16320, Cell division cycle protein 27 homolog A) has one of two homologs from *Aly* highly up regulated while all other genes are not differentially expressed. Moreover, N0.HOG0004098 (AT5G67100, *INCURVATA2*) codes for the DNA polymerase alpha catalytic subunit (Supplementary Figure 25). This gene family is highly expanded in *Aly* with a total of 5 homologs, of which one is highly conserved with the *Ath* homolog and is similarly expressed, while exactly one other homolog is up regulated under drought. The other three homologs are not expressed in this experiment and show higher divergence. The differentially expressed homolog and the closest related duplicate as well as the one conserved with *Ath* and the single ortholog from *Esa* show signatures of diversifying selection. It has “leaf morphogenesis” (GO:0009965) annotated ([37]). DNA Polymerase alpha is essential for the initiation of replication (*[37]*) and is involved in the maintenance of histone modifications ([38]).

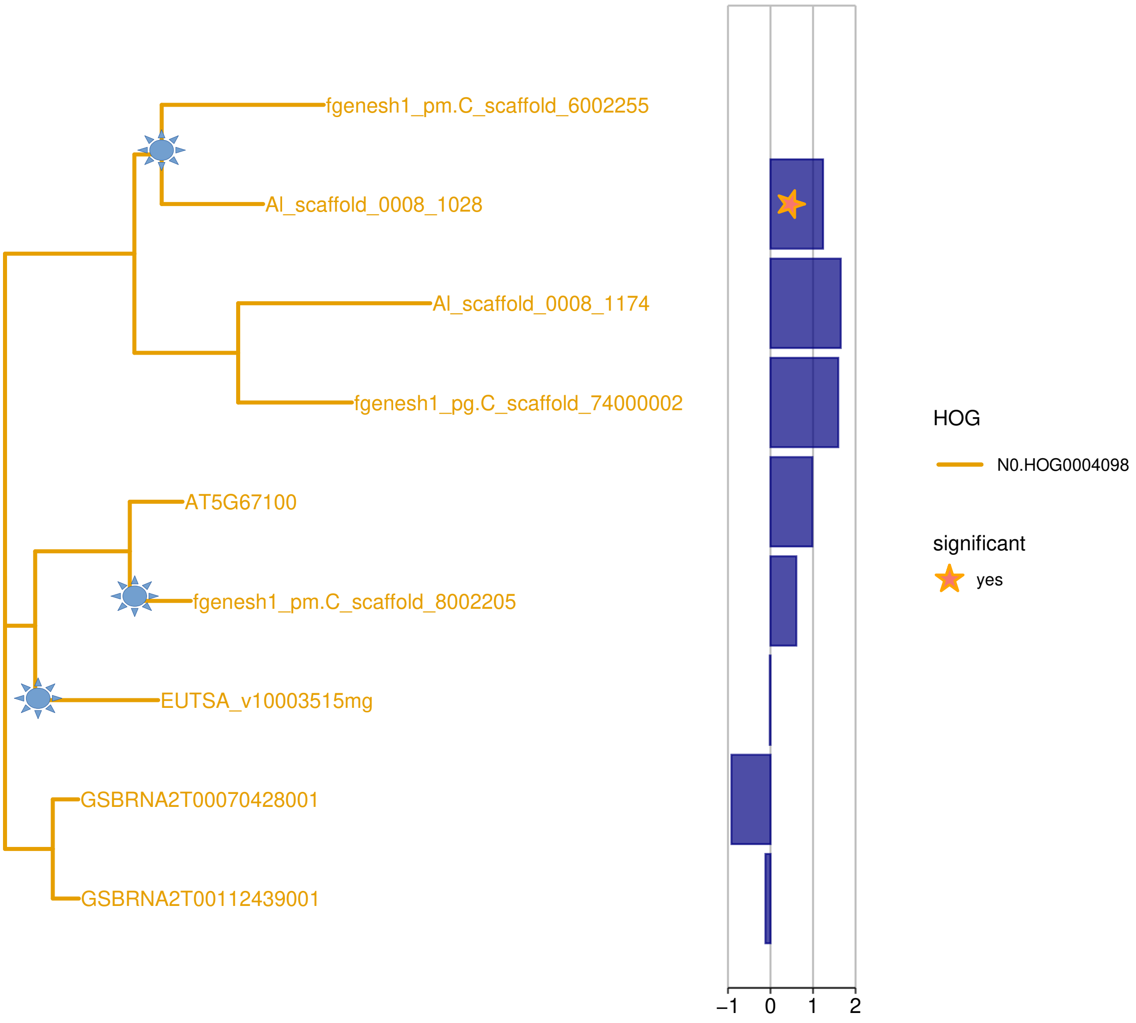

*Supplementary Figure 25: Phylogenetic tree of the genes from the four Brassicaceae species of the family N0.HOG0004098 (DNA polymerase alpha catalytic subunit).*

*Blue bars represent differential expression between drought and control (log2FC), where stars indicate a corrected p-value <= 0.1). Genes with IDs starting with "EUTSA" are from Esa, with "fgenesh" or "scaffold" are from Aly, with "AT" are from Ath and with "GSBRN" are from Bna. Sun symbols indicate genes or nodes which show signatures of diversifying selection.*

#### Diversifying selection in duplications in phylogenetically close species

*Supplementary Table 7: Number of genes for which diversifying selection was predicted by absrel ([7]) in expanded gene families.*

| Species | Gene family expansion in | Genes under diversifying selection | Genes under diversifying selection in background | $p_{upper}$ |
|---|---|---|---|---|
| *Bna* | *Bna* | 544 (10 %) | 2732 (11 %) | 0.07 |
| *Bna* or *Esa* | *Bna* and *Esa* | 34 (10 %) | 4103 (11 %) | 0.46 |
| *Ath* or *Aly* | *Ath* and *Aly* | 141 (13 %) | 2588 (9 %) | 6.9e-06 |
| *Ath, Aly* or *Esa* | *Ath, Aly* and *Esa* | 217 (12 %) | 3959 (9 %) | 5.7e-05 |

*For the given species, the number of genes showing diversifying selection is given in families expanded in the given species but not in any of the other species. Pupper = p-value from hypergeometric test for over representation in the set of expanded gene families, compared to the background of all genes from the given species in the Conserved Set.*

#### Diversifying selection in DEGs in gene families expanded in *Esa* and/or *Aly*

##### In *Esa* expanded gene families which have a DEG from *Esa* under diversifying selection

There are 39 gene families expanded in *Esa* with a total of 41 DEGs from *Esa* which show signatures of positive selection. Manual checking of the 39 HOGs showed that 16 (41 %) of them are most likely also expanded in a sensitive species or the duplicate is pseudogenizing because the multiple sequence alignment (MSA) of the peptide sequences is fragmented in one of the duplicates. Additionally, we cannot interpret the test for diversifying selection for 5 (12.8 %) additional HOGs, because the peptide MSA is strongly fragmented.

There are 18 (46 %) promising gene families with a total of 20 DEGs from *Esa* under diversifying selection. Detailed information including notes from manual evaluation are displayed in *Candidate_gene_families_and_tolerant_specific_expansions.xlsx*, Additional File 2, while we describe the most promising HOGs here.

Two interesting HOGs have two DEGs from *Esa* which show signatures of diversifying selection: The lysine-specific demethylase REF6 (N0.HOG0007350), which was already mentioned previously (Supplementary Figure 13) and N0.HOG0009382, which encodes a family of E3 ubiquitin-protein ligase PRT1 (PROTEOLYSIS1, AT3G24800, Supplementary Figure 26). Both homologs from *Esa* are up regulated and show signatures of diversifying selection.

Furthermore, there are several HOGs in which one DEG from *Esa* shows signatures of diversifying selection: 1) N0.HOG0005668 (AT1G6496, *HEB1*) codes for the Condensin II subunit CAP-G2. There are 4 homologs from *Esa*, all are differentially expressed and one of them shows signatures of diversifying selection. The single homolog from *Aly* is also differentially expressed. Over the whole MSA, there are several insertions and deletions, but these are not specific to the gene which shows signatures of diversifying selection, indicating that the amino acid sequences in this HOG are highly diverged. 2) Moreover, the already mentioned E3 UFM1-protein ligase 1 homolog (N0.HOG0008832, Supplementary Figure 9), which is expanded and up regulated in both tolerant species and functions in “modification-dependent protein catabolic process”, shows signatures of diversifying selection in the up regulated duplicate from *Esa* and in the non-regulated duplicate from *Aly*. 3) N0.HOG0009027 (AT1G12440) encodes a family of Zinc finger A20 and AN1 domain-containing stress-associated protein 1, which is duplicated in *Esa*. Both homologs, of which one shows signatures of diversifying selection, are down regulated. The single homolog from *Aly* is also down regulated. The homologs from the other species are not regulated. 4) N0.HOG0009134 (AT1G35530) encodes a DEAD/DEAH box RNA helicase family protein which has only both duplicates from *Esa* up regulated and one of them (EUTSA_v10001810mg) shows signatures of diversifying selection. 5) N0.HOG0009602 (AT3G54220, SCARECROW) encodes a transcription factor which regulates the radial pattern formation in roots and the radial organization of the shoot axial organs. It is expanded only in *Esa* and the differentially expressed homolog shows signatures of diversifying selection. 6) N0.HOG0012797 encodes Cryptochrome-2 (AT1G04400). It functions in leaf development and flowering time regulation via circadian clock. It is expanded only in *Esa* and up regulated in *Esa*, *Aly* and *Bna*. 7) N0.HOG0010317 encodes the uncharacterized protein OPA3-like protein (AT3G58150). Both homologs from *Esa* are up regulated. The homolog under diversifying selection (EUTSA_v10019778mg) is much more expressed than the other duplicate. The only homolog from *Aly* is also up regulated. 8) N0.HOG0015172 encodes an uncharacterized Protein (AT4G01870). Only the differentially expressed homolog from *Esa* is under diversifying selection. 9) We identified a candidate gene family which is known to be relevant for salt tolerance in *Esa*

[39]Monihan and colleagues showed that the duplication in the calcium sensor ATCBL10 is relevant for the salt tolerance of *Esa*. We identified this gene family, N0.HOG0009973, among our *Esa* specific candidate genes for drought adaptation: both homologs from *Esa* and the single homolog from *Ath* are down regulated under drought and additionally, EUTSA_v10026019mg shows signatures of diversifying selection. However, we would exclude this family upon manual inspection as there is also a duplication in the sensitive species *Bna*.

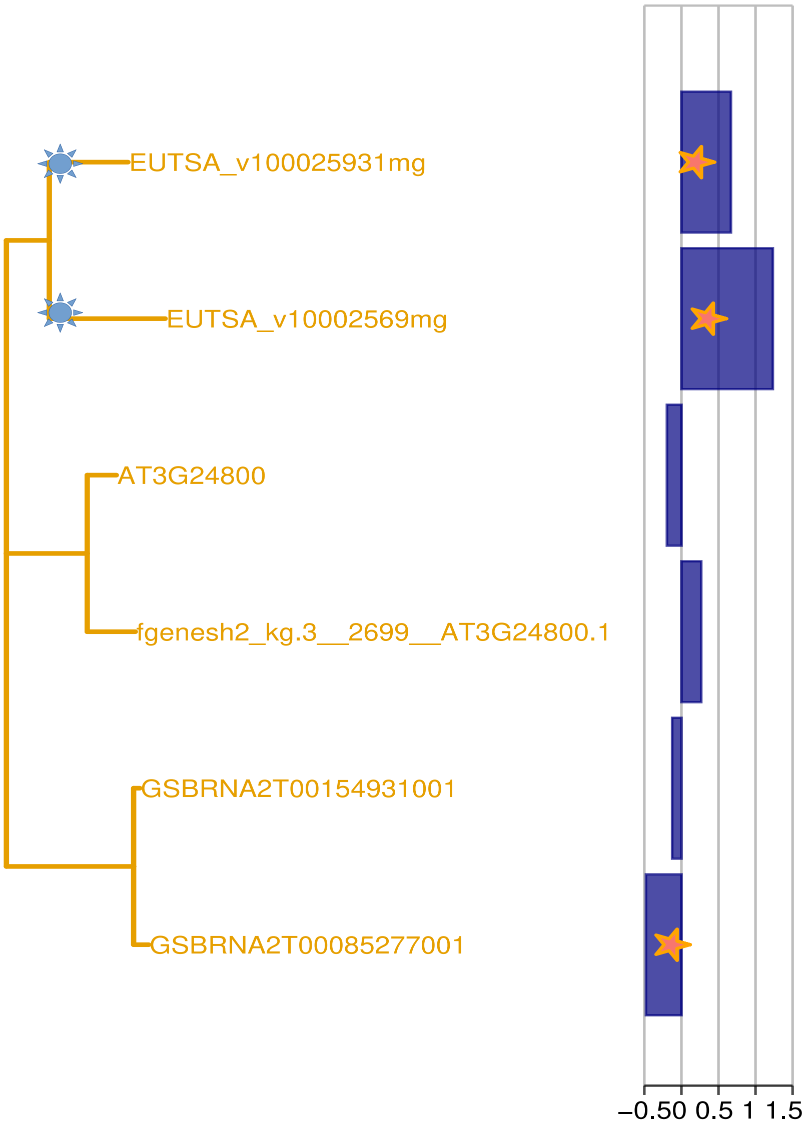

*Supplementary*

*Figure 26: Phylogenetic tree of the genes from the four Brassicaceae species of the family N0.HOG0009382 (E3 ubiquitin-protein ligase PRT1).*

*Blue bars represent differential expression between drought and control (log2FC), where stars indicate a corrected p-value <= 0.1). Genes with IDs starting with "EUTSA" are from Esa, with "fgenesh" or "scaffold" are from Aly, with "AT" are from Ath and with "GSBRN" are from Bna. Sun symbols indicate genes or nodes which show signatures of diversifying selection.*

##### In *Aly* expanded gene families which have a DEG from *Aly* under diversifying selection

There are 46 gene families expanded in *Aly* in which a DEG from *Aly shows* signatures of positive selection. Some interesting examples include 1) N0.HOG0004098, AT5G67100 (*INCURVATA2*) DNA polymerase alpha catalytic subunit, which is highly expanded in *Aly* and the only differentially expressed homolog from Aly and the single ortholog from *Esa* show signatures of diversifying selection (see above and Supplementary Figure 25). 2) **N0.HOG0008711** (AT3G16320, Cell division cycle protein 27 homolog A) functions in “anaphase-promoting complex-dependent catabolic process” and “positive regulation of mitotic metaphase/anaphase transition”. The up regulated homolog from *Aly* is under positive selection as well as one homolog from *Bna.* 3) N0.HOG0006612 encodes a cytoplasmic isoleucin-tRNA ligase (AT4G10320), which functions in “aminoacyl-tRNA metabolism involved in translational fidelity”. It has two duplications in *Aly*. Two of the three homologs are up regulated while the most diverged duplicate is not expressed in this experiment. All three homologs from *Aly* show signatures of positive selection.

#### Supplementary Discussion

##### Candidate gene families are enriched for genes which function in post translational regulation

Candidate gene families are uniquely enriched for “modification-dependent protein catabolism”. Similarly, [1] found that “ubiquitin-dependent protein modification” is the only enriched process in the gene families which are expanded in *Esa* compared to *Ath* using the program “i-ADHoRe 2.0” ([40], latest version is 3.0) to calculate the gene family expansion. We found “ubiquitin-dependent protein catabolic process”, which is a modification-dependent protein catabolism, enriched in the gene families which were expanded in both tolerant species or only in *Aly* regardless of a potential regulation under drought. “Protein ubiquitination” and “positive regulation of proteasomal ubiquitin-dependent protein catabolic process”, which regulates “modification-dependent protein catabolism”, were also enriched in the gene families which are expanded in *Esa.*

Importantly, within all gene families which are DE in any or in both tolerant species, no posttranslational regulatory process was enriched. The post transcriptional regulation of active protein abundance, mediated by E3 protein ligases, is a crucial regulatory step in most processes in plants (reviewed in [41]). F-box genes code for the substrate recognition sites of the SCF ubiquitin ligases and are organized into hundreds of F-box gene families. Some of these protein families show highly diverged domain organization, which indicates their relevance for adaptation ([42]). The expansion rate in the highly duplicating F-box gene families is even higher than in other rapidly duplicating genes such as receptor-like kinase genes or disease resistance (R) genes ([42]). It is clear that "modification-dependent protein catabolism" is a regulatory process important for adaptation as seen by the intensive duplication and domain shuffling in substrate recognition proteins. The results of the GO term enrichment analysis of expanded gene families from [1], which used a different method to define gene family expansion in *Esa* as we did, are comparable to ours. But beyond this, we could link this function to the evolution of drought tolerance because some of the gene families with this function are expanded in two drought adapted plant species and DE under drought. We could distinguish the drought adaptive genes, of which several function in post translational modification, from the overall drought reacting genes. The gene families in our study which do not react under drought but are expanded are most likely also adaptive but for other phenotypic differences between the studied species.

To summarize, gene families expanded in *Aly* and/or *Esa* are enriched for genes which function in post translational modification, and part of these are associated to drought adaptation because they are expanded in both drought adapted species and DE under drought.

#### Functions of species-specific candidate genes are diverse and some functions which are known for their relevance in drought tolerance and others which are not, are enriched in the species-specific candidate genes.

We identify enrichment of diverse processes in species-specific candidate genes of which some are already implicated in drought tolerance while others are not. We find stomatal complex development uniquely enriched in gene families which are expanded and DE in

*Esa* (*Esa*-specific candidate genes). But upon closer visual inspection, we find additional orthologs in *Bna* which were not predicted by Orthofinder. We manually verify that genes which function in stomatal complex development are not uniquely expanded in a tolerant species, but they might be adaptive for a conserved tolerance mechanism between *Bna* and *Esa*. Based on the functional annotation, several of the *Esa*-specific candidates function in processes which have already been implicated in drought tolerance.

*Esa*-specific candidate genes significantly more often function in “wax biosynthetic process” and “stomatal complex development”, processes which are relevant for drought avoidance traits. Intriguingly, [43] declares that the reduction of non-stomatal water loss is among the most important trait for plant production under drought because in contrast to reducing stomatal water loss by smaller stomatal aperture, it does not correlate with photosynthate assimilation. The relevance of the leaf wax composition for the amount of cuticular transpiration was shown e.g. in *Populus x canescens cer6* mutants ([44]) and in *Hordeum vulgare KAS1* mutants ([45]). The likewise halophytic relatives of *Esa* *Thelungiella parvula* and *Thelungiella halophila* show different leaf wax composition than *Ath* ([46]). It is hence probable that *Esa* has evolved a cuticular wax composition which could reduce the cuticular transpiration. Our *Esa*-specific candidate genes which function in wax biosynthetic process are likely relevant for drought avoidance by altered leaf wax composition in *Esa*.

*Esa*-specific candidate genes were significantly enriched for genes which function in “stomatal complex development”. Even though stomatal closure reduces photosynthesis, which leads to growth reduction in long periods of drought, it is a beneficial avoidance mechanism for short drought events ([47]). In gene families which function in stomatal complex development, duplications in *Esa* seem to be conserved with *Bna*, which suggests that they are relevant for common traits between the two species. On the other hand, the regulation under drought differs between the two species, which could reflect differences in the stress levels of the species, or actual differences in the velocity the plant reacts to drought. Moreover, the duplicates in *Bna* could be relevant for other than drought adaptation traits. It has been shown in several plant species that stomata reopen only partially after dehydration stress during watered recovery periods ([48], [49]). Beyond this drought reaction, the tolerant species *Aly* and *Esa* have their stomata less open even under well watered conditions and can react faster to water limitation ([5]). In *Ath* and *Aly,* a decrease in stomata size correlates with an increased water use efficiency under drought ([50], [51]), which is a drought avoidance trait but which might not necessarily be relevant for improved crop production ([43]). Yet, how stomata related traits can enhance drought tolerance without limiting plant production is not clear.

Candidate genes expanded and DE in *Aly* (*Aly*-specific candidate genes) are enriched for genes which function in “cell division”. Strikingly, [5] found that the growth reduction of leaves under drought was based on a higher reduction in cell proliferation in *Aly* compared to *Ath* and *Esa*. We can correlate our predicted *Aly*-specific candidate genes to this phenotypic observation. Two independent experiments showed that *Aly* responds earlier with growth reduction than *Ath* and recovers better at rewatering ([52], [5]). Moreover, [52] found that even in wilted leaves the photosynthetic capacity was relatively high in *Aly*. Based on these observations, we speculate that the early reduction in cell proliferation can keep the cells functional (quality over quantity) and facilitate the recovery at rewatering.

*Aly*-specific candidate genes are enriched for genes which function in “chromatin remodeling” and *Esa*-specific candidate genes are enriched for genes which function in “epigenetic regulation of gene expression”, which is a chromatin remodeling process. Several stress responses are activated by chromatin remodeling (reviewed in [53], [54]). For example, changes in the regulation of the cell-cycle can result in its arrest, leading to reduced growth under drought ([55]). The epigenetic regulators in *Aly* likely target, among many others, cell division genes as was discussed above. Another function of our candidate genes which are involved in chromatin remodeling might be epigenetic priming, a mechanism which is involved in stress tolerance ([56], [57], [58]). Epigenetically primed genes, also called memory genes ([59]), respond faster and / or stronger upon repeated stress ([53], [60], [61], [54]). We speculate that epigenetic priming is a relevant process for drought adaptation in *Aly* and in *Esa.* It is likely that several of our species-specific candidate genes are target genes for epigenetic priming (memory genes) in the tolerant species, as duplicated genes whose products are part of regulatory networks are more often retained ([62], [63], [64]).

*Esa*-specific candidate genes were also enriched for “regulation of ethylene-activated signaling pathway”. Several studies have shown that the modification of ethylene signaling pathway can enhance drought tolerance (reviewed in [65]).

The here mentioned processes are adaptations to drought, some of which differ between the two species.

To sum up, we identified several processes which are implicated in drought tolerance enriched in our species-specific candidate genes. We propose that the species-specific candidate genes which function in processes whose relevance has not yet been implicated in drought tolerance are likely to be relevant, too.

24. Ulm R, Baumann A, Oravecz A, Máté Z, Ádám É, Oakeley EJ, Schäfer E and Nagy F. Genome-wide analysis of gene expression reveals function of the bZIP transcription factor

HY5 in the UV-B response of Arabidopsis. Proceedings of the National Academy of Sciences. 2004;101:1397-1402. DOI: 10.1073/pnas.0308044100

53. Luo M, Liu X, Singh P, Cui Y, Zimmerli L and Wu K. Chromatin modifications and remodeling in plant abiotic stress responses. Biochimica et Biophysica Acta (BBA)-Gene

Regulatory Mechanisms. 2012;1819:129-136. DOI: 10.1016/j.bbagrm.2011.06.008
