## Additional File 3 for "Integration of differential expression under drought with gene family expansion unique to drought tolerant species predicts candidate genes for drought adaptation in Brassicaceae species": Analysis_A2TEA_output.R

```
# script to explore a2tea output BR-dr-03
library(Biostrings)
library(ggplot2)
library(reshape2)
library(tidyr)
library(dplyr)

setwd("/home/cropbio/sciebo/Documents/PhD/Brassicaceae-drought/BR-dr-03")
OUTDIR=paste0(getwd(), "/Analysis")
load("A2TEA_finished.RData")

#use the color palette Tyll used in the A2TEA publication
palette_OkabeIto_black <- c("#E69F00", "#56B4E9", "#009E73", "#F0E442", 
                            "#0072B2", "#D55E00", "#CC79A7", "#000000")

# write Conserved_dataset_matrix.tsv
Conserved <- HOG_level_list$hypothesis_1[HOG_level_list$hypothesis_1$Arabidopsis_lyrata_total>=1 & HOG_level_list$hypothesis_1$Arabidopsis_thaliana_total>=1 & HOG_level_list$hypothesis_1$Eutrema_salsugineum_total>=1 & HOG_level_list$hypothesis_1$Brassica_napus_total>=1,]$HOG
Dataset_matrix <- HOG_DE.a2tea[,c("HOG", "gene")]
Conserved_dataset_matrix <- Dataset_matrix[Dataset_matrix$HOG %in% Conserved,]
Conserved_dataset_matrix_ordered <- Conserved_dataset_matrix[order(Conserved_dataset_matrix$HOG),]

write.table(Conserved_dataset_matrix_ordered, paste0(OUTDIR, "/Conserved_dataset_matrix.tsv"), sep="\t", row.names=F, col.names=T, quote=F)


### Export tables which contain the GO terms and HRD for interesting sets of HOGs:
## save tables of these HOGs together with the genes, the respective GO terms and Human Readable Description:
my_list_of_interesting_HOGs <- (read.table("Analysis/GO_term_enrichment_analysis_webapp/sigs_OG level_GO0010025_BP_wax_biosyntheitic_process_enriched_in_Hypothesis_2_exp_and_DE.tsv", sep = "\t", header =T))$OG
Interesting_HOGs_and_genes_with_GO_and_HRD <- rbind(SFA$Arabidopsis_lyrata[SFA$Arabidopsis_lyrata$HOG %in% my_list_of_interesting_HOGs,],
                                                                       SFA$Arabidopsis_thaliana[SFA$Arabidopsis_thaliana$HOG %in% my_list_of_interesting_HOGs,],
                                                                       SFA$Eutrema_salsugineum[SFA$Eutrema_salsugineum$HOG %in% my_list_of_interesting_HOGs,],
                                                                       SFA$Brassica_napus[SFA$Brassica_napus$HOG %in% my_list_of_interesting_HOGs,])
Interesting_HOGs_and_genes_with_GO_and_HRD_ordered_by_HOG <- Interesting_HOGs_and_genes_with_GO_and_HRD[order(Interesting_HOGs_and_genes_with_GO_and_HRD$HOG),]
# name the table and export it
write.table(Interesting_HOGs_and_genes_with_GO_and_HRD_ordered_by_HOG, paste0("Analysis/GO_term_enrichment_analysis_webapp/", "sigs_OG level_GO0010025_BP_wax_biosyntheitic_process_enriched_in_Hypothesis_2_exp_and_DE_with_GOs_and_HRD.tsv"), sep="\t", row.names = F, col.names = T)


# Number of Expanded OGs per Hypothesis
hypotheses_names <- c()
hypotheses <- c()
sizes <- numeric()

for(x in HYPOTHESES.a2tea){
sizes <- append(sizes, length(x@expanded_OGs))
}

for (y in names(HYPOTHESES.a2tea)){
  hypotheses_names <- append(hypotheses_names, y)
}

for (z in HYPOTHESES.a2tea){
  hypotheses <- append(hypotheses, z@description)
}

sizes
# 144  787 1173   58  397 1865
hypotheses_names
hypotheses

HOGs_Eutrema <- HOG_level_list$all_species_overview[HOG_level_list$all_species_overview$Eutrema_salsugineum_total>0 ,]$HOG
HOGs_Alyrata <- HOG_level_list$all_species_overview[HOG_level_list$all_species_overview$Arabidopsis_lyrata_total >0 ,]$HOG
HOGs_Athaliana <- HOG_level_list$all_species_overview[HOG_level_list$all_species_overview$Arabidopsis_thaliana_total >0 ,]$HOG
HOGs_Bnapus <- HOG_level_list$all_species_overview[HOG_level_list$all_species_overview$Brassica_napus_total >0 ,]$HOG


### Plot the Intersect of all HOGs for all 4 species in an upset plot
library(UpSetR)
listInput <- list(Eutrema_salsugineum=HOGs_Eutrema,
                  Arabidopsis_lyrata=HOGs_Alyrata,
                  Arabidopsis_thaliana=HOGs_Athaliana,
                  Brassica_napus=HOGs_Bnapus)
sets <- c("Eutrema_salsugineum", "Arabidopsis_lyrata", "Arabidopsis_thaliana", "Brassica_napus")
names(listInput) <- c("Eutrema salsugineum",
                      "Arabidopsis lyrata",
                      "Arabidopsis thaliana",
                      "Brassica napus")


pdf(paste0(OUTDIR, "/UpSetPlot_HOGs_per_species_in_Total.pdf"), 10, 6, onefile = FALSE)
upset(fromList(listInput), nsets=4, nintersects=NA,
      set_size.show = TRUE,
      set_size.scale_max = 35000,
      sets.x.label = "Number of HOGs",
      mainbar.y.label = "Size of Intersection",
      keep.order = T, order.by = "freq",
      #      sets = sets, 
      number.angles = 0, 
      #show.numbers = T,
      point.size = 2, line.size = 1,
      text.scale = c(2,1.5,1.5,1.5,1.5,2),
      queries= list(list(query = intersects, params=c("Arabidopsis lyrata","Eutrema salsugineum", "Arabidopsis thaliana", "Brassica napus"), color= palette_OkabeIto_black[5], active = T)
      )
)
dev.off()


                      ### Select the CONSERVED data set!!! ###
### Select only conserved HOGs (at least 1 gene from each (of the compared) species)
HOG_level_list$hypothesis_1_cons <- HOG_level_list$hypothesis_1[HOG_level_list$hypothesis_1$Arabidopsis_lyrata_total>=1 & HOG_level_list$hypothesis_1$Arabidopsis_thaliana_total>=1 & HOG_level_list$hypothesis_1$Eutrema_salsugineum_total>=1 & HOG_level_list$hypothesis_1$Brassica_napus_total>=1,]
HOG_level_list$hypothesis_2_cons <- HOG_level_list$hypothesis_2[HOG_level_list$hypothesis_2$HOG %in% HOG_level_list$hypothesis_1_cons$HOG,]
HOG_level_list$hypothesis_3_cons <- HOG_level_list$hypothesis_3[HOG_level_list$hypothesis_3$HOG %in% HOG_level_list$hypothesis_1_cons$HOG,]
HOG_level_list$hypothesis_4_cons <- HOG_level_list$hypothesis_4[HOG_level_list$hypothesis_4$HOG %in% HOG_level_list$hypothesis_1_cons$HOG,]
HOG_level_list$hypothesis_5_cons <- HOG_level_list$hypothesis_5[HOG_level_list$hypothesis_5$HOG %in% HOG_level_list$hypothesis_1_cons$HOG,]
HOG_level_list$hypothesis_6_cons <- HOG_level_list$hypothesis_6[HOG_level_list$hypothesis_6$HOG %in% HOG_level_list$hypothesis_1_cons$HOG,]

#### How large is the effect if we select for Bnapus>=2 bc. Bnapus is tetraploid?
HOG_level_list$hypothesis_1_cons_2 <- HOG_level_list$hypothesis_1[HOG_level_list$hypothesis_1$Arabidopsis_lyrata_total>=1 & HOG_level_list$hypothesis_1$Arabidopsis_thaliana_total>=1 & HOG_level_list$hypothesis_1$Eutrema_salsugineum_total>=1 & HOG_level_list$hypothesis_1$Brassica_napus_total>=2,]
length(HOG_level_list$hypothesis_1_cons_2$HOG)
# 13555
# instead of
length(HOG_level_list$hypothesis_1_cons$HOG)
# 17488
#### select for Bnapus>=2 bc. Bnapus is tetraploid
HOG_level_list$hypothesis_2_cons_2 <- HOG_level_list$hypothesis_2[HOG_level_list$hypothesis_2$HOG %in% HOG_level_list$hypothesis_1_cons_2$HOG,]
HOG_level_list$hypothesis_3_cons_2 <- HOG_level_list$hypothesis_3[HOG_level_list$hypothesis_3$HOG %in% HOG_level_list$hypothesis_1_cons_2$HOG,]
HOG_level_list$hypothesis_4_cons_2 <- HOG_level_list$hypothesis_4[HOG_level_list$hypothesis_4$HOG %in% HOG_level_list$hypothesis_1_cons_2$HOG,]
HOG_level_list$hypothesis_5_cons_2 <- HOG_level_list$hypothesis_5[HOG_level_list$hypothesis_5$HOG %in% HOG_level_list$hypothesis_1_cons_2$HOG,]
HOG_level_list$hypothesis_6_cons_2 <- HOG_level_list$hypothesis_6[HOG_level_list$hypothesis_6$HOG %in% HOG_level_list$hypothesis_1_cons_2$HOG,]
# --> I did not use this "cons_2" dataset in the Analysis. This was only out of interest to check the effect.


#### How many HOGs are expanded in 1) Athal, Alyrata and Eutrema vs. Bnapus and 2) Athal and Alyrata vs. Eutrema and Bnapus? To find out, how important the phylogenetic 
# relationship is for the identification of gene duplications

HOGs_exp_3vsBnapus <- HOG_level_list$hypothesis_1_cons[HOG_level_list$hypothesis_1_cons$Arabidopsis_lyrata_total >= 2* (HOG_level_list$hypothesis_1_cons$Brassica_napus_total/2) &
                                  HOG_level_list$hypothesis_1_cons$Eutrema_salsugineum_total >= 2* (HOG_level_list$hypothesis_1_cons$Brassica_napus_total/2) &
                                   HOG_level_list$hypothesis_1_cons$Arabidopsis_thaliana_total >= 2* (HOG_level_list$hypothesis_1_cons$Brassica_napus_total/2) &
                                    HOG_level_list$hypothesis_1_cons$Arabidopsis_lyrata_total >= 2 & 
                                    HOG_level_list$hypothesis_1_cons$Eutrema_salsugineum_total >= 2 &
                                    HOG_level_list$hypothesis_1_cons$Arabidopsis_thaliana_total >= 2,]
length(HOGs_exp_3vsBnapus$HOG)
# 449
write.table(HOGs_exp_3vsBnapus, paste0(OUTDIR, "/HOGs_exp_3vsBnapus.tsv"), sep ="\t", row.names = F, col.names = T)

HOGs_exp_2ArabidopsisvsBnapusandEutrema <- HOG_level_list$hypothesis_1_cons[HOG_level_list$hypothesis_1_cons$Arabidopsis_lyrata_total>= 2* (HOG_level_list$hypothesis_1_cons$Brassica_napus_total/2) &
                                                                            HOG_level_list$hypothesis_1_cons$Arabidopsis_lyrata_total>= 2* HOG_level_list$hypothesis_1_cons$Eutrema_salsugineum_total &
                                                                              HOG_level_list$hypothesis_1_cons$Arabidopsis_thaliana_total >= 2* (HOG_level_list$hypothesis_1_cons$Brassica_napus_total/2) &
                                                                              HOG_level_list$hypothesis_1_cons$Arabidopsis_thaliana_total >= 2* HOG_level_list$hypothesis_1_cons$Eutrema_salsugineum_total &
                                                                              HOG_level_list$hypothesis_1_cons$Arabidopsis_lyrata_total>= 2 &
                                                                              HOG_level_list$hypothesis_1_cons$Arabidopsis_thaliana_total >= 2,]
length(HOGs_exp_2ArabidopsisvsBnapusandEutrema$HOG)
# 441
write.table(HOGs_exp_2ArabidopsisvsBnapusandEutrema, paste0(OUTDIR, "/HOGs_exp_2ArabidopsisvsBnapusandEutrema.tsv"), sep ="\t", row.names = F, col.names = T)

HOGs_Bnapusexp_vsall3 <- HOG_level_list$hypothesis_1_cons[HOG_level_list$hypothesis_1_cons$Brassica_napus_total/2 >= 2* (HOG_level_list$hypothesis_1_cons$Arabidopsis_lyrata_total) &
                                                         HOG_level_list$hypothesis_1_cons$Brassica_napus_total/2 >= 2* (HOG_level_list$hypothesis_1_cons$Eutrema_salsugineum_total) &
                                                         HOG_level_list$hypothesis_1_cons$Brassica_napus_total/2 >= 2* (HOG_level_list$hypothesis_1_cons$Arabidopsis_thaliana_total),]
length(HOGs_Bnapusexp_vsall3$HOG)
# 1762
write.table(HOGs_Bnapusexp_vsall3, paste0(OUTDIR, "/HOGs_Bnapusexp_vsall3.tsv"), sep ="\t", row.names = F, col.names = T)

HOGS_Esa_Bnaexp_vs_2_Arabidopsis <-  HOG_level_list$hypothesis_1_cons[HOG_level_list$hypothesis_1_cons$Brassica_napus_total/2 >= 2* (HOG_level_list$hypothesis_1_cons$Arabidopsis_lyrata_total) &
                                                                        HOG_level_list$hypothesis_1_cons$Brassica_napus_total/2 >= 2* (HOG_level_list$hypothesis_1_cons$Arabidopsis_thaliana_total) &
                                                                        HOG_level_list$hypothesis_1_cons$Eutrema_salsugineum_total >= 2* (HOG_level_list$hypothesis_1_cons$Arabidopsis_lyrata_total) &
                                                                        HOG_level_list$hypothesis_1_cons$Eutrema_salsugineum_total >= 2* (HOG_level_list$hypothesis_1_cons$Arabidopsis_thaliana_total),]
length(HOGS_Esa_Bnaexp_vs_2_Arabidopsis$HOG)
# 103
write.table(HOGS_Esa_Bnaexp_vs_2_Arabidopsis, paste0(OUTDIR, "/HOGS_Esa_Bnaexp_vs_2_Arabidopsis.tsv"), sep ="\t", row.names = F, col.names = T)

#### Plot the number of DEGs per species in the Conserved data set
Conserved <- HOG_level_list$hypothesis_1[HOG_level_list$hypothesis_1$Arabidopsis_lyrata_total>=1 & HOG_level_list$hypothesis_1$Arabidopsis_thaliana_total>=1 & HOG_level_list$hypothesis_1$Eutrema_salsugineum_total>=1 & HOG_level_list$hypothesis_1$Brassica_napus_total>=1,]
Conserved_with_expression <- merge((Conserved[,-c(6)]), HOG_DE.a2tea, by="HOG")

Sig_DEGs <- Conserved_with_expression[Conserved_with_expression$significant=="yes", ]

## Volcano plot of genes within conserved data set per species

pdf(paste0(OUTDIR, "/Volcano_plot_per_species_in_Conserved_set.pdf"), 6, 4)
ggplot(data=Conserved_with_expression[!is.na(Conserved_with_expression$pvalue),], aes(x=log2FoldChange, y=-log10(pvalue)))+
  geom_point()+
  xlim(-11, 11)+
  geom_hline(yintercept=-log10(0.05), col="red") +
  facet_wrap(~ species)
dev.off()

## Volcano plot of genes within conserved data set individually per species
pdf(paste0(OUTDIR, "/Volcano_plot_Eutrema_in_Conserved_set.pdf"), 6, 4)
ggplot(data=Conserved_with_expression[!is.na(Conserved_with_expression$pvalue) & Conserved_with_expression$species=="Eutrema_salsugineum",], aes(x=log2FoldChange, y=-log10(pvalue)))+
  geom_point()+
  xlim(-11, 11)+
  geom_hline(yintercept=-log10(0.05), col="red")
dev.off()

pdf(paste0(OUTDIR, "/Volcano_plot_Alyrata_in_Conserved_set.pdf"), 6, 4)
ggplot(data=Conserved_with_expression[!is.na(Conserved_with_expression$pvalue) & Conserved_with_expression$species=="Arabidopsis_lyrata",], aes(x=log2FoldChange, y=-log10(pvalue)))+
  geom_point()+
  xlim(-11, 11)+
  geom_hline(yintercept=-log10(0.05), col="red")
dev.off()

pdf(paste0(OUTDIR, "/Volcano_plot_Bnapus_in_Conserved_set.pdf"), 6, 4)
ggplot(data=Conserved_with_expression[!is.na(Conserved_with_expression$pvalue) & Conserved_with_expression$species=="Brassica_napus",], aes(x=log2FoldChange, y=-log10(pvalue)))+
  geom_point()+
  xlim(-11, 11)+
  geom_hline(yintercept=-log10(0.05), col="red")
dev.off()

pdf(paste0(OUTDIR, "/Volcano_plot_Athaliana_in_Conserved_set.pdf"), 6, 4)
ggplot(data=Conserved_with_expression[!is.na(Conserved_with_expression$pvalue) & Conserved_with_expression$species=="Arabidopsis_thaliana",], aes(x=log2FoldChange, y=-log10(pvalue)))+
  geom_point()+
  xlim(-11, 11)+
  geom_hline(yintercept=-log10(0.05), col="red")
dev.off()

## Volcano plot per species of genes within Total data set

Total_with_expression <- merge((HOG_level_list$hypothesis_1[,-c(6)]), HOG_DE.a2tea, by="HOG")

pdf(paste0(OUTDIR, "/Volcano_plot_per_species_in_Total_set.pdf"), 6, 4)
ggplot(data=Total_with_expression[!is.na(Total_with_expression$pvalue),], aes(x=log2FoldChange, y=-log10(pvalue)))+
  geom_point()+
  xlim(-11, 11)+
  geom_hline(yintercept=-log10(0.05), col="red") +
  facet_wrap(~ species)
dev.off()

### number of sig DEGs per species in Conserved
Eutrema_sig_DEGs <- length(Conserved_with_expression[Conserved_with_expression$significant=="yes" & Conserved_with_expression$species=="Eutrema_salsugineum", "HOG"])
Eutrema_sig_DEGs # 7790
Up <- length(Conserved_with_expression[Conserved_with_expression$significant=="yes" & Conserved_with_expression$species=="Eutrema_salsugineum" & Conserved_with_expression$log2FoldChange>0, "HOG"])
Up # 3674
Down <- length(Conserved_with_expression[Conserved_with_expression$significant=="yes" & Conserved_with_expression$species=="Eutrema_salsugineum" & Conserved_with_expression$log2FoldChange<0, "HOG"])
Down # 4116

Alyrata_sig_DEGs <- length(Conserved_with_expression[Conserved_with_expression$significant=="yes" & Conserved_with_expression$species=="Arabidopsis_lyrata", "HOG"])
Alyrata_sig_DEGs # 9478
Up <- length(Conserved_with_expression[Conserved_with_expression$significant=="yes" & Conserved_with_expression$species=="Arabidopsis_lyrata" & Conserved_with_expression$log2FoldChange>0, "HOG"])
Up # 4847
Down <- length(Conserved_with_expression[Conserved_with_expression$significant=="yes" & Conserved_with_expression$species=="Arabidopsis_lyrata" & Conserved_with_expression$log2FoldChange<0, "HOG"])
Down # 4631

Bnapus_sig_DEGs <- length(Conserved_with_expression[Conserved_with_expression$significant=="yes" & Conserved_with_expression$species=="Brassica_napus", "HOG"])
Bnapus_sig_DEGs # 10065
Up <- length(Conserved_with_expression[Conserved_with_expression$significant=="yes" & Conserved_with_expression$species=="Brassica_napus" & Conserved_with_expression$log2FoldChange>0, "HOG"])
Up # 4887
Down <- length(Conserved_with_expression[Conserved_with_expression$significant=="yes" & Conserved_with_expression$species=="Brassica_napus" & Conserved_with_expression$log2FoldChange<0, "HOG"])
Down # 5178

Athaliana_sig_DEGs <- length(Conserved_with_expression[Conserved_with_expression$significant=="yes" & Conserved_with_expression$species=="Arabidopsis_thaliana", "HOG"])
Athaliana_sig_DEGs # 3141
Up <- length(Conserved_with_expression[Conserved_with_expression$significant=="yes" & Conserved_with_expression$species=="Arabidopsis_thaliana" & Conserved_with_expression$log2FoldChange>0, "HOG"])
Up # 1185
Down <- length(Conserved_with_expression[Conserved_with_expression$significant=="yes" & Conserved_with_expression$species=="Arabidopsis_thaliana" & Conserved_with_expression$log2FoldChange<0, "HOG"])
Down # 1802


### Plot the Intersect of the HOGs with DEG for all 4 species in an upset plot
## HOGs with sig DEGs per species in Conserved

HOGs_Eutrema_sig_DEG <- unique(Conserved_with_expression[Conserved_with_expression$significant=="yes" & Conserved_with_expression$species=="Eutrema_salsugineum", "HOG"])
HOGs_Eutrema_sig_DEG # 7434
HOGs_Eutrema_sig_DEG_Up <- unique(Conserved_with_expression[Conserved_with_expression$significant=="yes" & Conserved_with_expression$species=="Eutrema_salsugineum" & Conserved_with_expression$log2FoldChange>0, "HOG"])
HOGs_Eutrema_sig_DEG_Up # 3562
HOGs_Eutrema_sig_DEG_Down <- unique(Conserved_with_expression[Conserved_with_expression$significant=="yes" & Conserved_with_expression$species=="Eutrema_salsugineum" & Conserved_with_expression$log2FoldChange<0, "HOG"])
HOGs_Eutrema_sig_DEG_Down # 3921

HOGs_Alyrata_sig_DEG <- unique(Conserved_with_expression[Conserved_with_expression$significant=="yes" & Conserved_with_expression$species=="Arabidopsis_lyrata", "HOG"])
HOGs_Alyrata_sig_DEG # 8866
HOGs_Alyrata_sig_DEG_Up <- unique(Conserved_with_expression[Conserved_with_expression$significant=="yes" & Conserved_with_expression$species=="Arabidopsis_lyrata" & Conserved_with_expression$log2FoldChange>0, "HOG"])
HOGs_Alyrata_sig_DEG_Up # 4582
HOGs_Alyrata_sig_DEG_Down <- unique(Conserved_with_expression[Conserved_with_expression$significant=="yes" & Conserved_with_expression$species=="Arabidopsis_lyrata" & Conserved_with_expression$log2FoldChange<0, "HOG"])
HOGs_Alyrata_sig_DEG_Down # 4360

HOGs_Athaliana_sig_DEG <- unique(Conserved_with_expression[Conserved_with_expression$significant=="yes" & Conserved_with_expression$species=="Arabidopsis_thaliana", "HOG"])
HOGs_Athaliana_sig_DEG # 2865
HOGs_Athaliana_sig_DEG_Up <- unique(Conserved_with_expression[Conserved_with_expression$significant=="yes" & Conserved_with_expression$species=="Arabidopsis_thaliana" & Conserved_with_expression$log2FoldChange>0, "HOG"])
HOGs_Athaliana_sig_DEG_Up # 1151
HOGs_Athaliana_sig_DEG_Down <- unique(Conserved_with_expression[Conserved_with_expression$significant=="yes" & Conserved_with_expression$species=="Arabidopsis_thaliana" & Conserved_with_expression$log2FoldChange<0, "HOG"])
HOGs_Athaliana_sig_DEG_Down # 1720

HOGs_Bnapus_sig_DEG <- unique(Conserved_with_expression[Conserved_with_expression$significant=="yes" & Conserved_with_expression$species=="Brassica_napus", "HOG"])
HOGs_Bnapus_sig_DEG # 6956
HOGs_Bnapus_sig_DEG_Up <- unique(Conserved_with_expression[Conserved_with_expression$significant=="yes" & Conserved_with_expression$species=="Brassica_napus" & Conserved_with_expression$log2FoldChange>0, "HOG"])
HOGs_Bnapus_sig_DEG_Up # 3541
HOGs_Bnapus_sig_DEG_Down <- unique(Conserved_with_expression[Conserved_with_expression$significant=="yes" & Conserved_with_expression$species=="Brassica_napus" & Conserved_with_expression$log2FoldChange<0, "HOG"])
HOGs_Bnapus_sig_DEG_Down # 3645

library(UpSetR)
listInput <- list(Eutrema_salsugineum=HOGs_Eutrema_sig_DEG,
                  Arabidopsis_lyrata=HOGs_Alyrata_sig_DEG,
                  Arabidopsis_thaliana=HOGs_Athaliana_sig_DEG,
                  Brassica_napus=HOGs_Bnapus_sig_DEG)
sets <- c("Eutrema_salsugineum", "Arabidopsis_lyrata", "Arabidopsis_thaliana", "Brassica_napus")
names(listInput) <- c("Eutrema salsugineum",
                      "Arabidopsis lyrata",
                      "Arabidopsis thaliana",
                      "Brassica napus")


pdf(paste0(OUTDIR, "/UpSetPlot_HOGs_with_DEG_per_species_in_Conserved.pdf"), 10, 6, onefile = FALSE)
upset(fromList(listInput), nsets=4, nintersects=NA,
      set_size.show = TRUE,
      set_size.scale_max = 11000,
      sets.x.label = "Number of HOGs",
      mainbar.y.label = "Size of Intersection",
      keep.order = T, order.by = "freq",
#      sets = sets, 
      number.angles = 0, 
      #show.numbers = T,
      point.size = 2, line.size = 1,
      text.scale = c(2,1.5,1.5,1.5,1.5,2),
      queries= list(list(query = intersects, params=c("Arabidopsis lyrata","Eutrema salsugineum"), color= palette_OkabeIto_black[5], active = T),
                    list(query = intersects, params=c("Arabidopsis lyrata","Eutrema salsugineum", "Arabidopsis thaliana"), color= palette_OkabeIto_black[5], active = T),
                    list(query = intersects, params=c("Arabidopsis lyrata","Eutrema salsugineum", "Brassica napus"), color= palette_OkabeIto_black[5], active = T),
                    list(query = intersects, params=c("Arabidopsis lyrata","Eutrema salsugineum", "Arabidopsis thaliana", "Brassica napus"), color= palette_OkabeIto_black[5], active = T)
                    )
)
dev.off()


### Do the same for HOGs with at least 1 upregulated gene per species

listInput <- list(Eutrema_salsugineum=HOGs_Eutrema_sig_DEG_Up,
                  Arabidopsis_lyrata=HOGs_Alyrata_sig_DEG_Up,
                  Arabidopsis_thaliana=HOGs_Athaliana_sig_DEG_Up,
                  Brassica_napus=HOGs_Bnapus_sig_DEG_Up)
sets <- c("Eutrema_salsugineum", "Arabidopsis_lyrata", "Arabidopsis_thaliana", "Brassica_napus")

pdf(paste0(OUTDIR, "/UpSetPlot_HOGs_with_upregulated_DEG_per_species_in_Conserved.pdf"), 9, 5)
upset(fromList(listInput), nsets=4, nintersects=NA, sets.x.label = "Number of HOGs with at least \n1 upregulated DEG from SPECIES", mainbar.y.label = "Intersection of HOGs",
      keep.order = T, order.by = "freq",
      sets =sets,
      number.angles = 0, 
      #show.numbers = T,
      point.size = 2, line.size = 1,
      text.scale = c(2,1.5,1,1.5,1.5,1.5)
)
dev.off()

### and for HOGs with at least 1 downregulated gene per species

listInput <- list(Eutrema_salsugineum=HOGs_Eutrema_sig_DEG_Down,
                  Arabidopsis_lyrata=HOGs_Alyrata_sig_DEG_Down,
                  Arabidopsis_thaliana=HOGs_Athaliana_sig_DEG_Down,
                  Brassica_napus=HOGs_Bnapus_sig_DEG_Down)
sets <- c("Eutrema_salsugineum", "Arabidopsis_lyrata", "Arabidopsis_thaliana", "Brassica_napus")

pdf(paste0(OUTDIR, "/UpSetPlot_HOGs_with_downregulated_DEG_per_species_in_Conserved.pdf"), 9, 5)
upset(fromList(listInput), nsets=4, nintersects=NA, sets.x.label = "Number of HOGs with at least \n1 upregulated DEG from SPECIES", mainbar.y.label = "Intersection of HOGs",
      keep.order = T, order.by = "freq",
      sets =sets,
      number.angles = 0, 
      #show.numbers = T,
      point.size = 2, line.size = 1,
      text.scale = c(2,1.5,1,1.5,1.5,1.5)
)
dev.off()


### Plot the same for the Hypotheses (expanded Sets of HOGs)
### Hypothesis 1
HOGs_all_tol_exp <- HOG_level_list$hypothesis_1_cons[HOG_level_list$hypothesis_1_cons$expansion == "yes",]$HOG

listInput <- list(Eutrema_salsugineum=intersect(HOGs_all_tol_exp, HOGs_Eutrema_sig_DEG),
                  Arabidopsis_lyrata=intersect(HOGs_all_tol_exp, HOGs_Alyrata_sig_DEG),
                  Arabidopsis_thaliana=intersect(HOGs_all_tol_exp, HOGs_Athaliana_sig_DEG),
                  Brassica_napus=intersect(HOGs_all_tol_exp, HOGs_Bnapus_sig_DEG))
sets <- c("Eutrema_salsugineum", "Arabidopsis_lyrata", "Arabidopsis_thaliana", "Brassica_napus")

names(listInput) <- c("Eutrema salsugineum",
                      "Arabidopsis lyrata",
                      "Arabidopsis thaliana",
                      "Brassica napus")

pdf(paste0(OUTDIR, "/UpSetPlot_HOGs_with_DEG_per_species_in_Hypothesis_1.pdf"), 10, 6, onefile = FALSE)
upset(fromList(listInput), nsets=4, nintersects=NA,
      set_size.show = TRUE,
      set_size.scale_max = 80,
      sets.x.label = "Number of HOGs",
      mainbar.y.label = "Size of Intersection",
      keep.order = T, order.by = "freq",
      #      sets = sets, 
      number.angles = 0, 
      #show.numbers = T,
      point.size = 2, line.size = 1,
      text.scale = c(2,1.5,1.5,1.5,1.5,2),
      queries= list(list(query = intersects, params=c("Arabidopsis lyrata","Eutrema salsugineum"), color= palette_OkabeIto_black[5], active = T),
                    list(query = intersects, params=c("Arabidopsis lyrata","Eutrema salsugineum", "Arabidopsis thaliana"), color= palette_OkabeIto_black[5], active = T),
                    list(query = intersects, params=c("Arabidopsis lyrata","Eutrema salsugineum", "Brassica napus"), color= palette_OkabeIto_black[5], active = T),
                    list(query = intersects, params=c("Arabidopsis lyrata","Eutrema salsugineum", "Arabidopsis thaliana", "Brassica napus"), color= palette_OkabeIto_black[5], active = T)
      )
)
dev.off()

### are the HOGs which are regulated only in both tolerant but not in the sensitive species significantly over represented 
# in the HOGs expanded in tolerant species 
# compared to the HOGs regulated only in both tolerant in the Conserved HOGs?

# subset: 28 out of 47
# bg: 1621 out of 4638
# (deduced from plots: subset = UpSetPlot_HOGs_with_DEG_per_species_in_Hypothesis_1.pdf and bg = UpSetPlot_HOGs_with_DEG_per_species_in_Conserved.pdf)

q <- 28
m <- 1621
n <- 4638-1621
k <- 47

p <- phyper((q-1), m, n, k, lower.tail=F)
p # 0.0004607957

### Hypothesis 4
HOGs_all_sens_exp <- HOG_level_list$hypothesis_4_cons[HOG_level_list$hypothesis_4_cons$expansion == "yes",]$HOG

listInput <- list(Eutrema_salsugineum=intersect(HOGs_all_sens_exp, HOGs_Eutrema_sig_DEG),
                  Arabidopsis_lyrata=intersect(HOGs_all_sens_exp, HOGs_Alyrata_sig_DEG),
                  Arabidopsis_thaliana=intersect(HOGs_all_sens_exp, HOGs_Athaliana_sig_DEG),
                  Brassica_napus=intersect(HOGs_all_sens_exp, HOGs_Bnapus_sig_DEG))
sets <- c("Eutrema_salsugineum", "Arabidopsis_lyrata", "Arabidopsis_thaliana", "Brassica_napus")

pdf(paste0(OUTDIR, "/UpSetPlot_HOGs_with_DEG_per_species_in_Hypothesis_4.pdf"), 9, 5)
upset(fromList(listInput), nsets=4, nintersects=NA, sets.x.label = "Number of HOGs with at least \n1 DEG from SPECIES", mainbar.y.label = "Intersection of HOGs",
      keep.order = T, order.by = "freq",
      sets =sets,
      number.angles = 0, 
      #show.numbers = T,
      point.size = 2, line.size = 1,
      text.scale = c(2,1.5,1,1.5,1.5,1.5)
)
dev.off()

### Hypothesis 2
HOGs_Eutrema_exp <- HOG_level_list$hypothesis_2_cons[HOG_level_list$hypothesis_2_cons$expansion == "yes",]$HOG

listInput <- list(Eutrema_salsugineum=intersect(HOGs_Eutrema_exp, HOGs_Eutrema_sig_DEG),
                  Arabidopsis_lyrata=intersect(HOGs_Eutrema_exp, HOGs_Alyrata_sig_DEG),
                  Arabidopsis_thaliana=intersect(HOGs_Eutrema_exp, HOGs_Athaliana_sig_DEG),
                  Brassica_napus=intersect(HOGs_Eutrema_exp, HOGs_Bnapus_sig_DEG))
sets <- c("Eutrema_salsugineum", "Arabidopsis_lyrata", "Arabidopsis_thaliana", "Brassica_napus")

pdf(paste0(OUTDIR, "/UpSetPlot_HOGs_with_DEG_per_species_in_Hypothesis_2.pdf"), 9, 5)
upset(fromList(listInput), nsets=4, nintersects=NA, sets.x.label = "Number of HOGs with at least \n1 DEG from SPECIES", mainbar.y.label = "Intersection of HOGs",
      keep.order = T, order.by = "freq",
      sets =sets,
      number.angles = 0, 
      #show.numbers = T,
      point.size = 2, line.size = 1,
      text.scale = c(2,1.5,1,1.5,1.5,1.5)
)
dev.off()

### Hypothesis 3
HOGs_Alyrata_exp <- HOG_level_list$hypothesis_3_cons[HOG_level_list$hypothesis_3_cons$expansion == "yes",]$HOG

listInput <- list(Eutrema_salsugineum=intersect(HOGs_Alyrata_exp, HOGs_Eutrema_sig_DEG),
                  Arabidopsis_lyrata=intersect(HOGs_Alyrata_exp, HOGs_Alyrata_sig_DEG),
                  Arabidopsis_thaliana=intersect(HOGs_Alyrata_exp, HOGs_Athaliana_sig_DEG),
                  Brassica_napus=intersect(HOGs_Alyrata_exp, HOGs_Bnapus_sig_DEG))
sets <- c("Eutrema_salsugineum", "Arabidopsis_lyrata", "Arabidopsis_thaliana", "Brassica_napus")

pdf(paste0(OUTDIR, "/UpSetPlot_HOGs_with_DEG_per_species_in_Hypothesis_3.pdf"), 9, 5)
upset(fromList(listInput), nsets=4, nintersects=NA, sets.x.label = "Number of HOGs with at least \n1 DEG from SPECIES", mainbar.y.label = "Intersection of HOGs",
      keep.order = T, order.by = "freq",
      sets =sets,
      number.angles = 0, 
      #show.numbers = T,
      point.size = 2, line.size = 1,
      text.scale = c(2,1.5,1,1.5,1.5,1.5)
)
dev.off()

### Hypothesis 5
HOGs_Athaliana_exp <- HOG_level_list$hypothesis_5_cons[HOG_level_list$hypothesis_5_cons$expansion == "yes",]$HOG

listInput <- list(Eutrema_salsugineum=intersect(HOGs_Athaliana_exp, HOGs_Eutrema_sig_DEG),
                  Arabidopsis_lyrata=intersect(HOGs_Athaliana_exp, HOGs_Alyrata_sig_DEG),
                  Arabidopsis_thaliana=intersect(HOGs_Athaliana_exp, HOGs_Athaliana_sig_DEG),
                  Brassica_napus=intersect(HOGs_Athaliana_exp, HOGs_Bnapus_sig_DEG))
sets <- c("Eutrema_salsugineum", "Arabidopsis_lyrata", "Arabidopsis_thaliana", "Brassica_napus")

pdf(paste0(OUTDIR, "/UpSetPlot_HOGs_with_DEG_per_species_in_Hypothesis_5.pdf"), 9, 5)
upset(fromList(listInput), nsets=4, nintersects=NA, sets.x.label = "Number of HOGs with at least \n1 DEG from SPECIES", mainbar.y.label = "Intersection of HOGs",
      keep.order = T, order.by = "freq",
      sets =sets,
      number.angles = 0, 
      #show.numbers = T,
      point.size = 2, line.size = 1,
      text.scale = c(2,1.5,1,1.5,1.5,1.5)
)
dev.off()


### Hypothesis 6
HOGs_Bnapus_exp <- HOG_level_list$hypothesis_6_cons[HOG_level_list$hypothesis_6_cons$expansion == "yes",]$HOG

listInput <- list(Eutrema_salsugineum=intersect(HOGs_Bnapus_exp, HOGs_Eutrema_sig_DEG),
                  Arabidopsis_lyrata=intersect(HOGs_Bnapus_exp, HOGs_Alyrata_sig_DEG),
                  Arabidopsis_thaliana=intersect(HOGs_Bnapus_exp, HOGs_Athaliana_sig_DEG),
                  Brassica_napus=intersect(HOGs_Bnapus_exp, HOGs_Bnapus_sig_DEG))
sets <- c("Eutrema_salsugineum", "Arabidopsis_lyrata", "Arabidopsis_thaliana", "Brassica_napus")

pdf(paste0(OUTDIR, "/UpSetPlot_HOGs_with_DEG_per_species_in_Hypothesis_6.pdf"), 9, 5)
upset(fromList(listInput), nsets=4, nintersects=NA, sets.x.label = "Number of HOGs with at least \n1 DEG from SPECIES", mainbar.y.label = "Intersection of HOGs",
      keep.order = T, order.by = "freq",
      sets =sets,
      number.angles = 0, 
      #show.numbers = T,
      point.size = 2, line.size = 1,
      text.scale = c(2,1.5,1,1.5,1.5,1.5)
)
dev.off()


#### Tests: Is DE more conserved or more species-specific?
# Test if HOGs with a DEG from exactly 1 species, from exactly 4 species or from exactly 0 species
# are more or less frequent than expected form the distribution of DEGs per each species

## From the numbers of HOGs with DEG per species: what is the probability to have a HOG with DEG from a species?
## Calculations:
# Number of HOGs with DEG (success in background)
# At = 2865, Al = 8866, Es = 7434, Bn = 6956
## probability for success = success in background/background = black
# At 2865/17488 = 0,164
# Al 8866/17488 = 0,507
# Es 7434/17488 = 0,425
# Bn 6956/17488 = 0,398
##probability for failure = 1 - probability for success = white
# At 1 - 0,164 = 0,836
# Al 1-0,507 = 0,493
# Es 1-0,425 = 0,575
# Bn 1-0,398 = 0,602

## Calculate the probability to get a HOG with DEG from exactly 1 species (there are 4 such possibilities, 
## the total probability is the sum of each of the possibility's probability):

#Possibility|Code(At, Al, Es, Bn) |Calculation|Probability|Ratio of OBSERVED HOGs with DE from exactly 1
#1 | black, white, white, white	0,164*0,493*0,575*0,602=	0,028	282/2865=0,098
#2 |	white, black, white, white	0,836*0,507*0,575*0,602=	0,147	2107/6688=0,315
#3 |	white, white, black, white	0,836*0,493*0,425*0,602=	0,105	1418/7434=0,191
#4 |	white, white, white, black	0,836*0,493*0,575*0,398=	0,094	1456/6956=0,210
# sum = 0.374

# OBSERVATION of exactly one
#2107 (Alyrata) + 1456 (Bnapus) + 1418 (Eutrema) + 282 (Athaliana) = 5,263
#(numbers deduced from UpSet Plot: Intersection of HOGs with DEG)

# Test if OBSERVED exactly one is larger or smaller than PROBABILITY of exactly one

# make a contingency table of the frequencies of HOGs meeting the criteria:
success_in_bg <- (0.028+0.147+0.105+0.094)*17488
# probability 0.374 * bg = expected
bg <- 17488
fail_in_bg <- (bg-success_in_bg)

success_in_sample <- (2107 + 1456 + 1418 + 282)
# in the real case, 5263 of 17488 HOGs have a DEG from exactly 1 species
sample <- 17488

# value derived from     yes   no
# observed              success_in_sample (sample-success_in_sample)
# expected              success_in_bg (sample-success_in_bg)

contingency_table_1 <- data.frame(c("observed", "expected"), c(success_in_sample, success_in_bg), c((sample-success_in_sample), (sample-success_in_bg)), row.names = T)
names(contingency_table_1) <- c("yes", "no")
#               yes       no
# observed 5263.000 12225.00
# expected 6540.512 10947.49


chisq.test(contingency_table_1)
# Pearson's Chi-squared test with Yates' continuity correction

# data:  contingency_table_1
# X-squared = 208.37, df = 1, p-value < 2.2e-16


# --> In the real case, 30,1 % of HOGs (5263/17488) have a DEG from exactly 1 species.
# This is sign. less than expected by chance (6540/17488) when considering the background distribution of 
# HOGs with DEG from each of the 4 species.


## Calculate the probability to get a HOG with DEG from 0 species (there is exactly one possibility):
  
# Possibility	Code (At, Al, Es, Bn)	Calculation	Probability
# 1 | white, white, white, white |0,836*0,493*0,575*0,602 =  0,143

# OBSERVATION of DEG from exactly 0 species: 17,488 - 13,531 = 3,957

# Test if OBSERVED exactly 0 is larger or smaller than PROBABILITY of exactly 0

success_in_bg <- 0.143*17488
# probability 0.143 * bg = expected
bg <- 17488
fail_in_bg <- (bg-success_in_bg)

success_in_sample <- 3957
# in the real case, 3957 of 17488 HOGs have a DEG from exactly 0 species
sample <- 17488

contingency_table_0 <- data.frame(c("observed", "expected"), c(success_in_sample, success_in_bg), c((sample-success_in_sample), (sample-success_in_bg)), row.names = T)
names(contingency_table_0) <- c("yes", "no")
#               yes       no
# observed 3957.000 13531.00
# expected 2500.784 14987.22

chisq.test(contingency_table_0)
#Pearson's Chi-squared test with Yates' continuity correction

# data:  contingency_table_0
# X-squared = 402.18, df = 1, p-value < 2.2e-16


## --> In the real case, 3957/17488 (22,6 %) of HOGs have no DEG.  
# This is sign. more than expected by chance 2500.784/17488 (0.143) when considering the background distribution of 
# HOGs with no DEG

## Calculate the probability to get a HOG with DEG from exactly 4 species (there is exactly one possibility):
# This is the reverse from "Calculate the probability to get a HOG with DEG from 0 species" → interchange "black" to "white"and vice versa in each of the possibilities:
  
# Possibility	Code (At, Al, Es, Bn)	Calculation	Probability
#1 | black, black, black, black	
# (0.164 * 0.507 * 0.425 * 0.398) = 0,014

# OBSERVATION of DEG from exactly 4 species: 846 (deduced from UpSet Plot: Intersection of HOGs with DEG)

# Test if OBSERVED exactly 4 is larger or smaller than PROBABILITY of exactly 4

success_in_bg <- 0.014*17488
# probability 0.014 * bg = expected (= 245)
bg <- 17488
fail_in_bg <- (bg-success_in_bg)

success_in_sample <- 846
# in the real case, 846 of 17488 HOGs have a DEG from exactly 4 species
sample <- 17488

contingency_table_4 <- data.frame(c("observed", "expected"), c(success_in_sample, success_in_bg), c((sample-success_in_sample), (sample-success_in_bg)), row.names = T)
names(contingency_table_4) <- c("yes", "no")
#               yes       no
# observed 846.000 16642.00
# expected 244.832 17243.17

chisq.test(contingency_table_4)
# Pearson's Chi-squared test with Yates' continuity correction

# data:  contingency_table_4
# X-squared = 340.84, df = 1, p-value < 2.2e-16

## --> In the real case, 846 (4.8 %) of 17488 HOGs have a DEG from all 4 species.
# this is sign. more than expected by chance (0.014) when considering the background distribution of 
# HOGs with DEG from 4 species


#### Tests for over representation of HOGs with DEG in expanded HOGs compared to the Conserved set

### Hypothesis 1 test
# 2.1)b)2) Number of OGs in "Conserved"
HOGs <- HOG_level_list$hypothesis_1_cons$HOG
# 2b) What is the frequency of OGs with at least 1 DEG in "Conserved" OGs?
HOGs_DEG <- HOG_level_list$hypothesis_1_cons[!is.na(HOG_level_list$hypothesis_1_cons$total_sigDE) & HOG_level_list$hypothesis_1_cons$total_sigDE>=1,]$HOG
####### length(HOGs_DEG) = 13531

# 2.1)d)1) What is the frequency of OGs with at least 1 DEG from any of the Hypothesis 1 expanded species in "Conserved" OGs?
HOGs_DEG_any_tol <- HOG_level_list$hypothesis_1_cons[HOG_level_list$hypothesis_1_cons$Arabidopsis_lyrata_sigDE>=1 & !is.na(HOG_level_list$hypothesis_1_cons$Arabidopsis_lyrata_sigDE)
                                                     |
                                                     HOG_level_list$hypothesis_1_cons$Eutrema_salsugineum_sigDE>=1 & !is.na(HOG_level_list$hypothesis_1_cons$Eutrema_salsugineum_sigDE)
                                                     ,]$HOG
# 2.1)e)1) What is the frequency of OGs with at least 1 DEG from both of the Hypothesis 1 expanded species in "Conserved" OGs?
HOGs_DEG_both_tol <- HOG_level_list$hypothesis_1_cons[HOG_level_list$hypothesis_1_cons$Arabidopsis_lyrata_sigDE>=1 & !is.na(HOG_level_list$hypothesis_1_cons$Arabidopsis_lyrata_sigDE)
                                                     &
                                                     HOG_level_list$hypothesis_1_cons$Eutrema_salsugineum_sigDE>=1 & !is.na(HOG_level_list$hypothesis_1_cons$Eutrema_salsugineum_sigDE)
                                                     ,]$HOG
# 2.1)c)2) Within "Conserved": Number of OGs with "Hypothesis 1 expanded==yes"
HOGs_all_tol_exp <- HOG_level_list$hypothesis_1_cons[HOG_level_list$hypothesis_1_cons$expansion == "yes",]$HOG
# 2c) Within "Conserved": What is the frequency of OGs with at least 1 DEG in "Hypothesis 1 expanded==yes" OGs?
HOGs_all_tol_exp_DEG_any <- HOG_level_list$hypothesis_1_cons[HOG_level_list$hypothesis_1_cons$expansion == "yes" & !is.na(HOG_level_list$hypothesis_1_cons$total_sigDE) & HOG_level_list$hypothesis_1_cons$total_sigDE>=1,]$HOG
# 2d) What is the frequency of OGs with at least 1 DEG from any of the expanded species in "Hypothesis 1 expanded==yes" OGs?
HOGs_all_tol_exp_DEG_any_tol <- HOG_level_list$hypothesis_1_cons[HOG_level_list$hypothesis_1_cons$expansion == "yes" & HOG_level_list$hypothesis_1_cons$Eutrema_salsugineum_sigDE>=1 & !is.na(HOG_level_list$hypothesis_1_cons$Eutrema_salsugineum_sigDE) 
                                                                 |
                                                                 HOG_level_list$hypothesis_1_cons$expansion == "yes" & HOG_level_list$hypothesis_1_cons$Arabidopsis_lyrata_sigDE>=1 & !is.na(HOG_level_list$hypothesis_1_cons$Arabidopsis_lyrata_sigDE)
                                                                 ,]$HOG
## export a matrix with these HOGs and Genes
write.table(HOG_level_list$hypothesis_1_cons[HOG_level_list$hypothesis_1_cons$HOG %in% HOGs_all_tol_exp_DEG_any_tol,-6], "Analysis/HOG_level_list_all_tol_exp_DEG_any_tol.tsv", row.names = F)
# 2e) What is the frequency of OGs with at least 1 DEG from both of the expanded species in "Hypothesis 1 expanded==yes" OGs?
HOGs_all_tol_exp_DEG_both_tol <- HOG_level_list$hypothesis_1_cons[HOG_level_list$hypothesis_1_cons$expansion == "yes" 
                                                                  & HOG_level_list$hypothesis_1_cons$Eutrema_salsugineum_sigDE>=1 & !is.na(HOG_level_list$hypothesis_1_cons$Eutrema_salsugineum_sigDE) 
                                                                  & HOG_level_list$hypothesis_1_cons$Arabidopsis_lyrata_sigDE>=1 & !is.na(HOG_level_list$hypothesis_1_cons$Arabidopsis_lyrata_sigDE)
                                                                  ,]$HOG
## export a matrix with these HOGs and Genes
write.table(HOG_level_list$hypothesis_1_cons[HOG_level_list$hypothesis_1_cons$HOG %in% HOGs_all_tol_exp_DEG_both_tol,-6], "Analysis/HOG_level_list_all_tol_exp_DEG_both_tol.tsv", row.names = F)

### Conserved Set numbers DEG:
# length(HOGs) = 17,488
# length(HOGs_DEG) = 13,531
# = 0.77 of HOGs have DEGs

## Select only HOGs with at least 2 DEGs! Threshold for "DEG" is padj<=0.1

Conserved_with_expression_at_least_2_DEG <- 
  Conserved_with_expression %>%
  group_by(HOG) %>%
  mutate(two_DEG=case_when(sum(na.omit(padj) <= 0.1)>=2  ~ "yes",
                           TRUE ~ "no"))
HOGs_2_DEG <- unique(Conserved_with_expression_at_least_2_DEG[ Conserved_with_expression_at_least_2_DEG$two_DEG=="yes",]$HOG)
# length(HOGs_2_DEG)
# 8800

# out of 13,531 HOGs with DEG, 8800 have at least 2 DEGs (0.65,
# which means that (13,531 - 8,800 = 4,731 HOGs) 0.36 of HOGs with DEG have exactly 1 DEG)

## This subset of HOGs with at least 2 DEGs was used in the script Analysis_expression_pattern_per_OG_only_at_least_2_sig.R


###Hypergeometric test for over representation of HOGs with DEG in HOG in "Conserved" Data Set
# Hypergeometric test (comparable to Fisher's exact test) for enrichment of exp.SPECIES OGs with DEGs over exp.SPECIES OGs in Conserved
# q = success in the sample
# m = success in population
# n = population - m
# k = sample

# Hypothesis 1: Tolerant vs. sensitive with DEG Any
# q = 2c)
q <- length(HOGs_all_tol_exp_DEG_any)
# m = 2b)
m <- length(HOGs_DEG)
# n = 2.1)b)2) - m
n <- (length(HOGs) - m)
# 2.1)c)2)
k <- length(HOGs_all_tol_exp)

p <- phyper((q-1), m, n, k, lower.tail=F)
p # 0.9599121

# Hypothesis 1: Tolerant vs. sensitive with DEG from at least 1 tol species
# q = 2d)
q <- length(HOGs_all_tol_exp_DEG_any_tol)
# m = 2.1)d)1)
m <- length(HOGs_DEG_any_tol)
# n = 2.1)b)2) - m
n <- (length(HOGs) - m)
# 2.1)c)2)
k <- length(HOGs_all_tol_exp)

p <- phyper((q-1), m, n, k, lower.tail=F)
p # 0.7362929

# Hypothesis 1: Tolerant vs. sensitive with DEG from both tol species
# 2e)
q <- length(HOGs_all_tol_exp_DEG_both_tol)
# m = 2.1)e)1)
m <- length(HOGs_DEG_both_tol)
# n = 2.1)b)2) - m
n <- (length(HOGs) - m)
# 2.1)c)2)
k <- length(HOGs_all_tol_exp)

p <- phyper((q-1), m, n, k, lower.tail=F)
p # 0.05981553


### Export a matrix with the HOGs and the Genes from Hypothesis_1 expanded OGs:
Hypothesis_1_Exp_matrix <- Conserved_dataset_matrix_ordered[Conserved_dataset_matrix_ordered$HOG %in% HOGs_all_tol_exp,]
write.table(Hypothesis_1_Exp_matrix, paste0(OUTDIR, "/Hypothesis_1_Exp_in_Conserved_dataset_matrix.tsv"), sep="\t", row.names=F, col.names=T, quote=F)

## extract the HOGs with their corresponding GO terms
HOGs_all_tol_exp_DEG_any_tol_with_GO <- SFA_OG_level$hypothesis_1[SFA_OG_level$hypothesis_1$HOG %in% HOGs_all_tol_exp_DEG_any_tol,]
HOGs_all_tol_exp_DEG_both_tol_with_GO <- SFA_OG_level$hypothesis_1[SFA_OG_level$hypothesis_1$HOG %in% HOGs_all_tol_exp_DEG_both_tol,]
# export the GO annotations of the subsets
write.table(HOGs_all_tol_exp_DEG_any_tol_with_GO, "./Analysis/GO_terms/HOGs_all_tol_exp_DEG_any_tol_with_GO.tsv", sep="\t", row.names = F, col.names = T)
write.table(HOGs_all_tol_exp_DEG_both_tol_with_GO, "./Analysis/GO_terms/HOGs_all_tol_exp_DEG_both_tol_with_GO.tsv", sep="\t", row.names = F, col.names = T)

# export a matrix with the HOGs and the Genes from the Hypothesis_1 expanded and DE HOGs with the genes function (human readable description (HRD))
## extract the HOGs with the HRD of each gene
HOGs_all_tol_exp_DEG_any_tol_with_HRD <- rbind(SFA$Arabidopsis_thaliana[SFA$Arabidopsis_thaliana$HOG %in% HOGs_all_tol_exp_DEG_any_tol,],
                                               SFA$Arabidopsis_lyrata[SFA$Arabidopsis_lyrata$HOG %in% HOGs_all_tol_exp_DEG_any_tol,],
                                               SFA$Eutrema_salsugineum[SFA$Eutrema_salsugineum$HOG %in% HOGs_all_tol_exp_DEG_any_tol,],
                                               SFA$Brassica_napus[SFA$Brassica_napus$HOG %in% HOGs_all_tol_exp_DEG_any_tol,])
HOGs_all_tol_exp_DEG_both_tol_with_HRD <- rbind(SFA$Arabidopsis_thaliana[SFA$Arabidopsis_thaliana$HOG %in% HOGs_all_tol_exp_DEG_both_tol,],
                                               SFA$Arabidopsis_lyrata[SFA$Arabidopsis_lyrata$HOG %in% HOGs_all_tol_exp_DEG_both_tol,],
                                               SFA$Eutrema_salsugineum[SFA$Eutrema_salsugineum$HOG %in% HOGs_all_tol_exp_DEG_both_tol,],
                                               SFA$Brassica_napus[SFA$Brassica_napus$HOG %in% HOGs_all_tol_exp_DEG_both_tol,])

# combine the table with the expression data
HOGs_all_tol_exp_DEG_any_tol_with_HRD_and_expression <- merge(HOGs_all_tol_exp_DEG_any_tol_with_HRD, Conserved_with_expression, by.x="Protein-Accession", by.y = "gene")
HOGs_all_tol_exp_DEG_any_tol_with_HRD_and_expression_publish <- HOGs_all_tol_exp_DEG_any_tol_with_HRD_and_expression[,c(1,2,5,22:24,27:29)]
HOGs_all_tol_exp_DEG_any_tol_with_HRD_and_expression_publish_ordered <- HOGs_all_tol_exp_DEG_any_tol_with_HRD_and_expression_publish[order(HOGs_all_tol_exp_DEG_any_tol_with_HRD_and_expression_publish$HOG.x, HOGs_all_tol_exp_DEG_any_tol_with_HRD_and_expression_publish$species),]

# export a table with all the information to publish
write.table(HOGs_all_tol_exp_DEG_any_tol_with_HRD_and_expression_publish_ordered, paste0(OUTDIR, "/Hypothesis_1_DEG_any_tol_with_HRD_and_expression.tsv"), sep="\t", row.names=F, col.names=T, quote=F)

# combine the table with the expression data
HOGs_all_tol_exp_DEG_both_tol_with_HRD_and_expression <- merge(HOGs_all_tol_exp_DEG_both_tol_with_HRD, Conserved_with_expression, by.x="Protein-Accession", by.y = "gene")
HOGs_all_tol_exp_DEG_both_tol_with_HRD_and_expression_publish <- HOGs_all_tol_exp_DEG_both_tol_with_HRD_and_expression[,c(1,2,5,22:24,27:29)]
HOGs_all_tol_exp_DEG_both_tol_with_HRD_and_expression_publish_ordered <- HOGs_all_tol_exp_DEG_both_tol_with_HRD_and_expression_publish[order(HOGs_all_tol_exp_DEG_both_tol_with_HRD_and_expression_publish$HOG.x, HOGs_all_tol_exp_DEG_both_tol_with_HRD_and_expression_publish$species),]

# export a table with all the information to publish
write.table(HOGs_all_tol_exp_DEG_both_tol_with_HRD_and_expression_publish_ordered, paste0(OUTDIR, "/Hypothesis_1_DEG_both_tol_with_HRD_and_expression.tsv"), sep="\t", row.names=F, col.names=T, quote=F)

#### test if the conservation of DE in both tolerant species (in contrast to DE in ANY of the both tolerant)
# is significantly enriched in the commonly expanded gene families compared to the background
# (Pie chart 2 vs. pie chart 1 which show the numbers of HOGs with DEG from both or from any tolerant in the expanded and in the conserved set)

q <- length(HOGs_all_tol_exp_DEG_both_tol)
# m = 2.1)e)1)
m <- length(HOGs_DEG_both_tol)
# n = 2.1)b)2) - m
n <- (length(HOGs_DEG_any_tol) - m)
# 2.1)c)2)
k <- length(HOGs_all_tol_exp_DEG_any_tol)

p <- phyper((q-1), m, n, k, lower.tail=F)
p # 0.02233873
## --> yes, the enrichment is significant! 


### How many of these HOGs are only UPregulated in both tolerant species and NOT regulated in the sensitive species?
Conserved_HOGs <- as.character(HOG_level_list$hypothesis_1_cons$HOG)
Conserved_with_expression <- HOG_DE.a2tea[HOG_DE.a2tea$HOG %in% Conserved_HOGs,]

Conserved_with_expression_with_species_specific_category <-
  Conserved_with_expression[!is.na(Conserved_with_expression$log2FoldChange) & !is.na(Conserved_with_expression$padj) & Conserved_with_expression$padj<=0.1,] %>%
  group_by(HOG, species) %>%
  mutate(category_of_regulation_per_species=case_when(all(log2FoldChange < 0) ~ "down" ,
                                                      all(log2FoldChange > 0) ~ "up",
                                                      any(log2FoldChange < 0) && any(log2FoldChange > 0) ~ "down_and_up"))

# species specific:
Conserved_with_expression_with_species_specific_category$category_of_regulation_per_species_with_name <-
  paste(Conserved_with_expression_with_species_specific_category$species, Conserved_with_expression_with_species_specific_category$category_of_regulation_per_species, sep=" ")

# now select only those HOGs which have at least 1 $category_of_regulation_per_species_with_name == "Arabidopsis_lyrata up" 
# and at least 1 $category_of_regulation_per_species_with_name=="Eutrema_salsugineum up"
# and do NOT contain $species=="Arabidopsis_thaliana" | $species=="Brassica_napus"

Conserved_with_expression_with_species_specific_category_only_both_tol_up <-
  Conserved_with_expression_with_species_specific_category %>%
  group_by(HOG) %>%
  filter(any(category_of_regulation_per_species_with_name == "Arabidopsis_lyrata up") & any(category_of_regulation_per_species_with_name=="Eutrema_salsugineum up")
         & !("Arabidopsis_thaliana" %in% species)
         &  !("Brassica_napus" %in% species))

length(unique(Conserved_with_expression_with_species_specific_category_only_both_tol_up$HOG))
# 777

# write the matrix of the sig. genes of the 777 HOGs
write.table(Conserved_with_expression_with_species_specific_category_only_both_tol_up, "/home/cropbio/sciebo/Documents/PhD/Brassicaceae-drought/BR-dr-03/Analysis/Conserved_with_expression_with_species_specific_category_only_both_tol_up.tsv", row.names = F, col.names = T, sep= "\t")

## extract the GO terms
Conserved_with_expression_with_species_specific_category_only_both_tol_up_with_GO <- SFA_OG_level$hypothesis_1[SFA_OG_level$hypothesis_1$HOG %in% Conserved_with_expression_with_species_specific_category_only_both_tol_up$HOG,]
# export the GO terms
write.table(Conserved_with_expression_with_species_specific_category_only_both_tol_up_with_GO, "/home/cropbio/sciebo/Documents/PhD/Brassicaceae-drought/BR-dr-03/Analysis/GO_terms/HOGs_only_both_tol_up_with_GO.tsv")


# select the expanded HOGs from here:
Conserved_with_expression_with_species_specific_category_only_both_tol_up_and_exp_both_tol <-
  Conserved_with_expression_with_species_specific_category_only_both_tol_up[
    Conserved_with_expression_with_species_specific_category_only_both_tol_up$HOG %in% HOGs_all_tol_exp,]

length(unique(Conserved_with_expression_with_species_specific_category_only_both_tol_up_and_exp_both_tol$HOG))
# 17 --> correct
# write the matrix of the sig. genes of these HOGs
write.table(Conserved_with_expression_with_species_specific_category_only_both_tol_up_and_exp_both_tol, "/home/cropbio/sciebo/Documents/PhD/Brassicaceae-drought/BR-dr-03/Analysis/Conserved_with_expression_with_species_specific_category_only_both_tol_up_and_exp_both_tol.tsv", row.names = F, col.names = T, sep= "\t")

## extract the GO terms
Conserved_with_expression_with_species_specific_category_only_both_tol_up_and_exp_both_tol_with_GO <- SFA_OG_level$hypothesis_1[SFA_OG_level$hypothesis_1$HOG %in% Conserved_with_expression_with_species_specific_category_only_both_tol_up_and_exp_both_tol$HOG,]
# export the GO terms
write.table(Conserved_with_expression_with_species_specific_category_only_both_tol_up_and_exp_both_tol_with_GO, "/home/cropbio/sciebo/Documents/PhD/Brassicaceae-drought/BR-dr-03/Analysis/GO_terms/HOGs_both_tol_exp_only_both_tol_up_with_GO.tsv")

### are the only in tolerant up regulated HOGs significantly over represented in the in tolerant species expanded HOGs which are only up, 
# compared to the only in tolerant up regulated HOGs in the Conserved HOGs which are only up regulated?

# 24 and 2768 only up regulated HOGs in the in both tolerant expanded and DE and in the Conserved Set were deduced from the script Analysis_expression_pattern_per_OG_only_at_least_2_sig.R
# 17/24 = 70.83 %
# 777/1472= 0.5279
q <- 17
m <- 777
n <- 1472-777
k <- 24

p <- phyper((q-1), m, n, k, lower.tail=F)
p # 0.05552217


### Hypothesis 4 test
# 2.1)b)2) Number of OGs in "Conserved"
HOGs <- HOG_level_list$hypothesis_1_cons$HOG
# 2b) What is the frequency of OGs with at least 1 DEG in "Conserved" OGs?
HOGs_DEG <- HOG_level_list$hypothesis_1_cons[!is.na(HOG_level_list$hypothesis_1_cons$total_sigDE) & HOG_level_list$hypothesis_1_cons$total_sigDE>=1,]$HOG
# 2.1)g)1) What is the frequency of OGs with at least 1 DEG from any of the Hypothesis 4 expanded species in "Conserved" OGs?
HOGs_DEG_any_sens <- HOG_level_list$hypothesis_4_cons[HOG_level_list$hypothesis_4_cons$Arabidopsis_thaliana_sigDE>=1 & !is.na(HOG_level_list$hypothesis_4_cons$Arabidopsis_thaliana_sigDE)
                                                     |
                                                     HOG_level_list$hypothesis_4_cons$Brassica_napus_sigDE>=1 & !is.na(HOG_level_list$hypothesis_4_cons$Brassica_napus_sigDE)
                                                     ,]$HOG
# 2.1)h)1) What is the frequency of OGs with at least 1 DEG from both of the Hypothesis 4 expanded species in "Conserved" OGs?
HOGs_DEG_both_sens <- HOG_level_list$hypothesis_4_cons[HOG_level_list$hypothesis_4_cons$Arabidopsis_thaliana_sigDE>=1 & !is.na(HOG_level_list$hypothesis_4_cons$Arabidopsis_thaliana_sigDE)
                                                      &
                                                      HOG_level_list$hypothesis_4_cons$Brassica_napus_sigDE>=1 & !is.na(HOG_level_list$hypothesis_4_cons$Brassica_napus_sigDE)
                                                      ,]$HOG
# 2.1)f)2) Within "Conserved": Number of OGs with "Hypothesis 4 expanded==yes"
HOGs_all_sens_exp <- HOG_level_list$hypothesis_4_cons[HOG_level_list$hypothesis_4_cons$expansion == "yes",]$HOG
# 2f) Within "Conserved": What is the frequency of OGs with at least 1 DEG in "Hypothesis 4 expanded==yes" OGs?
HOGs_all_sens_exp_DEG_any <- HOG_level_list$hypothesis_4_cons[HOG_level_list$hypothesis_4_cons$expansion == "yes" & !is.na(HOG_level_list$hypothesis_4_cons$total_sigDE) & HOG_level_list$hypothesis_4_cons$total_sigDE>=1,]$HOG
# 2g) What is the frequency of OGs with at least 1 DEG from any of the expanded species in "Hypothesis 4 expanded==yes" OGs?
HOGs_all_sens_exp_DEG_any_sens <- HOG_level_list$hypothesis_4_cons[HOG_level_list$hypothesis_4_cons$expansion == "yes" & HOG_level_list$hypothesis_4_cons$Arabidopsis_thaliana_sigDE>=1 & !is.na(HOG_level_list$hypothesis_4_cons$Arabidopsis_thaliana_sigDE) 
                                                                 |
                                                                 HOG_level_list$hypothesis_4_cons$expansion == "yes" & HOG_level_list$hypothesis_4_cons$Brassica_napus_sigDE>=1 & !is.na(HOG_level_list$hypothesis_4_cons$Brassica_napus_sigDE)
                                                                 ,]$HOG
# 2h) What is the frequency of OGs with at least 1 DEG from both of the expanded species in "Hypothesis 4 expanded==yes" OGs?
HOGs_all_sens_exp_DEG_both_sens <- HOG_level_list$hypothesis_4_cons[HOG_level_list$hypothesis_4_cons$expansion == "yes" 
                                                                  & HOG_level_list$hypothesis_4_cons$Arabidopsis_thaliana_sigDE>=1 & !is.na(HOG_level_list$hypothesis_4_cons$Arabidopsis_thaliana_sigDE) 
                                                                  & HOG_level_list$hypothesis_4_cons$Brassica_napus_sigDE>=1 & !is.na(HOG_level_list$hypothesis_4_cons$Brassica_napus_sigDE)
                                                                  ,]$HOG

###Hypergeometric test for over representation of HOGs with DEG in HOG in "Conserved" Data Set
# Hypergeometric test (comparable to Fisher's exact test) for enrichment of exp.SPECIES OGs with DEGs over exp.SPECIES OGs in Conserved
# q = OGs with diff exp genes in Hypothesis OGs (= success in the sample)
# m = OGs with diff exp genes in Conserved OGs (= success in population)
# n = Conserved OGs - m
# k = OGs in Hypothesis OGs (= sample)

# Hypothesis 4: Sensitive vs. tolerant with DEG Any
# q = 2f)
q <- length(HOGs_all_sens_exp_DEG_any)
# m = 2b)
m <- length(HOGs_DEG)
# n = 2.1)b)2) - m
n <- (length(HOGs) - m)
# 2.1)f)2)
k <- length(HOGs_all_sens_exp)

p <- phyper((q-1), m, n, k, lower.tail=F)
p # 0.3127017

# Hypothesis 4: Sensitive vs. tolerant with DEG from at least 1 sens species
# q = 2g)
q <- length(HOGs_all_sens_exp_DEG_any_sens)
# m= 2.1)g)1)
m <- length(HOGs_DEG_any_sens)
# n = 2.1)b)2) - m
n <- (length(HOGs) - m)
# 2.1)f)2)
k <- length(HOGs_all_sens_exp)

p <- phyper((q-1), m, n, k, lower.tail=F)
p # 0.002323285

# Hypothesis 4: Sensitive vs. tolerant with DEG from both sens species
# q = 2h)
q <- length(HOGs_all_sens_exp_DEG_both_sens)
# 2.1)h)1)
m <- length(HOGs_DEG_both_sens)
# n = 2.1)b)2) - m
n <- (length(HOGs) - m)
# 2.1)f)2)
k <- length(HOGs_all_sens_exp)

p <- phyper((q-1), m, n, k, lower.tail=F)
p # 0.340438


### Export a matrix with the HOGs and the Genes from Hypothesis_4 expanded OGs:
Hypothesis_4_Exp_matrix <- Conserved_dataset_matrix_ordered[Conserved_dataset_matrix_ordered$HOG %in% HOGs_all_sens_exp,]
write.table(Hypothesis_4_Exp_matrix, paste0(OUTDIR, "/Hypothesis_4_Exp_in_Conserved_dataset_matrix.tsv"), sep="\t", row.names=F, col.names=T, quote=F)


#### test if the conservation of DE in both sensitive species (in contrast to DE in ANY of the both sensitive)
# is significantly enriched in the commonly expanded gene families compared to the background
# (Pie chart 6 vs. pie chart 5 which show the numbers of HOGs with DEG from both or from any sensitive in the expanded and in the conserved set)

q <- length(HOGs_all_sens_exp_DEG_both_sens)
# m = 2.1)e)1)
m <- length(HOGs_DEG_both_sens)
# n = 2.1)b)2) - m
n <- (length(HOGs_DEG_any_sens) - m)
# 2.1)c)2)
k <- length(HOGs_all_sens_exp_DEG_any_sens)

p <- phyper((q-1), m, n, k, lower.tail=F)
p # 0.679


### Hypothesis 2 test
# 2.1)b)2) Number of OGs in "Conserved"
HOGs <- HOG_level_list$hypothesis_1_cons$HOG
# 2b) What is the frequency of OGs with at least 1 DEG in "Conserved" OGs?
HOGs_DEG <- HOG_level_list$hypothesis_1_cons[!is.na(HOG_level_list$hypothesis_1_cons$total_sigDE) & HOG_level_list$hypothesis_1_cons$total_sigDE>=1,]$HOG
# 2.1)j)1) What is the frequency of OGs with at least 1 DEG from the Hypothesis 2 expanded species in "Conserved" OGs?
HOGs_DEG_Eutrema <- HOG_level_list$hypothesis_2_cons[HOG_level_list$hypothesis_2_cons$Eutrema_salsugineum_sigDE>=1 & !is.na(HOG_level_list$hypothesis_2_cons$Eutrema_salsugineum_sigDE)
                                                     ,]$HOG
# 2.1)i)2) Within "Conserved": Number of OGs with "Hypothesis 2 expanded==yes"
HOGs_Eutrema_exp <- HOG_level_list$hypothesis_2_cons[HOG_level_list$hypothesis_2_cons$expansion == "yes",]$HOG
# 2i) Within "Conserved": What is the frequency of OGs with at least 1 DEG in "Hypothesis 2 expanded==yes" OGs?
cons_1_and_2_merged_2 <-  merge(HOG_level_list$hypothesis_1_cons[,c("HOG", "total_sigDE")], HOG_level_list$hypothesis_2_cons, by="HOG")
# select for total_sigDE.x because we are interested in total_sigDE from all 4 species (inferred from data frame HOG_level_list$hypothesis_1_cons, which contains all 4 species)
HOGs_Eutrema_exp_DEG_any <- cons_1_and_2_merged_2[cons_1_and_2_merged_2$expansion == "yes" & !is.na(cons_1_and_2_merged_2$total_sigDE.x) & cons_1_and_2_merged_2$total_sigDE.x>=1,]$HOG
# 2j) What is the frequency of OGs with at least 1 DEG from any of the expanded species in "Hypothesis 2 expanded==yes" OGs?
HOGs_Eutrema_exp_DEG_Eutrema <- HOG_level_list$hypothesis_2_cons[HOG_level_list$hypothesis_2_cons$expansion == "yes" & HOG_level_list$hypothesis_2_cons$Eutrema_salsugineum_sigDE>=1 & !is.na(HOG_level_list$hypothesis_2_cons$Eutrema_salsugineum_sigDE) 
                                                                 ,]$HOG
## export a matrix with these HOGs and Genes
write.table(HOG_level_list$hypothesis_1_cons[HOG_level_list$hypothesis_1_cons$HOG %in% HOGs_Eutrema_exp_DEG_Eutrema,-6], "Analysis/HOG_level_list_Eutrema_exp_DEG_Eutrema.tsv", row.names = F)

###Hypergeometric test for over representation of HOGs with DEG in HOG in "Conserved" Data Set
# Hypergeometric test (comparable to Fisher's exact test) for enrichment of exp.SPECIES OGs with DEGs over exp.SPECIES OGs in Conserved
# q = OGs with diff exp genes in Hypothesis OGs (= success in the sample)
# m = OGs with diff exp genes in Conserved OGs (= population)
# n = Conserved OGs - m
# k = OGs in Hypothesis OGs (= sample)

# Hypothesis 2: Eutrema vs. sensitive with DEG Any
# q = 2i)
q <- length(HOGs_Eutrema_exp_DEG_any)
# m = 2b)
m <- length(HOGs_DEG)
# n = 2.1)b)2) - m
n <- (length(HOGs) - m)
# 2.1)i)2)
k <- length(HOGs_Eutrema_exp)

p <- phyper((q-1), m, n, k, lower.tail=F)
p # 0.9920274

# Hypothesis 2: Eutrema vs. sensitive with DEG Eutrema
# q = 2j)
q <- length(HOGs_Eutrema_exp_DEG_Eutrema)
# m= 2.1)j)1)
m <- length(HOGs_DEG_Eutrema)
# n = 2.1)b)2) - m
n <- (length(HOGs) - m)
# 2.1)i)2)
k <- length(HOGs_Eutrema_exp)

p <- phyper((q-1), m, n, k, lower.tail=F)
p # 3.165845e-05

### Export a matrix with the HOGs and the Genes from Hypothesis_2 expanded OGs:
Hypothesis_2_Exp_matrix <- Conserved_dataset_matrix_ordered[Conserved_dataset_matrix_ordered$HOG %in% HOGs_Eutrema_exp,]
write.table(Hypothesis_2_Exp_matrix, paste0(OUTDIR, "/Hypothesis_2_Exp_in_Conserved_dataset_matrix.tsv"), sep="\t", row.names=F, col.names=T, quote=F)

# export a matrix with the HOGs and the Genes from the Hypothesis_2 expanded and DE HOGs with the genes function (human readable description (HRD))
## extract the HOGs with the HRD of each gene
HOGs_Eutrema_exp_DEG_Eutrema_with_HRD <- rbind(SFA$Arabidopsis_thaliana[SFA$Arabidopsis_thaliana$HOG %in% HOGs_Eutrema_exp_DEG_Eutrema,],
                                               SFA$Arabidopsis_lyrata[SFA$Arabidopsis_lyrata$HOG %in% HOGs_Eutrema_exp_DEG_Eutrema,],
                                               SFA$Eutrema_salsugineum[SFA$Eutrema_salsugineum$HOG %in% HOGs_Eutrema_exp_DEG_Eutrema,],
                                               SFA$Brassica_napus[SFA$Brassica_napus$HOG %in% HOGs_Eutrema_exp_DEG_Eutrema,])

# combine the table with the expression data
HOGs_Eutrema_exp_DEG_Eutrema_with_HRD_and_expression <- merge(HOGs_Eutrema_exp_DEG_Eutrema_with_HRD, Conserved_with_expression, by.x="Protein-Accession", by.y = "gene")
HOGs_Eutrema_exp_DEG_Eutrema_with_HRD_and_expression_publish <- HOGs_Eutrema_exp_DEG_Eutrema_with_HRD_and_expression[,c(1,2,5,7:9,12:14)]
HOGs_Eutrema_exp_DEG_Eutrema_with_HRD_and_expression_publish_ordered <- HOGs_Eutrema_exp_DEG_Eutrema_with_HRD_and_expression_publish[order(HOGs_Eutrema_exp_DEG_Eutrema_with_HRD_and_expression_publish$HOG.x, HOGs_Eutrema_exp_DEG_Eutrema_with_HRD_and_expression_publish$species),]

# export a table with all the information to publish
write.table(HOGs_Eutrema_exp_DEG_Eutrema_with_HRD_and_expression_publish_ordered, paste0(OUTDIR, "/HOGs_Eutrema_exp_DEG_Eutrema_with_HRD_and_expression.tsv"), sep="\t", row.names=F, col.names=T, quote=F)


### Hypothesis 3 test
# 2.1)b)2) Number of OGs in "Conserved"
HOGs <- HOG_level_list$hypothesis_1_cons$HOG
# 2b) What is the frequency of OGs with at least 1 DEG in "Conserved" OGs?
HOGs_DEG <- HOG_level_list$hypothesis_1_cons[!is.na(HOG_level_list$hypothesis_1_cons$total_sigDE) & HOG_level_list$hypothesis_1_cons$total_sigDE>=1,]$HOG
# 2.1)l)1) What is the frequency of OGs with at least 1 DEG from the Hypothesis 3 expanded species in "Conserved" OGs?
HOGs_DEG_Alyrata <- HOG_level_list$hypothesis_3_cons[HOG_level_list$hypothesis_3_cons$Arabidopsis_lyrata_sigDE>=1 & !is.na(HOG_level_list$hypothesis_3_cons$Arabidopsis_lyrata_sigDE)
                                                     ,]$HOG
# 2.1)k)2) Within "Conserved": Number of OGs with "Hypothesis 3 expanded==yes"
HOGs_Alyrata_exp <- HOG_level_list$hypothesis_3_cons[HOG_level_list$hypothesis_3_cons$expansion == "yes",]$HOG
# 2k) Within "Conserved": What is the frequency of OGs with at least 1 DEG in "Hypothesis 3 expanded==yes" OGs?
cons_1_and_3_merged_2 <-  merge(HOG_level_list$hypothesis_1_cons[,c("HOG",  "total_sigDE")], HOG_level_list$hypothesis_3_cons, by="HOG")
# select for total_sigDE.x because we are interested in total_sigDE from all 4 species (inferred from data frame HOG_level_list$hypothesis_1_cons, which contains all 4 species)
HOGs_Alyrata_exp_DEG_any <- cons_1_and_3_merged_2[cons_1_and_3_merged_2$expansion == "yes" & !is.na(cons_1_and_3_merged_2$total_sigDE.x) & cons_1_and_3_merged_2$total_sigDE.x>=1,]$HOG
# 2l) What is the frequency of OGs with at least 1 DEG from any of the expanded species in "Hypothesis 3 expanded==yes" OGs?
HOGs_Alyrata_exp_DEG_Alyrata <- HOG_level_list$hypothesis_3_cons[HOG_level_list$hypothesis_3_cons$expansion == "yes" & HOG_level_list$hypothesis_3_cons$Arabidopsis_lyrata_sigDE>=1 & !is.na(HOG_level_list$hypothesis_3_cons$Arabidopsis_lyrata_sigDE) 
                                                                 ,]$HOG
## export a matrix with these HOGs and Genes
write.table(HOG_level_list$hypothesis_1_cons[HOG_level_list$hypothesis_1_cons$HOG %in% HOGs_Alyrata_exp_DEG_Alyrata,-6], "Analysis/HOG_level_list_Alyrata_exp_DEG_Alyrata.tsv", row.names = F)

###Hypergeometric test for over representation of HOGs with DEG in HOG in "Conserved" Data Set
# Hypergeometric test (comparable to Fisher's exact test) for enrichment of exp.SPECIES OGs with DEGs over exp.SPECIES OGs in Conserved
# q = OGs with diff exp genes in Hypothesis OGs (= success in the sample)
# m = OGs with diff exp genes in Conserved OGs (= population)
# n = Conserved OGs - m
# k = OGs in Hypothesis OGs (= sample)

# Hypothesis 3: Alyrata vs. sensitive with DEG Any
# q = 2k)
q <- length(HOGs_Alyrata_exp_DEG_any)
# m = 2b)
m <- length(HOGs_DEG)
# n = 2.1)b)2) - m
n <- (length(HOGs) - m)
# 2.1)k)2)
k <- length(HOGs_Alyrata_exp)

p <- phyper((q-1), m, n, k, lower.tail=F)
p # 0.2101261

# Hypothesis 3: Alyrata vs. sensitive with DEG Alyrata
# q = 2l)
q <- length(HOGs_Alyrata_exp_DEG_Alyrata)
# m= 2.1)l)1)
m <- length(HOGs_DEG_Alyrata)
# n = 2.1)b)2) - m
n <- (length(HOGs) - m)
# 2.1)k)2)
k <- length(HOGs_Alyrata_exp)

p <- phyper((q-1), m, n, k, lower.tail=F)
p # 1.032192e-05


### Export a matrix with the HOGs and the Genes from Hypothesis_3 expanded OGs:
Hypothesis_3_Exp_matrix <- Conserved_dataset_matrix_ordered[Conserved_dataset_matrix_ordered$HOG %in% HOGs_Alyrata_exp,]
write.table(Hypothesis_3_Exp_matrix, paste0(OUTDIR, "/Hypothesis_3_Exp_in_Conserved_dataset_matrix.tsv"), sep="\t", row.names=F, col.names=T, quote=F)

# export a matrix with the HOGs and the Genes from the Hypothesis_3 expanded and DE HOGs with the genes function (human readable description (HRD))
## extract the HOGs with the HRD of each gene
HOGs_Alyrata_exp_DEG_Alyrata_with_HRD <- rbind(SFA$Arabidopsis_thaliana[SFA$Arabidopsis_thaliana$HOG %in% HOGs_Alyrata_exp_DEG_Alyrata,],
                                               SFA$Arabidopsis_lyrata[SFA$Arabidopsis_lyrata$HOG %in% HOGs_Alyrata_exp_DEG_Alyrata,],
                                               SFA$Eutrema_salsugineum[SFA$Eutrema_salsugineum$HOG %in% HOGs_Alyrata_exp_DEG_Alyrata,],
                                               SFA$Brassica_napus[SFA$Brassica_napus$HOG %in% HOGs_Alyrata_exp_DEG_Alyrata,])

# combine the table with the expression data
HOGs_Alyrata_exp_DEG_Alyrata_with_HRD_and_expression <- merge(HOGs_Alyrata_exp_DEG_Alyrata_with_HRD, Conserved_with_expression, by.x="Protein-Accession", by.y = "gene")
HOGs_Alyrata_exp_DEG_Alyrata_with_HRD_and_expression_publish <- HOGs_Alyrata_exp_DEG_Alyrata_with_HRD_and_expression[,c(1,2,5,7:9,12:14)]
HOGs_Alyrata_exp_DEG_Alyrata_with_HRD_and_expression_publish_ordered <- HOGs_Alyrata_exp_DEG_Alyrata_with_HRD_and_expression_publish[order(HOGs_Alyrata_exp_DEG_Alyrata_with_HRD_and_expression_publish$HOG.x, HOGs_Alyrata_exp_DEG_Alyrata_with_HRD_and_expression_publish$species),]

# export a table with all the information to publish
write.table(HOGs_Alyrata_exp_DEG_Alyrata_with_HRD_and_expression_publish_ordered, paste0(OUTDIR, "/HOGs_Alyrata_exp_DEG_Alyrata_with_HRD_and_expression.tsv"), sep="\t", row.names=F, col.names=T, quote=F)


### Hypothesis 5 test
# 2.1)b)2) Number of OGs in "Conserved"
HOGs <- HOG_level_list$hypothesis_1_cons$HOG
# 2b) What is the frequency of OGs with at least 1 DEG in "Conserved" OGs?
HOGs_DEG <- HOG_level_list$hypothesis_1_cons[!is.na(HOG_level_list$hypothesis_1_cons$total_sigDE) & HOG_level_list$hypothesis_1_cons$total_sigDE>=1,]$HOG
# 2.1)n)1) What is the frequency of OGs with at least 1 DEG from the Hypothesis 5 expanded species in "Conserved" OGs?
HOGs_DEG_Athaliana <- HOG_level_list$hypothesis_5_cons[HOG_level_list$hypothesis_5_cons$Arabidopsis_thaliana_sigDE>=1 & !is.na(HOG_level_list$hypothesis_5_cons$Arabidopsis_thaliana_sigDE)
                                                       ,]$HOG
# 2.1)m)2) Within "Conserved": Number of OGs with "Hypothesis 5 expanded==yes"
HOGs_Athaliana_exp <- HOG_level_list$hypothesis_5_cons[HOG_level_list$hypothesis_5_cons$expansion == "yes",]$HOG
# 2m) Within "Conserved": What is the frequency of OGs with at least 1 DEG in "Hypothesis 5 expanded==yes" OGs?
cons_1_and_5_merged_2 <-  merge(HOG_level_list$hypothesis_1_cons[,c("HOG", "total_sigDE")], HOG_level_list$hypothesis_5_cons, by="HOG")
# select for total_sigDE.x because we are interested in total_sigDE from all 4 species (inferred from data frame HOG_level_list$hypothesis_1_cons, which contains all 4 species)
HOGs_Athaliana_exp_DEG_any <- cons_1_and_5_merged_2[cons_1_and_5_merged_2$expansion == "yes" & !is.na(cons_1_and_5_merged_2$total_sigDE.x) & cons_1_and_5_merged_2$total_sigDE.x>=1,]$HOG
# 2n) What is the frequency of OGs with at least 1 DEG from any of the expanded species in "Hypothesis 5 expanded==yes" OGs?
HOGs_Athaliana_exp_DEG_Athaliana <- HOG_level_list$hypothesis_5_cons[HOG_level_list$hypothesis_5_cons$expansion == "yes" & HOG_level_list$hypothesis_5_cons$Arabidopsis_thaliana_sigDE>=1 & !is.na(HOG_level_list$hypothesis_5_cons$Arabidopsis_thaliana_sigDE)
                                                                     ,]$HOG
###Hypergeometric test for over representation of HOGs with DEG in HOG in "Conserved" Data Set
# Hypergeometric test (comparable to Fisher's exact test) for enrichment of exp.SPECIES OGs with DEGs over exp.SPECIES OGs in Conserved
# q = OGs with diff exp genes in Hypothesis OGs (= success in the sample)
# m = OGs with diff exp genes in Conserved OGs (= population)
# n = Conserved OGs - m
# k = OGs in Hypothesis OGs (= sample)

# Hypothesis 5: Athaliana vs. tolerant with DEG Any
# q = 2m)
q <- length(HOGs_Athaliana_exp_DEG_any)
# m = 2b)
m <- length(HOGs_DEG)
# n = 2.1)b)2) - m
n <- (length(HOGs) - m)
# 2.1)m)2)
k <- length(HOGs_Athaliana_exp)

p <- phyper((q-1), m, n, k, lower.tail=F)
p # 0.8848153

# Hypothesis 5: Athaliana vs. tolerant with DEG Athaliana
# q = 2n)
q <- length(HOGs_Athaliana_exp_DEG_Athaliana)
# m= 2.1)n)1)
m <- length(HOGs_DEG_Athaliana)
# n = 2.1)b)2) - m
n <- (length(HOGs) - m)
# 2.1)m)2)
k <- length(HOGs_Athaliana_exp)

p <- phyper((q-1), m, n, k, lower.tail=F)
p # 0.000350742


### Export a matrix with the HOGs and the Genes from Hypothesis_5 expanded OGs:
Hypothesis_5_Exp_matrix <- Conserved_dataset_matrix_ordered[Conserved_dataset_matrix_ordered$HOG %in% HOGs_Athaliana_exp,]
write.table(Hypothesis_5_Exp_matrix, paste0(OUTDIR, "/Hypothesis_5_Exp_in_Conserved_dataset_matrix.tsv"), sep="\t", row.names=F, col.names=T, quote=F)


### Hypothesis 6 test
# 2.1)b)2) Number of OGs in "Conserved"
HOGs <- HOG_level_list$hypothesis_1_cons$HOG
# 2b) What is the frequency of OGs with at least 1 DEG in "Conserved" OGs?
HOGs_DEG <- HOG_level_list$hypothesis_1_cons[!is.na(HOG_level_list$hypothesis_1_cons$total_sigDE) & HOG_level_list$hypothesis_1_cons$total_sigDE>=1,]$HOG
# 2.1)p)1) What is the frequency of OGs with at least 1 DEG from the Hypothesis 6 expanded species in "Conserved" OGs?
HOGs_DEG_Bnapus <- HOG_level_list$hypothesis_6_cons[HOG_level_list$hypothesis_6_cons$Brassica_napus_sigDE>=1 & !is.na(HOG_level_list$hypothesis_6_cons$Brassica_napus_sigDE)
                                                    ,]$HOG
# 2.1)o)2) Within "Conserved": Number of OGs with "Hypothesis 6 expanded==yes"
HOGs_Bnapus_exp <- HOG_level_list$hypothesis_6_cons[HOG_level_list$hypothesis_6_cons$expansion == "yes",]$HOG
# 2o) Within "Conserved": What is the frequency of OGs with at least 1 DEG in "Hypothesis 6 expanded==yes" OGs?
cons_1_and_6_merged_2 <-  merge(HOG_level_list$hypothesis_1_cons[,c("HOG", "total_sigDE")], HOG_level_list$hypothesis_6_cons, by="HOG")
# select for total_sigDE.x because we are interested in total_sigDE from all 4 species (inferred from data frame HOG_level_list$hypothesis_1_cons, which contains all 4 species)
HOGs_Bnapus_exp_DEG_any <- cons_1_and_6_merged_2[cons_1_and_6_merged_2$expansion == "yes" & !is.na(cons_1_and_6_merged_2$total_sigDE.x) & cons_1_and_6_merged_2$total_sigDE.x>=1,]$HOG
# 2p) What is the frequency of OGs with at least 1 DEG from any of the expanded species in "Hypothesis 6 expanded==yes" OGs?
HOGs_Bnapus_exp_DEG_Bnapus <- HOG_level_list$hypothesis_6_cons[HOG_level_list$hypothesis_6_cons$expansion == "yes" & HOG_level_list$hypothesis_6_cons$Brassica_napus_sigDE>=1 & !is.na(HOG_level_list$hypothesis_6_cons$Brassica_napus_sigDE)
                                                               ,]$HOG
###Hypergeometric test for over representation of HOGs with DEG in HOG in "Conserved" Data Set
# Hypergeometric test (comparable to Fisher's exact test) for enrichment of exp.SPECIES OGs with DEGs over exp.SPECIES OGs in Conserved
# q = OGs with diff exp genes in Hypothesis OGs (= success in the sample)
# m = OGs with diff exp genes in Conserved OGs (= population)
# n = Conserved OGs - m
# k = OGs in Hypothesis OGs (= sample)

# Hypothesis 6: Bnapus vs. tolerant with DEG Any
# q = 2o)
q <- length(HOGs_Bnapus_exp_DEG_any)
# m = 2b)
m <- length(HOGs_DEG)
# n = 2.1)b)2) - m
n <- (length(HOGs) - m)
# 2.1)o)2)
k <- length(HOGs_Bnapus_exp)

p <- phyper((q-1), m, n, k, lower.tail=F)
p # 0.0002281104

# Hypothesis 6: Bnapus vs. tolerant with DEG Bnapus
# q = 2p)
q <- length(HOGs_Bnapus_exp_DEG_Bnapus)
# m= 2.1)p)1)
m <- length(HOGs_DEG_Bnapus)
# n = 2.1)b)2) - m
n <- (length(HOGs) - m)
# 2.1)o)2)
k <- length(HOGs_Bnapus_exp)

p <- phyper((q-1), m, n, k, lower.tail=F)
p # 3.794396e-34

### Export a matrix with the HOGs and the Genes from Hypothesis_6 expanded OGs:
Hypothesis_6_Exp_matrix <- Conserved_dataset_matrix_ordered[Conserved_dataset_matrix_ordered$HOG %in% HOGs_Bnapus_exp,]
write.table(Hypothesis_6_Exp_matrix, paste0(OUTDIR, "/Hypothesis_6_Exp_in_Conserved_dataset_matrix.tsv"), sep="\t", row.names=F, col.names=T, quote=F)


                          ### Plot HOG SIZES ###
######## IMPORTANT NOTE: Please skip to and run "Conserved Set" in order to interprete results of the upcoming plots!!

## "ALL" Set
HOG_level_list$hypothesis_1$HOG_sizes <- apply(HOG_level_list$hypothesis_1[,c("Arabidopsis_lyrata_total", "Arabidopsis_thaliana_total", "Eutrema_salsugineum_total", "Brassica_napus_total")], 1, sum)
pdf(paste0(OUTDIR, "/Boxplot_OG_sizes_total_set.pdf"), 6, 4)
boxplot(HOG_level_list$hypothesis_1$HOG_sizes, ylab="OG sizes", main="Total Set")
dev.off()
summary(HOG_level_list$hypothesis_1$HOG_sizes)
hist(HOG_level_list$hypothesis_1$HOG_sizes)

## "Conserved" Set
HOG_level_list$hypothesis_1_cons$HOG_sizes <- apply(HOG_level_list$hypothesis_1_cons[,c("Arabidopsis_lyrata_total", "Arabidopsis_thaliana_total", "Eutrema_salsugineum_total", "Brassica_napus_total")], 1, sum)
pdf(paste0(OUTDIR, "/Boxplot_OG_sizes_Conserved_set.pdf"), 6, 4)
boxplot(HOG_level_list$hypothesis_1_cons$HOG_sizes, ylab="OG sizes", main="Conserved Set")
dev.off()
hist(HOG_level_list$hypothesis_1_cons$HOG_sizes, xlab = "Sizes of OGs in Conserved", main=NA) #breaks = seq(0, 50, 2), freq = F,
summary(HOG_level_list$hypothesis_1_cons$HOG_sizes)
# -> NOTE: the outliers from "ALL" Set are mainly removed when focusing on Conserved HOGs
## "DEG" in Conserved Set
HOGs_DEG <- HOG_level_list$hypothesis_1_cons[!is.na(HOG_level_list$hypothesis_1_cons$total_sigDE) & HOG_level_list$hypothesis_1_cons$total_sigDE>=1,]$HOG
HOG_level_list$hypothesis_1_cons$HOG_sizes <- apply(HOG_level_list$hypothesis_1_cons[,c("Arabidopsis_lyrata_total", "Arabidopsis_thaliana_total", "Eutrema_salsugineum_total", "Brassica_napus_total")], 1, sum)
pdf(paste0(OUTDIR, "/Boxplot_OG_sizes_DEG_in_Conserved_set.pdf"), 6, 4)
boxplot(HOG_level_list$hypothesis_1_cons[HOG_level_list$hypothesis_1_cons$HOG %in% HOGs_DEG,]$HOG_sizes)
dev.off()
hist(HOG_level_list$hypothesis_1_cons[HOG_level_list$hypothesis_1_cons$HOG %in% HOGs_DEG,]$HOG_sizes, xlab = "Sizes of OGs with DEG in Conserved", main=NA) #, breaks = seq(0, 50, 2), freq = F, ylim = c(0,0.4)
summary(HOG_level_list$hypothesis_1_cons[HOG_level_list$hypothesis_1_cons$HOG %in% HOGs_DEG,]$HOG_sizes)

## "Conserved_2" Set (BC. BNAPUS IS TETRAPLOID)
#HOG_level_list$hypothesis_1_cons_2$HOG_sizes <- apply(HOG_level_list$hypothesis_1_cons_2[,c("Arabidopsis_lyrata_total", "Arabidopsis_thaliana_total", "Eutrema_salsugineum_total", "Brassica_napus_total")], 1, sum)
#boxplot(HOG_level_list$hypothesis_1_cons_2$HOG_sizes)
#hist(HOG_level_list$hypothesis_1_cons_2$HOG_sizes, xlab = "Sizes of OGs in Conserved_2", main=NA) # breaks = seq(0, 50, 2), freq = F, ylim = c(0,0.4),
#summary(HOG_level_list$hypothesis_1_cons_2$HOG_sizes)


# Stacked histogram Conserved without DEG and Conserved with DEG OG sizes
Cons_with_DEG <- HOG_level_list$hypothesis_1_cons[HOG_level_list$hypothesis_1_cons$HOG %in% HOGs_DEG,]$HOG_sizes
Cons_without_DEG <- HOG_level_list$hypothesis_1_cons[!(HOG_level_list$hypothesis_1_cons$HOG %in% HOGs_DEG),]$HOG_sizes
bucket <- list(Cons_with_DEG, Cons_without_DEG)  # this puts all values in one list
# the melt command flattens the 'bucket' list into value/vectorname pairs
# the 2 columns are called 'value' and 'L1' by default
# 'fill' will color bars differently depending on L1 group
pdf(paste0(OUTDIR, "/Histogram_stacked_OG_sizes_with_and_without_DEG_in_Conserved_set.pdf"), 6, 4)
ggplot(melt(bucket), aes(value, fill = as.factor(L1))) + 
  #call geom_histogram with position="dodge" to offset the bars and manual binwidth of 2
#  geom_histogram(aes(y = ..density..), binwidth = 2)+
  geom_histogram(binwidth = 1)+
  scale_fill_manual(name="Dataset",labels = c("with DEG", "no DEG"), values = c("1", "2"))+
  xlab("OG size")+
  xlim(c(0,30))
dev.off()

# Stacked histogram Conserved_2 without DEG and Conserved_2 with DEG OG sizes
#Cons_2_with_DEG <- HOG_level_list$hypothesis_1_cons_2[HOG_level_list$hypothesis_1_cons_2$HOG %in% HOGs_DEG,]$HOG_sizes
#Cons_2_without_DEG <- HOG_level_list$hypothesis_1_cons_2[!(HOG_level_list$hypothesis_1_cons_2$HOG %in% HOGs_DEG),]$HOG_sizes
#bucket <- list(Cons_2_with_DEG, Cons_2_without_DEG)  # this puts all values in one list
# the melt command flattens the 'bucket' list into value/vector name pairs
# the 2 columns are called 'value' and 'L1' by default
# 'fill' will color bars differently depending on L1 group
#ggplot(melt(bucket), aes(value, fill = as.factor(L1))) + 
  #call geom_histogram with position="dodge" to offset the bars and manual binwidth of 2
  #  geom_histogram(aes(y = ..density..), binwidth = 2)+
#  geom_histogram(binwidth = 1)+
#  scale_fill_manual(name="Dataset",labels = c("with DEG", "no DEG"), values = c("1", "2"))+
#  xlab("OG size")+
#  xlim(c(0,30))

### order the data.frame of HOGs labeled with "DEG" or "no DEG" by HOG size
n <- 50
DEG_label <- rep("DEG", times=length(Cons_with_DEG))
no_DEG_label <- rep("noDEG", times=length(Cons_without_DEG))
data <- data.frame(HOG_sizes=c(Cons_with_DEG, Cons_without_DEG), label=c(DEG_label, no_DEG_label))
ordered_data <- data[order(data$HOG_sizes),]
grouping <- factor(character(length = length(ordered_data$HOG_sizes)))
grouping <- rep(seq(1,ceiling(length(grouping)/n)),each=n)
grouping <- as.factor(as.character(grouping[1:17488]))

ordered_data$grouping <- grouping

count_of_OGs_with_DEG_per_grouping<- table(ordered_data$label=="DEG", grouping)[2,]
count_df <- data.frame(unclass(count_of_OGs_with_DEG_per_grouping))
count_df$group <- row.names(count_df)
mean_OG_size_of_all_OGs_per_grouping<- tapply(ordered_data$HOG_sizes, ordered_data$grouping, mean)
mean_df <- data.frame(unclass(mean_OG_size_of_all_OGs_per_grouping))
mean_df$group <- row.names(mean_df)

count_and_mean <- merge(count_df, mean_df, by="group")
names(count_and_mean) <- c("group", "count_of_OGs_with_DEG", "mean_OG_size")
count_and_mean$percent_OGs_with_DEG <- (count_and_mean$count_of_OGs_with_DEG/n*100)

pdf(paste0(OUTDIR, "/Jitter_plot_with_regression_of_mean_OG_sizes_of_350_groups_of_size_50_Conserved_DEG_Any.pdf"), 6, 4)
ggplot(count_and_mean, aes(x=mean_OG_size, y=percent_OGs_with_DEG))+
  #  geom_point(aes(x=mean_OG_size, y=percent_OGs_with_DEG), shape=21, fill=3, alpha=0.3)
  geom_jitter(shape=21, fill=1, alpha=0.3)+
  geom_smooth(method="lm", color=1)
dev.off()


## Hypothesis 1 Set
Hypothesis_1_Cons_HOGs_all_tol_exp <- HOG_level_list$hypothesis_1_cons[HOG_level_list$hypothesis_1_cons$expansion == "yes",]
Hypothesis_1_Cons_HOGs_all_tol_exp$HOG_sizes <- apply(Hypothesis_1_Cons_HOGs_all_tol_exp[,c("Arabidopsis_lyrata_total", "Arabidopsis_thaliana_total", "Eutrema_salsugineum_total", "Brassica_napus_total")], 1, sum)
boxplot(Hypothesis_1_Cons_HOGs_all_tol_exp$HOG_sizes)
hist(Hypothesis_1_Cons_HOGs_all_tol_exp$HOG_sizes, breaks = seq(0, 50, 2), freq = F, ylim = c(0,0.4), xlab = "Sizes of OGs expanded in Tolerant", main=NA)
summary(Hypothesis_1_Cons_HOGs_all_tol_exp$HOG_sizes)

# Stacked histogram Hypothesis 1 without DEG and Hypothesis 1 with DEG Any OG sizes
Hypo1_with_DEG <- HOG_level_list$hypothesis_1_cons[(HOG_level_list$hypothesis_1_cons$HOG %in% HOGs_DEG) & HOG_level_list$hypothesis_1_cons$expansion=="yes" ,]$HOG_sizes
Hypo1_without_DEG <- HOG_level_list$hypothesis_1_cons[(!(HOG_level_list$hypothesis_1_cons$HOG %in% HOGs_DEG)) & HOG_level_list$hypothesis_1_cons$expansion=="yes",]$HOG_sizes
bucket <- list(Hypo1_with_DEG, Hypo1_without_DEG)  # this puts all values in one list
# the melt command flattens the 'bucket' list into value/vectorname pairs
# the 2 columns are called 'value' and 'L1' by default
# 'fill' will color bars differently depending on L1 group
pdf(paste0(OUTDIR, "/Histogram_stacked_OG_sizes_with_and_without_DEG_in_Hypothesis_1_set.pdf"), 6, 4)
ggplot(melt(bucket), aes(value, fill = as.factor(L1))) + 
  #call geom_histogram with position="dodge" to offset the bars and manual binwidth of 2
#  geom_histogram(aes(y = ..density..), binwidth = 1)+
  geom_histogram(binwidth = 1)+
  scale_fill_manual(name="Dataset",labels = c("with DEG", "no DEG"), values = c("1", "2"))+
  xlab("OG size")+
  xlim(c(0,30))
dev.off()

### order the data.frame of HOGs labeled with "DEG" or "no DEG" by HOG size
n <- 5
DEG_label <- rep("DEG", times=length(Hypo1_with_DEG))
no_DEG_label <- rep("noDEG", times=length(Hypo1_without_DEG))
data <- data.frame(HOG_sizes=c(Hypo1_with_DEG, Hypo1_without_DEG), label=c(DEG_label, no_DEG_label))
ordered_data <- data[order(data$HOG_sizes),]
grouping <- factor(character(length = length(ordered_data$HOG_sizes)))
grouping <- rep(seq(1,ceiling(length(grouping)/n)),each=n)
grouping <- as.factor(as.character(grouping[1:144]))

ordered_data$grouping <- grouping

count_of_OGs_with_DEG_per_grouping<- table(ordered_data$label=="DEG", grouping)[2,]
count_df <- data.frame(unclass(count_of_OGs_with_DEG_per_grouping))
count_df$group <- row.names(count_df)
mean_OG_size_of_all_OGs_per_grouping<- tapply(ordered_data$HOG_sizes, ordered_data$grouping, mean)
mean_df <- data.frame(unclass(mean_OG_size_of_all_OGs_per_grouping))
mean_df$group <- row.names(mean_df)

count_and_mean <- merge(count_df, mean_df, by="group")
names(count_and_mean) <- c("group", "count_of_OGs_with_DEG", "mean_OG_size")
count_and_mean$percent_OGs_with_DEG <- (count_and_mean$count_of_OGs_with_DEG/n*100)

pdf(paste0(OUTDIR, "/Jitter_plot_with_regression_of_mean_OG_sizes_of_29_groups_of_size_5_Hypothesis_1_DEG_Any.pdf"), 6, 4)
ggplot(count_and_mean, aes(x=mean_OG_size, y=percent_OGs_with_DEG))+
  #  geom_point(aes(x=mean_OG_size, y=percent_OGs_with_DEG), shape=21, fill=3, alpha=0.3)
  geom_jitter(shape=21, fill=1, alpha=0.3)+
  geom_smooth(method="lm", color=1)
dev.off()

# Stacked histogram Conserved without DEG from both tolerant and Conserved with DEG from both tolerant OG sizes
Cons_with_DEG_both_tol <- HOG_level_list$hypothesis_1_cons[HOG_level_list$hypothesis_1_cons$HOG %in% HOGs_DEG_both_tol,]$HOG_sizes
Cons_without_DEG_both_tol <- HOG_level_list$hypothesis_1_cons[!(HOG_level_list$hypothesis_1_cons$HOG %in% HOGs_DEG_both_tol),]$HOG_sizes
bucket <- list(Cons_with_DEG_both_tol, Cons_without_DEG_both_tol)  # this puts all values in one list
# the melt command flattens the 'bucket' list into value/vectorname pairs
# the 2 columns are called 'value' and 'L1' by default
# 'fill' will color bars differently depending on L1 group
pdf(paste0(OUTDIR, "/Histogram_stacked_OG_sizes_with_and_without_DEG_from_both_tol_in_Conserved_set.pdf"), 6, 4)
ggplot(melt(bucket), aes(value, fill = as.factor(L1))) + 
  #call geom_histogram with position="dodge" to offset the bars and manual binwidth of 2
  #  geom_histogram(aes(y = ..density..), binwidth = 2)+
  geom_histogram(binwidth = 1)+
  scale_fill_manual(name="Dataset",labels = c("with DEG from both tol", "no DEG from both tol"), values = c("1", "2"))+
  xlab("OG size")+
  xlim(c(0,30))
dev.off()

### order the data.frame of HOGs labeled with "DEG" or "no DEG" by HOG size
n <- 50
DEG_label <- rep("DEG", times=length(Cons_with_DEG_both_tol))
no_DEG_label <- rep("noDEG", times=length(Cons_without_DEG_both_tol))
data <- data.frame(HOG_sizes=c(Cons_with_DEG_both_tol, Cons_without_DEG_both_tol), label=c(DEG_label, no_DEG_label))
ordered_data <- data[order(data$HOG_sizes),]
grouping <- factor(character(length = length(ordered_data$HOG_sizes)))
grouping <- rep(seq(1,ceiling(length(grouping)/n)),each=n)
grouping <- as.factor(as.character(grouping[1:17488]))

ordered_data$grouping <- grouping

count_of_OGs_with_DEG_per_grouping<- table(ordered_data$label=="DEG", grouping)[2,]
count_df <- data.frame(unclass(count_of_OGs_with_DEG_per_grouping))
count_df$group <- row.names(count_df)
mean_OG_size_of_all_OGs_per_grouping<- tapply(ordered_data$HOG_sizes, ordered_data$grouping, mean)
mean_df <- data.frame(unclass(mean_OG_size_of_all_OGs_per_grouping))
mean_df$group <- row.names(mean_df)

count_and_mean <- merge(count_df, mean_df, by="group")
names(count_and_mean) <- c("group", "count_of_OGs_with_DEG", "mean_OG_size")
count_and_mean$percent_OGs_with_DEG <- (count_and_mean$count_of_OGs_with_DEG/n*100)

pdf(paste0(OUTDIR, "/Jitter_plot_with_regression_of_mean_OG_sizes_of_350_groups_of_size_50_Conserved_DEG_from_both_tolerant.pdf"), 6, 4)
ggplot(count_and_mean, aes(x=mean_OG_size, y=percent_OGs_with_DEG))+
  #  geom_point(aes(x=mean_OG_size, y=percent_OGs_with_DEG), shape=21, fill=3, alpha=0.3)
  geom_jitter(shape=21, fill=1, alpha=0.3)+
  geom_smooth(method="lm", color=1)
dev.off()

# Stacked histogram Hypothesis 1 without DEG from both tolerant and Hypothesis 1 with DEG from both tolerant OG sizes
Hypo1_with_DEG_both_tol <- HOG_level_list$hypothesis_1_cons[HOG_level_list$hypothesis_1_cons$HOG %in% HOGs_all_tol_exp_DEG_both_tol,]$HOG_sizes
Hypo1_without_DEG_both_tol <- HOG_level_list$hypothesis_1_cons[HOG_level_list$hypothesis_1_cons$HOG %in% HOGs_all_tol_exp & !(HOG_level_list$hypothesis_1_cons$HOG %in% HOGs_DEG_both_tol),]$HOG_sizes
bucket <- list(Hypo1_with_DEG_both_tol, Hypo1_without_DEG_both_tol)  # this puts all values in one list
# the melt command flattens the 'bucket' list into value/vectorname pairs
# the 2 columns are called 'value' and 'L1' by default
# 'fill' will color bars differently depending on L1 group
pdf(paste0(OUTDIR, "/Histogram_stacked_OG_sizes_with_and_without_DEG_from_both_tol_in_Hypothesis_1_set.pdf"), 6, 4)
ggplot(melt(bucket), aes(value, fill = as.factor(L1))) + 
  #call geom_histogram with position="dodge" to offset the bars and manual binwidth of 2
  #  geom_histogram(aes(y = ..density..), binwidth = 1)+
  geom_histogram(binwidth = 1)+
  scale_fill_manual(name="Dataset",labels = c("with DEG from both tol", "no DEG from both tol"), values = c("1", "2"))+
  xlab("OG size")+
  xlim(c(0,30))
dev.off()


### order the data.frame of HOGs labeled with "DEG" or "no DEG" by HOG size
n <- 5
DEG_label <- rep("DEG", times=length(Hypo1_with_DEG_both_tol))
no_DEG_label <- rep("noDEG", times=length(Hypo1_without_DEG_both_tol))
data <- data.frame(HOG_sizes=c(Hypo1_with_DEG_both_tol, Hypo1_without_DEG_both_tol), label=c(DEG_label, no_DEG_label))
ordered_data <- data[order(data$HOG_sizes),]
grouping <- factor(character(length = length(ordered_data$HOG_sizes)))
grouping <- rep(seq(1,ceiling(length(grouping)/n)),each=n)
grouping <- as.factor(as.character(grouping[1:144]))

ordered_data$grouping <- grouping

count_of_OGs_with_DEG_per_grouping<- table(ordered_data$label=="DEG", grouping)[2,]
count_df <- data.frame(unclass(count_of_OGs_with_DEG_per_grouping))
count_df$group <- row.names(count_df)
mean_OG_size_of_all_OGs_per_grouping<- tapply(ordered_data$HOG_sizes, ordered_data$grouping, mean)
mean_df <- data.frame(unclass(mean_OG_size_of_all_OGs_per_grouping))
mean_df$group <- row.names(mean_df)

count_and_mean <- merge(count_df, mean_df, by="group")
names(count_and_mean) <- c("group", "count_of_OGs_with_DEG", "mean_OG_size")
count_and_mean$percent_OGs_with_DEG <- (count_and_mean$count_of_OGs_with_DEG/n*100)

pdf(paste0(OUTDIR, "/Jitter_plot_with_regression_of_mean_OG_sizes_of_29_groups_of_size_5_Hypothesis_1_DEG_from_both_tol.pdf"), 6, 4)
ggplot(count_and_mean, aes(x=mean_OG_size, y=percent_OGs_with_DEG))+
  #  geom_point(aes(x=mean_OG_size, y=percent_OGs_with_DEG), shape=21, fill=3, alpha=0.3)
  geom_jitter(shape=21, fill=1, alpha=0.3)+
  geom_smooth(method="lm", color=1)
dev.off()


## Hypothesis 4 Set
hypothesis_4_Cons_HOGs_all_sens_exp <- HOG_level_list$hypothesis_4_cons[HOG_level_list$hypothesis_4_cons$expansion == "yes",]
hypothesis_4_Cons_HOGs_all_sens_exp$HOG_sizes <- apply(hypothesis_4_Cons_HOGs_all_sens_exp[,c("Arabidopsis_lyrata_total", "Arabidopsis_thaliana_total", "Eutrema_salsugineum_total", "Brassica_napus_total")], 1, sum)
boxplot(hypothesis_4_Cons_HOGs_all_sens_exp$HOG_sizes)
hist(hypothesis_4_Cons_HOGs_all_sens_exp$HOG_sizes, breaks = seq(0, 50, 2), freq = F, ylim = c(0,0.4), xlab = "Sizes of OGs expanded in Sensitive", main=NA)
summary(hypothesis_4_Cons_HOGs_all_sens_exp$HOG_sizes)

# Stacked histogram Hypothesis 4 without DEG and Hypothesis 4 with DEG Any OG sizes
Hypo4_with_DEG <- hypothesis_4_Cons_HOGs_all_sens_exp[hypothesis_4_Cons_HOGs_all_sens_exp$HOG %in% HOGs_DEG ,]$HOG_sizes
Hypo4_without_DEG <-hypothesis_4_Cons_HOGs_all_sens_exp[!(hypothesis_4_Cons_HOGs_all_sens_exp$HOG %in% HOGs_DEG) ,]$HOG_sizes
bucket <- list(Hypo4_with_DEG, Hypo4_without_DEG)  # this puts all values in one list
# the melt command flattens the 'bucket' list into value/vectorname pairs
# the 2 columns are called 'value' and 'L1' by default
# 'fill' will color bars differently depending on L1 group
pdf(paste0(OUTDIR, "/Histogram_stacked_OG_sizes_with_and_without_DEG_in_Hypothesis_4_set.pdf"), 6, 4)
ggplot(melt(bucket), aes(value, fill = as.factor(L1))) + 
  #call geom_histogram with position="dodge" to offset the bars and manual binwidth of 2
  #  geom_histogram(aes(y = ..density..), binwidth = 1)+
  geom_histogram(binwidth = 1)+
  scale_fill_manual(name="Dataset",labels = c("with DEG", "no DEG"), values = c("1", "2"))+
  xlab("OG size")+
  xlim(c(0,30))
dev.off()

### order the data.frame of HOGs labeled with "DEG" or "no DEG" by HOG size
n <- 5
DEG_label <- rep("DEG", times=length(Hypo4_with_DEG))
no_DEG_label <- rep("noDEG", times=length(Hypo4_without_DEG))
data <- data.frame(HOG_sizes=c(Hypo4_with_DEG, Hypo4_without_DEG), label=c(DEG_label, no_DEG_label))
ordered_data <- data[order(data$HOG_sizes),]
grouping <- factor(character(length = length(ordered_data$HOG_sizes)))
grouping <- rep(seq(1,ceiling(length(grouping)/n)),each=n)
grouping <- as.factor(as.character(grouping[1:58]))

ordered_data$grouping <- grouping

count_of_OGs_with_DEG_per_grouping<- table(ordered_data$label=="DEG", grouping)[2,]
count_df <- data.frame(unclass(count_of_OGs_with_DEG_per_grouping))
count_df$group <- row.names(count_df)
mean_OG_size_of_all_OGs_per_grouping<- tapply(ordered_data$HOG_sizes, ordered_data$grouping, mean)
mean_df <- data.frame(unclass(mean_OG_size_of_all_OGs_per_grouping))
mean_df$group <- row.names(mean_df)

count_and_mean <- merge(count_df, mean_df, by="group")
names(count_and_mean) <- c("group", "count_of_OGs_with_DEG", "mean_OG_size")
count_and_mean$percent_OGs_with_DEG <- (count_and_mean$count_of_OGs_with_DEG/n*100)

pdf(paste0(OUTDIR, "/Jitter_plot_with_regression_of_mean_OG_sizes_of_12_groups_of_size_5_Hypothesis_4_DEG_Any.pdf"), 6, 4)
ggplot(count_and_mean, aes(x=mean_OG_size, y=percent_OGs_with_DEG))+
  #  geom_point(aes(x=mean_OG_size, y=percent_OGs_with_DEG), shape=21, fill=3, alpha=0.3)
  geom_jitter(shape=21, fill=1, alpha=0.3)+
  geom_smooth(method="lm", color=1)
dev.off()

# Stacked histogram Conserved without DEG from both sensitive and Conserved with DEG from both sensitive OG sizes
Cons_with_DEG_both_sens <- HOG_level_list$hypothesis_1_cons[HOG_level_list$hypothesis_1_cons$HOG %in% HOGs_DEG_both_sens,]$HOG_sizes
Cons_without_DEG_both_sens <- HOG_level_list$hypothesis_1_cons[!(HOG_level_list$hypothesis_1_cons$HOG %in% HOGs_DEG_both_sens),]$HOG_sizes
bucket <- list(Cons_with_DEG_both_sens, Cons_without_DEG_both_sens)  # this puts all values in one list
# the melt command flattens the 'bucket' list into value/vectorname pairs
# the 2 columns are called 'value' and 'L1' by default
# 'fill' will color bars differently depending on L1 group
pdf(paste0(OUTDIR, "/Histogram_stacked_OG_sizes_with_and_without_DEG_from_both_sens_in_Conserved_set.pdf"), 6, 4)
ggplot(melt(bucket), aes(value, fill = as.factor(L1))) + 
  #call geom_histogram with position="dodge" to offset the bars and manual binwidth of 2
  #  geom_histogram(aes(y = ..density..), binwidth = 2)+
  geom_histogram(binwidth = 1)+
  scale_fill_manual(name="Dataset",labels = c("with DEG from both sens", "no DEG from both sens"), values = c("1", "2"))+
  xlab("OG size")+
  xlim(c(0,30))
dev.off()

### order the data.frame of HOGs labeled with "DEG" or "no DEG" by HOG size
n <- 50
DEG_label <- rep("DEG", times=length(Cons_with_DEG_both_sens))
no_DEG_label <- rep("noDEG", times=length(Cons_without_DEG_both_sens))
data <- data.frame(HOG_sizes=c(Cons_with_DEG_both_sens, Cons_without_DEG_both_sens), label=c(DEG_label, no_DEG_label))
ordered_data <- data[order(data$HOG_sizes),]
grouping <- factor(character(length = length(ordered_data$HOG_sizes)))
grouping <- rep(seq(1,ceiling(length(grouping)/n)),each=n)
grouping <- as.factor(as.character(grouping[1:17488]))

ordered_data$grouping <- grouping

count_of_OGs_with_DEG_per_grouping<- table(ordered_data$label=="DEG", grouping)[2,]
count_df <- data.frame(unclass(count_of_OGs_with_DEG_per_grouping))
count_df$group <- row.names(count_df)
mean_OG_size_of_all_OGs_per_grouping<- tapply(ordered_data$HOG_sizes, ordered_data$grouping, mean)
mean_df <- data.frame(unclass(mean_OG_size_of_all_OGs_per_grouping))
mean_df$group <- row.names(mean_df)

count_and_mean <- merge(count_df, mean_df, by="group")
names(count_and_mean) <- c("group", "count_of_OGs_with_DEG", "mean_OG_size")
count_and_mean$percent_OGs_with_DEG <- (count_and_mean$count_of_OGs_with_DEG/n*100)

pdf(paste0(OUTDIR, "/Jitter_plot_with_regression_of_mean_OG_sizes_of_350_groups_of_size_50_Conserved_DEG_from_both_sensitive.pdf"), 6, 4)
ggplot(count_and_mean, aes(x=mean_OG_size, y=percent_OGs_with_DEG))+
  #  geom_point(aes(x=mean_OG_size, y=percent_OGs_with_DEG), shape=21, fill=3, alpha=0.3)
  geom_jitter(shape=21, fill=1, alpha=0.3)+
  geom_smooth(method="lm", color=1)
dev.off()


# Stacked histogram Hypothesis 4 without DEG from both sensitive and Hypothesis 4 with DEG from both sensitive OG sizes
Hypo4_with_DEG_both_sens <- HOG_level_list$hypothesis_1_cons[HOG_level_list$hypothesis_1_cons$HOG %in% HOGs_all_sens_exp_DEG_both_sens,]$HOG_sizes
Hypo4_without_DEG_both_sens <- HOG_level_list$hypothesis_1_cons[HOG_level_list$hypothesis_1_cons$HOG %in% HOGs_all_sens_exp & !(HOG_level_list$hypothesis_1_cons$HOG %in% HOGs_DEG_both_sens),]$HOG_sizes
bucket <- list(Hypo4_with_DEG_both_sens, Hypo4_without_DEG_both_sens)  # this puts all values in one list
# the melt command flattens the 'bucket' list into value/vectorname pairs
# the 2 columns are called 'value' and 'L1' by default
# 'fill' will color bars differently depending on L1 group
pdf(paste0(OUTDIR, "/Histogram_stacked_OG_sizes_with_and_without_DEG_from_both_sens_in_Hypothesis_4_set.pdf"), 6, 4)
ggplot(melt(bucket), aes(value, fill = as.factor(L1))) + 
  #call geom_histogram with position="dodge" to offset the bars and manual binwidth of 2
  #  geom_histogram(aes(y = ..density..), binwidth = 1)+
  geom_histogram(binwidth = 1)+
  scale_fill_manual(name="Dataset",labels = c("with DEG from both sens", "no DEG from both sens"), values = c("1", "2"))+
  xlab("OG size")+
  xlim(c(0,30))
dev.off()

### order the data.frame of HOGs labeled with "DEG" or "no DEG" by HOG size
n <- 5
DEG_label <- rep("DEG", times=length(Hypo4_with_DEG_both_sens))
no_DEG_label <- rep("noDEG", times=length(Hypo4_without_DEG_both_sens))
data <- data.frame(HOG_sizes=c(Hypo4_with_DEG_both_sens, Hypo4_without_DEG_both_sens), label=c(DEG_label, no_DEG_label))
ordered_data <- data[order(data$HOG_sizes),]
grouping <- factor(character(length = length(ordered_data$HOG_sizes)))
grouping <- rep(seq(1,ceiling(length(grouping)/n)),each=n)
grouping <- as.factor(as.character(grouping[1:58]))

ordered_data$grouping <- grouping

count_of_OGs_with_DEG_per_grouping<- table(ordered_data$label=="DEG", grouping)[2,]
count_df <- data.frame(unclass(count_of_OGs_with_DEG_per_grouping))
count_df$group <- row.names(count_df)
mean_OG_size_of_all_OGs_per_grouping<- tapply(ordered_data$HOG_sizes, ordered_data$grouping, mean)
mean_df <- data.frame(unclass(mean_OG_size_of_all_OGs_per_grouping))
mean_df$group <- row.names(mean_df)

count_and_mean <- merge(count_df, mean_df, by="group")
names(count_and_mean) <- c("group", "count_of_OGs_with_DEG", "mean_OG_size")
count_and_mean$percent_OGs_with_DEG <- (count_and_mean$count_of_OGs_with_DEG/n*100)

pdf(paste0(OUTDIR, "/Jitter_plot_with_regression_of_mean_OG_sizes_of_12_groups_of_size_5_Hypothesis_4_DEG_from_both_sens.pdf"), 6, 4)
ggplot(count_and_mean, aes(x=mean_OG_size, y=percent_OGs_with_DEG))+
  #  geom_point(aes(x=mean_OG_size, y=percent_OGs_with_DEG), shape=21, fill=3, alpha=0.3)
  geom_jitter(shape=21, fill=1, alpha=0.3)+
  geom_smooth(method="lm", color=1)
dev.off()


## Hypothesis 2 Set

hypothesis_2_Cons_HOGs_Eutrema_exp <- HOG_level_list$hypothesis_1_cons[HOG_level_list$hypothesis_1_cons$HOG %in% HOGs_Eutrema_exp ,]
hypothesis_2_Cons_HOGs_Eutrema_exp$HOG_sizes <- apply(hypothesis_2_Cons_HOGs_Eutrema_exp[,c("Arabidopsis_lyrata_total", "Arabidopsis_thaliana_total", "Eutrema_salsugineum_total", "Brassica_napus_total")], 1, sum)
boxplot(hypothesis_2_Cons_HOGs_Eutrema_exp$HOG_sizes)
hist(hypothesis_2_Cons_HOGs_Eutrema_exp$HOG_sizes, breaks = seq(0, 50, 2), freq = F, ylim = c(0,0.4), xlab = "Sizes of OGs expanded in Eutrema", main=NA)
summary(hypothesis_2_Cons_HOGs_Eutrema_exp$HOG_sizes)


# Stacked histogram Conserved without DEG from Eutrema and Conserved with DEG from Eutrema OG sizes
Cons_with_DEG_Eutrema <- HOG_level_list$hypothesis_1_cons[HOG_level_list$hypothesis_1_cons$HOG %in% HOGs_DEG_Eutrema,]$HOG_sizes
Cons_without_DEG_Eutrema <- HOG_level_list$hypothesis_1_cons[!(HOG_level_list$hypothesis_1_cons$HOG %in% HOGs_DEG_Eutrema),]$HOG_sizes
bucket <- list(Cons_with_DEG_Eutrema, Cons_without_DEG_Eutrema)  # this puts all values in one list
# the melt command flattens the 'bucket' list into value/vectorname pairs
# the 2 columns are called 'value' and 'L1' by default
# 'fill' will color bars differently depending on L1 group
pdf(paste0(OUTDIR, "/Histogram_stacked_OG_sizes_with_and_without_DEG_from_Eutrema_in_Conserved_set.pdf"), 6, 4)
ggplot(melt(bucket), aes(value, fill = as.factor(L1))) + 
  #call geom_histogram with position="dodge" to offset the bars and manual binwidth of 2
  #  geom_histogram(aes(y = ..density..), binwidth = 2)+
  geom_histogram(binwidth = 1)+
  scale_fill_manual(name="Dataset",labels = c("with DEG from Eutrema", "no DEG from Eutrema"), values = c("1", "2"))+
  xlab("OG size")+
  xlim(c(0,30))
dev.off()

### order the data.frame of HOGs labeled with "DEG" or "no DEG" by HOG size
n <- 50
DEG_label <- rep("DEG", times=length(Cons_with_DEG_Eutrema))
no_DEG_label <- rep("noDEG", times=length(Cons_without_DEG_Eutrema))
data <- data.frame(HOG_sizes=c(Cons_with_DEG_Eutrema, Cons_without_DEG_Eutrema), label=c(DEG_label, no_DEG_label))
ordered_data <- data[order(data$HOG_sizes),]
grouping <- factor(character(length = length(ordered_data$HOG_sizes)))
grouping <- rep(seq(1,ceiling(length(grouping)/n)),each=n)
grouping <- as.factor(as.character(grouping[1:17488]))

ordered_data$grouping <- grouping

count_of_OGs_with_DEG_per_grouping<- table(ordered_data$label=="DEG", grouping)[2,]
count_df <- data.frame(unclass(count_of_OGs_with_DEG_per_grouping))
count_df$group <- row.names(count_df)
mean_OG_size_of_all_OGs_per_grouping<- tapply(ordered_data$HOG_sizes, ordered_data$grouping, mean)
mean_df <- data.frame(unclass(mean_OG_size_of_all_OGs_per_grouping))
mean_df$group <- row.names(mean_df)

count_and_mean <- merge(count_df, mean_df, by="group")
names(count_and_mean) <- c("group", "count_of_OGs_with_DEG", "mean_OG_size")
count_and_mean$percent_OGs_with_DEG <- (count_and_mean$count_of_OGs_with_DEG/n*100)

pdf(paste0(OUTDIR, "/Jitter_plot_with_regression_of_mean_OG_sizes_of_350_groups_of_size_50_Conserved_DEG_from_Eutrema.pdf"), 6, 4)
ggplot(count_and_mean, aes(x=mean_OG_size, y=percent_OGs_with_DEG))+
  #  geom_point(aes(x=mean_OG_size, y=percent_OGs_with_DEG), shape=21, fill=3, alpha=0.3)
  geom_jitter(shape=21, fill=1, alpha=0.3)+
  geom_smooth(method="lm", color=1)
dev.off()


# Stacked histogram Hypothesis_2 without DEG from Eutrema and Hypothesis_2 with DEG from Eutrema OG sizes
Hypo2_with_DEG_Eutrema <- HOG_level_list$hypothesis_1_cons[HOG_level_list$hypothesis_1_cons$HOG %in% HOGs_Eutrema_exp_DEG_Eutrema,]$HOG_sizes
Hypo2_without_DEG_Eutrema <- HOG_level_list$hypothesis_1_cons[HOG_level_list$hypothesis_1_cons$HOG %in% HOGs_Eutrema_exp & !(HOG_level_list$hypothesis_1_cons$HOG %in% HOGs_DEG_Eutrema),]$HOG_sizes
bucket <- list(Hypo2_with_DEG_Eutrema, Hypo2_without_DEG_Eutrema)  # this puts all values in one list
# the melt command flattens the 'bucket' list into value/vectorname pairs
# the 2 columns are called 'value' and 'L1' by default
# 'fill' will color bars differently depending on L1 group
pdf(paste0(OUTDIR, "/Histogram_stacked_OG_sizes_with_and_without_DEG_from_Eutrema_in_Hypothesis_2_set.pdf"), 6, 4)
ggplot(melt(bucket), aes(value, fill = as.factor(L1))) + 
  #call geom_histogram with position="dodge" to offset the bars and manual binwidth of 2
  #  geom_histogram(aes(y = ..density..), binwidth = 2)+
  geom_histogram(binwidth = 1)+
  scale_fill_manual(name="Dataset",labels = c("with DEG from Eutrema", "no DEG from Eutrema"), values = c("1", "2"))+
  xlab("OG size")+
  xlim(c(0,30))
dev.off()

### order the data.frame of HOGs labeled with "DEG" or "no DEG" by HOG size
n <- 10
DEG_label <- rep("DEG", times=length(Hypo2_with_DEG_Eutrema))
no_DEG_label <- rep("noDEG", times=length(Hypo2_without_DEG_Eutrema))
data <- data.frame(HOG_sizes=c(Hypo2_with_DEG_Eutrema, Hypo2_without_DEG_Eutrema), label=c(DEG_label, no_DEG_label))
ordered_data <- data[order(data$HOG_sizes),]
grouping <- factor(character(length = length(ordered_data$HOG_sizes)))
grouping <- rep(seq(1,ceiling(length(grouping)/n)),each=n)
grouping <- as.factor(as.character(grouping[1:727]))

ordered_data$grouping <- grouping

count_of_OGs_with_DEG_per_grouping<- table(ordered_data$label=="DEG", grouping)[2,]
count_df <- data.frame(unclass(count_of_OGs_with_DEG_per_grouping))
count_df$group <- row.names(count_df)
mean_OG_size_of_all_OGs_per_grouping<- tapply(ordered_data$HOG_sizes, ordered_data$grouping, mean)
mean_df <- data.frame(unclass(mean_OG_size_of_all_OGs_per_grouping))
mean_df$group <- row.names(mean_df)

count_and_mean <- merge(count_df, mean_df, by="group")
names(count_and_mean) <- c("group", "count_of_OGs_with_DEG", "mean_OG_size")
count_and_mean$percent_OGs_with_DEG <- (count_and_mean$count_of_OGs_with_DEG/n*100)

pdf(paste0(OUTDIR, "/Jitter_plot_with_regression_of_mean_OG_sizes_of_73_groups_of_size_10_Hypothesis_2_DEG_from_Eutrema.pdf"), 6, 4)
ggplot(count_and_mean, aes(x=mean_OG_size, y=percent_OGs_with_DEG))+
  #  geom_point(aes(x=mean_OG_size, y=percent_OGs_with_DEG), shape=21, fill=3, alpha=0.3)
  geom_jitter(shape=21, fill=1, alpha=0.3)+
  geom_smooth(method="lm", color=1)
dev.off()


# Stacked histogram Hypothesis_2 without DEG and Hypothesis_2 with DEG Any OG sizes
Hypo2_with_DEG <- hypothesis_2_Cons_HOGs_Eutrema_exp[hypothesis_2_Cons_HOGs_Eutrema_exp$HOG %in% HOGs_DEG ,]$HOG_sizes
Hypo2_without_DEG <-hypothesis_2_Cons_HOGs_Eutrema_exp[!(hypothesis_2_Cons_HOGs_Eutrema_exp$HOG %in% HOGs_DEG) ,]$HOG_sizes
bucket <- list(Hypo2_with_DEG, Hypo2_without_DEG)  # this puts all values in one list
# the melt command flattens the 'bucket' list into value/vectorname pairs
# the 2 columns are called 'value' and 'L1' by default
# 'fill' will color bars differently depending on L1 group
pdf(paste0(OUTDIR, "/Histogram_stacked_OG_sizes_with_and_without_DEG_in_Hypothesis_2_set.pdf"), 6, 4)
ggplot(melt(bucket), aes(value, fill = as.factor(L1))) + 
  #call geom_histogram with position="dodge" to offset the bars and manual binwidth of 2
  #  geom_histogram(aes(y = ..density..), binwidth = 1)+
  geom_histogram(binwidth = 1)+
  scale_fill_manual(name="Dataset",labels = c("with DEG", "no DEG"), values = c("1", "2"))+
  xlab("OG size")+
  xlim(c(0,30))
dev.off()

### order the data.frame of HOGs labeled with "DEG" or "no DEG" by HOG size
n <- 10
DEG_label <- rep("DEG", times=length(Hypo2_with_DEG))
no_DEG_label <- rep("noDEG", times=length(Hypo2_without_DEG))
data <- data.frame(HOG_sizes=c(Hypo2_with_DEG, Hypo2_without_DEG), label=c(DEG_label, no_DEG_label))
ordered_data <- data[order(data$HOG_sizes),]
grouping <- factor(character(length = length(ordered_data$HOG_sizes)))
grouping <- rep(seq(1,ceiling(length(grouping)/n)),each=n)
grouping <- as.factor(as.character(grouping[1:727]))

ordered_data$grouping <- grouping

count_of_OGs_with_DEG_per_grouping<- table(ordered_data$label=="DEG", grouping)[2,]
count_df <- data.frame(unclass(count_of_OGs_with_DEG_per_grouping))
count_df$group <- row.names(count_df)
mean_OG_size_of_all_OGs_per_grouping<- tapply(ordered_data$HOG_sizes, ordered_data$grouping, mean)
mean_df <- data.frame(unclass(mean_OG_size_of_all_OGs_per_grouping))
mean_df$group <- row.names(mean_df)

count_and_mean <- merge(count_df, mean_df, by="group")
names(count_and_mean) <- c("group", "count_of_OGs_with_DEG", "mean_OG_size")
count_and_mean$percent_OGs_with_DEG <- (count_and_mean$count_of_OGs_with_DEG/n*100)

pdf(paste0(OUTDIR, "/Jitter_plot_with_regression_of_mean_OG_sizes_of_73_groups_of_size_10_Hypothesis_2_DEG_Any.pdf"), 6, 4)
ggplot(count_and_mean, aes(x=mean_OG_size, y=percent_OGs_with_DEG))+
  #  geom_point(aes(x=mean_OG_size, y=percent_OGs_with_DEG), shape=21, fill=3, alpha=0.3)
  geom_jitter(shape=21, fill=1, alpha=0.3)+
  geom_smooth(method="lm", color=1)
dev.off()


## Hypothesis 3 Set

hypothesis_3_Cons_HOGs_Alyrata_exp <- HOG_level_list$hypothesis_1_cons[HOG_level_list$hypothesis_1_cons$HOG %in% HOGs_Alyrata_exp ,]
hypothesis_3_Cons_HOGs_Alyrata_exp$HOG_sizes <- apply(hypothesis_3_Cons_HOGs_Alyrata_exp[,c("Arabidopsis_lyrata_total", "Arabidopsis_thaliana_total", "Eutrema_salsugineum_total", "Brassica_napus_total")], 1, sum)
boxplot(hypothesis_3_Cons_HOGs_Alyrata_exp$HOG_sizes)
hist(hypothesis_3_Cons_HOGs_Alyrata_exp$HOG_sizes, breaks = seq(0, 50, 2), freq = F, ylim = c(0,0.4), xlab = "Sizes of OGs expanded in Alyrata", main=NA)
summary(hypothesis_3_Cons_HOGs_Alyrata_exp$HOG_sizes)


# Stacked histogram Conserved without DEG from Alyrata and Conserved with DEG from Alyrata OG sizes
Cons_with_DEG_Alyrata <- HOG_level_list$hypothesis_1_cons[HOG_level_list$hypothesis_1_cons$HOG %in% HOGs_DEG_Alyrata,]$HOG_sizes
Cons_without_DEG_Alyrata <- HOG_level_list$hypothesis_1_cons[!(HOG_level_list$hypothesis_1_cons$HOG %in% HOGs_DEG_Alyrata),]$HOG_sizes
bucket <- list(Cons_with_DEG_Alyrata, Cons_without_DEG_Alyrata)  # this puts all values in one list
# the melt command flattens the 'bucket' list into value/vectorname pairs
# the 2 columns are called 'value' and 'L1' by default
# 'fill' will color bars differently depending on L1 group
pdf(paste0(OUTDIR, "/Histogram_stacked_OG_sizes_with_and_without_DEG_from_Alyrata_in_Conserved_set.pdf"), 6, 4)
ggplot(melt(bucket), aes(value, fill = as.factor(L1))) + 
  #call geom_histogram with position="dodge" to offset the bars and manual binwidth of 2
  #  geom_histogram(aes(y = ..density..), binwidth = 2)+
  geom_histogram(binwidth = 1)+
  scale_fill_manual(name="Dataset",labels = c("with DEG from Alyrata", "no DEG from Alyrata"), values = c("1", "2"))+
  xlab("OG size")+
  xlim(c(0,30))
dev.off()

### order the data.frame of HOGs labeled with "DEG" or "no DEG" by HOG size
n <- 50
DEG_label <- rep("DEG", times=length(Cons_with_DEG_Alyrata))
no_DEG_label <- rep("noDEG", times=length(Cons_without_DEG_Alyrata))
data <- data.frame(HOG_sizes=c(Cons_with_DEG_Alyrata, Cons_without_DEG_Alyrata), label=c(DEG_label, no_DEG_label))
ordered_data <- data[order(data$HOG_sizes),]
grouping <- factor(character(length = length(ordered_data$HOG_sizes)))
grouping <- rep(seq(1,ceiling(length(grouping)/n)),each=n)
grouping <- as.factor(as.character(grouping[1:17488]))

ordered_data$grouping <- grouping

count_of_OGs_with_DEG_per_grouping<- table(ordered_data$label=="DEG", grouping)[2,]
count_df <- data.frame(unclass(count_of_OGs_with_DEG_per_grouping))
count_df$group <- row.names(count_df)
mean_OG_size_of_all_OGs_per_grouping<- tapply(ordered_data$HOG_sizes, ordered_data$grouping, mean)
mean_df <- data.frame(unclass(mean_OG_size_of_all_OGs_per_grouping))
mean_df$group <- row.names(mean_df)

count_and_mean <- merge(count_df, mean_df, by="group")
names(count_and_mean) <- c("group", "count_of_OGs_with_DEG", "mean_OG_size")
count_and_mean$percent_OGs_with_DEG <- (count_and_mean$count_of_OGs_with_DEG/n*100)

pdf(paste0(OUTDIR, "/Jitter_plot_with_regression_of_mean_OG_sizes_of_350_groups_of_size_50_Conserved_DEG_from_Alyrata.pdf"), 6, 4)
ggplot(count_and_mean, aes(x=mean_OG_size, y=percent_OGs_with_DEG))+
  #  geom_point(aes(x=mean_OG_size, y=percent_OGs_with_DEG), shape=21, fill=3, alpha=0.3)
  geom_jitter(shape=21, fill=1, alpha=0.3)+
  geom_smooth(method="lm", color=1)
dev.off()


# Stacked histogram Hypothesis_3 without DEG from Alyrata and Hypothesis_3 with DEG from Alyrata OG sizes
Hypo3_with_DEG_Alyrata <- HOG_level_list$hypothesis_1_cons[HOG_level_list$hypothesis_1_cons$HOG %in% HOGs_Alyrata_exp_DEG_Alyrata,]$HOG_sizes
Hypo3_without_DEG_Alyrata <- HOG_level_list$hypothesis_1_cons[HOG_level_list$hypothesis_1_cons$HOG %in% HOGs_Alyrata_exp & !(HOG_level_list$hypothesis_1_cons$HOG %in% HOGs_DEG_Alyrata),]$HOG_sizes
bucket <- list(Hypo3_with_DEG_Alyrata, Hypo3_without_DEG_Alyrata)  # this puts all values in one list
# the melt command flattens the 'bucket' list into value/vectorname pairs
# the 2 columns are called 'value' and 'L1' by default
# 'fill' will color bars differently depending on L1 group
pdf(paste0(OUTDIR, "/Histogram_stacked_OG_sizes_with_and_without_DEG_from_Alyrata_in_Hypothesis_3_set.pdf"), 6, 4)
ggplot(melt(bucket), aes(value, fill = as.factor(L1))) + 
  #call geom_histogram with position="dodge" to offset the bars and manual binwidth of 2
  #  geom_histogram(aes(y = ..density..), binwidth = 2)+
  geom_histogram(binwidth = 1)+
  scale_fill_manual(name="Dataset",labels = c("with DEG from Alyrata", "no DEG from Alyrata"), values = c("1", "2"))+
  xlab("OG size")+
  xlim(c(0,30))
dev.off()

### order the data.frame of HOGs labeled with "DEG" or "no DEG" by HOG size
n <- 10
DEG_label <- rep("DEG", times=length(Hypo3_with_DEG_Alyrata))
no_DEG_label <- rep("noDEG", times=length(Hypo3_without_DEG_Alyrata))
data <- data.frame(HOG_sizes=c(Hypo3_with_DEG_Alyrata, Hypo3_without_DEG_Alyrata), label=c(DEG_label, no_DEG_label))
ordered_data <- data[order(data$HOG_sizes),]
grouping <- factor(character(length = length(ordered_data$HOG_sizes)))
grouping <- rep(seq(1,ceiling(length(grouping)/n)),each=n)
grouping <- as.factor(as.character(grouping[1:1075]))

ordered_data$grouping <- grouping

count_of_OGs_with_DEG_per_grouping<- table(ordered_data$label=="DEG", grouping)[2,]
count_df <- data.frame(unclass(count_of_OGs_with_DEG_per_grouping))
count_df$group <- row.names(count_df)
mean_OG_size_of_all_OGs_per_grouping<- tapply(ordered_data$HOG_sizes, ordered_data$grouping, mean)
mean_df <- data.frame(unclass(mean_OG_size_of_all_OGs_per_grouping))
mean_df$group <- row.names(mean_df)

count_and_mean <- merge(count_df, mean_df, by="group")
names(count_and_mean) <- c("group", "count_of_OGs_with_DEG", "mean_OG_size")
count_and_mean$percent_OGs_with_DEG <- (count_and_mean$count_of_OGs_with_DEG/n*100)

pdf(paste0(OUTDIR, "/Jitter_plot_with_regression_of_mean_OG_sizes_of_108_groups_of_size_10_Hypothesis_3_DEG_from_Alyrata.pdf"), 6, 4)
ggplot(count_and_mean, aes(x=mean_OG_size, y=percent_OGs_with_DEG))+
  #  geom_point(aes(x=mean_OG_size, y=percent_OGs_with_DEG), shape=21, fill=3, alpha=0.3)
  geom_jitter(shape=21, fill=1, alpha=0.3)+
  geom_smooth(method="lm", color=1)
dev.off()


# Stacked histogram Hypothesis_3 without DEG and Hypothesis_3 with DEG Any OG sizes
Hypo3_with_DEG <- hypothesis_3_Cons_HOGs_Alyrata_exp[hypothesis_3_Cons_HOGs_Alyrata_exp$HOG %in% HOGs_DEG ,]$HOG_sizes
Hypo3_without_DEG <-hypothesis_3_Cons_HOGs_Alyrata_exp[!(hypothesis_3_Cons_HOGs_Alyrata_exp$HOG %in% HOGs_DEG) ,]$HOG_sizes
bucket <- list(Hypo3_with_DEG, Hypo3_without_DEG)  # this puts all values in one list
# the melt command flattens the 'bucket' list into value/vectorname pairs
# the 2 columns are called 'value' and 'L1' by default
# 'fill' will color bars differently depending on L1 group
pdf(paste0(OUTDIR, "/Histogram_stacked_OG_sizes_with_and_without_DEG_in_Hypothesis_3_set.pdf"), 6, 4)
ggplot(melt(bucket), aes(value, fill = as.factor(L1))) + 
  #call geom_histogram with position="dodge" to offset the bars and manual binwidth of 2
  #  geom_histogram(aes(y = ..density..), binwidth = 1)+
  geom_histogram(binwidth = 1)+
  scale_fill_manual(name="Dataset",labels = c("with DEG", "no DEG"), values = c("1", "2"))+
  xlab("OG size")+
  xlim(c(0,30))
dev.off()


### order the data.frame of HOGs labeled with "DEG" or "no DEG" by HOG size
n <- 10
DEG_label <- rep("DEG", times=length(Hypo3_with_DEG))
no_DEG_label <- rep("noDEG", times=length(Hypo3_without_DEG))
data <- data.frame(HOG_sizes=c(Hypo3_with_DEG, Hypo3_without_DEG), label=c(DEG_label, no_DEG_label))
ordered_data <- data[order(data$HOG_sizes),]
grouping <- factor(character(length = length(ordered_data$HOG_sizes)))
grouping <- rep(seq(1,ceiling(length(grouping)/n)),each=n)
grouping <- as.factor(as.character(grouping[1:1075]))

ordered_data$grouping <- grouping

count_of_OGs_with_DEG_per_grouping<- table(ordered_data$label=="DEG", grouping)[2,]
count_df <- data.frame(unclass(count_of_OGs_with_DEG_per_grouping))
count_df$group <- row.names(count_df)
mean_OG_size_of_all_OGs_per_grouping<- tapply(ordered_data$HOG_sizes, ordered_data$grouping, mean)
mean_df <- data.frame(unclass(mean_OG_size_of_all_OGs_per_grouping))
mean_df$group <- row.names(mean_df)

count_and_mean <- merge(count_df, mean_df, by="group")
names(count_and_mean) <- c("group", "count_of_OGs_with_DEG", "mean_OG_size")
count_and_mean$percent_OGs_with_DEG <- (count_and_mean$count_of_OGs_with_DEG/n*100)

pdf(paste0(OUTDIR, "/Jitter_plot_with_regression_of_mean_OG_sizes_of_108_groups_of_size_10_Hypothesis_3_DEG_Any.pdf"), 6, 4)
ggplot(count_and_mean, aes(x=mean_OG_size, y=percent_OGs_with_DEG))+
  #  geom_point(aes(x=mean_OG_size, y=percent_OGs_with_DEG), shape=21, fill=3, alpha=0.3)
  geom_jitter(shape=21, fill=1, alpha=0.3)+
  geom_smooth(method="lm", color=1)
dev.off()


## Hypothesis 5 Set


hypothesis_5_Cons_HOGs_Athaliana_exp <- HOG_level_list$hypothesis_1_cons[HOG_level_list$hypothesis_1_cons$HOG %in% HOGs_Athaliana_exp ,]
hypothesis_5_Cons_HOGs_Athaliana_exp$HOG_sizes <- apply(hypothesis_5_Cons_HOGs_Athaliana_exp[,c("Arabidopsis_lyrata_total", "Arabidopsis_thaliana_total", "Eutrema_salsugineum_total", "Brassica_napus_total")], 1, sum)
boxplot(hypothesis_5_Cons_HOGs_Athaliana_exp$HOG_sizes)
hist(hypothesis_5_Cons_HOGs_Athaliana_exp$HOG_sizes, breaks = seq(0, 50, 2), freq = F, ylim = c(0,0.4), xlab = "Sizes of OGs expanded in Athaliana", main=NA)
summary(hypothesis_5_Cons_HOGs_Athaliana_exp$HOG_sizes)


# Stacked histogram Conserved without DEG from Athaliana and Conserved with DEG from Athaliana OG sizes
Cons_with_DEG_Athaliana <- HOG_level_list$hypothesis_1_cons[HOG_level_list$hypothesis_1_cons$HOG %in% HOGs_DEG_Athaliana,]$HOG_sizes
Cons_without_DEG_Athaliana <- HOG_level_list$hypothesis_1_cons[!(HOG_level_list$hypothesis_1_cons$HOG %in% HOGs_DEG_Athaliana),]$HOG_sizes
bucket <- list(Cons_with_DEG_Athaliana, Cons_without_DEG_Athaliana)  # this puts all values in one list
# the melt command flattens the 'bucket' list into value/vectorname pairs
# the 2 columns are called 'value' and 'L1' by default
# 'fill' will color bars differently depending on L1 group
pdf(paste0(OUTDIR, "/Histogram_stacked_OG_sizes_with_and_without_DEG_from_Athaliana_in_Conserved_set.pdf"), 6, 4)
ggplot(melt(bucket), aes(value, fill = as.factor(L1))) + 
  #call geom_histogram with position="dodge" to offset the bars and manual binwidth of 2
  #  geom_histogram(aes(y = ..density..), binwidth = 2)+
  geom_histogram(binwidth = 1)+
  scale_fill_manual(name="Dataset",labels = c("with DEG from Athliana", "no DEG from Athaliana"), values = c("1", "2"))+
  xlab("OG size")+
  xlim(c(0,30))
dev.off()

### order the data.frame of HOGs labeled with "DEG" or "no DEG" by HOG size
n <- 50
DEG_label <- rep("DEG", times=length(Cons_with_DEG_Athaliana))
no_DEG_label <- rep("noDEG", times=length(Cons_without_DEG_Athaliana))
data <- data.frame(HOG_sizes=c(Cons_with_DEG_Athaliana, Cons_without_DEG_Athaliana), label=c(DEG_label, no_DEG_label))
ordered_data <- data[order(data$HOG_sizes),]
grouping <- factor(character(length = length(ordered_data$HOG_sizes)))
grouping <- rep(seq(1,ceiling(length(grouping)/n)),each=n)
grouping <- as.factor(as.character(grouping[1:17488]))

ordered_data$grouping <- grouping

count_of_OGs_with_DEG_per_grouping<- table(ordered_data$label=="DEG", grouping)[2,]
count_df <- data.frame(unclass(count_of_OGs_with_DEG_per_grouping))
count_df$group <- row.names(count_df)
mean_OG_size_of_all_OGs_per_grouping<- tapply(ordered_data$HOG_sizes, ordered_data$grouping, mean)
mean_df <- data.frame(unclass(mean_OG_size_of_all_OGs_per_grouping))
mean_df$group <- row.names(mean_df)

count_and_mean <- merge(count_df, mean_df, by="group")
names(count_and_mean) <- c("group", "count_of_OGs_with_DEG", "mean_OG_size")
count_and_mean$percent_OGs_with_DEG <- (count_and_mean$count_of_OGs_with_DEG/n*100)

pdf(paste0(OUTDIR, "/Jitter_plot_with_regression_of_mean_OG_sizes_of_350_groups_of_size_50_Conserved_DEG_from_Athaliana.pdf"), 6, 4)
ggplot(count_and_mean, aes(x=mean_OG_size, y=percent_OGs_with_DEG))+
  #  geom_point(aes(x=mean_OG_size, y=percent_OGs_with_DEG), shape=21, fill=3, alpha=0.3)
  geom_jitter(shape=21, fill=1, alpha=0.3)+
  geom_smooth(method="lm", color=1)
dev.off()


# Stacked histogram Hypothesis_5 without DEG from Athaliana and Hypothesis_5 with DEG from Athaliana OG sizes
Hypo5_with_DEG_Athaliana <- HOG_level_list$hypothesis_1_cons[HOG_level_list$hypothesis_1_cons$HOG %in% HOGs_Athaliana_exp_DEG_Athaliana,]$HOG_sizes
Hypo5_without_DEG_Athaliana <- HOG_level_list$hypothesis_1_cons[HOG_level_list$hypothesis_1_cons$HOG %in% HOGs_Athaliana_exp & !(HOG_level_list$hypothesis_1_cons$HOG %in% HOGs_DEG_Athaliana),]$HOG_sizes
bucket <- list(Hypo5_with_DEG_Athaliana, Hypo5_without_DEG_Athaliana)  # this puts all values in one list
# the melt command flattens the 'bucket' list into value/vectorname pairs
# the 2 columns are called 'value' and 'L1' by default
# 'fill' will color bars differently depending on L1 group
pdf(paste0(OUTDIR, "/Histogram_stacked_OG_sizes_with_and_without_DEG_from_Athaliana_in_Hypothesis_5_set.pdf"), 6, 4)
ggplot(melt(bucket), aes(value, fill = as.factor(L1))) + 
  #call geom_histogram with position="dodge" to offset the bars and manual binwidth of 2
  #  geom_histogram(aes(y = ..density..), binwidth = 2)+
  geom_histogram(binwidth = 1)+
  scale_fill_manual(name="Dataset",labels = c("with DEG from Athaliana", "no DEG from Athaliana"), values = c("1", "2"))+
  xlab("OG size")+
  xlim(c(0,30))
dev.off()

### order the data.frame of HOGs labeled with "DEG" or "no DEG" by HOG size
n <- 10
DEG_label <- rep("DEG", times=length(Hypo5_with_DEG_Athaliana))
no_DEG_label <- rep("noDEG", times=length(Hypo5_without_DEG_Athaliana))
data <- data.frame(HOG_sizes=c(Hypo5_with_DEG_Athaliana, Hypo5_without_DEG_Athaliana), label=c(DEG_label, no_DEG_label))
ordered_data <- data[order(data$HOG_sizes),]
grouping <- factor(character(length = length(ordered_data$HOG_sizes)))
grouping <- rep(seq(1,ceiling(length(grouping)/n)),each=n)
grouping <- as.factor(as.character(grouping[1:375]))

ordered_data$grouping <- grouping

count_of_OGs_with_DEG_per_grouping<- table(ordered_data$label=="DEG", grouping)[2,]
count_df <- data.frame(unclass(count_of_OGs_with_DEG_per_grouping))
count_df$group <- row.names(count_df)
mean_OG_size_of_all_OGs_per_grouping<- tapply(ordered_data$HOG_sizes, ordered_data$grouping, mean)
mean_df <- data.frame(unclass(mean_OG_size_of_all_OGs_per_grouping))
mean_df$group <- row.names(mean_df)

count_and_mean <- merge(count_df, mean_df, by="group")
names(count_and_mean) <- c("group", "count_of_OGs_with_DEG", "mean_OG_size")
count_and_mean$percent_OGs_with_DEG <- (count_and_mean$count_of_OGs_with_DEG/n*100)

pdf(paste0(OUTDIR, "/Jitter_plot_with_regression_of_mean_OG_sizes_of_38_groups_of_size_10_Hypothesis_5_DEG_from_Athaliana.pdf"), 6, 4)
ggplot(count_and_mean, aes(x=mean_OG_size, y=percent_OGs_with_DEG))+
  #  geom_point(aes(x=mean_OG_size, y=percent_OGs_with_DEG), shape=21, fill=3, alpha=0.3)
  geom_jitter(shape=21, fill=1, alpha=0.3)+
  geom_smooth(method="lm", color=1)
dev.off()


# Stacked histogram Hypothesis_5 without DEG and Hypothesis_5 with DEG Any OG sizes
Hypo5_with_DEG <- hypothesis_5_Cons_HOGs_Athaliana_exp[hypothesis_5_Cons_HOGs_Athaliana_exp$HOG %in% HOGs_DEG ,]$HOG_sizes
Hypo5_without_DEG <-hypothesis_5_Cons_HOGs_Athaliana_exp[!(hypothesis_5_Cons_HOGs_Athaliana_exp$HOG %in% HOGs_DEG) ,]$HOG_sizes
bucket <- list(Hypo5_with_DEG, Hypo5_without_DEG)  # this puts all values in one list
# the melt command flattens the 'bucket' list into value/vectorname pairs
# the 2 columns are called 'value' and 'L1' by default
# 'fill' will color bars differently depending on L1 group
pdf(paste0(OUTDIR, "/Histogram_stacked_OG_sizes_with_and_without_DEG_in_Hypothesis_5_set.pdf"), 6, 4)
ggplot(melt(bucket), aes(value, fill = as.factor(L1))) + 
  #call geom_histogram with position="dodge" to offset the bars and manual binwidth of 2
  #  geom_histogram(aes(y = ..density..), binwidth = 1)+
  geom_histogram(binwidth = 1)+
  scale_fill_manual(name="Dataset",labels = c("with DEG", "no DEG"), values = c("1", "2"))+
  xlab("OG size")+
  xlim(c(0,30))
dev.off()

### order the data.frame of HOGs labeled with "DEG" or "no DEG" by HOG size
n <- 10
DEG_label <- rep("DEG", times=length(Hypo5_with_DEG))
no_DEG_label <- rep("noDEG", times=length(Hypo5_without_DEG))
data <- data.frame(HOG_sizes=c(Hypo5_with_DEG, Hypo5_without_DEG), label=c(DEG_label, no_DEG_label))
ordered_data <- data[order(data$HOG_sizes),]
grouping <- factor(character(length = length(ordered_data$HOG_sizes)))
grouping <- rep(seq(1,ceiling(length(grouping)/n)),each=n)
grouping <- as.factor(as.character(grouping[1:375]))

ordered_data$grouping <- grouping

count_of_OGs_with_DEG_per_grouping<- table(ordered_data$label=="DEG", grouping)[2,]
count_df <- data.frame(unclass(count_of_OGs_with_DEG_per_grouping))
count_df$group <- row.names(count_df)
mean_OG_size_of_all_OGs_per_grouping<- tapply(ordered_data$HOG_sizes, ordered_data$grouping, mean)
mean_df <- data.frame(unclass(mean_OG_size_of_all_OGs_per_grouping))
mean_df$group <- row.names(mean_df)

count_and_mean <- merge(count_df, mean_df, by="group")
names(count_and_mean) <- c("group", "count_of_OGs_with_DEG", "mean_OG_size")
count_and_mean$percent_OGs_with_DEG <- (count_and_mean$count_of_OGs_with_DEG/n*100)

pdf(paste0(OUTDIR, "/Jitter_plot_with_regression_of_mean_OG_sizes_of_38_groups_of_size_10_Hypothesis_5_DEG_Any.pdf"), 6, 4)
ggplot(count_and_mean, aes(x=mean_OG_size, y=percent_OGs_with_DEG))+
  #  geom_point(aes(x=mean_OG_size, y=percent_OGs_with_DEG), shape=21, fill=3, alpha=0.3)
  geom_jitter(shape=21, fill=1, alpha=0.3)+
  geom_smooth(method="lm", color=1)
dev.off()


## Hypothesis 6 Set

hypothesis_6_Cons_HOGs_Bnapus_exp <- HOG_level_list$hypothesis_1_cons[HOG_level_list$hypothesis_1_cons$HOG %in% HOGs_Bnapus_exp ,]
hypothesis_6_Cons_HOGs_Bnapus_exp$HOG_sizes <- apply(hypothesis_6_Cons_HOGs_Bnapus_exp[,c("Arabidopsis_lyrata_total", "Arabidopsis_thaliana_total", "Eutrema_salsugineum_total", "Brassica_napus_total")], 1, sum)
boxplot(hypothesis_6_Cons_HOGs_Bnapus_exp$HOG_sizes)
hist(hypothesis_6_Cons_HOGs_Bnapus_exp$HOG_sizes, breaks = seq(0, 60, 2), freq = F, ylim = c(0,0.4), xlab = "Sizes of OGs expanded in Bnapus", main=NA)
summary(hypothesis_6_Cons_HOGs_Bnapus_exp$HOG_sizes)


# Stacked histogram Conserved without DEG from Bnapus and Conserved with DEG from Bnapus OG sizes
Cons_with_DEG_Bnapus <- HOG_level_list$hypothesis_1_cons[HOG_level_list$hypothesis_1_cons$HOG %in% HOGs_DEG_Bnapus,]$HOG_sizes
Cons_without_DEG_Bnapus <- HOG_level_list$hypothesis_1_cons[!(HOG_level_list$hypothesis_1_cons$HOG %in% HOGs_DEG_Bnapus),]$HOG_sizes
bucket <- list(Cons_with_DEG_Bnapus, Cons_without_DEG_Bnapus)  # this puts all values in one list
# the melt command flattens the 'bucket' list into value/vectorname pairs
# the 2 columns are called 'value' and 'L1' by default
# 'fill' will color bars differently depending on L1 group
pdf(paste0(OUTDIR, "/Histogram_stacked_OG_sizes_with_and_without_DEG_from_Bnapus_in_Conserved_set.pdf"), 6, 4)
ggplot(melt(bucket), aes(value, fill = as.factor(L1))) + 
  #call geom_histogram with position="dodge" to offset the bars and manual binwidth of 2
  #  geom_histogram(aes(y = ..density..), binwidth = 2)+
  geom_histogram(binwidth = 1)+
  scale_fill_manual(name="Dataset",labels = c("with DEG from Bnapus", "no DEG from Bnapus"), values = c("1", "2"))+
  xlab("OG size")+
  xlim(c(0,30))
dev.off()

### order the data.frame of HOGs labeled with "DEG" or "no DEG" by HOG size
n <- 50
DEG_label <- rep("DEG", times=length(Cons_with_DEG_Bnapus))
no_DEG_label <- rep("noDEG", times=length(Cons_without_DEG_Bnapus))
data <- data.frame(HOG_sizes=c(Cons_with_DEG_Bnapus, Cons_without_DEG_Bnapus), label=c(DEG_label, no_DEG_label))
ordered_data <- data[order(data$HOG_sizes),]
grouping <- factor(character(length = length(ordered_data$HOG_sizes)))
grouping <- rep(seq(1,ceiling(length(grouping)/n)),each=n)
grouping <- as.factor(as.character(grouping[1:17488]))

ordered_data$grouping <- grouping

count_of_OGs_with_DEG_per_grouping<- table(ordered_data$label=="DEG", grouping)[2,]
count_df <- data.frame(unclass(count_of_OGs_with_DEG_per_grouping))
count_df$group <- row.names(count_df)
mean_OG_size_of_all_OGs_per_grouping<- tapply(ordered_data$HOG_sizes, ordered_data$grouping, mean)
mean_df <- data.frame(unclass(mean_OG_size_of_all_OGs_per_grouping))
mean_df$group <- row.names(mean_df)

count_and_mean <- merge(count_df, mean_df, by="group")
names(count_and_mean) <- c("group", "count_of_OGs_with_DEG", "mean_OG_size")
count_and_mean$percent_OGs_with_DEG <- (count_and_mean$count_of_OGs_with_DEG/n*100)

pdf(paste0(OUTDIR, "/Jitter_plot_with_regression_of_mean_OG_sizes_of_350_groups_of_size_50_Conserved_DEG_from_Bnapus.pdf"), 6, 4)
ggplot(count_and_mean, aes(x=mean_OG_size, y=percent_OGs_with_DEG))+
  #  geom_point(aes(x=mean_OG_size, y=percent_OGs_with_DEG), shape=21, fill=3, alpha=0.3)
  geom_jitter(shape=21, fill=1, alpha=0.3)+
  geom_smooth(method="lm", color=1)
dev.off()


# Stacked histogram Hypothesis_6 without DEG from Bnapus and Hypothesis_6 with DEG from Bnapus OG sizes
Hypo6_with_DEG_Bnapus <- HOG_level_list$hypothesis_1_cons[HOG_level_list$hypothesis_1_cons$HOG %in% HOGs_Bnapus_exp_DEG_Bnapus,]$HOG_sizes
Hypo6_without_DEG_Bnapus <- HOG_level_list$hypothesis_1_cons[HOG_level_list$hypothesis_1_cons$HOG %in% HOGs_Bnapus_exp & !(HOG_level_list$hypothesis_1_cons$HOG %in% HOGs_DEG_Bnapus),]$HOG_sizes
bucket <- list(Hypo6_with_DEG_Bnapus, Hypo6_without_DEG_Bnapus)  # this puts all values in one list
# the melt command flattens the 'bucket' list into value/vectorname pairs
# the 2 columns are called 'value' and 'L1' by default
# 'fill' will color bars differently depending on L1 group
pdf(paste0(OUTDIR, "/Histogram_stacked_OG_sizes_with_and_without_DEG_from_Bnapus_in_Hypothesis_6_set.pdf"), 6, 4)
ggplot(melt(bucket), aes(value, fill = as.factor(L1))) + 
  #call geom_histogram with position="dodge" to offset the bars and manual binwidth of 2
  #  geom_histogram(aes(y = ..density..), binwidth = 2)+
  geom_histogram(binwidth = 1)+
  scale_fill_manual(name="Dataset",labels = c("with DEG from Bnapus", "no DEG from Bnapus"), values = c("1", "2"))+
  xlab("OG size")+
  xlim(c(0,30))
dev.off()

### order the data.frame of HOGs labeled with "DEG" or "no DEG" by HOG size
n <- 15
DEG_label <- rep("DEG", times=length(Hypo6_with_DEG_Bnapus))
no_DEG_label <- rep("noDEG", times=length(Hypo6_without_DEG_Bnapus))
data <- data.frame(HOG_sizes=c(Hypo6_with_DEG_Bnapus, Hypo6_without_DEG_Bnapus), label=c(DEG_label, no_DEG_label))
ordered_data <- data[order(data$HOG_sizes),]
grouping <- factor(character(length = length(ordered_data$HOG_sizes)))
grouping <- rep(seq(1,ceiling(length(grouping)/n)),each=n)
grouping <- as.factor(as.character(grouping[1:1825]))

ordered_data$grouping <- grouping

count_of_OGs_with_DEG_per_grouping<- table(ordered_data$label=="DEG", grouping)[2,]
count_df <- data.frame(unclass(count_of_OGs_with_DEG_per_grouping))
count_df$group <- row.names(count_df)
mean_OG_size_of_all_OGs_per_grouping<- tapply(ordered_data$HOG_sizes, ordered_data$grouping, mean)
mean_df <- data.frame(unclass(mean_OG_size_of_all_OGs_per_grouping))
mean_df$group <- row.names(mean_df)

count_and_mean <- merge(count_df, mean_df, by="group")
names(count_and_mean) <- c("group", "count_of_OGs_with_DEG", "mean_OG_size")
count_and_mean$percent_OGs_with_DEG <- (count_and_mean$count_of_OGs_with_DEG/n*100)

pdf(paste0(OUTDIR, "/Jitter_plot_with_regression_of_mean_OG_sizes_of_122_groups_of_size_15_Hypothesis_6_DEG_from_Bnapus.pdf"), 6, 4)
ggplot(count_and_mean, aes(x=mean_OG_size, y=percent_OGs_with_DEG))+
  #  geom_point(aes(x=mean_OG_size, y=percent_OGs_with_DEG), shape=21, fill=3, alpha=0.3)
  geom_jitter(shape=21, fill=1, alpha=0.3)+
  geom_smooth(method="lm", color=1)
dev.off()


# Stacked histogram Hypothesis_6 without DEG and Hypothesis_6 with DEG Any OG sizes
Hypo6_with_DEG <- hypothesis_6_Cons_HOGs_Bnapus_exp[hypothesis_6_Cons_HOGs_Bnapus_exp$HOG %in% HOGs_DEG ,]$HOG_sizes
Hypo6_without_DEG <-hypothesis_6_Cons_HOGs_Bnapus_exp[!(hypothesis_6_Cons_HOGs_Bnapus_exp$HOG %in% HOGs_DEG) ,]$HOG_sizes
bucket <- list(Hypo6_with_DEG, Hypo6_without_DEG)  # this puts all values in one list
# the melt command flattens the 'bucket' list into value/vectorname pairs
# the 2 columns are called 'value' and 'L1' by default
# 'fill' will color bars differently depending on L1 group
pdf(paste0(OUTDIR, "/Histogram_stacked_OG_sizes_with_and_without_DEG_in_Hypothesis_6_set.pdf"), 6, 4)
ggplot(melt(bucket), aes(value, fill = as.factor(L1))) + 
  #call geom_histogram with position="dodge" to offset the bars and manual binwidth of 2
  #  geom_histogram(aes(y = ..density..), binwidth = 1)+
  geom_histogram(binwidth = 1)+
  scale_fill_manual(name="Dataset",labels = c("with DEG", "no DEG"), values = c("1", "2"))+
  xlab("OG size")+
  xlim(c(0,30))
dev.off()


### order the data.frame of HOGs labeled with "DEG" or "no DEG" by HOG size
n <- 15
DEG_label <- rep("DEG", times=length(Hypo6_with_DEG))
no_DEG_label <- rep("noDEG", times=length(Hypo6_without_DEG))
data <- data.frame(HOG_sizes=c(Hypo6_with_DEG, Hypo6_without_DEG), label=c(DEG_label, no_DEG_label))
ordered_data <- data[order(data$HOG_sizes),]
grouping <- factor(character(length = length(ordered_data$HOG_sizes)))
grouping <- rep(seq(1,ceiling(length(grouping)/n)),each=n)
grouping <- as.factor(as.character(grouping[1:1825]))

ordered_data$grouping <- grouping

count_of_OGs_with_DEG_per_grouping<- table(ordered_data$label=="DEG", grouping)[2,]
count_df <- data.frame(unclass(count_of_OGs_with_DEG_per_grouping))
count_df$group <- row.names(count_df)
mean_OG_size_of_all_OGs_per_grouping<- tapply(ordered_data$HOG_sizes, ordered_data$grouping, mean)
mean_df <- data.frame(unclass(mean_OG_size_of_all_OGs_per_grouping))
mean_df$group <- row.names(mean_df)

count_and_mean <- merge(count_df, mean_df, by="group")
names(count_and_mean) <- c("group", "count_of_OGs_with_DEG", "mean_OG_size")
count_and_mean$percent_OGs_with_DEG <- (count_and_mean$count_of_OGs_with_DEG/n*100)

pdf(paste0(OUTDIR, "/Jitter_plot_with_regression_of_mean_OG_sizes_of_122_groups_of_size_15_Hypothesis_6_DEG_Any.pdf"), 6, 4)
ggplot(count_and_mean, aes(x=mean_OG_size, y=percent_OGs_with_DEG))+
  #  geom_point(aes(x=mean_OG_size, y=percent_OGs_with_DEG), shape=21, fill=3, alpha=0.3)
  geom_jitter(shape=21, fill=1, alpha=0.3)+
  geom_smooth(method="lm", color=1)
dev.off()

### UpSetR plot for All OGs
library("UpSetR")
Eutrema <- HOG_level_list$hypothesis_1[(HOG_level_list$hypothesis_1$Eutrema_salsugineum_total>=1) & (!is.na(HOG_level_list$hypothesis_1$Eutrema_salsugineum_total)),]$HOG
Alyrata <- HOG_level_list$hypothesis_1[(HOG_level_list$hypothesis_1$Arabidopsis_lyrata_total>=1) & (!is.na(HOG_level_list$hypothesis_1$Arabidopsis_lyrata_total)),]$HOG
Athaliana <-  HOG_level_list$hypothesis_1[(HOG_level_list$hypothesis_1$Arabidopsis_thaliana_total>=1) & (!is.na(HOG_level_list$hypothesis_1$Arabidopsis_thaliana_total)),]$HOG
Bnapus <-  HOG_level_list$hypothesis_1[(HOG_level_list$hypothesis_1$Brassica_napus_total>=1) & (!is.na(HOG_level_list$hypothesis_1$Brassica_napus_total)),]$HOG

listInput <- list(Eutrema=Eutrema,Alyrata=Alyrata,Athaliana=Athaliana,Bnapus=Bnapus)
pdf(paste0(OUTDIR, "/Intersection_of_OGs_total_set.pdf"), 6, 4)
upset(fromList(listInput), nsets=4, order.by = "freq", sets.x.label = "Number of OGs", mainbar.y.label = "Intersection of OGs",
      number.angles = 0, 
      point.size = 4, line.size = 1.5,
      text.scale = c(1, 1, 1, 1, 1.5, 1.5)
)
dev.off()

### UpSetR plot for expanded OGs
library("UpSetR")
Exp_Eutrema <- HOG_level_list$hypothesis_2_cons[HOG_level_list$hypothesis_2_cons$expansion == "yes",]$HOG
Exp_Alyrata <- HOG_level_list$hypothesis_3_cons[HOG_level_list$hypothesis_3_cons$expansion == "yes",]$HOG
Exp_Athaliana <- HOG_level_list$hypothesis_5_cons[HOG_level_list$hypothesis_5_cons$expansion == "yes",]$HOG
Exp_Bnapus <- HOG_level_list$hypothesis_6_cons[HOG_level_list$hypothesis_6_cons$expansion == "yes",]$HOG

listInput <- list(Eutrema=Exp_Eutrema,Alyrata=Exp_Alyrata,Athaliana=Exp_Athaliana,Bnapus=Exp_Bnapus)
pdf(paste0(OUTDIR, "/Intersection_of_expanded_OGs.pdf"), 6, 4)
upset(fromList(listInput), nsets=4, order.by = "freq", sets.x.label = "Number of expanded OGs", mainbar.y.label = "Intersection of OGs",
      number.angles = 0, 
      point.size = 4, line.size = 1.5,
      text.scale = c(1, 1, 1, 1, 1.5, 1.5)
)
dev.off()


### find the HOGs in the candidate gene set with the GO term "metaxylem development"
candidates <- read.table("Analysis/HOG_level_list_all_tol_exp_DEG_any_tol.tsv", header =T)$HOG
SFA_OG_level$hypothesis_1[SFA_OG_level$hypothesis_1$HOG %in% candidates
                          ,]$HOG
SFA_OG_level$hypothesis_1[grep("GO:0090058", SFA_OG_level$hypothesis_1$`Gene-Ontology-Term`) 
                          ,]$HOG
intersect(SFA_OG_level$hypothesis_1[SFA_OG_level$hypothesis_1$HOG %in% candidates
                                    ,]$HOG,
          SFA_OG_level$hypothesis_1[grep("GO:0090058", SFA_OG_level$hypothesis_1$`Gene-Ontology-Term`) 
                                    ,]$HOG)
```
