## Additional File 3 for "Integration of differential expression under drought with gene family expansion unique to drought tolerant species predicts candidate genes for drought adaptation in Brassicaceae species": Analysis_A2TEA_output_with_Selection_v04_with_corrected_pvalues.R

```
# script to explore a2tea output BR-dr-03 together with results from Selection Analysis and GO term enrichment analyses
### I did not use this script for the over representation analyses of subsets with pos. selection in the Conserved set
# the script for these analyses is Selection_overrepresentation_v04_with_corrected_pvalues.Rmd
#library(readr)
library(Biostrings)
#library(ape)
library(ggplot2)
library(reshape2)
library(tidyr)
library(dplyr)
library(topGO)

setwd("/home/cropbio/sciebo/Documents/PhD/Brassicaceae-drought/BR-dr-03/Selection/Analysis/")

load("/home/cropbio/sciebo/Documents/PhD/Brassicaceae-drought/BR-dr-03/A2TEA_finished.RData")

### Count the number of genes which were in the !!! input !!! dataset per species (the ones which are in the Conserved OGs file)
Conserved_Selection_input_dataset <-  read.table("./results_20221027_Conserved_Dataset/Conserved_dataset_matrix.tsv", sep = "\t", header=T)
Conserved_genes_HOG_DE <- merge(Conserved_Selection_input_dataset, HOG_DE.a2tea, by=c("gene", "HOG"))

length(Conserved_genes_HOG_DE[Conserved_genes_HOG_DE$species=="Eutrema_salsugineum",]$gene)
# 19990
length(Conserved_genes_HOG_DE[Conserved_genes_HOG_DE$species=="Arabidopsis_lyrata",]$gene)
# 21173
length(Conserved_genes_HOG_DE[Conserved_genes_HOG_DE$species=="Arabidopsis_thaliana",]$gene)
# 19919
length(Conserved_genes_HOG_DE[Conserved_genes_HOG_DE$species=="Brassica_napus",]$gene)
# 38861

## -> the input is more ore less even (Bnapus has double as it is tetraploid)


                      ### And from that the HOGs that were evaluated (!!! results !!!) in the Selection Analysis ###
#load the results from the Selection Analysis
Conserved_Selection_results_numbers <- read.table("./results_20221027_Conserved_Dataset/OGs_number_of_branches_under_pos_selection.tsv", sep = "\t", header=F)
names(Conserved_Selection_results_numbers) <- c("HOG", "number_of_pos_sel")
Conserved_Selection_results_genes_pvalues <- read.table("./results_20221027_Conserved_Dataset/OGs_pos_selection_pvalues.tsv", sep = "\t", header=F)
names(Conserved_Selection_results_genes_pvalues) <- c("HOG", "gene", "uncorrected_selection_pvalue", "selection_pvalue")

HOGs_in_results_selection_analysis <- Conserved_Selection_results_numbers$HOG

## For all calculations on gene level, merge Conserved_Selection_results_genes_pvalues with HOG_DE.a2tea (also to obtain the species per gene)
#Problem: in Selection results, some IDs of Alyrata are with "_1" in the end or with "pg." or with "kg." in the middle. In DE results, the IDs of Alyrata are with ".1" in the end or "pg." or "kg."-->
  
#  change the "." used as separator in some Alyrata IDs to "_" in the HOG_DE.a2tea dataset

HOG_DE.a2tea_edited_01<- HOG_DE.a2tea
HOG_DE.a2tea_edited_01$gene <- gsub(".", "_", HOG_DE.a2tea_edited_01$gene, fixed = T)


Conserved_Selection_results_genes_pvalues_HOG_DE <- merge(Conserved_Selection_results_genes_pvalues, HOG_DE.a2tea_edited_01, by=c("gene", "HOG"))


### Count the number of genes which were evaluated per species (the ones which are in the results files and as such have a p-value)
length(Conserved_Selection_results_genes_pvalues_HOG_DE[Conserved_Selection_results_genes_pvalues_HOG_DE$species=="Eutrema_salsugineum",]$gene)
# 14412
length(Conserved_Selection_results_genes_pvalues_HOG_DE[Conserved_Selection_results_genes_pvalues_HOG_DE$species=="Arabidopsis_lyrata",]$gene)
# 14974
length(Conserved_Selection_results_genes_pvalues_HOG_DE[Conserved_Selection_results_genes_pvalues_HOG_DE$species=="Arabidopsis_thaliana",]$gene)
# 14408
length(Conserved_Selection_results_genes_pvalues_HOG_DE[Conserved_Selection_results_genes_pvalues_HOG_DE$species=="Brassica_napus",]$gene)
# 27962

### Count the number of genes with pos. Sel. per species per HOG
Conserved_Selection_results_genes_pvalues_HOG_DE_At <-
  Conserved_Selection_results_genes_pvalues_HOG_DE %>% 
  group_by(HOG) %>%
  summarise(Arabidopsis_thaliana_pos_sel = sum(species== "Arabidopsis_thaliana" & selection_pvalue<=0.05))

Conserved_Selection_results_genes_pvalues_HOG_DE_Al <-
  Conserved_Selection_results_genes_pvalues_HOG_DE %>% 
  group_by(HOG) %>%
  summarise(Arabidopsis_lyrata_pos_sel = sum(species== "Arabidopsis_lyrata" & selection_pvalue<=0.05))

Conserved_Selection_results_genes_pvalues_HOG_DE_Es <-
  Conserved_Selection_results_genes_pvalues_HOG_DE %>% 
  group_by(HOG) %>%
  summarise(Eutrema_salsugineum_pos_sel = sum(species== "Eutrema_salsugineum" & selection_pvalue<=0.05))

Conserved_Selection_results_genes_pvalues_HOG_DE_Bn <-
  Conserved_Selection_results_genes_pvalues_HOG_DE %>% 
  group_by(HOG) %>%
  summarise(Brassica_napus_pos_sel = sum(species== "Brassica_napus" & selection_pvalue<=0.05))

merge(merge(Conserved_Selection_results_genes_pvalues_HOG_DE_Al, Conserved_Selection_results_genes_pvalues_HOG_DE_At), merge(Conserved_Selection_results_genes_pvalues_HOG_DE_Bn, Conserved_Selection_results_genes_pvalues_HOG_DE_Es))


### Select only conserved HOGs (at least 1 gene from each (of the compared) species)
# (this is redundant because the input set consists already of Conserved HOGs, but I kept this step because the following parts depend on the variable names)
### and hereof only the ones which were evaluated in the Selection Analysis
HOG_level_list$hypothesis_1_cons <- HOG_level_list$hypothesis_1[HOG_level_list$hypothesis_1$Arabidopsis_lyrata_total>=1 & HOG_level_list$hypothesis_1$Arabidopsis_thaliana_total>=1 & HOG_level_list$hypothesis_1$Eutrema_salsugineum_total>=1 & HOG_level_list$hypothesis_1$Brassica_napus_total>=1,]
HOG_level_list$hypothesis_1_cons <- HOG_level_list$hypothesis_1_cons[HOG_level_list$hypothesis_1_cons$HOG %in% HOGs_in_results_selection_analysis,]
HOG_level_list$hypothesis_2_cons <- HOG_level_list$hypothesis_2[HOG_level_list$hypothesis_2$HOG %in% HOG_level_list$hypothesis_1_cons$HOG,]
HOG_level_list$hypothesis_3_cons <- HOG_level_list$hypothesis_3[HOG_level_list$hypothesis_3$HOG %in% HOG_level_list$hypothesis_1_cons$HOG,]
HOG_level_list$hypothesis_4_cons <- HOG_level_list$hypothesis_4[HOG_level_list$hypothesis_4$HOG %in% HOG_level_list$hypothesis_1_cons$HOG,]
HOG_level_list$hypothesis_5_cons <- HOG_level_list$hypothesis_5[HOG_level_list$hypothesis_5$HOG %in% HOG_level_list$hypothesis_1_cons$HOG,]
HOG_level_list$hypothesis_6_cons <- HOG_level_list$hypothesis_6[HOG_level_list$hypothesis_6$HOG %in% HOG_level_list$hypothesis_1_cons$HOG,]

### merge the data frames from A2TEA analysis with the ones from the Selection Analysis
# NOTE that Conserved_Selection_results_numbers contains the total number of genes or NODES which are under pos. Sel.
# when choosing per species, only the genes under pos. Sel. are considered, but not the NODES. Hence, the column "number_of_pos_sel" is not
# necessarily the sum of the values per HOG of the columns "{$species}_pos_sel"
HOG_level_list$hypothesis_1_cons_with_numbers_pos_sel <- merge(HOG_level_list$hypothesis_1_cons, Conserved_Selection_results_numbers, by="HOG")
#put all data frames into list
df_list <- list(HOG_level_list$hypothesis_1_cons_with_numbers_pos_sel, Conserved_Selection_results_genes_pvalues_HOG_DE_At, Conserved_Selection_results_genes_pvalues_HOG_DE_Al, Conserved_Selection_results_genes_pvalues_HOG_DE_Es, Conserved_Selection_results_genes_pvalues_HOG_DE_Bn)
#merge all data frames in list
HOG_level_list$hypothesis_1_cons_with_numbers_pos_sel_per_species <- Reduce(function(x, y) merge(x, y, all=TRUE), df_list)


##### Plot the overlaps of DEG and exp in the OGs with pos. Sel.

#### Select the genes from Es or Al which are 1) pos. Sel, 2) DE and 3) exp.
# load the list of expanded OGs
Eutrema_exp <- read.table("/home/cropbio/sciebo/Documents/PhD/Brassicaceae-drought/BR-dr-03/Analysis/Hypothesis_2_Exp_in_Conserved_dataset_matrix.tsv", sep="\t", header=T)
Alyrata_exp <- read.table("/home/cropbio/sciebo/Documents/PhD/Brassicaceae-drought/BR-dr-03/Analysis/Hypothesis_3_Exp_in_Conserved_dataset_matrix.tsv", sep="\t", header=T)
Athaliana_exp <- read.table("/home/cropbio/sciebo/Documents/PhD/Brassicaceae-drought/BR-dr-03/Analysis/Hypothesis_5_Exp_in_Conserved_dataset_matrix.tsv", sep="\t", header=T)
Bnapus_exp <- read.table("/home/cropbio/sciebo/Documents/PhD/Brassicaceae-drought/BR-dr-03/Analysis/Hypothesis_6_Exp_in_Conserved_dataset_matrix.tsv", sep="\t", header=T)


df <- data.frame(abbrev = c("Al", "At", "Es", "Bn"), species = unique(Conserved_Selection_results_genes_pvalues_HOG_DE$species))
#define the expanded HOGs
Al_exp <- unique(Alyrata_exp$HOG)
At_exp <- unique(Athaliana_exp$HOG)
Es_exp <- unique(Eutrema_exp$HOG)
Bn_exp <- unique(Bnapus_exp$HOG)

for (x in seq_len(nrow(df))) {
  print(paste0("Conserved_Selection_results_genes_pvalues_HOG_DE_", df[x,1], "_pos", " species is ", df[x,2]))
  print(df[x,2])
  a <- df[x,1]
  b <- df[x,2]
  
  print(head(get(paste0(a, "_exp"))))
  # 1. Select the dataframes with genes from $species which are 1) pos. Sel
  mame <- paste0("Conserved_Selection_results_genes_pvalues_HOG_DE_", a, "_pos") 
  data <-
    Conserved_Selection_results_genes_pvalues_HOG_DE[Conserved_Selection_results_genes_pvalues_HOG_DE$species==b 
                                                     & Conserved_Selection_results_genes_pvalues_HOG_DE$selection_pvalue<=0.05,]
  #print(head(assign(mame, data)))
  # 1a. Select the dataframes with genes from$species which are 2) DE
  mame2 <- paste0("Conserved_Selection_results_genes_pvalues_HOG_DE_", a,"_DE")
  data2 <- 
    Conserved_Selection_results_genes_pvalues_HOG_DE[Conserved_Selection_results_genes_pvalues_HOG_DE$species==b 
                                                     & (!is.na(Conserved_Selection_results_genes_pvalues_HOG_DE$padj)) 
                                                     & Conserved_Selection_results_genes_pvalues_HOG_DE$padj<=0.1,]
  #print(head(assign(mame2, data2)))
  
  # 1b. Select the dataframes with all genes from $species from all OGs which have at least one 
  # 1) pos. Sel and at least one 2) DE from $species
  mame2a <- paste0("Conserved_Selection_results_genes_pvalues_HOG_DE_", a,"_pos_and_DE")
  data2a <- 
    Conserved_Selection_results_genes_pvalues_HOG_DE[Conserved_Selection_results_genes_pvalues_HOG_DE$HOG %in% data2$HOG
                                                     & Conserved_Selection_results_genes_pvalues_HOG_DE$species==b 
                                                     & Conserved_Selection_results_genes_pvalues_HOG_DE$selection_pvalue<=0.05,]
  #print(head(assign(mame2a, data2a)))
  
  # More detailed:
  # Select the dataframes with genes from $species which are 1) pos. Sel and 2) DE
  mame3 <- paste0("Conserved_Selection_results_genes_pvalues_HOG_DE_", a, "_DEG_is_pos")
  data3 <-
    data2a[(!is.na(data2a$padj)) & data2a$padj<=0.1,]
  #print(head(assign(mame3, data3)))
  
  # Select the dataframes with genes from $species where the pos. Sel. is NON-DE
  mame3a <- paste0("Conserved_Selection_results_genes_pvalues_HOG_DE_", a, "_DE_and_NON_DEG_is_pos")
  data3a <-
    data2a[((is.na(data2a$padj)) | data2a$padj>0.1),]
  #print(head(assign(mame3a, data3a)))
  
  # Select the dataframes with genes from $species which are 1) pos. Sel and 2) exp
  mame4 <- paste0("Conserved_Selection_results_genes_pvalues_HOG_DE_", a, "_exp_and_pos") 
  data4 <-
    data[data$HOG %in% get(paste0(a, "_exp")),]
  #print(head(assign(mame4, data4)))
  
  # Select the dataframes with genes from $species which are 1) pos. Sel and 2) exp and 3) have a DE from $species
  # (it is not necessarily the one under pos. Sel.)
  mame4a <- paste0("Conserved_Selection_results_genes_pvalues_HOG_DE_", a, "_exp_and_DE_and_pos") 
  data4a <- data4[data4$HOG %in% data2$HOG,]
  #print(head(assign(mame4a, data4a)))
  
  # More detailed:
  # 3. Select the dataframes with genes from $species which are 1) pos. Sel and 2) the same gene is DE and 3) exp. in $species
  mame5 <- paste0("Conserved_Selection_results_genes_pvalues_HOG_DE_", a, "_exp_and_DEG_is_pos")
  data5 <-
    data3[data3$HOG %in% get(paste0(a, "_exp")),]
  #print(head(assign(mame5, data5)))
  # 3x. Select the dataframes with genes from $species which are in a DE in $species HOG,
  # and 1) pos. Sel and 2) a NON-DE gene from $species is pos. sel. and 3) exp. in $species
  mame5a <- paste0("Conserved_Selection_results_genes_pvalues_HOG_DE_", a, "_exp_and_DE_and_NON-DEG_is_pos")
  data5a <-
    data3a[data3a$HOG %in% get(paste0(a, "_exp")),]
  #print(head(assign(mame5a, data5a)))
  
  # i missed the simple case:
  mame5b <- paste0("Conserved_Selection_results_genes_pvalues_HOG_DE_", a, "_exp_and_DE")
  data5b <- Conserved_Selection_results_genes_pvalues_HOG_DE[Conserved_Selection_results_genes_pvalues_HOG_DE$HOG %in% get(paste0(a, "_exp"))
                                                             & Conserved_Selection_results_genes_pvalues_HOG_DE$HOG %in% data2$HOG,]
  
  # 4. count number of HOGs (or genes) in the subsets 1) pos. Sel and 2) DE and 3) exp. per $species
  
  #1)
  mame6 <- paste0("number_of_OGs_", a, "_pos")
  data6 <- length(unique(data$HOG))
  #print(mame6)
  #print(data6)
  # 2573
  
  mame7 <- paste0("genes_", a, "_pos") 
  data7 <- length(unique(data[data$species==b,]$gene))
  # 2687
  
  #2a)
  mame8 <- paste0("number_of_OGs_", a, "_DEG_is_pos")
  data8 <- length(unique(data3$HOG))
  # 996
  # --> here is missing at least 1 true positive, which is found in above way of subsetting the combination of Pos. Sel. and DE:
  ## N0.HOG0000076 is pos. Sel. in Eutrema and DE in Eutrema
  
  ##### !!!!! The subsetting HERE takes into account the gene level: so the GENE under pos. Sel. must also be DE !!!!!!!!!! ######
  
  mame9 <- paste0("genes_", a, "_DEG_is_pos") 
  data9 <- length(unique(data3[data3$species==b,]$gene))
  # 1004
  
  # calculate the number of HOGs within the HOGs DE in species, where the pos. sel. gene from species is NOT DE
  mame10 <- paste0("number_of_OGs_", a, "_DE_and_NON-DEG_is_pos")
  data10 <- length(unique(data3a$HOG))
  # gene level
  mame11 <- paste0("genes_", a, "_DE_and_NON-DEG_is_pos")
  data11 <- length(unique(data3a[data3a$species==b,]$gene))
  
  
  # more general background:
  mame12 <- paste0("number_of_OGs_", a, "_DE")
  data12 <- length(unique(data2$HOG))
  # gene level:
  mame13 <- paste0("genes_", a, "_DE")
  data13 <-length(unique(Conserved_Selection_results_genes_pvalues_HOG_DE[Conserved_Selection_results_genes_pvalues_HOG_DE$HOG %in% data2$HOG & Conserved_Selection_results_genes_pvalues_HOG_DE$species==b,]$gene))
  # DEGs:
  mame14 <- paste0("DEGs_", a, "_DE")
  data14 <- length(unique(data2[data2$species==b,]$gene))
  
  
  #more general with pos sel.:
  mame15 <- paste0("number_of_OGs_", a, "_pos_and_DE")
  data15 <- length(unique(data2a$HOG))
  # gene level:
  mame16 <- paste0("genes_", a, "_in_pos_and_DE")
  data16 <- length(unique(data2a[data2a$species==b,]$gene))
  
  #2b)
  mame17 <- paste0("number_of_OGs_", a, "_pos_and_exp")
  data17 <- length(unique(data4$HOG))
  # 239
  # gene level:
  mame18 <- paste0("genes_", a, "_pos_and_exp")
  data18 <- length(unique(data4[data4$species==b,]$gene))
  # 306
  
  # exp and DE and pos
  mame19 <- paste0("number_of_OGs_", a, "_exp_and_DE_and_pos")
  data19 <- length(unique(data4a$HOG))
  # gene level:
  mame20 <- paste0("genes_", a, "_in_exp_and_DE_and_pos")
  data20 <- length(unique(data4a[data4a$species==b,]$gene))
  
  
  #3) 
  mame21 <- paste0("number_of_OGs_", a, "_exp_and_DE_is_pos")
  data21 <- length(unique(data5$HOG))
  # 58
  # gene level
  mame22 <- paste0("genes_", a, "_exp_and_DE_is_pos")
  data22 <- length(unique(data5$gene))
  # 64
  
  # the contrast: exp and DE and the NON-DE is pos
  mame23 <- paste0("number_of_OGs_", a, "_exp_and_DE_and_NON-DE_is_pos")
  data23 <- length(unique(data5a$HOG))
  # 
  # gene level
  mame24 <- paste0("genes_", a, "_exp_and_DE_and_NON-DE_is_pos")
  data24 <- length(unique(data5a$gene))
  # 
  
  
  # background set (all conserved HOGs which were evaluated for pos. sel.)
  mame25 <- paste0("number_of_OGs_", a, "_in_Conserved")
  data25 <- length(unique(Conserved_Selection_results_genes_pvalues_HOG_DE$HOG))
  # same on gene level
  mame26 <- paste0("genes_", a, "_in_Conserved")
  data26 <- length(unique(Conserved_Selection_results_genes_pvalues_HOG_DE[
    Conserved_Selection_results_genes_pvalues_HOG_DE$species==b,]$gene))
  
  # background exp
  mame27 <- paste0("number_of_OGs_", a, "_exp")
  data27 <- length(unique(Conserved_Selection_results_genes_pvalues_HOG_DE[
    Conserved_Selection_results_genes_pvalues_HOG_DE$HOG %in% get(paste0(a, "_exp")),]$HOG))
  
  mame28 <- paste0("genes_", a, "_exp")
  data28 <- length(unique(Conserved_Selection_results_genes_pvalues_HOG_DE[
    Conserved_Selection_results_genes_pvalues_HOG_DE$HOG %in% get(paste0(a, "_exp"))
    & Conserved_Selection_results_genes_pvalues_HOG_DE$species==b,]$gene))
  
  mame29 <- paste0("number_of_OGs_", a, "_exp_and_DE")
  data29 <- length(unique(data5b$HOG))
  # gene level:
  mame30 <- paste0("genes_", a, "_exp_and_DE")
  data30 <- length(unique(data5b[data5b$species==b,]$gene))
  # only the DEGs:
  mame31 <- paste0("DEGs_", a, "_exp_and_DE")
  data31 <- length(unique(data5b[data5b$species==b
                                 & (!is.na(data5b$padj)) 
                                 & data5b$padj<=0.1,]$gene))
  # only the NON-DEGs:
  mame32 <- paste0("NON-DEGs_", a, "_exp_and_DE")
  data32 <- length(unique(data5b[data5b$species==b
                                 & (is.na(data5b$padj) 
                                    | data5b$padj>0.1),]$gene))
  
  
  # put the names of the sets into a character vector
  set_names <- c(paste0("mame", seq(6,32)))
  #print(set_names)
  print(as.character(sapply(set_names, get)))
  
  # put the counts of the sets into a numeric vector
  set_values <- c(paste0("data", seq(6,32)))
  #print(set_values)
  print(as.numeric(sapply(set_values, get)))
  
  # combine the names and the counts of the sets to a data.frame
  df_name <- paste0("counts_pos_sel_", a)
  output_df <- data.frame(set_names=as.character(sapply(set_names, get)), counts=as.numeric(sapply(set_values, get)))
  # save the output_df in a variable
  assign(df_name, output_df)
  # export the data.frame
  write.table(output_df, paste0(df_name, ".tsv"), sep="\t", row.names = F, col.names = T)
}

### the numbers of genes under pos selection were counted in the above loop over all hypotheses. 
# The following lines are per species, the counts I had used previously but NOT in the results of the manuscript.
# The table 20231027_Conserved_Selection_results_genes_pvalues_HOG_DE_Es_pos_and_DE_and_exp.tsv which I used for manual exploration of the candidate gene families might miss Info on some HOGs

# 1. Select the dataframes with genes from Es or Al which are 1) pos. Sel
Conserved_Selection_results_genes_pvalues_HOG_DE_Es_pos <-
  Conserved_Selection_results_genes_pvalues_HOG_DE[Conserved_Selection_results_genes_pvalues_HOG_DE$species=="Eutrema_salsugineum" & Conserved_Selection_results_genes_pvalues_HOG_DE$selection_pvalue<=0.05,]
Conserved_Selection_results_genes_pvalues_HOG_DE_Al_pos <-
  Conserved_Selection_results_genes_pvalues_HOG_DE[Conserved_Selection_results_genes_pvalues_HOG_DE$species=="Arabidopsis_lyrata" & Conserved_Selection_results_genes_pvalues_HOG_DE$selection_pvalue<=0.05,]
# 1a. Select the dataframes with genes from Es or Al which are 2) DE
Conserved_Selection_results_genes_pvalues_HOG_DE_Es_DE <- 
  Conserved_Selection_results_genes_pvalues_HOG_DE[Conserved_Selection_results_genes_pvalues_HOG_DE$species=="Eutrema_salsugineum" 
                                                   & (!is.na(Conserved_Selection_results_genes_pvalues_HOG_DE$padj)) 
                                                   & Conserved_Selection_results_genes_pvalues_HOG_DE$padj<=0.1,]
length(unique(Conserved_Selection_results_genes_pvalues_HOG_DE_Es_DE$HOG))
# 5750

# 2a. Select the dataframes with genes from Es or Al which are 1) pos. Sel and 2) DE
Conserved_Selection_results_genes_pvalues_HOG_DE_Es_pos_and_DE <-
  Conserved_Selection_results_genes_pvalues_HOG_DE_Es_pos[(!is.na(Conserved_Selection_results_genes_pvalues_HOG_DE_Es_pos$padj)) & Conserved_Selection_results_genes_pvalues_HOG_DE_Es_pos$padj<=0.1,]
Conserved_Selection_results_genes_pvalues_HOG_DE_Al_pos_and_DE <-
  Conserved_Selection_results_genes_pvalues_HOG_DE_Al_pos[!is.na(Conserved_Selection_results_genes_pvalues_HOG_DE_Al_pos$padj) & Conserved_Selection_results_genes_pvalues_HOG_DE_Al_pos$padj<=0.1,]
# 2b. Select the dataframes with genes from Es or Al which are 1) pos. Sel and 2) exp
Conserved_Selection_results_genes_pvalues_HOG_DE_Es_pos_and_exp <-
  Conserved_Selection_results_genes_pvalues_HOG_DE_Es_pos[Conserved_Selection_results_genes_pvalues_HOG_DE_Es_pos$HOG %in% Eutrema_exp$HOG,]

Conserved_Selection_results_genes_pvalues_HOG_DE_Al_pos_and_exp <-
  Conserved_Selection_results_genes_pvalues_HOG_DE_Al_pos[Conserved_Selection_results_genes_pvalues_HOG_DE_Al_pos$HOG %in% Alyrata_exp$HOG,]

# 3. Select the dataframes with genes from Es or Al which are 1) pos. Sel and 2) DE and 3) exp. in this species
Conserved_Selection_results_genes_pvalues_HOG_DE_Es_pos_and_DE_and_exp <-
  Conserved_Selection_results_genes_pvalues_HOG_DE_Es_pos_and_DE[Conserved_Selection_results_genes_pvalues_HOG_DE_Es_pos_and_DE$HOG %in% Eutrema_exp$HOG,]
Conserved_Selection_results_genes_pvalues_HOG_DE_Al_pos_and_DE_and_exp <-
  Conserved_Selection_results_genes_pvalues_HOG_DE_Al_pos_and_DE[Conserved_Selection_results_genes_pvalues_HOG_DE_Al_pos_and_DE$HOG %in% Alyrata_exp$HOG,]

# 4. count number of HOGs (or genes) in the subsets 1) pos. Sel and 2) DE and 3) exp. in this species

########### Eutrema

#1)
number_of_OGs_Es_pos <- length(unique(Conserved_Selection_results_genes_pvalues_HOG_DE_Es_pos$HOG))
# 1318
genes_Es_pos <-length(unique(Conserved_Selection_results_genes_pvalues_HOG_DE_Es_pos$gene))
# 1371

#2a)
number_of_OGs_Es_pos_and_DE <- length(unique(Conserved_Selection_results_genes_pvalues_HOG_DE_Es_pos_and_DE$HOG))
# 486
## --> The subsetting HERE takes into account the gene level: so the GENE under pos. Sel. must also be DE

genes_Es_pos_and_DE <- length(unique(Conserved_Selection_results_genes_pvalues_HOG_DE_Es_pos_and_DE$gene))
# 489

#2b)
number_of_OGs_Es_pos_and_exp <- length(unique(Conserved_Selection_results_genes_pvalues_HOG_DE_Es_pos_and_exp$HOG))
# 179
genes_Es_pos_and_exp <- length(unique(Conserved_Selection_results_genes_pvalues_HOG_DE_Es_pos_and_exp$gene))
# 209

#3) 
number_of_OGs_Es_pos_and_DE_and_exp <- length(unique(Conserved_Selection_results_genes_pvalues_HOG_DE_Es_pos_and_DE_and_exp$HOG))
# 39
genes_Es_pos_and_DE_and_exp <- length(unique(Conserved_Selection_results_genes_pvalues_HOG_DE_Es_pos_and_DE_and_exp$gene))
# 41

### extract the Athaliana gene ID in these HOGs of interest
OGs_of_interest_Es <- read.table("20231027_Conserved_Selection_results_genes_pvalues_HOG_DE_Es_pos_and_DE_and_exp.tsv", sep="\t", header=T)$HOG
AT_genes_in_OGs_of_interest_Es <- HOG_DE.a2tea[HOG_DE.a2tea$HOG %in% OGs_of_interest_Es & HOG_DE.a2tea$species=="Arabidopsis_thaliana",]$gene

# export this as list with Athaliana genes of HOGs of interest to blast the CDS against the Eutrema genome (to verify the Eutrema annotation)
# write.table(AT_genes_in_OGs_of_interest_Es, "/home/cropbio/sciebo/Documents/PhD/Brassicaceae-drought/BR-dr-03/Selection/Analysis/List_of_AT_genes_in_OGs_of_interest_Es.txt"
#            , row.names = F, col.names = F, quote=F)

### Plot the intersect
library(UpSetR)
listInput <- list(Es_DE = Conserved_Selection_results_genes_pvalues_HOG_DE_Es_pos_and_DE$HOG,
                  Es_exp = Conserved_Selection_results_genes_pvalues_HOG_DE_Es_pos_and_exp$HOG)

pdf("Intersection_of_OGs_with_pos_selection_in_Eutrema_and_DE_and_exp.pdf", 7, 5)
upset(fromList(listInput), nsets=2, nintersects=NA, order.by = "freq", sets.x.label = "Number of OGs with at least \n1 pos. Sel. gene from Eutrema", mainbar.y.label = "Intersection of OGs",
      number.angles = 0, 
      point.size = 2, line.size = 1,
      text.scale = c(2,1.5,1,1.5,1.5,1.5)
)
dev.off()

########### Alyrata
#1)
number_of_OGs_Al_pos <- length(unique(Conserved_Selection_results_genes_pvalues_HOG_DE_Al_pos$HOG))
# 1335
genes_Al_pos <-length(unique(Conserved_Selection_results_genes_pvalues_HOG_DE_Al_pos$gene))
# 1423

#2a)
number_of_OGs_Al_pos_and_DE <- length(unique(Conserved_Selection_results_genes_pvalues_HOG_DE_Al_pos_and_DE$HOG))
# 536
genes_Al_pos_and_DE <- length(unique(Conserved_Selection_results_genes_pvalues_HOG_DE_Al_pos_and_DE$gene))
# 544

#2b)
number_of_OGs_Al_pos_and_exp <- length(unique(Conserved_Selection_results_genes_pvalues_HOG_DE_Al_pos_and_exp$HOG))
# 272
genes_Al_pos_and_exp <- length(unique(Conserved_Selection_results_genes_pvalues_HOG_DE_Al_pos_and_exp$gene))
# 326

#3) 
number_of_OGs_Al_pos_and_DE_and_exp <- length(unique(Conserved_Selection_results_genes_pvalues_HOG_DE_Al_pos_and_DE_and_exp$HOG))
# 64
genes_Al_pos_and_DE_and_exp <- length(unique(Conserved_Selection_results_genes_pvalues_HOG_DE_Al_pos_and_DE_and_exp$gene))
# 70

listInput <- list(Al_pos_and_DE = Conserved_Selection_results_genes_pvalues_HOG_DE_Al_pos_and_DE$HOG,
                  Al_pos_and_exp = Conserved_Selection_results_genes_pvalues_HOG_DE_Al_pos_and_exp$HOG)

pdf("Intersection_of_OGs_with_pos_selection_in_Alyrata_and_DE_and_exp.pdf", 6, 4)
upset(fromList(listInput), nsets=2, nintersects=NA, order.by = "freq", sets.x.label = "Number of OGs", mainbar.y.label = "Intersection of OGs",
      number.angles = 0, 
      point.size = 2, line.size = 1,
      text.scale = c(1.5, 0.8,0.8, 0.8,0.8, 0.8)
)
dev.off()

### Plot the Intersect of Es and Al

listInput <- list(Es_pos_and_DE = Conserved_Selection_results_genes_pvalues_HOG_DE_Es_pos_and_DE$HOG,
                  Es_pos_and_exp = Conserved_Selection_results_genes_pvalues_HOG_DE_Es_pos_and_exp$HOG,
                  Al_pos_and_DE = Conserved_Selection_results_genes_pvalues_HOG_DE_Al_pos_and_DE$HOG,
                  Al_pos_and_exp = Conserved_Selection_results_genes_pvalues_HOG_DE_Al_pos_and_exp$HOG)

pdf("Intersection_of_OGs_with_pos_selection_in_Eutrema_and_Alyrata_and_DE_and_exp.pdf", 6, 4)
upset(fromList(listInput), nsets=4, nintersects=NA, order.by = "freq", sets.x.label = "Number of OGs", mainbar.y.label = "Intersection of OGs",
      number.angles = 0, 
      point.size = 2, line.size = 1,
      text.scale = c(1.5, 0.8,0.8, 0.8,0.8, 0.8)
)
dev.off()

### Do the same for the sensitive species
###### Select the genes from At or Bn which are 1) pos. Sel, 2) DE and 3) exp.
# load the list of expanded OGs
Athaliana_exp <- read.table("/home/cropbio/sciebo/Documents/PhD/Brassicaceae-drought/BR-dr-03/Analysis/Hypothesis_5_Exp_in_Conserved_dataset_matrix.tsv", sep="\t", header=T)
Bnapus_exp <- read.table("/home/cropbio/sciebo/Documents/PhD/Brassicaceae-drought/BR-dr-03/Analysis/Hypothesis_6_Exp_in_Conserved_dataset_matrix.tsv", sep="\t", header=T)


# 1. Select the dataframes with genes from At or Bn which are 1) pos. Sel
Conserved_Selection_results_genes_pvalues_HOG_DE_At_pos <-
  Conserved_Selection_results_genes_pvalues_HOG_DE[Conserved_Selection_results_genes_pvalues_HOG_DE$species=="Arabidopsis_thaliana" & Conserved_Selection_results_genes_pvalues_HOG_DE$selection_pvalue<=0.05,]
Conserved_Selection_results_genes_pvalues_HOG_DE_Bn_pos <-
  Conserved_Selection_results_genes_pvalues_HOG_DE[Conserved_Selection_results_genes_pvalues_HOG_DE$species=="Brassica_napus" & Conserved_Selection_results_genes_pvalues_HOG_DE$selection_pvalue<=0.05,]
# 2a. Select the dataframes with genes from At or Bn which are 1) pos. Sel and 2) DE
Conserved_Selection_results_genes_pvalues_HOG_DE_At_pos_and_DE <-
  Conserved_Selection_results_genes_pvalues_HOG_DE_At_pos[!is.na(Conserved_Selection_results_genes_pvalues_HOG_DE_At_pos$padj) & Conserved_Selection_results_genes_pvalues_HOG_DE_At_pos$padj<=0.1,]
Conserved_Selection_results_genes_pvalues_HOG_DE_Bn_pos_and_DE <-
  Conserved_Selection_results_genes_pvalues_HOG_DE_Bn_pos[!is.na(Conserved_Selection_results_genes_pvalues_HOG_DE_Bn_pos$padj) & Conserved_Selection_results_genes_pvalues_HOG_DE_Bn_pos$padj<=0.1,]
# 2b. Select the dataframes with genes from At or Bn which are 1) pos. Sel and 2) exp
Conserved_Selection_results_genes_pvalues_HOG_DE_At_pos_and_exp <-
  Conserved_Selection_results_genes_pvalues_HOG_DE_At_pos[Conserved_Selection_results_genes_pvalues_HOG_DE_At_pos$HOG %in% Athaliana_exp$HOG,]
Conserved_Selection_results_genes_pvalues_HOG_DE_Bn_pos_and_exp <-
  Conserved_Selection_results_genes_pvalues_HOG_DE_Bn_pos[Conserved_Selection_results_genes_pvalues_HOG_DE_Bn_pos$HOG %in% Bnapus_exp$HOG,]

# 3. Select the dataframes with genes from At or Bn which are 1) pos. Sel and 2) DE and 3) exp. in this species
Conserved_Selection_results_genes_pvalues_HOG_DE_At_pos_and_DE_and_exp <-
  Conserved_Selection_results_genes_pvalues_HOG_DE_At_pos_and_DE[Conserved_Selection_results_genes_pvalues_HOG_DE_At_pos_and_DE$HOG %in% Athaliana_exp$HOG,]
Conserved_Selection_results_genes_pvalues_HOG_DE_Bn_pos_and_DE_and_exp <-
  Conserved_Selection_results_genes_pvalues_HOG_DE_Bn_pos_and_DE[Conserved_Selection_results_genes_pvalues_HOG_DE_Bn_pos_and_DE$HOG %in% Bnapus_exp$HOG,]

# 4. count number of HOGs (or genes) in the subsets 1) pos. Sel and 2) DE and 3) exp. in this species

########### Athaliana

#1)
number_of_OGs_At_pos <- length(unique(Conserved_Selection_results_genes_pvalues_HOG_DE_At_pos$HOG))
# 1127
genes_At_pos <-length(unique(Conserved_Selection_results_genes_pvalues_HOG_DE_At_pos$gene))
# 1165

#2a)
number_of_OGs_At_pos_and_DE <- length(unique(Conserved_Selection_results_genes_pvalues_HOG_DE_At_pos_and_DE$HOG))
# 133
genes_At_pos_and_DE <- length(unique(Conserved_Selection_results_genes_pvalues_HOG_DE_At_pos_and_DE$gene))
# 133

#2b)
number_of_OGs_At_pos_and_exp <- length(unique(Conserved_Selection_results_genes_pvalues_HOG_DE_At_pos_and_exp$HOG))
# 83
genes_At_pos_and_exp <- length(unique(Conserved_Selection_results_genes_pvalues_HOG_DE_At_pos_and_exp$gene))
# 97

#3) 
number_of_OGs_At_pos_and_DE_and_exp <- length(unique(Conserved_Selection_results_genes_pvalues_HOG_DE_At_pos_and_DE_and_exp$HOG))
# 6
genes_At_pos_and_DE_and_exp <- length(unique(Conserved_Selection_results_genes_pvalues_HOG_DE_At_pos_and_DE_and_exp$gene))
# 6

### Plot the intersect
library(UpSetR)
listInput <- list(At_pos_and_DE = Conserved_Selection_results_genes_pvalues_HOG_DE_At_pos_and_DE$HOG,
                  At_pos_and_exp = Conserved_Selection_results_genes_pvalues_HOG_DE_At_pos_and_exp$HOG)

pdf("Intersection_of_OGs_with_pos_selection_in_Athaliana_and_DE_and_exp.pdf", 6, 4)
upset(fromList(listInput), nsets=2, nintersects=NA, order.by = "freq", sets.x.label = "Number of OGs", mainbar.y.label = "Intersection of OGs",
      number.angles = 0, 
      point.size = 2, line.size = 1,
      text.scale = c(1.5, 0.8,0.8, 0.8,0.8, 0.8)
)
dev.off()

########### Bnapus

#1)
number_of_OGs_Bn_pos <- length(unique(Conserved_Selection_results_genes_pvalues_HOG_DE_Bn_pos$HOG))
# 2320
genes_Bn_pos <-length(unique(Conserved_Selection_results_genes_pvalues_HOG_DE_Bn_pos$gene))
# 2732

#2a)
number_of_OGs_Bn_pos_and_DE <- length(unique(Conserved_Selection_results_genes_pvalues_HOG_DE_Bn_pos_and_DE$HOG))
# 558
genes_Bn_pos_and_DE <- length(unique(Conserved_Selection_results_genes_pvalues_HOG_DE_Bn_pos_and_DE$gene))
# 596

#2b)
number_of_OGs_Bn_pos_and_exp <- length(unique(Conserved_Selection_results_genes_pvalues_HOG_DE_Bn_pos_and_exp$HOG))
# 405
genes_Bn_pos_and_exp <- length(unique(Conserved_Selection_results_genes_pvalues_HOG_DE_Bn_pos_and_exp$gene))
# 565

#3) 
number_of_OGs_Bn_pos_and_DE_and_exp <- length(unique(Conserved_Selection_results_genes_pvalues_HOG_DE_Bn_pos_and_DE_and_exp$HOG))
# 95
genes_Bn_pos_and_DE_and_exp <- length(unique(Conserved_Selection_results_genes_pvalues_HOG_DE_Bn_pos_and_DE_and_exp$gene))
# 109

listInput <- list(Bn_pos_and_DE = Conserved_Selection_results_genes_pvalues_HOG_DE_Bn_pos_and_DE$HOG,
                  Bn_pos_and_exp = Conserved_Selection_results_genes_pvalues_HOG_DE_Bn_pos_and_exp$HOG)

pdf("Intersection_of_OGs_with_pos_selection_in_Bnapus_and_DE_and_exp.pdf", 6, 4)
upset(fromList(listInput), nsets=2, nintersects=NA, order.by = "freq", sets.x.label = "Number of OGs", mainbar.y.label = "Intersection of OGs",
      number.angles = 0, 
      point.size = 2, line.size = 1,
      text.scale = c(1.5, 0.8,0.8, 0.8,0.8, 0.8)
)
dev.off()

### Plot the Intersect of At and Bn
library(UpSetR)
listInput <- list(At_pos_and_DE = Conserved_Selection_results_genes_pvalues_HOG_DE_At_pos_and_DE$HOG,
                  At_pos_and_exp = Conserved_Selection_results_genes_pvalues_HOG_DE_At_pos_and_exp$HOG,
                  Bn_pos_and_DE = Conserved_Selection_results_genes_pvalues_HOG_DE_Bn_pos_and_DE$HOG,
                  Bn_pos_and_exp = Conserved_Selection_results_genes_pvalues_HOG_DE_Bn_pos_and_exp$HOG)

pdf("Intersection_of_OGs_with_pos_selection_in_Athaliana_and_Bnapus_and_DE_and_exp.pdf", 6, 4)
upset(fromList(listInput), nsets=4, nintersects=NA, order.by = "freq", sets.x.label = "Number of OGs with at least \n1 pos. Sel. gene from SPECIES", mainbar.y.label = "Intersection of OGs",
      number.angles = 0, 
      point.size = 2, line.size = 1,
      text.scale = c(1.5, 0.8,0.8, 0.8,0.8, 0.8)
)
dev.off()

### Plot the Intersect of all 4 species
library(UpSetR)
listInput <- list(Es_DE = Conserved_Selection_results_genes_pvalues_HOG_DE_Es_pos_and_DE$HOG,
                  Es_exp = Conserved_Selection_results_genes_pvalues_HOG_DE_Es_pos_and_exp$HOG,
                  Al_DE = Conserved_Selection_results_genes_pvalues_HOG_DE_Al_pos_and_DE$HOG,
                  Al_exp = Conserved_Selection_results_genes_pvalues_HOG_DE_Al_pos_and_exp$HOG,
                  At_DE = Conserved_Selection_results_genes_pvalues_HOG_DE_At_pos_and_DE$HOG,
                  At_exp = Conserved_Selection_results_genes_pvalues_HOG_DE_At_pos_and_exp$HOG,
                  Bn_DE = Conserved_Selection_results_genes_pvalues_HOG_DE_Bn_pos_and_DE$HOG,
                  Bn_exp = Conserved_Selection_results_genes_pvalues_HOG_DE_Bn_pos_and_exp$HOG)
sets <- c("Es_DE", "Es_exp", "Al_DE", "Al_exp", "At_DE", "At_exp", "Bn_DE", "Bn_exp")

pdf("Intersection_of_OGs_with_pos_selection_in_All_4_species_and_DE_and_exp.pdf", 8, 6)
upset(fromList(listInput), nsets=8, nintersects=NA, sets.x.label = "Number of OGs with at least \n1 pos. Sel. gene from SPECIES", mainbar.y.label = "Intersection of OGs",
      keep.order = T, order.by = "degree",
      sets =sets,
      number.angles = 0, show.numbers = F,
      point.size = 2, line.size = 1,
      text.scale = c(2,1.5,1,1.5,1.5,1.5)
)
dev.off()

pdf("Intersection_of_OGs_with_pos_selection_in_All_4_species_and_DE_and_exp_ordered_by_freq.pdf", 8, 6)
upset(fromList(listInput), nsets=8, nintersects=NA, sets.x.label = "Number of OGs with at least \n1 pos. Sel. gene from SPECIES", mainbar.y.label = "Intersection of OGs",
      keep.order = T, order.by = "freq",
      sets =sets,
      number.angles = 0, show.numbers = F,
      point.size = 2, line.size = 1,
      text.scale = c(2,1.5,1,1.5,1.5,1.5)
)
dev.off()

# Find the focus set of 17 candidate HOGs which are exp in both tolerant and uniquely up regulated in both tolerant in the selection results:
HOGs_tol_uniquely_up  <- unique(read.table("/home/cropbio/sciebo/Documents/PhD/Brassicaceae-drought/BR-dr-03/Analysis/Conserved_with_expression_with_species_specific_category_only_both_tol_up_and_exp_both_tol.tsv", header=T, sep ="\t")$HOG)
# extract the respective HOGs from the selection and DE results combination 
Conserved_Selection_results_genes_pvalues_HOG_DE_HOGs_tol_uniquely_up <- Conserved_Selection_results_genes_pvalues_HOG_DE[Conserved_Selection_results_genes_pvalues_HOG_DE$HOG %in% HOGs_tol_uniquely_up,]
Conserved_Selection_results_genes_pvalues_HOG_DE_HOGs_tol_uniquely_up_sorted <- Conserved_Selection_results_genes_pvalues_HOG_DE_HOGs_tol_uniquely_up[order(Conserved_Selection_results_genes_pvalues_HOG_DE_HOGs_tol_uniquely_up$HOG, Conserved_Selection_results_genes_pvalues_HOG_DE_HOGs_tol_uniquely_up$species),]

# how many HOGs were evaluated?
length(unique(Conserved_Selection_results_genes_pvalues_HOG_DE_HOGs_tol_uniquely_up_sorted$HOG))
# 7
DEGs_under_pos_sel <- Conserved_Selection_results_genes_pvalues_HOG_DE_HOGs_tol_uniquely_up_sorted[!is.na(Conserved_Selection_results_genes_pvalues_HOG_DE_HOGs_tol_uniquely_up_sorted$selection_pvalue)
                                                                                                            &
                                                                                                            Conserved_Selection_results_genes_pvalues_HOG_DE_HOGs_tol_uniquely_up_sorted$selection_pvalue <= 0.05
                                                                                                            &
                                                                                                            !is.na(Conserved_Selection_results_genes_pvalues_HOG_DE_HOGs_tol_uniquely_up_sorted$padj)
                                                                                                            &
                                                                                                            Conserved_Selection_results_genes_pvalues_HOG_DE_HOGs_tol_uniquely_up_sorted$padj <= 0.1
                                                                                                  ,]
Conserved_Selection_results_genes_pvalues_HOG_DE_HOGs_tol_uniquely_up_sorted_where_DEGs_under_pos_sel <- 
  Conserved_Selection_results_genes_pvalues_HOG_DE_HOGs_tol_uniquely_up_sorted[Conserved_Selection_results_genes_pvalues_HOG_DE_HOGs_tol_uniquely_up_sorted$HOG %in% DEGs_under_pos_sel$HOG,]
# How many have a DEG under pos sel?  
length(unique(Conserved_Selection_results_genes_pvalues_HOG_DE_HOGs_tol_uniquely_up_sorted_where_DEGs_under_pos_sel$HOG))
# 4


################# GO term enrichment analysis of HOGs with Selection in 1) Eutrema or 2) Alyrata

### find overrep. GO terms in the subset of OGs with Selection
library("topGO")
setwd("/home/cropbio/sciebo/Documents/PhD/Brassicaceae-drought/BR-dr-03/Selection/GO_term_enrichment_analyses_Selection_with_corrected_pvalues/")
load("/home/cropbio/sciebo/Documents/PhD/Brassicaceae-drought/BR-dr-03/A2TEA_finished.RData")

# SFA_OG_level$hypothesis_1 contains GO terms per all OGs. 
# Select conserved set from SFA_OG_level$hypothesis_1
Conserved <- HOG_level_list$hypothesis_1[HOG_level_list$hypothesis_1$Arabidopsis_lyrata_total>=1 & HOG_level_list$hypothesis_1$Arabidopsis_thaliana_total>=1 & HOG_level_list$hypothesis_1$Eutrema_salsugineum_total>=1 & HOG_level_list$hypothesis_1$Brassica_napus_total>=1,]$HOG
GO_Conserved <- SFA_OG_level$hypothesis_1[SFA_OG_level$hypothesis_1$HOG %in% Conserved,]
write.table(GO_Conserved, "GO_annotations_Conserved.tsv", row.names = F, col.names = T, sep="\t", quote=F)

### Select only conserved HOGs (at least 1 gene from each (of the compared) species)
### and hereof only the ones which were evaluated in the Selection Analysis
HOG_level_list$hypothesis_1_cons <- HOG_level_list$hypothesis_1[HOG_level_list$hypothesis_1$Arabidopsis_lyrata_total>=1 & HOG_level_list$hypothesis_1$Arabidopsis_thaliana_total>=1 & HOG_level_list$hypothesis_1$Eutrema_salsugineum_total>=1 & HOG_level_list$hypothesis_1$Brassica_napus_total>=1,]
HOG_level_list$hypothesis_1_cons <- HOG_level_list$hypothesis_1_cons[HOG_level_list$hypothesis_1_cons$HOG %in% HOGs_in_results_selection_analysis,]
HOG_level_list$hypothesis_2_cons <- HOG_level_list$hypothesis_2[HOG_level_list$hypothesis_2$HOG %in% HOG_level_list$hypothesis_1_cons$HOG,]
HOG_level_list$hypothesis_3_cons <- HOG_level_list$hypothesis_3[HOG_level_list$hypothesis_3$HOG %in% HOG_level_list$hypothesis_1_cons$HOG,]
HOG_level_list$hypothesis_4_cons <- HOG_level_list$hypothesis_4[HOG_level_list$hypothesis_4$HOG %in% HOG_level_list$hypothesis_1_cons$HOG,]
HOG_level_list$hypothesis_5_cons <- HOG_level_list$hypothesis_5[HOG_level_list$hypothesis_5$HOG %in% HOG_level_list$hypothesis_1_cons$HOG,]
HOG_level_list$hypothesis_6_cons <- HOG_level_list$hypothesis_6[HOG_level_list$hypothesis_6$HOG %in% HOG_level_list$hypothesis_1_cons$HOG,]

### Subset the Hypotheses HOG sets. Numbers of parts refer to numbers of parts in "Hypotheses_to_test_with_selection.txt"

### Select the CONSERVED data set!!! ###
### And from that the OGs that were evaluated in the Selection Analysis ###
#load the results from the Selection Analysis
Conserved_Selection_results_numbers <- read.table("/home/cropbio/sciebo/Documents/PhD/Brassicaceae-drought/BR-dr-03/Selection/Analysis/results_20221027_Conserved_Dataset/OGs_number_of_branches_under_pos_selection.tsv", sep = "\t", header=F)
names(Conserved_Selection_results_numbers) <- c("HOG", "number_of_pos_sel")
Conserved_Selection_results_genes_pvalues <- read.table("/home/cropbio/sciebo/Documents/PhD/Brassicaceae-drought/BR-dr-03/Selection/Analysis/results_20221027_Conserved_Dataset/OGs_pos_selection_pvalues.tsv", sep = "\t", header=F)
names(Conserved_Selection_results_genes_pvalues) <- c("HOG", "gene", "uncorrected_pvalue", "selection_pvalue")

HOGs_in_results_selection_analysis <- Conserved_Selection_results_numbers$HOG
length(HOGs_in_results_selection_analysis)
# 13257
### merge the data frames from A2TEA analysis with the ones from the Selection Analysis
HOG_level_list$hypothesis_1_cons_with_numbers_pos_sel <- merge(HOG_level_list$hypothesis_1_cons, Conserved_Selection_results_numbers, by="HOG")
#put all data frames into list
df_list <- list(HOG_level_list$hypothesis_1_cons_with_numbers_pos_sel, Conserved_Selection_results_genes_pvalues_HOG_DE_At, Conserved_Selection_results_genes_pvalues_HOG_DE_Al, Conserved_Selection_results_genes_pvalues_HOG_DE_Es, Conserved_Selection_results_genes_pvalues_HOG_DE_Bn)
#merge all data frames in list
HOG_level_list$hypothesis_1_cons_with_numbers_pos_sel_per_species <- Reduce(function(x, y) merge(x, y, all=TRUE), df_list)


### Hypothesis 2
# 6.1)b)2) OGs in "Conserved"
HOGs <- HOG_level_list$hypothesis_1_cons$HOG
# 6b) OGs with at least 1 pos. sel. in "Conserved" OGs?
HOGs_pos_sel <- HOG_level_list$hypothesis_1_cons_with_numbers_pos_sel_per_species[!is.na(HOG_level_list$hypothesis_1_cons_with_numbers_pos_sel_per_species$number_of_pos_sel) & HOG_level_list$hypothesis_1_cons_with_numbers_pos_sel_per_species$number_of_pos_sel>=1,]$HOG
# 6.1)j)1) OGs with at least 1 pos. sel. from the Hypothesis 2 expanded species in "Conserved" OGs?
HOGs_pos_sel_Eutrema <- HOG_level_list$hypothesis_1_cons_with_numbers_pos_sel_per_species[HOG_level_list$hypothesis_1_cons_with_numbers_pos_sel_per_species$Eutrema_salsugineum_pos_sel>=1 & !is.na(HOG_level_list$hypothesis_1_cons_with_numbers_pos_sel_per_species$Eutrema_salsugineum_pos_sel)
                                                                                          ,]$HOG
# 6.1)i)2) Within "Conserved": OGs with "Hypothesis 2 expanded==yes"
HOGs_Eutrema_exp <- HOG_level_list$hypothesis_2_cons[HOG_level_list$hypothesis_2_cons$expansion == "yes",]$HOG
# 6i) Within "Conserved": OGs with at least 1 pos. sel. in "Hypothesis 2 expanded==yes" OGs?
HOGs_Eutrema_exp_pos_sel_any <- HOG_level_list$hypothesis_1_cons_with_numbers_pos_sel_per_species[HOG_level_list$hypothesis_1_cons_with_numbers_pos_sel_per_species$HOG %in% HOGs_Eutrema_exp
                                                                                                  &
                                                                                                    HOG_level_list$hypothesis_1_cons_with_numbers_pos_sel_per_species$number_of_pos_sel>=1 & !is.na(HOG_level_list$hypothesis_1_cons_with_numbers_pos_sel_per_species$number_of_pos_sel),]$HOG
# 6j) OGs with at least 1 pos. sel. from any of the expanded species in "Hypothesis 2 expanded==yes" OGs?
HOGs_Eutrema_exp_pos_sel_Eutrema <- HOG_level_list$hypothesis_1_cons_with_numbers_pos_sel_per_species[HOG_level_list$hypothesis_1_cons_with_numbers_pos_sel_per_species$HOG %in% HOGs_Eutrema_exp 
                                                                                                      & 
                                                                                                        HOG_level_list$hypothesis_1_cons_with_numbers_pos_sel_per_species$Eutrema_salsugineum_pos_sel>=1 & !is.na(HOG_level_list$hypothesis_1_cons_with_numbers_pos_sel_per_species$Eutrema_salsugineum_pos_sel) 
                                                                                                      ,]$HOG

##Eutrema pos. Sel. in Conserved

#Make a list of the OGs that are in your gene set
genesOfInterest <- as.character(HOGs_pos_sel_Eutrema)
#set the universe for GO analysis (map GO term IDs (right) to Gene IDs (left))
geneID2GO <- readMappings(file = "GO_annotations_Conserved.tsv", sep="\t")
geneUniverse <- names(geneID2GO)
#Then we need to tell TopGO where these interesting genes appear in the 'geneUniverse' vector:
geneList <- factor(as.integer(geneUniverse %in% genesOfInterest))
names(geneList) <- geneUniverse # The geneList object tells TopGO which genes in the gene universe are your genes of interest.
#Putting the data together into an R object: We need to put our data in an object of type 'topGOdata'. This will contain the list of genes of interest, the GO annotations, and the GO hierarchy itself.

# Make the analysis for GO = BP
myGOdata <- new("topGOdata", description="Eutrema pos. Sel. in Conserved", ontology="BP", allGenes=geneList,  annot = annFUN.gene2GO, gene2GO = geneID2GO) # The 'ontology' argument can be 'BP' (biological process), 'MF' (molecular function), or 'CC' (cellular component).
myGOdata
#Perform Fisher's exact test:
resultFisher <- runTest(myGOdata, algorithm="weight01", statistic="fisher")
resultFisher
#We can list the top significant results found:
allRes <- GenTable(myGOdata, classicFisher = resultFisher, orderBy = "resultFisher", ranksOf = "classicFisher", topNodes = 7)
allRes <- allRes[allRes$Significant>1,]
#And Save the table of sign. overrepresented GO terms
write.table(allRes, "Sig_GO_terms_BP_Eutrema_pos_sel_in_Conserved.txt", sep="\t", quote = F)

# Make the analysis for GO = MF
myGOdata <- new("topGOdata", description="Eutrema pos. Sel. in Conserved", ontology="MF", allGenes=geneList,  annot = annFUN.gene2GO, gene2GO = geneID2GO) # The 'ontology' argument can be 'BP' (biological process), 'MF' (molecular function), or 'CC' (cellular component).
myGOdata
#Perform Fisher's exact test:
resultFisher <- runTest(myGOdata, algorithm="weight01", statistic="fisher")
resultFisher
#We can list the top significant results found:
allRes <- GenTable(myGOdata, classicFisher = resultFisher, orderBy = "resultFisher", ranksOf = "classicFisher", topNodes = 7)
allRes <- allRes[allRes$Significant>1,]
#And Save the table of sign. overrepresented GO terms
write.table(allRes, "Sig_GO_terms_MF_Eutrema_pos_sel_in_Conserved.txt", sep="\t", quote = F)

# Make the analysis for GO = CC
myGOdata <- new("topGOdata", description="Eutrema pos. Sel. in Conserved", ontology="CC", allGenes=geneList,  annot = annFUN.gene2GO, gene2GO = geneID2GO) # The 'ontology' argument can be 'BP' (biological process), 'MF' (molecular function), or 'CC' (cellular component).
myGOdata
#Perform Fisher's exact test:
resultFisher <- runTest(myGOdata, algorithm="weight01", statistic="fisher")
resultFisher
#We can list the top significant results found:
allRes <- GenTable(myGOdata, classicFisher = resultFisher, orderBy = "resultFisher", ranksOf = "classicFisher", topNodes = 3)
allRes <- allRes[allRes$Significant>1,]
#And Save the table of sign. overrepresented GO terms
write.table(allRes, "Sig_GO_terms_CC_Eutrema_pos_sel_in_Conserved.txt", sep="\t", quote = F)

##Eutrema pos. Sel. AND exp. in Conserved

#Make a list of the OGs that are in your gene set
genesOfInterest <- as.character(HOGs_Eutrema_exp_pos_sel_Eutrema)
#set the universe for GO analysis (map GO term IDs (right) to Gene IDs (left))
geneID2GO <- readMappings(file = "GO_annotations_Conserved.tsv", sep="\t")
geneUniverse <- names(geneID2GO)
#Then we need to tell TopGO where these interesting genes appear in the 'geneUniverse' vector:
geneList <- factor(as.integer(geneUniverse %in% genesOfInterest))
names(geneList) <- geneUniverse # The geneList object tells TopGO which genes in the gene universe are your genes of interest.
#Putting the data together into an R object: We need to put our data in an object of type 'topGOdata'. This will contain the list of genes of interest, the GO annotations, and the GO hierarchy itself.

# Make the analysis for GO = BP
myGOdata <- new("topGOdata", description="Eutrema pos. Sel. and exp. in Conserved", ontology="BP", allGenes=geneList,  annot = annFUN.gene2GO, gene2GO = geneID2GO) # The 'ontology' argument can be 'BP' (biological process), 'MF' (molecular function), or 'CC' (cellular component).
myGOdata
#Perform Fisher's exact test:
resultFisher <- runTest(myGOdata, algorithm="weight01", statistic="fisher")
resultFisher
#We can list the top significant results found:
allRes <- GenTable(myGOdata, classicFisher = resultFisher, orderBy = "resultFisher", ranksOf = "classicFisher", topNodes = 3)
allRes <- allRes[allRes$Significant>1,]
#And Save the table of sign. overrepresented GO terms
write.table(allRes, "Sig_GO_terms_BP_Eutrema_pos_sel_and_exp_in_Conserved.txt", sep="\t", quote = F)

# Make the analysis for GO = MF
myGOdata <- new("topGOdata", description="Eutrema pos. Sel. and exp. in Conserved", ontology="MF", allGenes=geneList,  annot = annFUN.gene2GO, gene2GO = geneID2GO) # The 'ontology' argument can be 'BP' (biological process), 'MF' (molecular function), or 'CC' (cellular component).
myGOdata
#Perform Fisher's exact test:
resultFisher <- runTest(myGOdata, algorithm="weight01", statistic="fisher")
resultFisher
#We can list the top significant results found:
allRes <- GenTable(myGOdata, classicFisher = resultFisher, orderBy = "resultFisher", ranksOf = "classicFisher", topNodes = 3)
allRes <- allRes[allRes$Significant>1,]
#And Save the table of sign. overrepresented GO terms
write.table(allRes, "Sig_GO_terms_MF_Eutrema_pos_sel_and_exp_in_Conserved.txt", sep="\t", quote = F)

# Make the analysis for GO = CC
myGOdata <- new("topGOdata", description="Eutrema pos. Sel. and exp. in Conserved", ontology="CC", allGenes=geneList,  annot = annFUN.gene2GO, gene2GO = geneID2GO) # The 'ontology' argument can be 'BP' (biological process), 'MF' (molecular function), or 'CC' (cellular component).
myGOdata
#Perform Fisher's exact test:
resultFisher <- runTest(myGOdata, algorithm="weight01", statistic="fisher")
resultFisher
#We can list the top significant results found:
allRes <- GenTable(myGOdata, classicFisher = resultFisher, orderBy = "resultFisher", ranksOf = "classicFisher", topNodes = 3)
allRes <- allRes[allRes$Significant>1,]
#And Save the table of sign. overrepresented GO terms
write.table(allRes, "Sig_GO_terms_CC_Eutrema_pos_sel_and_exp_in_Conserved.txt", sep="\t", quote = F)


# Eutrema pos. Sel. AND Eutrema DEG

  # 7.1)b)2) Number of OGs in "Conserved"
HOGs <- HOG_level_list$hypothesis_1_cons$HOG
# 7b) What is the frequency of OGs with at least 1 pos. Sel. in "Conserved" OGs?
HOGs_pos_sel <- HOG_level_list$hypothesis_1_cons_with_numbers_pos_sel_per_species[!is.na(HOG_level_list$hypothesis_1_cons_with_numbers_pos_sel_per_species$number_of_pos_sel) & HOG_level_list$hypothesis_1_cons_with_numbers_pos_sel_per_species$number_of_pos_sel>=1,]$HOG
# 7.1)j)1) What is the frequency of OGs with at least 1 pos. Sel. from any of the Hypothesis 2 expanded species in "Conserved" OGs?
HOGs_pos_sel_Eutrema <- HOG_level_list$hypothesis_1_cons_with_numbers_pos_sel_per_species[HOG_level_list$hypothesis_1_cons_with_numbers_pos_sel_per_species$Eutrema_salsugineum_pos_sel>=1 & !is.na(HOG_level_list$hypothesis_1_cons_with_numbers_pos_sel_per_species$Eutrema_salsugineum_pos_sel)
                                                                                          ,]$HOG


# 7.1)i)2) Within "Conserved": Number of OGs with Hypothesis 2 species DEG
HOGs_Eutrema_DEG <- HOG_level_list$hypothesis_1_cons_with_numbers_pos_sel_per_species[HOG_level_list$hypothesis_1_cons_with_numbers_pos_sel_per_species$Eutrema_salsugineum_sigDE >= 1
                                                                                      &
                                                                                        !is.na(HOG_level_list$hypothesis_1_cons_with_numbers_pos_sel_per_species$Eutrema_salsugineum_sigDE)
                                                                                      ,]$HOG
# 7i) Within "Conserved": What is the frequency of OGs with at least 1 pos. Sel. from any species in OGs with DEG from Hypothesis 2 expanded species?
HOGs_Eutrema_DEG_pos_sel_any <- HOG_level_list$hypothesis_1_cons_with_numbers_pos_sel_per_species[HOG_level_list$hypothesis_1_cons_with_numbers_pos_sel_per_species$Eutrema_salsugineum_sigDE >= 1
                                                                                                  &
                                                                                                    !is.na(HOG_level_list$hypothesis_1_cons_with_numbers_pos_sel_per_species$Eutrema_salsugineum_sigDE)
                                                                                                  & 
                                                                                                    !is.na(HOG_level_list$hypothesis_1_cons_with_numbers_pos_sel_per_species$number_of_pos_sel) & HOG_level_list$hypothesis_1_cons_with_numbers_pos_sel_per_species$number_of_pos_sel>=1,]$HOG

#7j) What is the frequency of OGs with at least 1 pos. sel. from any of the expanded species in Hypothesis 2 species DEG OGs?
HOGs_Eutrema_DEG_pos_sel_Eutrema <- HOG_level_list$hypothesis_1_cons_with_numbers_pos_sel_per_species[HOG_level_list$hypothesis_1_cons_with_numbers_pos_sel_per_species$Eutrema_salsugineum_sigDE >= 1
                                                                                                      &
                                                                                                        !is.na(HOG_level_list$hypothesis_1_cons_with_numbers_pos_sel_per_species$Eutrema_salsugineum_sigDE)
                                                                                                      & 
                                                                                                        HOG_level_list$hypothesis_1_cons_with_numbers_pos_sel_per_species$Eutrema_salsugineum_pos_sel>=1 
                                                                                                      &
                                                                                                        !is.na(HOG_level_list$hypothesis_1_cons_with_numbers_pos_sel_per_species$Eutrema_salsugineum_pos_sel) 
                                                                                                      ,]$HOG


#Make a list of the OGs that are in your gene set
genesOfInterest <- as.character(HOGs_Eutrema_DEG_pos_sel_Eutrema)
#set the universe for GO analysis (map GO term IDs (right) to Gene IDs (left))
geneID2GO <- readMappings(file = "GO_annotations_Conserved.tsv", sep="\t")
geneUniverse <- names(geneID2GO)
#Then we need to tell TopGO where these interesting genes appear in the 'geneUniverse' vector:
geneList <- factor(as.integer(geneUniverse %in% genesOfInterest))
names(geneList) <- geneUniverse # The geneList object tells TopGO which genes in the gene universe are your genes of interest.
#Putting the data together into an R object: We need to put our data in an object of type 'topGOdata'. This will contain the list of genes of interest, the GO annotations, and the GO hierarchy itself.

# Make the analysis for GO = BP
myGOdata <- new("topGOdata", description="Eutrema pos. Sel. and DEG Eutrema in Conserved", ontology="BP", allGenes=geneList,  annot = annFUN.gene2GO, gene2GO = geneID2GO) # The 'ontology' argument can be 'BP' (biological process), 'MF' (molecular function), or 'CC' (cellular component).
myGOdata
#Perform Fisher's exact test:
resultFisher <- runTest(myGOdata, algorithm="weight01", statistic="fisher")
resultFisher
#We can list the top significant results found:
allRes <- GenTable(myGOdata, classicFisher = resultFisher, orderBy = "resultFisher", ranksOf = "classicFisher", topNodes = 10)
allRes <- allRes[allRes$Significant>1,]
#And Save the table of sign. overrepresented GO terms
write.table(allRes, "Sig_GO_terms_BP_Eutrema_pos_sel_and_DEG_Eutrema_in_Conserved.txt", sep="\t", quote = F)

# Make the analysis for GO = MF
myGOdata <- new("topGOdata", description="Eutrema pos. Sel. and DEG Eutrema in Conserved", ontology="MF", allGenes=geneList,  annot = annFUN.gene2GO, gene2GO = geneID2GO) # The 'ontology' argument can be 'BP' (biological process), 'MF' (molecular function), or 'CC' (cellular component).
myGOdata
#Perform Fisher's exact test:
resultFisher <- runTest(myGOdata, algorithm="weight01", statistic="fisher")
resultFisher
#We can list the top significant results found:
allRes <- GenTable(myGOdata, classicFisher = resultFisher, orderBy = "resultFisher", ranksOf = "classicFisher", topNodes = 7)
allRes <- allRes[allRes$Significant>1,]
#And Save the table of sign. overrepresented GO terms
write.table(allRes, "Sig_GO_terms_MF_Eutrema_pos_sel_and_DEG_Eutrema_in_Conserved.txt", sep="\t", quote = F)

# Make the analysis for GO = CC
myGOdata <- new("topGOdata", description="Eutrema pos. Sel. and DEG Eutrema in Conserved", ontology="CC", allGenes=geneList,  annot = annFUN.gene2GO, gene2GO = geneID2GO) # The 'ontology' argument can be 'BP' (biological process), 'MF' (molecular function), or 'CC' (cellular component).
myGOdata
#Perform Fisher's exact test:
resultFisher <- runTest(myGOdata, algorithm="weight01", statistic="fisher")
resultFisher
#We can list the top significant results found:
allRes <- GenTable(myGOdata, classicFisher = resultFisher, orderBy = "resultFisher", ranksOf = "classicFisher", topNodes = 4)
allRes <- allRes[allRes$Significant>1,]
#And Save the table of sign. overrepresented GO terms
write.table(allRes, "Sig_GO_terms_CC_Eutrema_pos_sel_and_DEG_Eutrema_in_Conserved.txt", sep="\t", quote = F)


# Eutrema pos. Sel. AND exp. AND Eutrema DEG SAME GENE

### Hypothesis 2: Eutrema exp. and DEG

# 8.1)b)2) Number of OGs in "Conserved"
HOGs <- HOGs_in_results_selection_analysis
# 8b) What is the frequency of OGs with at least 1 pos. Sel. in "Conserved" OGs?
HOGs_pos_sel <- unique(Conserved_Selection_results_genes_pvalues_HOG_DE[!is.na(Conserved_Selection_results_genes_pvalues_HOG_DE$selection_pvalue) & (Conserved_Selection_results_genes_pvalues_HOG_DE$selection_pvalue <= 0.05),]$HOG)
# 8.1)d)1) What is the frequency of OGs with at least 1 pos. Sel. from Eutrema in "Conserved" OGs?
HOGs_pos_sel_Eutrema <- unique(Conserved_Selection_results_genes_pvalues_HOG_DE[!is.na(Conserved_Selection_results_genes_pvalues_HOG_DE$selection_pvalue) & (Conserved_Selection_results_genes_pvalues_HOG_DE$selection_pvalue <= 0.05)
                                                                                & Conserved_Selection_results_genes_pvalues_HOG_DE$species=="Eutrema_salsugineum"
                                                                                ,]$HOG)

# 8.1)c)2) Within "Conserved": Number of OGs with DEG from Eutrema
HOGs_Eutrema_DEG <- unique(Conserved_Selection_results_genes_pvalues_HOG_DE[!is.na(Conserved_Selection_results_genes_pvalues_HOG_DE$padj) & (Conserved_Selection_results_genes_pvalues_HOG_DE$padj <= 0.1)
                                                                            & Conserved_Selection_results_genes_pvalues_HOG_DE$species=="Eutrema_salsugineum"
                                                                            ,]$HOG)
# Expanded in Eutrema
HOGs_Eutrema_exp <- unique(Eutrema_exp$HOG)

# 8.1)c)2) Within "Conserved": Number of OGs with DEG from Eutrema in the in Eutrema expanded OGs
HOGs_Eutrema_DEG_and_exp_Eutrema <- intersect(HOGs_Eutrema_DEG, HOGs_Eutrema_exp)

# 8c) Within "Conserved": What is the frequency of OGs with at least 1 pos. Sel. in in Eutrema expanded OGs with Eutrema DEG?
HOGs_Eutrema_DEG_and_exp_Eutrema_pos_sel_any <- intersect(HOGs_Eutrema_DEG_and_exp_Eutrema, HOGs_pos_sel)
# 8d) What is the frequency of OGs with at least 1 pos. sel. from Eutrema in in Eutrema expanded OGs with Eutrema DEG?
HOGs_Eutrema_DEG_and_exp_Eutrema_pos_sel_Eutrema <- intersect(HOGs_Eutrema_DEG_and_exp_Eutrema, HOGs_pos_sel_Eutrema)

# More specific: The DE gene must be the pos. Sel. gene
HOGs_Eutrema_DEG_and_exp_Eutrema_pos_sel_same_gene_as_DEG <- intersect(
  Conserved_Selection_results_genes_pvalues_HOG_DE[!is.na(Conserved_Selection_results_genes_pvalues_HOG_DE$padj) & (Conserved_Selection_results_genes_pvalues_HOG_DE$padj <= 0.1)
                                                   & Conserved_Selection_results_genes_pvalues_HOG_DE$species=="Eutrema_salsugineum"
                                                   & !is.na(Conserved_Selection_results_genes_pvalues_HOG_DE$selection_pvalue) & (Conserved_Selection_results_genes_pvalues_HOG_DE$selection_pvalue <= 0.05)
                                                   ,]$HOG, 
  HOGs_Eutrema_exp)


#Make a list of the OGs that are in your gene set
genesOfInterest <- as.character(HOGs_Eutrema_DEG_and_exp_Eutrema_pos_sel_same_gene_as_DEG)
#set the universe for GO analysis (map GO term IDs (right) to Gene IDs (left))
geneID2GO <- readMappings(file = "GO_annotations_Conserved.tsv", sep="\t")
geneUniverse <- names(geneID2GO)
#Then we need to tell TopGO where these interesting genes appear in the 'geneUniverse' vector:
geneList <- factor(as.integer(geneUniverse %in% genesOfInterest))
names(geneList) <- geneUniverse # The geneList object tells TopGO which genes in the gene universe are your genes of interest.
#Putting the data together into an R object: We need to put our data in an object of type 'topGOdata'. This will contain the list of genes of interest, the GO annotations, and the GO hierarchy itself.

# Make the analysis for GO = BP
myGOdata <- new("topGOdata", description="Eutrema pos. Sel. and exp Eutrema and DEG Eutrema same gene in Conserved", ontology="BP", allGenes=geneList,  annot = annFUN.gene2GO, gene2GO = geneID2GO) # The 'ontology' argument can be 'BP' (biological process), 'MF' (molecular function), or 'CC' (cellular component).
myGOdata
#Perform Fisher's exact test:
resultFisher <- runTest(myGOdata, algorithm="weight01", statistic="fisher")
resultFisher
#We can list the top significant results found:
allRes <- GenTable(myGOdata, classicFisher = resultFisher, orderBy = "resultFisher", ranksOf = "classicFisher", topNodes = 29)
allRes <- allRes[allRes$Significant>1,]
#And Save the table of sign. overrepresented GO terms
write.table(allRes, "Sig_GO_terms_BP_Eutrema_pos_sel_and_exp_Eutrema_and_DEG_Eutrema_same_gene_in_Conserved.txt", sep="\t", quote = F)

### extract the OGs with the significant GO term annotated:
write.csv(intersect(unlist(genesInTerm(myGOdata)["GO:0042245"]), genesOfInterest), "HOGs_annotated_with_GO:0042245_RNA repair.csv")
#write.csv(intersect(unlist(genesInTerm(myGOdata)["GO:0048366"]), genesOfInterest), "HOGs_annotated_with_GO:0048366_leaf development.csv")
#write.csv(intersect(unlist(genesInTerm(myGOdata)["GO:0010182"]), genesOfInterest), "HOGs_annotated_with_GO:0010182_sugar mediated signaling pathway.csv")
# GO:0070734 histone H3-K27 methylation was excluded because according to REVIGO it is a molecular function and not a BP


# Make the analysis for GO = MF
myGOdata <- new("topGOdata", description="Eutrema pos. Sel. and exp Eutrema and DEG Eutrema same gene in Conserved", ontology="MF", allGenes=geneList,  annot = annFUN.gene2GO, gene2GO = geneID2GO) # The 'ontology' argument can be 'BP' (biological process), 'MF' (molecular function), or 'CC' (cellular component).
myGOdata
#Perform Fisher's exact test:
resultFisher <- runTest(myGOdata, algorithm="weight01", statistic="fisher")
resultFisher
#We can list the top significant results found:
allRes <- GenTable(myGOdata, classicFisher = resultFisher, orderBy = "resultFisher", ranksOf = "classicFisher", topNodes = 17)
allRes <- allRes[allRes$Significant>1,]
#And Save the table of sign. overrepresented GO terms
write.table(allRes, "Sig_GO_terms_MF_Eutrema_pos_sel_and_exp_Eutrema_and_DEG_Eutrema_same_gene_in_Conserved.txt", sep="\t", quote = F)

# Make the analysis for GO = CC
myGOdata <- new("topGOdata", description="Eutrema pos. Sel. and exp Eutrema and DEG Eutrema same gene in Conserved", ontology="CC", allGenes=geneList,  annot = annFUN.gene2GO, gene2GO = geneID2GO) # The 'ontology' argument can be 'BP' (biological process), 'MF' (molecular function), or 'CC' (cellular component).
myGOdata
#Perform Fisher's exact test:
resultFisher <- runTest(myGOdata, algorithm="weight01", statistic="fisher")
resultFisher
#We can list the top significant results found:
allRes <- GenTable(myGOdata, classicFisher = resultFisher, orderBy = "resultFisher", ranksOf = "classicFisher", topNodes = 6)
allRes <- allRes[allRes$Significant>1,]
#And Save the table of sign. overrepresented GO terms
write.table(allRes, "Sig_GO_terms_CC_Eutrema_pos_sel_and_exp_Eutrema_and_DEG_Eutrema_same_gene_in_Conserved.txt", sep="\t", quote = F)


## 20240912 ##

################# GO term enrichment analysis of HOGs with Selection in Alyrata
### change because we have some more!!!

### Hypothesis 3
# 6.1)b)2) OGs in "Conserved"
HOGs <- HOG_level_list$hypothesis_1_cons$HOG
# 6b) OGs with at least 1 pos. sel. in "Conserved" OGs?
HOGs_pos_sel <- HOG_level_list$hypothesis_1_cons_with_numbers_pos_sel_per_species[!is.na(HOG_level_list$hypothesis_1_cons_with_numbers_pos_sel_per_species$number_of_pos_sel) & HOG_level_list$hypothesis_1_cons_with_numbers_pos_sel_per_species$number_of_pos_sel>=1,]$HOG
# 6.1)j)1) OGs with at least 1 pos. sel. from the Hypothesis 2 expanded species in "Conserved" OGs?
HOGs_pos_sel_Alyrata <- HOG_level_list$hypothesis_1_cons_with_numbers_pos_sel_per_species[HOG_level_list$hypothesis_1_cons_with_numbers_pos_sel_per_species$Arabidopsis_lyrata_pos_sel>=1 & !is.na(HOG_level_list$hypothesis_1_cons_with_numbers_pos_sel_per_species$Arabidopsis_lyrata_pos_sel)
                                                                                          ,]$HOG
# 6.1)i)2) Within "Conserved": OGs with "Hypothesis 3 expanded==yes"
HOGs_Alyrata_exp <- HOG_level_list$hypothesis_3_cons[HOG_level_list$hypothesis_3_cons$expansion == "yes",]$HOG
# 6i) Within "Conserved": OGs with at least 1 pos. sel. in "Hypothesis 3 expanded==yes" OGs?
HOGs_Alyrata_exp_pos_sel_any <- HOG_level_list$hypothesis_1_cons_with_numbers_pos_sel_per_species[HOG_level_list$hypothesis_1_cons_with_numbers_pos_sel_per_species$HOG %in% HOGs_Alyrata_exp
                                                                                                  &
                                                                                                    HOG_level_list$hypothesis_1_cons_with_numbers_pos_sel_per_species$number_of_pos_sel>=1 & !is.na(HOG_level_list$hypothesis_1_cons_with_numbers_pos_sel_per_species$number_of_pos_sel),]$HOG
# 6j) OGs with at least 1 pos. sel. from any of the expanded species in "Hypothesis 3 expanded==yes" OGs?
HOGs_Alyrata_exp_pos_sel_Alyrata <- HOG_level_list$hypothesis_1_cons_with_numbers_pos_sel_per_species[HOG_level_list$hypothesis_1_cons_with_numbers_pos_sel_per_species$HOG %in% HOGs_Alyrata_exp 
                                                                                                      & 
                                                                                                        HOG_level_list$hypothesis_1_cons_with_numbers_pos_sel_per_species$Arabidopsis_lyrata_pos_sel>=1 & !is.na(HOG_level_list$hypothesis_1_cons_with_numbers_pos_sel_per_species$Arabidopsis_lyrata_pos_sel) 
                                                                                                      ,]$HOG

##Test Alyrata pos. Sel. in Conserved

#Make a list of the OGs that are in your gene set
genesOfInterest <- as.character(HOGs_pos_sel_Alyrata)
#set the universe for GO analysis (map GO term IDs (right) to Gene IDs (left))
geneID2GO <- readMappings(file = "GO_annotations_Conserved.tsv", sep="\t")
geneUniverse <- names(geneID2GO)
#Then we need to tell TopGO where these interesting genes appear in the 'geneUniverse' vector:
geneList <- factor(as.integer(geneUniverse %in% genesOfInterest))
names(geneList) <- geneUniverse # The geneList object tells TopGO which genes in the gene universe are your genes of interest.
#Putting the data together into an R object: We need to put our data in an object of type 'topGOdata'. This will contain the list of genes of interest, the GO annotations, and the GO hierarchy itself.

# Make the analysis for GO = BP
myGOdata <- new("topGOdata", description="Alyrata pos. Sel. in Conserved", ontology="BP", allGenes=geneList,  annot = annFUN.gene2GO, gene2GO = geneID2GO) # The 'ontology' argument can be 'BP' (biological process), 'MF' (molecular function), or 'CC' (cellular component).
myGOdata
#Perform Fisher's exact test:
resultFisher <- runTest(myGOdata, algorithm="weight01", statistic="fisher")
resultFisher
#We can list the top significant results found:
allRes <- GenTable(myGOdata, classicFisher = resultFisher, orderBy = "resultFisher", ranksOf = "classicFisher", topNodes = 15)
allRes <- allRes[allRes$Significant>1,]
#And Save the table of sign. overrepresented GO terms
write.table(allRes, "Sig_GO_terms_BP_Alyrata_pos_sel_in_Conserved.txt", sep="\t", quote = F)

# Make the analysis for GO = MF
myGOdata <- new("topGOdata", description="Alyrata pos. Sel. in Conserved", ontology="MF", allGenes=geneList,  annot = annFUN.gene2GO, gene2GO = geneID2GO) # The 'ontology' argument can be 'BP' (biological process), 'MF' (molecular function), or 'CC' (cellular component).
myGOdata
#Perform Fisher's exact test:
resultFisher <- runTest(myGOdata, algorithm="weight01", statistic="fisher")
resultFisher
#We can list the top significant results found:
allRes <- GenTable(myGOdata, classicFisher = resultFisher, orderBy = "resultFisher", ranksOf = "classicFisher", topNodes = 15)
allRes <- allRes[allRes$Significant>1,]
#And Save the table of sign. overrepresented GO terms
write.table(allRes, "Sig_GO_terms_MF_Alyrata_pos_sel_in_Conserved.txt", sep="\t", quote = F)

# Make the analysis for GO = CC
myGOdata <- new("topGOdata", description="Alyrata pos. Sel. in Conserved", ontology="CC", allGenes=geneList,  annot = annFUN.gene2GO, gene2GO = geneID2GO) # The 'ontology' argument can be 'BP' (biological process), 'MF' (molecular function), or 'CC' (cellular component).
myGOdata
#Perform Fisher's exact test:
resultFisher <- runTest(myGOdata, algorithm="weight01", statistic="fisher")
resultFisher
#We can list the top significant results found:
allRes <- GenTable(myGOdata, classicFisher = resultFisher, orderBy = "resultFisher", ranksOf = "classicFisher", topNodes = 3)
allRes <- allRes[allRes$Significant>1,]
#And Save the table of sign. overrepresented GO terms
write.table(allRes, "Sig_GO_terms_CC_Alyrata_pos_sel_in_Conserved.txt", sep="\t", quote = F)

##Alyrata pos. Sel. AND exp. in Conserved

#Make a list of the OGs that are in your gene set
genesOfInterest <- as.character(HOGs_Alyrata_exp_pos_sel_Alyrata)
#set the universe for GO analysis (map GO term IDs (right) to Gene IDs (left))
geneID2GO <- readMappings(file = "GO_annotations_Conserved.tsv", sep="\t")
geneUniverse <- names(geneID2GO)
#Then we need to tell TopGO where these interesting genes appear in the 'geneUniverse' vector:
geneList <- factor(as.integer(geneUniverse %in% genesOfInterest))
names(geneList) <- geneUniverse # The geneList object tells TopGO which genes in the gene universe are your genes of interest.
#Putting the data together into an R object: We need to put our data in an object of type 'topGOdata'. This will contain the list of genes of interest, the GO annotations, and the GO hierarchy itself.

# Make the analysis for GO = BP
myGOdata <- new("topGOdata", description="Alyrata pos. Sel. and exp. in Conserved", ontology="BP", allGenes=geneList,  annot = annFUN.gene2GO, gene2GO = geneID2GO) # The 'ontology' argument can be 'BP' (biological process), 'MF' (molecular function), or 'CC' (cellular component).
myGOdata
#Perform Fisher's exact test:
resultFisher <- runTest(myGOdata, algorithm="weight01", statistic="fisher")
resultFisher
#We can list the top significant results found:
allRes <- GenTable(myGOdata, classicFisher = resultFisher, orderBy = "resultFisher", ranksOf = "classicFisher", topNodes = 4)
allRes <- allRes[allRes$Significant>1,]
#And Save the table of sign. overrepresented GO terms
write.table(allRes, "Sig_GO_terms_BP_Alyrata_pos_sel_and_exp_in_Conserved.txt", sep="\t", quote = F)

# Make the analysis for GO = MF
myGOdata <- new("topGOdata", description="Alyrata pos. Sel. and exp. in Conserved", ontology="MF", allGenes=geneList,  annot = annFUN.gene2GO, gene2GO = geneID2GO) # The 'ontology' argument can be 'BP' (biological process), 'MF' (molecular function), or 'CC' (cellular component).
myGOdata
#Perform Fisher's exact test:
resultFisher <- runTest(myGOdata, algorithm="weight01", statistic="fisher")
resultFisher
#We can list the top significant results found:
allRes <- GenTable(myGOdata, classicFisher = resultFisher, orderBy = "resultFisher", ranksOf = "classicFisher", topNodes = 3)
allRes <- allRes[allRes$Significant>1,]
#And Save the table of sign. overrepresented GO terms
write.table(allRes, "Sig_GO_terms_MF_Alyrata_pos_sel_and_exp_in_Conserved.txt", sep="\t", quote = F)

# Make the analysis for GO = CC
myGOdata <- new("topGOdata", description="Alyrata pos. Sel. and exp. in Conserved", ontology="CC", allGenes=geneList,  annot = annFUN.gene2GO, gene2GO = geneID2GO) # The 'ontology' argument can be 'BP' (biological process), 'MF' (molecular function), or 'CC' (cellular component).
myGOdata
#Perform Fisher's exact test:
resultFisher <- runTest(myGOdata, algorithm="weight01", statistic="fisher")
resultFisher
#We can list the top significant results found:
allRes <- GenTable(myGOdata, classicFisher = resultFisher, orderBy = "resultFisher", ranksOf = "classicFisher", topNodes = 2)
allRes <- allRes[allRes$Significant>1,]
#And Save the table of sign. overrepresented GO terms
write.table(allRes, "Sig_GO_terms_CC_Alyrata_pos_sel_and_exp_in_Conserved.txt", sep="\t", quote = F)


# Alyrata pos. Sel. AND Alyrata DEG

# 7.1)b)2) Number of OGs in "Conserved"
HOGs <- HOG_level_list$hypothesis_1_cons$HOG
# 7b) What is the frequency of OGs with at least 1 pos. Sel. in "Conserved" OGs?
HOGs_pos_sel <- HOG_level_list$hypothesis_1_cons_with_numbers_pos_sel_per_species[!is.na(HOG_level_list$hypothesis_1_cons_with_numbers_pos_sel_per_species$number_of_pos_sel) & HOG_level_list$hypothesis_1_cons_with_numbers_pos_sel_per_species$number_of_pos_sel>=1,]$HOG
# 7.1)j)1) What is the frequency of OGs with at least 1 pos. Sel. from any of the Hypothesis 2 expanded species in "Conserved" OGs?
HOGs_pos_sel_Alyrata <- HOG_level_list$hypothesis_1_cons_with_numbers_pos_sel_per_species[HOG_level_list$hypothesis_1_cons_with_numbers_pos_sel_per_species$Arabidopsis_lyrata_pos_sel>=1 & !is.na(HOG_level_list$hypothesis_1_cons_with_numbers_pos_sel_per_species$Arabidopsis_lyrata_pos_sel)
                                                                                          ,]$HOG


# 7.1)i)2) Within "Conserved": Number of OGs with Hypothesis 3 species DEG
HOGsAlyrata_DEG <- HOG_level_list$hypothesis_1_cons_with_numbers_pos_sel_per_species[HOG_level_list$hypothesis_1_cons_with_numbers_pos_sel_per_species$Arabidopsis_lyrata_sigDE >= 1
                                                                                      &
                                                                                        !is.na(HOG_level_list$hypothesis_1_cons_with_numbers_pos_sel_per_species$Arabidopsis_lyrata_sigDE)
                                                                                      ,]$HOG
# 7i) Within "Conserved": What is the frequency of OGs with at least 1 pos. Sel. from any species in OGs with DEG from Hypothesis 3 expanded species?
HOGs_Alyrata_DEG_pos_sel_any <- HOG_level_list$hypothesis_1_cons_with_numbers_pos_sel_per_species[HOG_level_list$hypothesis_1_cons_with_numbers_pos_sel_per_species$Arabidopsis_lyrata_sigDE >= 1
                                                                                                  &
                                                                                                    !is.na(HOG_level_list$hypothesis_1_cons_with_numbers_pos_sel_per_species$Arabidopsis_lyrata_sigDE)
                                                                                                  & 
                                                                                                    !is.na(HOG_level_list$hypothesis_1_cons_with_numbers_pos_sel_per_species$number_of_pos_sel) & HOG_level_list$hypothesis_1_cons_with_numbers_pos_sel_per_species$number_of_pos_sel>=1,]$HOG

#7j) What is the frequency of OGs with at least 1 pos. sel. from any of the expanded species in Hypothesis 3 species DEG OGs?
HOGs_Alyrata_DEG_pos_sel_Alyrata <- HOG_level_list$hypothesis_1_cons_with_numbers_pos_sel_per_species[HOG_level_list$hypothesis_1_cons_with_numbers_pos_sel_per_species$Arabidopsis_lyrata_sigDE >= 1
                                                                                                      &
                                                                                                        !is.na(HOG_level_list$hypothesis_1_cons_with_numbers_pos_sel_per_species$Arabidopsis_lyrata_sigDE)
                                                                                                      & 
                                                                                                        HOG_level_list$hypothesis_1_cons_with_numbers_pos_sel_per_species$Arabidopsis_lyrata_pos_sel>=1 
                                                                                                      &
                                                                                                        !is.na(HOG_level_list$hypothesis_1_cons_with_numbers_pos_sel_per_species$Arabidopsis_lyrata_pos_sel) 
                                                                                                      ,]$HOG


#Make a list of the OGs that are in your gene set
genesOfInterest <- as.character(HOGs_Alyrata_DEG_pos_sel_Alyrata)
#set the universe for GO analysis (map GO term IDs (right) to Gene IDs (left))
geneID2GO <- readMappings(file = "GO_annotations_Conserved.tsv", sep="\t")
geneUniverse <- names(geneID2GO)
#Then we need to tell TopGO where these interesting genes appear in the 'geneUniverse' vector:
geneList <- factor(as.integer(geneUniverse %in% genesOfInterest))
names(geneList) <- geneUniverse # The geneList object tells TopGO which genes in the gene universe are your genes of interest.
#Putting the data together into an R object: We need to put our data in an object of type 'topGOdata'. This will contain the list of genes of interest, the GO annotations, and the GO hierarchy itself.

# Make the analysis for GO = BP
myGOdata <- new("topGOdata", description="Alyrata pos. Sel. and DEG Alyrata in Conserved", ontology="BP", allGenes=geneList,  annot = annFUN.gene2GO, gene2GO = geneID2GO) # The 'ontology' argument can be 'BP' (biological process), 'MF' (molecular function), or 'CC' (cellular component).
myGOdata
#Perform Fisher's exact test:
resultFisher <- runTest(myGOdata, algorithm="weight01", statistic="fisher")
resultFisher
#We can list the top significant results found:
allRes <- GenTable(myGOdata, classicFisher = resultFisher, orderBy = "resultFisher", ranksOf = "classicFisher", topNodes = 15)
allRes <- allRes[allRes$Significant>1,]
#And Save the table of sign. overrepresented GO terms
write.table(allRes, "Sig_GO_terms_BP_Alyrata_pos_sel_and_DEG_Alyrata_in_Conserved.txt", sep="\t", quote = F)

# Make the analysis for GO = MF
myGOdata <- new("topGOdata", description="Alyrata pos. Sel. and DEG Alyrata in Conserved", ontology="MF", allGenes=geneList,  annot = annFUN.gene2GO, gene2GO = geneID2GO) # The 'ontology' argument can be 'BP' (biological process), 'MF' (molecular function), or 'CC' (cellular component).
myGOdata
#Perform Fisher's exact test:
resultFisher <- runTest(myGOdata, algorithm="weight01", statistic="fisher")
resultFisher
#We can list the top significant results found:
allRes <- GenTable(myGOdata, classicFisher = resultFisher, orderBy = "resultFisher", ranksOf = "classicFisher", topNodes = 13)
allRes <- allRes[allRes$Significant>1,]
#And Save the table of sign. overrepresented GO terms
write.table(allRes, "Sig_GO_terms_MF_Alyrata_pos_sel_and_DEG_Alyrata_in_Conserved.txt", sep="\t", quote = F)

# Make the analysis for GO = CC
myGOdata <- new("topGOdata", description="Alyrata pos. Sel. and DEG Alyrata in Conserved", ontology="CC", allGenes=geneList,  annot = annFUN.gene2GO, gene2GO = geneID2GO) # The 'ontology' argument can be 'BP' (biological process), 'MF' (molecular function), or 'CC' (cellular component).
myGOdata
#Perform Fisher's exact test:
resultFisher <- runTest(myGOdata, algorithm="weight01", statistic="fisher")
resultFisher
#We can list the top significant results found:
allRes <- GenTable(myGOdata, classicFisher = resultFisher, orderBy = "resultFisher", ranksOf = "classicFisher", topNodes = 3)
allRes <- allRes[allRes$Significant>1,]
#And Save the table of sign. overrepresented GO terms
write.table(allRes, "Sig_GO_terms_CC_Alyrata_pos_sel_and_DEG_Alyrata_in_Conserved.txt", sep="\t", quote = F)


# Alyrata pos. Sel. AND exp. AND Alyrata DEG SAME GENE

### Hypothesis 3: Alyrata exp. and DEG

# 8.1)b)2) Number of OGs in "Conserved"
HOGs <- HOGs_in_results_selection_analysis
# 8b) What is the frequency of OGs with at least 1 pos. Sel. in "Conserved" OGs?
HOGs_pos_sel <- unique(Conserved_Selection_results_genes_pvalues_HOG_DE[!is.na(Conserved_Selection_results_genes_pvalues_HOG_DE$selection_pvalue) & (Conserved_Selection_results_genes_pvalues_HOG_DE$selection_pvalue <= 0.05),]$HOG)
# 8.1)d)1) What is the frequency of OGs with at least 1 pos. Sel. from Alyrata in "Conserved" OGs?
HOGs_pos_sel_Alyrata <- unique(Conserved_Selection_results_genes_pvalues_HOG_DE[!is.na(Conserved_Selection_results_genes_pvalues_HOG_DE$selection_pvalue) & (Conserved_Selection_results_genes_pvalues_HOG_DE$selection_pvalue <= 0.05)
                                                                                & Conserved_Selection_results_genes_pvalues_HOG_DE$species=="Arabidopsis_lyrata"
                                                                                ,]$HOG)

# 8.1)c)2) Within "Conserved": Number of OGs with DEG from Alyrata
HOGs_Alyrata_DEG <- unique(Conserved_Selection_results_genes_pvalues_HOG_DE[!is.na(Conserved_Selection_results_genes_pvalues_HOG_DE$padj) & (Conserved_Selection_results_genes_pvalues_HOG_DE$padj <= 0.1)
                                                                            & Conserved_Selection_results_genes_pvalues_HOG_DE$species=="Arabidopsis_lyrata"
                                                                            ,]$HOG)
# Expanded in Alyrata
HOGs_Alyrata_exp <- unique(Alyrata_exp$HOG)

# 8.1)c)2) Within "Conserved": Number of OGs with DEG from Alyrata in the in Alyrata expanded OGs
HOGs_Alyrata_DEG_and_exp_Alyrata <- intersect(HOGs_Alyrata_DEG, HOGs_Alyrata_exp)

# 8c) Within "Conserved": What is the frequency of OGs with at least 1 pos. Sel. in in Alyrata expanded OGs with Alyrata DEG?
HOGs_Alyrata_DEG_and_exp_Alyrata_pos_sel_any <- intersect(HOGs_Alyrata_DEG_and_exp_Alyrata, HOGs_pos_sel)
# 8d) What is the frequency of OGs with at least 1 pos. sel. from Alyrata in in Alyrata expanded OGs with Alyrata DEG?
HOGs_Alyrata_DEG_and_exp_Alyrata_pos_sel_Alyrata <- intersect(HOGs_Alyrata_DEG_and_exp_Alyrata, HOGs_pos_sel_Alyrata)

# More specific: The DE gene must be the pos. Sel. gene
HOGs_Alyrata_DEG_and_exp_Alyrata_pos_sel_same_gene_as_DEG <- intersect(
  Conserved_Selection_results_genes_pvalues_HOG_DE[!is.na(Conserved_Selection_results_genes_pvalues_HOG_DE$padj) & (Conserved_Selection_results_genes_pvalues_HOG_DE$padj <= 0.1)
                                                   & Conserved_Selection_results_genes_pvalues_HOG_DE$species=="Arabidopsis_lyrata"
                                                   & !is.na(Conserved_Selection_results_genes_pvalues_HOG_DE$selection_pvalue) & (Conserved_Selection_results_genes_pvalues_HOG_DE$selection_pvalue <= 0.05)
                                                   ,]$HOG, 
  HOGs_Alyrata_exp)


#Make a list of the OGs that are in your gene set
genesOfInterest <- as.character(HOGs_Alyrata_DEG_and_exp_Alyrata_pos_sel_same_gene_as_DEG)
#set the universe for GO analysis (map GO term IDs (right) to Gene IDs (left))
geneID2GO <- readMappings(file = "GO_annotations_Conserved.tsv", sep="\t")
geneUniverse <- names(geneID2GO)
#Then we need to tell TopGO where these interesting genes appear in the 'geneUniverse' vector:
geneList <- factor(as.integer(geneUniverse %in% genesOfInterest))
names(geneList) <- geneUniverse # The geneList object tells TopGO which genes in the gene universe are your genes of interest.
#Putting the data together into an R object: We need to put our data in an object of type 'topGOdata'. This will contain the list of genes of interest, the GO annotations, and the GO hierarchy itself.

# Make the analysis for GO = BP
myGOdata <- new("topGOdata", description="Alyrata pos. Sel. and exp Alyrata and DEG Alyrata same gene in Conserved", ontology="BP", allGenes=geneList,  annot = annFUN.gene2GO, gene2GO = geneID2GO) # The 'ontology' argument can be 'BP' (biological process), 'MF' (molecular function), or 'CC' (cellular component).
myGOdata
#Perform Fisher's exact test:
resultFisher <- runTest(myGOdata, algorithm="weight01", statistic="fisher")
resultFisher
#We can list the top significant results found:
allRes <- GenTable(myGOdata, classicFisher = resultFisher, orderBy = "resultFisher", ranksOf = "classicFisher", topNodes = 13)
allRes <- allRes[allRes$Significant>1,]
#And Save the table of sign. overrepresented GO terms
write.table(allRes, "Sig_GO_terms_BP_Alyrata_pos_sel_and_exp_Alyrata_and_DEG_Alyrata_same_gene_in_Conserved.txt", sep="\t", quote = F)

### extract the OGs with the significant GO terms annotated:
write.csv(intersect(unlist(genesInTerm(myGOdata)["GO:0045842"]), genesOfInterest), "HOGs_annotated_with_GO:0045842_positive regulation of mitotic metaphase anaphase transition.csv")
write.csv(intersect(unlist(genesInTerm(myGOdata)["GO:0106074"]), genesOfInterest), "HOGs_annotated_with_GO:0106074_aminoacyl-tRNA metabolism involved in translation.csv")
write.csv(intersect(unlist(genesInTerm(myGOdata)["GO:0031145"]), genesOfInterest), "HOGs_annotated_with_GO:0031145_anaphase-promoting complex-dependent catabolism.csv")
write.csv(intersect(unlist(genesInTerm(myGOdata)["GO:0006388"]), genesOfInterest), "HOGs_annotated_with_GO:0006388_tRNA splicing, via endonucleolytic cleav.csv")
write.csv(intersect(unlist(genesInTerm(myGOdata)["GO:0010206"]), genesOfInterest), "HOGs_annotated_with_GO:0010206_photosystem_II_repair.csv")


# Make the analysis for GO = MF
myGOdata <- new("topGOdata", description="Alyrata pos. Sel. and exp Alyrata and DEG Alyrata same gene in Conserved", ontology="MF", allGenes=geneList,  annot = annFUN.gene2GO, gene2GO = geneID2GO) # The 'ontology' argument can be 'BP' (biological process), 'MF' (molecular function), or 'CC' (cellular component).
myGOdata
#Perform Fisher's exact test:
resultFisher <- runTest(myGOdata, algorithm="weight01", statistic="fisher")
resultFisher
#We can list the top significant results found:
allRes <- GenTable(myGOdata, classicFisher = resultFisher, orderBy = "resultFisher", ranksOf = "classicFisher", topNodes = 13)
allRes <- allRes[allRes$Significant>1,]
#And Save the table of sign. overrepresented GO terms
write.table(allRes, "Sig_GO_terms_MF_Alyrata_pos_sel_and_exp_Alyrata_and_DEG_Alyrata_same_gene_in_Conserved.txt", sep="\t", quote = F)

# Make the analysis for GO = CC
myGOdata <- new("topGOdata", description="Alyrata pos. Sel. and exp Alyrata and DEG Alyrata same gene in Conserved", ontology="CC", allGenes=geneList,  annot = annFUN.gene2GO, gene2GO = geneID2GO) # The 'ontology' argument can be 'BP' (biological process), 'MF' (molecular function), or 'CC' (cellular component).
myGOdata
#Perform Fisher's exact test:
resultFisher <- runTest(myGOdata, algorithm="weight01", statistic="fisher")
resultFisher
#We can list the top significant results found:
allRes <- GenTable(myGOdata, classicFisher = resultFisher, orderBy = "resultFisher", ranksOf = "classicFisher", topNodes = 1)
allRes <- allRes[allRes$Significant>1,]
#And Save the table of sign. overrepresented GO terms
write.table(allRes, "Sig_GO_terms_CC_Alyrata_pos_sel_and_exp_Alyrata_and_DEG_Alyrata_same_gene_in_Conserved.txt", sep="\t", quote = F)
```
