## Additional File 3 for "Integration of differential expression under drought with gene family expansion unique to drought tolerant species predicts candidate genes for drought adaptation in Brassicaceae species": Analysis_expression_pattern_per_OG_only_at_least_2_sig.R

```
# script to explore a2tea output BR-dr-03
library(Biostrings)
library(ggplot2)
library(reshape2)
library(dplyr)
library(gridExtra)
library(RColorBrewer)

setwd("/home/cropbio/sciebo/Documents/PhD/Brassicaceae-drought/BR-dr-03")
OUTDIR=paste0(getwd(), "/Analysis/")
load("A2TEA_finished.RData")

#use the color palette Tyll used in the A2TEA publication
palette_OkabeIto_black <- c("#E69F00", "#56B4E9", "#009E73", "#F0E442", 
                           "#0072B2", "#D55E00", "#CC79A7", "#000000")

#### merge gene level expression data with hypotheses data

### Conserved
# Conserved HOGs
Conserved <- HOG_level_list$hypothesis_1[HOG_level_list$hypothesis_1$Arabidopsis_lyrata_total>=1 & HOG_level_list$hypothesis_1$Arabidopsis_thaliana_total>=1 & HOG_level_list$hypothesis_1$Eutrema_salsugineum_total>=1 & HOG_level_list$hypothesis_1$Brassica_napus_total>=1,]
Conserved_with_expression <- merge((Conserved[,-c(6)]), HOG_DE.a2tea, by="HOG")

## Select only OGs with at least 2 expressed Genes! Threshold for "expressed" is 10

Conserved_with_expression_at_least_2_expressed <- 
  Conserved_with_expression %>%
  group_by(HOG) %>%
  mutate(two_expressed=case_when(sum(na.omit(baseMean) >= 10)>=2  ~ "yes",
                                 TRUE ~ "no"))

## Select only OGs with at least 2 DEG Genes! Threshold for "DEG" is padj<=0.1

Conserved_with_expression_at_least_2_expressed_and_at_least_2_DEG <- 
  Conserved_with_expression_at_least_2_expressed %>%
  group_by(HOG) %>%
  mutate(two_DEG=case_when(sum(na.omit(padj) <= 0.1)>=2  ~ "yes",
                                 TRUE ~ "no"))
### How Many HOGs have at least 2 expressed and at least 2 DE genes?
### length(unique(Conserved_with_expression_at_least_2_expressed_and_at_least_2_DEG[Conserved_with_expression_at_least_2_expressed_and_at_least_2_DEG$two_DEG=="yes" & Conserved_with_expression_at_least_2_expressed_and_at_least_2_DEG$two_expressed=="yes",]$HOG))
# 8704
### what difference does it make to include also HOGs which do NOT have at least 2 expressed genes?
### == How many HOGs have at least 2 DEGs?
### length(unique(Conserved_with_expression_at_least_2_expressed_and_at_least_2_DEG[Conserved_with_expression_at_least_2_expressed_and_at_least_2_DEG$two_DEG=="yes",]$HOG))
# 8800

### This here is over all 4 species. More downstream in this script, I performed the analysis also per species and plotted in a UpSetR plot

# For each HOG, the value of the new column "category_of_regulation" is -1 where all values of "log2FoldChange" are <0 for this HOG
# and select only those gene families which contain at least 1 DEG

Conserved_with_expression_at_least_2_expressed_and_at_least_2_DEG_and_category <-
  Conserved_with_expression_at_least_2_expressed_and_at_least_2_DEG[Conserved_with_expression_at_least_2_expressed_and_at_least_2_DEG$two_expressed=="yes" & Conserved_with_expression_at_least_2_expressed_and_at_least_2_DEG$two_DEG=="yes" & !is.na(Conserved_with_expression_at_least_2_expressed$log2FoldChange) & !is.na(Conserved_with_expression_at_least_2_expressed$padj) & Conserved_with_expression_at_least_2_expressed$padj<=0.1,] %>%
  group_by(HOG) %>%
  mutate(category_of_regulation=case_when(all(log2FoldChange < 0) ~ "down" ,
                                          all(log2FoldChange > 0) ~ "up",
                                          any(log2FoldChange < 0) && any(log2FoldChange > 0) ~ "down and up"))

Conserved_with_expression_at_least_2_expressed_and_at_least_2_DEG_and_category_unique_HOGs <-
  Conserved_with_expression_at_least_2_expressed_and_at_least_2_DEG[Conserved_with_expression_at_least_2_expressed_and_at_least_2_DEG$two_expressed=="yes" & Conserved_with_expression_at_least_2_expressed_and_at_least_2_DEG$two_DEG=="yes" & !is.na(Conserved_with_expression_at_least_2_expressed$log2FoldChange) & !is.na(Conserved_with_expression_at_least_2_expressed$padj) & Conserved_with_expression_at_least_2_expressed$padj<=0.1,] %>%
  group_by(HOG) %>%
  summarize(category_of_regulation=case_when(all(log2FoldChange < 0) ~ "down" ,
                                          all(log2FoldChange > 0) ~ "up",
                                          any(log2FoldChange < 0) && any(log2FoldChange > 0) ~ "down and up"))

### length(unique(Conserved_with_expression_at_least_2_expressed_and_at_least_2_DEG_and_category_unique_HOGs$HOG))
# 8704

### -> why are some OGs lost when adding the categories??? Because: From 8800 (without selecting two_expressed!!!) to only 8704 OGs (with 2 expressed and 2 diff. exp.).

#### 20240206 ####### see what difference it makes when not excluding the HOGs which do not have at least 2 expressed genes:
Not_min_2_expr_Conserved_with_expression_at_least_2_expressed_and_at_least_2_DEG_and_category <-
  Conserved_with_expression_at_least_2_expressed_and_at_least_2_DEG[Conserved_with_expression_at_least_2_expressed_and_at_least_2_DEG$two_DEG=="yes" & !is.na(Conserved_with_expression_at_least_2_expressed$log2FoldChange) & !is.na(Conserved_with_expression_at_least_2_expressed$padj) & Conserved_with_expression_at_least_2_expressed$padj<=0.1,] %>%
  group_by(HOG) %>%
  mutate(category_of_regulation=case_when(all(log2FoldChange < 0) ~ "down" ,
                                          all(log2FoldChange > 0) ~ "up",
                                          any(log2FoldChange < 0) && any(log2FoldChange > 0) ~ "down and up"))

Not_min_2_expr_Conserved_with_expression_at_least_2_expressed_and_at_least_2_DEG_and_category_unique_HOGs <-
  Conserved_with_expression_at_least_2_expressed_and_at_least_2_DEG[Conserved_with_expression_at_least_2_expressed_and_at_least_2_DEG$two_DEG=="yes" & !is.na(Conserved_with_expression_at_least_2_expressed$log2FoldChange) & !is.na(Conserved_with_expression_at_least_2_expressed$padj) & Conserved_with_expression_at_least_2_expressed$padj<=0.1,] %>%
  group_by(HOG) %>%
  summarize(category_of_regulation=case_when(all(log2FoldChange < 0) ~ "down" ,
                                             all(log2FoldChange > 0) ~ "up",
                                             any(log2FoldChange < 0) && any(log2FoldChange > 0) ~ "down and up"))

### Conserved Set
title="HOGs_Conserved"
a <- ggplot(Not_min_2_expr_Conserved_with_expression_at_least_2_expressed_and_at_least_2_DEG_and_category_unique_HOGs, aes(x="", fill=as.factor(category_of_regulation)))+
  geom_bar(width=1)+
  ggtitle(title)+
  coord_polar("y")+
  scale_fill_manual(values = palette_OkabeIto_black[1:3])
a
#### 20240206 ####### does not make a difference... I tested the same for all subsets (see downstream of the main analysis)
## and I used the calculations and plot which DO NOT take into account the basemean for the publication, 
# so switch to second part of the script which starts with: #### 20240206 ####### I tested all subsets if selecting or not "min 2 expr" with a baseMean threshold of 10...


#### Plot the distribution of HOGs with either up- / down- or both-regulated genes under drought ####

### Select only conserved HOGs (at least 1 gene from each (of the compared) species)
HOG_level_list$hypothesis_1_cons <- HOG_level_list$hypothesis_1[HOG_level_list$hypothesis_1$Arabidopsis_lyrata_total>=1 & HOG_level_list$hypothesis_1$Arabidopsis_thaliana_total>=1 & HOG_level_list$hypothesis_1$Eutrema_salsugineum_total>=1 & HOG_level_list$hypothesis_1$Brassica_napus_total>=1,]
HOG_level_list$hypothesis_2_cons <- HOG_level_list$hypothesis_2[HOG_level_list$hypothesis_2$HOG %in% HOG_level_list$hypothesis_1_cons$HOG,]
HOG_level_list$hypothesis_3_cons <- HOG_level_list$hypothesis_3[HOG_level_list$hypothesis_3$HOG %in% HOG_level_list$hypothesis_1_cons$HOG,]
HOG_level_list$hypothesis_4_cons <- HOG_level_list$hypothesis_4[HOG_level_list$hypothesis_4$HOG %in% HOG_level_list$hypothesis_1_cons$HOG,]
HOG_level_list$hypothesis_5_cons <- HOG_level_list$hypothesis_5[HOG_level_list$hypothesis_5$HOG %in% HOG_level_list$hypothesis_1_cons$HOG,]
HOG_level_list$hypothesis_6_cons <- HOG_level_list$hypothesis_6[HOG_level_list$hypothesis_6$HOG %in% HOG_level_list$hypothesis_1_cons$HOG,]


### Conserved Set with DEG from any species
# 2b) What is the frequency of OGs with at least 1 DEG in "Conserved" OGs?
HOGs_DEG <- HOG_level_list$hypothesis_1_cons[!is.na(HOG_level_list$hypothesis_1_cons$total_sigDE) & HOG_level_list$hypothesis_1_cons$total_sigDE>=1,]$HOG

### change the dataset to plot here ###
dataset=HOGs_DEG
title="HOGs_Conserved_DEG"
pdf(paste0(OUTDIR, "/Regulation_category_at_least_2_expressed_and_at_least_2_DEG_in_Conserved_set_only_padj_less_0.1.pdf"), 12, 8)
b <- ggplot(Conserved_with_expression_at_least_2_expressed_and_at_least_2_DEG_and_category_unique_HOGs[Conserved_with_expression_at_least_2_expressed_and_at_least_2_DEG_and_category_unique_HOGs$HOG %in% dataset,], aes(x="", fill=as.factor(category_of_regulation)))+
  geom_bar(width=1)+
  ggtitle(title)+
  coord_polar("y")+
  scale_fill_brewer(palette = "Set2")
b
dev.off()

# how many are only up?
length(Conserved_with_expression_at_least_2_expressed_and_at_least_2_DEG_and_category_unique_HOGs[Conserved_with_expression_at_least_2_expressed_and_at_least_2_DEG_and_category_unique_HOGs$HOG %in% dataset & Conserved_with_expression_at_least_2_expressed_and_at_least_2_DEG_and_category_unique_HOGs$category_of_regulation=="up",]$HOG)
# 2722

# from how many categorized HOGs in total?
length(unique(Conserved_with_expression_at_least_2_expressed_and_at_least_2_DEG_and_category_unique_HOGs[Conserved_with_expression_at_least_2_expressed_and_at_least_2_DEG_and_category_unique_HOGs$HOG %in% dataset,]$HOG))
# 8704

# 2722 / 8704 = 0.318


### Hypothesis 1

# 2.1)d)1) What is the frequency of OGs with at least 1 DEG from any of the Hypothesis 1 expanded species in "Conserved" OGs?
HOGs_DEG_any_tol <- HOG_level_list$hypothesis_1_cons[HOG_level_list$hypothesis_1_cons$Arabidopsis_lyrata_sigDE>=1 & !is.na(HOG_level_list$hypothesis_1_cons$Arabidopsis_lyrata_sigDE)
                                                     |
                                                       HOG_level_list$hypothesis_1_cons$Eutrema_salsugineum_sigDE>=1 & !is.na(HOG_level_list$hypothesis_1_cons$Eutrema_salsugineum_sigDE)
                                                     ,]$HOG

### change the dataset to plot here ###
dataset=HOGs_DEG_any_tol
title="HOGs_DEG_any_tol"
a <- ggplot(Conserved_with_expression_at_least_2_expressed_and_at_least_2_DEG_and_category_unique_HOGs[Conserved_with_expression_at_least_2_expressed_and_at_least_2_DEG_and_category_unique_HOGs$HOG %in% dataset,], aes(x="", fill=as.factor(category_of_regulation)))+
  geom_bar(width=1)+
  theme(legend.position="none")+
  ggtitle(title)+
  coord_polar("y")
a

# 2.1)e)1) What is the frequency of OGs with at least 1 DEG from both of the Hypothesis 1 expanded species in "Conserved" OGs?
HOGs_DEG_both_tol <- HOG_level_list$hypothesis_1_cons[HOG_level_list$hypothesis_1_cons$Arabidopsis_lyrata_sigDE>=1 & !is.na(HOG_level_list$hypothesis_1_cons$Arabidopsis_lyrata_sigDE)
                                                      &
                                                        HOG_level_list$hypothesis_1_cons$Eutrema_salsugineum_sigDE>=1 & !is.na(HOG_level_list$hypothesis_1_cons$Eutrema_salsugineum_sigDE)
                                                      ,]$HOG

### change the dataset to plot here ###
dataset=HOGs_DEG_both_tol
title="HOGs_DEG_both_tol"
b <- ggplot(Conserved_with_expression_at_least_2_expressed_and_at_least_2_DEG_and_category_unique_HOGs[Conserved_with_expression_at_least_2_expressed_and_at_least_2_DEG_and_category_unique_HOGs$HOG %in% dataset,], aes(x="", fill=as.factor(category_of_regulation)))+
  geom_bar(width=1)+
  theme(legend.position="none")+
  ggtitle(title)+
  coord_polar("y")
b
# 2.1)c)2) Within "Conserved": Number of OGs with "Hypothesis 1 expanded==yes"
HOGs_all_tol_exp <- HOG_level_list$hypothesis_1_cons[HOG_level_list$hypothesis_1_cons$expansion == "yes",]$HOG

### change the dataset to plot here ###
dataset=HOGs_all_tol_exp
title="HOGs_all_tol_exp"
c <- ggplot(Conserved_with_expression_at_least_2_expressed_and_at_least_2_DEG_and_category_unique_HOGs[Conserved_with_expression_at_least_2_expressed_and_at_least_2_DEG_and_category_unique_HOGs$HOG %in% dataset,], aes(x="", fill=as.factor(category_of_regulation)))+
  geom_bar(width=1)+
  theme(legend.position="none")+
  ggtitle(title)+
  coord_polar("y")
c

# 2c) Within "Conserved": What is the frequency of OGs with at least 1 DEG in "Hypothesis 1 expanded==yes" OGs?
HOGs_all_tol_exp_DEG_any <- HOG_level_list$hypothesis_1_cons[HOG_level_list$hypothesis_1_cons$expansion == "yes" & !is.na(HOG_level_list$hypothesis_1_cons$total_sigDE) & HOG_level_list$hypothesis_1_cons$total_sigDE>=1,]$HOG

### change the dataset to plot here ###
dataset=HOGs_all_tol_exp_DEG_any
title="HOGs_all_tol_exp_DEG_any"
d <- ggplot(Conserved_with_expression_at_least_2_expressed_and_at_least_2_DEG_and_category_unique_HOGs[Conserved_with_expression_at_least_2_expressed_and_at_least_2_DEG_and_category_unique_HOGs$HOG %in% dataset,], aes(x="", fill=as.factor(category_of_regulation)))+
  geom_bar(width=1)+
  theme(legend.position="none")+
  ggtitle(title)+
  coord_polar("y")
d

# 2d) What is the frequency of OGs with at least 1 DEG from any of the expanded species in "Hypothesis 1 expanded==yes" OGs?
HOGs_all_tol_exp_DEG_any_tol <- HOG_level_list$hypothesis_1_cons[HOG_level_list$hypothesis_1_cons$expansion == "yes" & HOG_level_list$hypothesis_1_cons$Eutrema_salsugineum_sigDE>=1 & !is.na(HOG_level_list$hypothesis_1_cons$Eutrema_salsugineum_sigDE) 
                                                                 |
                                                                   HOG_level_list$hypothesis_1_cons$expansion == "yes" & HOG_level_list$hypothesis_1_cons$Arabidopsis_lyrata_sigDE>=1 & !is.na(HOG_level_list$hypothesis_1_cons$Arabidopsis_lyrata_sigDE)
                                                                 ,]$HOG

### change the dataset to plot here ###
dataset=HOGs_all_tol_exp_DEG_any_tol
title="HOGs_all_tol_exp_DEG_any_tol"
e <- ggplot(Conserved_with_expression_at_least_2_expressed_and_at_least_2_DEG_and_category_unique_HOGs[Conserved_with_expression_at_least_2_expressed_and_at_least_2_DEG_and_category_unique_HOGs$HOG %in% dataset,], aes(x="", fill=as.factor(category_of_regulation)))+
  geom_bar(width=1)+
  theme(legend.position="none")+
  ggtitle(title)+
  coord_polar("y")
e

# 2e) What is the frequency of OGs with at least 1 DEG from both of the expanded species in "Hypothesis 1 expanded==yes" OGs?
HOGs_all_tol_exp_DEG_both_tol <- HOG_level_list$hypothesis_1_cons[HOG_level_list$hypothesis_1_cons$expansion == "yes" 
                                                                  & HOG_level_list$hypothesis_1_cons$Eutrema_salsugineum_sigDE>=1 & !is.na(HOG_level_list$hypothesis_1_cons$Eutrema_salsugineum_sigDE) 
                                                                  & HOG_level_list$hypothesis_1_cons$Arabidopsis_lyrata_sigDE>=1 & !is.na(HOG_level_list$hypothesis_1_cons$Arabidopsis_lyrata_sigDE)
                                                                  ,]$HOG
### change the dataset and title here ###
dataset=HOGs_all_tol_exp_DEG_both_tol
title="HOGs_all_tol_exp_DEG_both_tol"
f <- ggplot(Conserved_with_expression_at_least_2_expressed_and_at_least_2_DEG_and_category_unique_HOGs[Conserved_with_expression_at_least_2_expressed_and_at_least_2_DEG_and_category_unique_HOGs$HOG %in% dataset,], aes(x="", fill=as.factor(category_of_regulation)))+
  geom_bar(width=1)+
  ggtitle(title)+
  coord_polar("y")
f

pdf(paste0(OUTDIR, "/Regulation_category_at_least_2_expressed_and_at_least_2_DEG_with_and_without_DEGs_in_Hypothesis_1_set_only_padj_less_0.1.pdf"), 12, 8)
ggarrange(a, b, c, d, e, f, ncol=3, nrow=2, common.legend = TRUE, legend="bottom")
dev.off()

# how many are only "up"?
length(Conserved_with_expression_at_least_2_expressed_and_at_least_2_DEG_and_category_unique_HOGs[Conserved_with_expression_at_least_2_expressed_and_at_least_2_DEG_and_category_unique_HOGs$HOG %in% dataset & Conserved_with_expression_at_least_2_expressed_and_at_least_2_DEG_and_category_unique_HOGs$category_of_regulation=="up",]$HOG)
# 24

# from how many categorized HOGs in total?
length(unique(Conserved_with_expression_at_least_2_expressed_and_at_least_2_DEG_and_category_unique_HOGs[Conserved_with_expression_at_least_2_expressed_and_at_least_2_DEG_and_category_unique_HOGs$HOG %in% dataset,]$HOG))
# 46

# 24 / 46 = 0.522

# test if this is a significant enrichment of HOGs which are only up regulated compared to the Conserved set:
q <- 24
m <- 2722
n <- (8704 - m)
k <- 46

p <- phyper((q-1), m, n, k, lower.tail=F)
p # 0.002491145


### Hypothesis 4
# 2.1)g)1) What is the frequency of OGs with at least 1 DEG from any of the Hypothesis 4 expanded species in "Conserved" OGs?
HOGs_DEG_any_sens <- HOG_level_list$hypothesis_4_cons[HOG_level_list$hypothesis_4_cons$Arabidopsis_thaliana_sigDE>=1 & !is.na(HOG_level_list$hypothesis_4_cons$Arabidopsis_thaliana_sigDE)
                                                     |
                                                     HOG_level_list$hypothesis_4_cons$Brassica_napus_sigDE>=1 & !is.na(HOG_level_list$hypothesis_4_cons$Brassica_napus_sigDE)
                                                     ,]$HOG

### change the dataset to plot here ###
dataset=HOGs_DEG_any_sens
title="HOGs_DEG_any_sens"
a <- ggplot(Conserved_with_expression_at_least_2_expressed_and_at_least_2_DEG_and_category_unique_HOGs[Conserved_with_expression_at_least_2_expressed_and_at_least_2_DEG_and_category_unique_HOGs$HOG %in% dataset,], aes(x="", fill=as.factor(category_of_regulation)))+
  geom_bar(width=1)+
  ggtitle(title)+
  coord_polar("y")

# 2.1)h)1) What is the frequency of OGs with at least 1 DEG from both of the Hypothesis 4 expanded species in "Conserved" OGs?
HOGs_DEG_both_sens <- HOG_level_list$hypothesis_4_cons[HOG_level_list$hypothesis_4_cons$Arabidopsis_thaliana_sigDE>=1 & !is.na(HOG_level_list$hypothesis_4_cons$Arabidopsis_thaliana_sigDE)
                                                      &
                                                      HOG_level_list$hypothesis_4_cons$Brassica_napus_sigDE>=1 & !is.na(HOG_level_list$hypothesis_4_cons$Brassica_napus_sigDE)
                                                      ,]$HOG
### change the dataset to plot here ###
dataset=HOGs_DEG_both_sens
title="HOGs_DEG_both_sens"
b <- ggplot(Conserved_with_expression_at_least_2_expressed_and_at_least_2_DEG_and_category_unique_HOGs[Conserved_with_expression_at_least_2_expressed_and_at_least_2_DEG_and_category_unique_HOGs$HOG %in% dataset,], aes(x="", fill=as.factor(category_of_regulation)))+
  geom_bar(width=1)+
  ggtitle(title)+
  coord_polar("y")

# 2.1)f)2) Within "Conserved": Number of OGs with "Hypothesis 4 expanded==yes"
HOGs_all_sens_exp <- HOG_level_list$hypothesis_4_cons[HOG_level_list$hypothesis_4_cons$expansion == "yes",]$HOG

### change the dataset to plot here ###
dataset=HOGs_all_sens_exp
title="HOGs_all_sens_exp"
c <- ggplot(Conserved_with_expression_at_least_2_expressed_and_at_least_2_DEG_and_category_unique_HOGs[Conserved_with_expression_at_least_2_expressed_and_at_least_2_DEG_and_category_unique_HOGs$HOG %in% dataset,], aes(x="", fill=as.factor(category_of_regulation)))+
  geom_bar(width=1)+
  ggtitle(title)+
  coord_polar("y")

# 2f) Within "Conserved": What is the frequency of OGs with at least 1 DEG in "Hypothesis 4 expanded==yes" OGs?
HOGs_all_sens_exp_DEG_any <- HOG_level_list$hypothesis_4_cons[HOG_level_list$hypothesis_4_cons$expansion == "yes" & !is.na(HOG_level_list$hypothesis_4_cons$total_sigDE) & HOG_level_list$hypothesis_4_cons$total_sigDE>=1,]$HOG

### change the dataset to plot here ###
dataset=HOGs_all_sens_exp_DEG_any
title="HOGs_all_sens_exp_DEG_any"
d <- ggplot(Conserved_with_expression_at_least_2_expressed_and_at_least_2_DEG_and_category_unique_HOGs[Conserved_with_expression_at_least_2_expressed_and_at_least_2_DEG_and_category_unique_HOGs$HOG %in% dataset,], aes(x="", fill=as.factor(category_of_regulation)))+
  geom_bar(width=1)+
  ggtitle(title)+
  coord_polar("y")

# 2g) What is the frequency of OGs with at least 1 DEG from any of the expanded species in "Hypothesis 4 expanded==yes" OGs?
HOGs_all_sens_exp_DEG_any_sens <- HOG_level_list$hypothesis_4_cons[HOG_level_list$hypothesis_4_cons$expansion == "yes" & HOG_level_list$hypothesis_4_cons$Arabidopsis_thaliana_sigDE>=1 & !is.na(HOG_level_list$hypothesis_4_cons$Arabidopsis_thaliana_sigDE) 
                                                                 |
                                                                 HOG_level_list$hypothesis_4_cons$expansion == "yes" & HOG_level_list$hypothesis_4_cons$Brassica_napus_sigDE>=1 & !is.na(HOG_level_list$hypothesis_4_cons$Brassica_napus_sigDE)
                                                                 ,]$HOG
### change the dataset to plot here ###
dataset=HOGs_all_sens_exp_DEG_any_sens
title="HOGs_all_sens_exp_DEG_any_sens"
e <- ggplot(Conserved_with_expression_at_least_2_expressed_and_at_least_2_DEG_and_category_unique_HOGs[Conserved_with_expression_at_least_2_expressed_and_at_least_2_DEG_and_category_unique_HOGs$HOG %in% dataset,], aes(x="", fill=as.factor(category_of_regulation)))+
  geom_bar(width=1)+
  ggtitle(title)+
  coord_polar("y")


# 2h) What is the frequency of OGs with at least 1 DEG from both of the expanded species in "Hypothesis 4 expanded==yes" OGs?
HOGs_all_sens_exp_DEG_both_sens <- HOG_level_list$hypothesis_4_cons[HOG_level_list$hypothesis_4_cons$expansion == "yes" 
                                                                  & HOG_level_list$hypothesis_4_cons$Arabidopsis_thaliana_sigDE>=1 & !is.na(HOG_level_list$hypothesis_4_cons$Arabidopsis_thaliana_sigDE) 
                                                                  & HOG_level_list$hypothesis_4_cons$Brassica_napus_sigDE>=1 & !is.na(HOG_level_list$hypothesis_4_cons$Brassica_napus_sigDE)
                                                                  ,]$HOG

### change the dataset and title here ###
dataset=HOGs_all_sens_exp_DEG_both_sens
title="HOGs_all_sens_exp_DEG_both_sens"
f <- ggplot(Conserved_with_expression_at_least_2_expressed_and_at_least_2_DEG_and_category_unique_HOGs[Conserved_with_expression_at_least_2_expressed_and_at_least_2_DEG_and_category_unique_HOGs$HOG %in% dataset,], aes(x="", fill=as.factor(category_of_regulation)))+
  geom_bar(width=1)+
  ggtitle(title)+
  coord_polar("y")

pdf(paste0(OUTDIR, "/Regulation_category_at_least_2_expressed_and_at_least_2_DEG_with_and_without_DEGs_in_Hypothesis_4_set_only_padj_less_0.1.pdf"), 12, 8)
ggarrange(a, b, c, d, e, f, ncol=3, nrow=2, common.legend = TRUE, legend="bottom")
dev.off()


### Hypothesis 2
# 2.1)j)1) What is the frequency of OGs with at least 1 DEG from the Hypothesis 2 expanded species in "Conserved" OGs?
HOGs_DEG_Eutrema <- HOG_level_list$hypothesis_2_cons[HOG_level_list$hypothesis_2_cons$Eutrema_salsugineum_sigDE>=1 & !is.na(HOG_level_list$hypothesis_2_cons$Eutrema_salsugineum_sigDE)
                                                      ,]$HOG
### change the dataset and title here ###
dataset=HOGs_DEG_Eutrema
title="HOGs_DEG_Eutrema"
a <- ggplot(Conserved_with_expression_at_least_2_expressed_and_at_least_2_DEG_and_category_unique_HOGs[Conserved_with_expression_at_least_2_expressed_and_at_least_2_DEG_and_category_unique_HOGs$HOG %in% dataset,], aes(x="", fill=as.factor(category_of_regulation)))+
  geom_bar(width=1)+
  ggtitle(title)+
  coord_polar("y")

# 2.1)i)2) Within "Conserved": Number of OGs with "Hypothesis 2 expanded==yes"
HOGs_Eutrema_exp <- HOG_level_list$hypothesis_2_cons[HOG_level_list$hypothesis_2_cons$expansion == "yes",]$HOG

### change the dataset and title here ###
dataset=HOGs_Eutrema_exp
title="HOGs_Eutrema_exp"
b <- ggplot(Conserved_with_expression_at_least_2_expressed_and_at_least_2_DEG_and_category_unique_HOGs[Conserved_with_expression_at_least_2_expressed_and_at_least_2_DEG_and_category_unique_HOGs$HOG %in% dataset,], aes(x="", fill=as.factor(category_of_regulation)))+
  geom_bar(width=1)+
  ggtitle(title)+
  coord_polar("y")

# 2i) Within "Conserved": What is the frequency of OGs with at least 1 DEG in "Hypothesis 2 expanded==yes" OGs?
cons_1_and_2_merged_2 <-  merge(HOG_level_list$hypothesis_1_cons[,c("HOG", "total_sigDE")], HOG_level_list$hypothesis_2_cons, by="HOG")
# select for total_sigDE.x because we are interested in total_sigDE from all 4 species (inferred from data frame HOG_level_list$hypothesis_1_cons, which contains all 4 species)
HOGs_Eutrema_exp_DEG_any <- cons_1_and_2_merged_2[cons_1_and_2_merged_2$expansion == "yes" & !is.na(cons_1_and_2_merged_2$total_sigDE.x) & cons_1_and_2_merged_2$total_sigDE.x>=1,]$HOG

### change the dataset and title here ###
dataset=HOGs_Eutrema_exp_DEG_any
title="HOGs_Eutrema_exp_DEG_any"
c <- ggplot(Conserved_with_expression_at_least_2_expressed_and_at_least_2_DEG_and_category_unique_HOGs[Conserved_with_expression_at_least_2_expressed_and_at_least_2_DEG_and_category_unique_HOGs$HOG %in% dataset,], aes(x="", fill=as.factor(category_of_regulation)))+
  geom_bar(width=1)+
  ggtitle(title)+
  coord_polar("y")

# 2j) What is the frequency of OGs with at least 1 DEG from any of the expanded species in "Hypothesis 2 expanded==yes" OGs?
HOGs_Eutrema_exp_DEG_Eutrema <- HOG_level_list$hypothesis_2_cons[HOG_level_list$hypothesis_2_cons$expansion == "yes" & HOG_level_list$hypothesis_2_cons$Eutrema_salsugineum_sigDE>=1 & !is.na(HOG_level_list$hypothesis_2_cons$Eutrema_salsugineum_sigDE) 
                                                                ,]$HOG

### change the dataset and title here ###
dataset=HOGs_Eutrema_exp_DEG_Eutrema
title="HOGs_Eutrema_exp_DEG_Eutrema"
d <- ggplot(Conserved_with_expression_at_least_2_expressed_and_at_least_2_DEG_and_category_unique_HOGs[Conserved_with_expression_at_least_2_expressed_and_at_least_2_DEG_and_category_unique_HOGs$HOG %in% dataset,], aes(x="", fill=as.factor(category_of_regulation)))+
  geom_bar(width=1)+
  ggtitle(title)+
  coord_polar("y")

pdf(paste0(OUTDIR, "/Regulation_category_at_least_2_expressed_and_at_least_2_DEG_with_and_without_DEGs_in_Hypothesis_2_set_only_padj_less_0.1.pdf"), 12, 8)
ggarrange(a, b, c, d, ncol=2, nrow=2, common.legend = TRUE, legend="bottom")
dev.off()


### Hypothesis 3
# 2.1)l)1) What is the frequency of OGs with at least 1 DEG from the Hypothesis 3 expanded species in "Conserved" OGs?
HOGs_DEG_Alyrata <- HOG_level_list$hypothesis_3_cons[HOG_level_list$hypothesis_3_cons$Arabidopsis_lyrata_sigDE>=1 & !is.na(HOG_level_list$hypothesis_3_cons$Arabidopsis_lyrata_sigDE)
                                                     ,]$HOG

### change the dataset and title here ###
dataset=HOGs_DEG_Alyrata
title="HOGs_DEG_Alyrata"
a <- ggplot(Conserved_with_expression_at_least_2_expressed_and_at_least_2_DEG_and_category_unique_HOGs[Conserved_with_expression_at_least_2_expressed_and_at_least_2_DEG_and_category_unique_HOGs$HOG %in% dataset,], aes(x="", fill=as.factor(category_of_regulation)))+
  geom_bar(width=1)+
  ggtitle(title)+
  coord_polar("y")

# 2.1)k)2) Within "Conserved": Number of OGs with "Hypothesis 3 expanded==yes"
HOGs_Alyrata_exp <- HOG_level_list$hypothesis_3_cons[HOG_level_list$hypothesis_3_cons$expansion == "yes",]$HOG

### change the dataset and title here ###
dataset=HOGs_Alyrata_exp
title="HOGs_Alyrata_exp"
b <- ggplot(Conserved_with_expression_at_least_2_expressed_and_at_least_2_DEG_and_category_unique_HOGs[Conserved_with_expression_at_least_2_expressed_and_at_least_2_DEG_and_category_unique_HOGs$HOG %in% dataset,], aes(x="", fill=as.factor(category_of_regulation)))+
  geom_bar(width=1)+
  ggtitle(title)+
  coord_polar("y")

# 2k) Within "Conserved": What is the frequency of OGs with at least 1 DEG in "Hypothesis 3 expanded==yes" OGs?
cons_1_and_3_merged_2 <-  merge(HOG_level_list$hypothesis_1_cons[,c("HOG", "total_sigDE")], HOG_level_list$hypothesis_3_cons, by="HOG")
# select for total_sigDE.x because we are interested in total_sigDE from all 4 species (inferred from data frame HOG_level_list$hypothesis_1_cons, which contains all 4 species)
HOGs_Alyrata_exp_DEG_any <- cons_1_and_3_merged_2[cons_1_and_3_merged_2$expansion == "yes" & !is.na(cons_1_and_3_merged_2$total_sigDE.x) & cons_1_and_3_merged_2$total_sigDE.x>=1,]$HOG

### change the dataset and title here ###
dataset=HOGs_Alyrata_exp_DEG_any
title="HOGs_Alyrata_exp_DEG_any"
c <- ggplot(Conserved_with_expression_at_least_2_expressed_and_at_least_2_DEG_and_category_unique_HOGs[Conserved_with_expression_at_least_2_expressed_and_at_least_2_DEG_and_category_unique_HOGs$HOG %in% dataset,], aes(x="", fill=as.factor(category_of_regulation)))+
  geom_bar(width=1)+
  ggtitle(title)+
  coord_polar("y")

# 2l) What is the frequency of OGs with at least 1 DEG from any of the expanded species in "Hypothesis 3 expanded==yes" OGs?
HOGs_Alyrata_exp_DEG_Alyrata <- HOG_level_list$hypothesis_3_cons[HOG_level_list$hypothesis_3_cons$expansion == "yes" & HOG_level_list$hypothesis_3_cons$Arabidopsis_lyrata_sigDE>=1 & !is.na(HOG_level_list$hypothesis_3_cons$Arabidopsis_lyrata_sigDE) 
                                                                 ,]$HOG

### change the dataset and title here ###
dataset=HOGs_Alyrata_exp_DEG_Alyrata
title="HOGs_Alyrata_exp_DEG_Alyrata"
d <- ggplot(Conserved_with_expression_at_least_2_expressed_and_at_least_2_DEG_and_category_unique_HOGs[Conserved_with_expression_at_least_2_expressed_and_at_least_2_DEG_and_category_unique_HOGs$HOG %in% dataset,], aes(x="", fill=as.factor(category_of_regulation)))+
  geom_bar(width=1)+
  ggtitle(title)+
  coord_polar("y")

pdf(paste0(OUTDIR, "/Regulation_category_at_least_2_expressed_and_at_least_2_DEG_with_and_without_DEGs_in_Hypothesis_3_set_only_padj_less_0.1.pdf"), 12, 8)
ggarrange(a, b, c, d, ncol=2, nrow=2, common.legend = TRUE, legend="bottom")
dev.off()

### Hypothesis 4
# 2.1)g)1) What is the frequency of OGs with at least 1 DEG from any of the Hypothesis 4 expanded species in "Conserved" OGs?
HOGs_DEG_any_sens <- HOG_level_list$hypothesis_4_cons[HOG_level_list$hypothesis_4_cons$Arabidopsis_thaliana_sigDE>=1 & !is.na(HOG_level_list$hypothesis_4_cons$Arabidopsis_thaliana_sigDE)
                                                     |
                                                     HOG_level_list$hypothesis_4_cons$Brassica_napus_sigDE>=1 & !is.na(HOG_level_list$hypothesis_4_cons$Brassica_napus_sigDE)
                                                     ,]$HOG

### change the dataset to plot here ###
dataset=HOGs_DEG_any_sens
title="HOGs_DEG_any_sens"
a <- ggplot(Conserved_with_expression_at_least_2_expressed_and_at_least_2_DEG_and_category_unique_HOGs[Conserved_with_expression_at_least_2_expressed_and_at_least_2_DEG_and_category_unique_HOGs$HOG %in% dataset,], aes(x="", fill=as.factor(category_of_regulation)))+
  geom_bar(width=1)+
  ggtitle(title)+
  coord_polar("y")

# 2.1)h)1) What is the frequency of OGs with at least 1 DEG from both of the Hypothesis 4 expanded species in "Conserved" OGs?
HOGs_DEG_both_sens <- HOG_level_list$hypothesis_4_cons[HOG_level_list$hypothesis_4_cons$Arabidopsis_thaliana_sigDE>=1 & !is.na(HOG_level_list$hypothesis_4_cons$Arabidopsis_thaliana_sigDE)
                                                      &
                                                      HOG_level_list$hypothesis_4_cons$Brassica_napus_sigDE>=1 & !is.na(HOG_level_list$hypothesis_4_cons$Brassica_napus_sigDE)
                                                      ,]$HOG
### change the dataset to plot here ###
dataset=HOGs_DEG_both_sens
title="HOGs_DEG_both_sens"
b <- ggplot(Conserved_with_expression_at_least_2_expressed_and_at_least_2_DEG_and_category_unique_HOGs[Conserved_with_expression_at_least_2_expressed_and_at_least_2_DEG_and_category_unique_HOGs$HOG %in% dataset,], aes(x="", fill=as.factor(category_of_regulation)))+
  geom_bar(width=1)+
  ggtitle(title)+
  coord_polar("y")

# 2.1)f)2) Within "Conserved": Number of OGs with "Hypothesis 4 expanded==yes"
HOGs_all_sens_exp <- HOG_level_list$hypothesis_4_cons[HOG_level_list$hypothesis_4_cons$expansion == "yes",]$HOG

### change the dataset to plot here ###
dataset=HOGs_all_sens_exp
title="HOGs_all_sens_exp"
c <- ggplot(Conserved_with_expression_at_least_2_expressed_and_at_least_2_DEG_and_category_unique_HOGs[Conserved_with_expression_at_least_2_expressed_and_at_least_2_DEG_and_category_unique_HOGs$HOG %in% dataset,], aes(x="", fill=as.factor(category_of_regulation)))+
  geom_bar(width=1)+
  ggtitle(title)+
  coord_polar("y")

# 2f) Within "Conserved": What is the frequency of OGs with at least 1 DEG in "Hypothesis 4 expanded==yes" OGs?
HOGs_all_sens_exp_DEG_any <- HOG_level_list$hypothesis_4_cons[HOG_level_list$hypothesis_4_cons$expansion == "yes" & !is.na(HOG_level_list$hypothesis_4_cons$total_sigDE) & HOG_level_list$hypothesis_4_cons$total_sigDE>=1,]$HOG

### change the dataset to plot here ###
dataset=HOGs_all_sens_exp_DEG_any
title="HOGs_all_sens_exp_DEG_any"
d <- ggplot(Conserved_with_expression_at_least_2_expressed_and_at_least_2_DEG_and_category_unique_HOGs[Conserved_with_expression_at_least_2_expressed_and_at_least_2_DEG_and_category_unique_HOGs$HOG %in% dataset,], aes(x="", fill=as.factor(category_of_regulation)))+
  geom_bar(width=1)+
  ggtitle(title)+
  coord_polar("y")

# 2g) What is the frequency of OGs with at least 1 DEG from any of the expanded species in "Hypothesis 4 expanded==yes" OGs?
HOGs_all_sens_exp_DEG_any_sens <- HOG_level_list$hypothesis_4_cons[HOG_level_list$hypothesis_4_cons$expansion == "yes" & HOG_level_list$hypothesis_4_cons$Arabidopsis_thaliana_sigDE>=1 & !is.na(HOG_level_list$hypothesis_4_cons$Arabidopsis_thaliana_sigDE) 
                                                                 |
                                                                 HOG_level_list$hypothesis_4_cons$expansion == "yes" & HOG_level_list$hypothesis_4_cons$Brassica_napus_sigDE>=1 & !is.na(HOG_level_list$hypothesis_4_cons$Brassica_napus_sigDE)
                                                                 ,]$HOG
### change the dataset to plot here ###
dataset=HOGs_all_sens_exp_DEG_any_sens
title="HOGs_all_sens_exp_DEG_any_sens"
e <- ggplot(Conserved_with_expression_at_least_2_expressed_and_at_least_2_DEG_and_category_unique_HOGs[Conserved_with_expression_at_least_2_expressed_and_at_least_2_DEG_and_category_unique_HOGs$HOG %in% dataset,], aes(x="", fill=as.factor(category_of_regulation)))+
  geom_bar(width=1)+
  ggtitle(title)+
  coord_polar("y")


# 2h) What is the frequency of OGs with at least 1 DEG from both of the expanded species in "Hypothesis 4 expanded==yes" OGs?
HOGs_all_sens_exp_DEG_both_sens <- HOG_level_list$hypothesis_4_cons[HOG_level_list$hypothesis_4_cons$expansion == "yes" 
                                                                  & HOG_level_list$hypothesis_4_cons$Arabidopsis_thaliana_sigDE>=1 & !is.na(HOG_level_list$hypothesis_4_cons$Arabidopsis_thaliana_sigDE) 
                                                                  & HOG_level_list$hypothesis_4_cons$Brassica_napus_sigDE>=1 & !is.na(HOG_level_list$hypothesis_4_cons$Brassica_napus_sigDE)
                                                                  ,]$HOG

### change the dataset and title here ###
dataset=HOGs_all_sens_exp_DEG_both_sens
title="HOGs_all_sens_exp_DEG_both_sens"
f <- ggplot(Conserved_with_expression_at_least_2_expressed_and_at_least_2_DEG_and_category_unique_HOGs[Conserved_with_expression_at_least_2_expressed_and_at_least_2_DEG_and_category_unique_HOGs$HOG %in% dataset,], aes(x="", fill=as.factor(category_of_regulation)))+
  geom_bar(width=1)+
  ggtitle(title)+
  coord_polar("y")

pdf(paste0(OUTDIR, "/Regulation_category_at_least_2_expressed_and_at_least_2_DEG_with_and_without_DEGs_in_Hypothesis_4_set_only_padj_less_0.1.pdf"), 12, 8)
ggarrange(a, b, c, d, e, f, ncol=3, nrow=2, common.legend = TRUE, legend="bottom")
dev.off()


### Hypothesis 5
# 2.1)n)1) What is the frequency of OGs with at least 1 DEG from the Hypothesis 5 expanded species in "Conserved" OGs?
HOGs_DEG_Athaliana <- HOG_level_list$hypothesis_5_cons[HOG_level_list$hypothesis_5_cons$Arabidopsis_thaliana_sigDE>=1 & !is.na(HOG_level_list$hypothesis_5_cons$Arabidopsis_thaliana_sigDE)
                                                     ,]$HOG
### change the dataset and title here ###
dataset=HOGs_DEG_Athaliana
title="HOGs_DEG_Athaliana"
a <- ggplot(Conserved_with_expression_at_least_2_expressed_and_at_least_2_DEG_and_category_unique_HOGs[Conserved_with_expression_at_least_2_expressed_and_at_least_2_DEG_and_category_unique_HOGs$HOG %in% dataset,], aes(x="", fill=as.factor(category_of_regulation)))+
  geom_bar(width=1)+
  ggtitle(title)+
  coord_polar("y")

# 2.1)m)2) Within "Conserved": Number of OGs with "Hypothesis 5 expanded==yes"
HOGs_Athaliana_exp <- HOG_level_list$hypothesis_5_cons[HOG_level_list$hypothesis_5_cons$expansion == "yes",]$HOG

### change the dataset and title here ###
dataset=HOGs_Athaliana_exp
title="HOGs_Athaliana_exp"
b <- ggplot(Conserved_with_expression_at_least_2_expressed_and_at_least_2_DEG_and_category_unique_HOGs[Conserved_with_expression_at_least_2_expressed_and_at_least_2_DEG_and_category_unique_HOGs$HOG %in% dataset,], aes(x="", fill=as.factor(category_of_regulation)))+
  geom_bar(width=1)+
  ggtitle(title)+
  coord_polar("y")

# 2m) Within "Conserved": What is the frequency of OGs with at least 1 DEG in "Hypothesis 5 expanded==yes" OGs?
cons_1_and_5_merged_2 <-  merge(HOG_level_list$hypothesis_1_cons[,c("HOG", "total_sigDE")], HOG_level_list$hypothesis_5_cons, by="HOG")
# select for total_sigDE.x because we are interested in total_sigDE from all 4 species (inferred from data frame HOG_level_list$hypothesis_1_cons, which contains all 4 species)
HOGs_Athaliana_exp_DEG_any <- cons_1_and_5_merged_2[cons_1_and_5_merged_2$expansion == "yes" & !is.na(cons_1_and_5_merged_2$total_sigDE.x) & cons_1_and_5_merged_2$total_sigDE.x>=1,]$HOG

### change the dataset and title here ###
dataset=HOGs_Athaliana_exp_DEG_any
title="HOGs_Athaliana_exp_DEG_any"
c <- ggplot(Conserved_with_expression_at_least_2_expressed_and_at_least_2_DEG_and_category_unique_HOGs[Conserved_with_expression_at_least_2_expressed_and_at_least_2_DEG_and_category_unique_HOGs$HOG %in% dataset,], aes(x="", fill=as.factor(category_of_regulation)))+
  geom_bar(width=1)+
  ggtitle(title)+
  coord_polar("y")

# 2n) What is the frequency of OGs with at least 1 DEG from any of the expanded species in "Hypothesis 5 expanded==yes" OGs?
HOGs_Athaliana_exp_DEG_Athaliana <- HOG_level_list$hypothesis_5_cons[HOG_level_list$hypothesis_5_cons$expansion == "yes" & HOG_level_list$hypothesis_5_cons$Arabidopsis_thaliana_sigDE>=1 & !is.na(HOG_level_list$hypothesis_5_cons$Arabidopsis_thaliana_sigDE)
                                                                 ,]$HOG

### change the dataset and title here ###
dataset=HOGs_Athaliana_exp_DEG_Athaliana
title="HOGs_Athaliana_exp_DEG_Athaliana"
d <- ggplot(Conserved_with_expression_at_least_2_expressed_and_at_least_2_DEG_and_category_unique_HOGs[Conserved_with_expression_at_least_2_expressed_and_at_least_2_DEG_and_category_unique_HOGs$HOG %in% dataset,], aes(x="", fill=as.factor(category_of_regulation)))+
  geom_bar(width=1)+
  ggtitle(title)+
  coord_polar("y")

pdf(paste0(OUTDIR, "/Regulation_category_at_least_2_expressed_and_at_least_2_DEG_with_and_without_DEGs_in_Hypothesis_5_set_only_padj_less_0.1.pdf"), 12, 8)
ggarrange(a, b, c, d, ncol=2, nrow=2, common.legend = TRUE, legend="bottom")
dev.off()


### Hypothesis 6
# 2.1)p)1) What is the frequency of OGs with at least 1 DEG from the Hypothesis 6 expanded species in "Conserved" OGs?
HOGs_DEG_Bnapus <- HOG_level_list$hypothesis_6_cons[HOG_level_list$hypothesis_6_cons$Brassica_napus_sigDE>=1 & !is.na(HOG_level_list$hypothesis_6_cons$Brassica_napus_sigDE)
                                                       ,]$HOG

### change the dataset and title here ###
dataset=HOGs_DEG_Bnapus
title="HOGs_DEG_Bnapus"
a <- ggplot(Conserved_with_expression_at_least_2_expressed_and_at_least_2_DEG_and_category_unique_HOGs[Conserved_with_expression_at_least_2_expressed_and_at_least_2_DEG_and_category_unique_HOGs$HOG %in% dataset,], aes(x="", fill=as.factor(category_of_regulation)))+
  geom_bar(width=1)+
  ggtitle(title)+
  coord_polar("y")

# 2.1)o)2) Within "Conserved": Number of OGs with "Hypothesis 6 expanded==yes"
HOGs_Bnapus_exp <- HOG_level_list$hypothesis_6_cons[HOG_level_list$hypothesis_6_cons$expansion == "yes",]$HOG

### change the dataset and title here ###
dataset=HOGs_Bnapus_exp
title="HOGs_Bnapus_exp"
b <- ggplot(Conserved_with_expression_at_least_2_expressed_and_at_least_2_DEG_and_category_unique_HOGs[Conserved_with_expression_at_least_2_expressed_and_at_least_2_DEG_and_category_unique_HOGs$HOG %in% dataset,], aes(x="", fill=as.factor(category_of_regulation)))+
  geom_bar(width=1)+
  ggtitle(title)+
  coord_polar("y")

# 2o) Within "Conserved": What is the frequency of OGs with at least 1 DEG in "Hypothesis 6 expanded==yes" OGs?
cons_1_and_6_merged_2 <-  merge(HOG_level_list$hypothesis_1_cons[,c("HOG", "total_sigDE")], HOG_level_list$hypothesis_6_cons, by="HOG")
# select for total_sigDE.x because we are interested in total_sigDE from all 4 species (inferred from data frame HOG_level_list$hypothesis_1_cons, which contains all 4 species)
HOGs_Bnapus_exp_DEG_any <- cons_1_and_6_merged_2[cons_1_and_6_merged_2$expansion == "yes" & !is.na(cons_1_and_6_merged_2$total_sigDE.x) & cons_1_and_6_merged_2$total_sigDE.x>=1,]$HOG

### change the dataset and title here ###
dataset=HOGs_Bnapus_exp_DEG_any
title="HOGs_Bnapus_exp_DEG_any"
c <- ggplot(Conserved_with_expression_at_least_2_expressed_and_at_least_2_DEG_and_category_unique_HOGs[Conserved_with_expression_at_least_2_expressed_and_at_least_2_DEG_and_category_unique_HOGs$HOG %in% dataset,], aes(x="", fill=as.factor(category_of_regulation)))+
  geom_bar(width=1)+
  ggtitle(title)+
  coord_polar("y")

# 2p) What is the frequency of OGs with at least 1 DEG from any of the expanded species in "Hypothesis 6 expanded==yes" OGs?
HOGs_Bnapus_exp_DEG_Bnapus <- HOG_level_list$hypothesis_6_cons[HOG_level_list$hypothesis_6_cons$expansion == "yes" & HOG_level_list$hypothesis_6_cons$Brassica_napus_sigDE>=1 & !is.na(HOG_level_list$hypothesis_6_cons$Brassica_napus_sigDE)
                                                                     ,]$HOG
### change the dataset and title here ###
dataset=HOGs_Bnapus_exp_DEG_Bnapus
title="HOGs_Bnapus_exp_DEG_Bnapus"
d <- ggplot(Conserved_with_expression_at_least_2_expressed_and_at_least_2_DEG_and_category_unique_HOGs[Conserved_with_expression_at_least_2_expressed_and_at_least_2_DEG_and_category_unique_HOGs$HOG %in% dataset,], aes(x="", fill=as.factor(category_of_regulation)))+
  geom_bar(width=1)+
  ggtitle(title)+
  coord_polar("y")

pdf(paste0(OUTDIR, "/Regulation_category_at_least_2_expressed_and_at_least_2_DEG_with_and_without_DEGs_in_Hypothesis_6_set_only_padj_less_0.1.pdf"), 12, 8)
ggarrange(a, b, c, d, ncol=2, nrow=2, common.legend = TRUE, legend="bottom")
dev.off()


#### 20240206 ####### I tested all subsets if selecting or not "min 2 expr" with a baseMean threshold of 10
# makes a difference in the distribution of regulation categories
## --> it DOES NOT. 
###### IMPORTANT######### :
# I used the plots which do not take into account the "min 2 expressed" for the publication
### Conserved Set
title="HOGs_Conserved"
a <- ggplot(Not_min_2_expr_Conserved_with_expression_at_least_2_expressed_and_at_least_2_DEG_and_category_unique_HOGs, aes(x="", fill=as.factor(category_of_regulation)))+
  geom_bar(width=1)+
  ggtitle(title)+
  coord_polar("y")
a
### Conserved Set with DEG any
# 2b) What is the frequency of OGs with at least 1 DEG in "Conserved" OGs?
HOGs_DEG <- HOG_level_list$hypothesis_1_cons[!is.na(HOG_level_list$hypothesis_1_cons$total_sigDE) & HOG_level_list$hypothesis_1_cons$total_sigDE>=1,]$HOG

### change the dataset to plot here ###
dataset=HOGs_DEG
title="HOGs_Conserved_DEG"
pdf(paste0(OUTDIR, "/Regulation_category_ONLY_at_least_2_DEG_with_DEG_any_in_Conserved_set_only_padj_less_0.1.pdf"), 12, 8)
b <- ggplot(Not_min_2_expr_Conserved_with_expression_at_least_2_expressed_and_at_least_2_DEG_and_category_unique_HOGs[Not_min_2_expr_Conserved_with_expression_at_least_2_expressed_and_at_least_2_DEG_and_category_unique_HOGs$HOG %in% dataset,], aes(x="", fill=as.factor(category_of_regulation)))+
  geom_bar(width=1)+
  ggtitle(title)+
  coord_polar("y")+
  scale_fill_manual(values = palette_OkabeIto_black[1:3])
b
dev.off()

total=length(Not_min_2_expr_Conserved_with_expression_at_least_2_expressed_and_at_least_2_DEG_and_category_unique_HOGs$HOG)
geneset1 <- names(summary(as.factor(Not_min_2_expr_Conserved_with_expression_at_least_2_expressed_and_at_least_2_DEG_and_category_unique_HOGs$category_of_regulation)))
group1 <- c(total, total, total)
At1 <- as.integer(c(summary(as.factor(Not_min_2_expr_Conserved_with_expression_at_least_2_expressed_and_at_least_2_DEG_and_category_unique_HOGs$category_of_regulation))))
at <- data.frame(geneset1, group1, At1)
#make 2 stacked bar charts and then make the y-axis with polar coordinates with log10 x-axis
pdf(paste0(OUTDIR, "/Regulation_category_ONLY_at_least_2_DEG_in_Conserved_set_only_padj_less_0.1.pdf"), 12, 8)
a <- ggplot(at)+
  geom_bar(aes(x=log10(total)/2, y=At1, fill=geneset1), width = log10(total), position="fill", stat="identity", color="white", alpha=1)+
  coord_polar(theta = "y", start=0)+
  scale_y_continuous(name ="Conserved and at least 2 DEGs from Any Species", limits= NULL, breaks=NULL, expand = expansion(0))+
  scale_x_continuous(name ="Proportion of HOGs per regulation category", limits= c(0,log10(total)), breaks = NULL, expand = expansion(0))+
  labs(fill="Regulation category")+
  theme(
    legend.position = "right",
    panel.grid.major = element_line(linetype = "dotted", colour="white"),
    panel.background = element_rect(fill="transparent", colour="black"))+
  geom_label(aes(x=log10(total)-log10(total)/6, y=0,label=paste(total)), color="black", size=7)+
  scale_fill_manual(values = palette_OkabeIto_black[1:3])
a
dev.off()


# how many are only up?
length(Not_min_2_expr_Conserved_with_expression_at_least_2_expressed_and_at_least_2_DEG_and_category_unique_HOGs[Not_min_2_expr_Conserved_with_expression_at_least_2_expressed_and_at_least_2_DEG_and_category_unique_HOGs$HOG %in% dataset & Not_min_2_expr_Conserved_with_expression_at_least_2_expressed_and_at_least_2_DEG_and_category_unique_HOGs$category_of_regulation=="up",]$HOG)
# 2768

# from how many categorized HOGs in total?
length(unique(Not_min_2_expr_Conserved_with_expression_at_least_2_expressed_and_at_least_2_DEG_and_category_unique_HOGs[Not_min_2_expr_Conserved_with_expression_at_least_2_expressed_and_at_least_2_DEG_and_category_unique_HOGs$HOG %in% dataset,]$HOG))
# 8800

## --> little difference for Conserved Set (2768 vs. 2722 are only up), but the ratio between the regulation categories is the same in both cases
# from 8704 or 8800 categorized HOGs in total

# 2768 / 8800 = 0.315

### Hypothesis 1

# 2.1)d)1) What is the frequency of OGs with at least 1 DEG from any of the Hypothesis 1 expanded species in "Conserved" OGs?
HOGs_DEG_any_tol <- HOG_level_list$hypothesis_1_cons[HOG_level_list$hypothesis_1_cons$Arabidopsis_lyrata_sigDE>=1 & !is.na(HOG_level_list$hypothesis_1_cons$Arabidopsis_lyrata_sigDE)
                                                     |
                                                       HOG_level_list$hypothesis_1_cons$Eutrema_salsugineum_sigDE>=1 & !is.na(HOG_level_list$hypothesis_1_cons$Eutrema_salsugineum_sigDE)
                                                     ,]$HOG

### change the dataset to plot here ###
dataset=HOGs_DEG_any_tol
title="HOGs_DEG_any_tol"
a <- ggplot(Not_min_2_expr_Conserved_with_expression_at_least_2_expressed_and_at_least_2_DEG_and_category_unique_HOGs[Not_min_2_expr_Conserved_with_expression_at_least_2_expressed_and_at_least_2_DEG_and_category_unique_HOGs$HOG %in% dataset,], aes(x="", fill=as.factor(category_of_regulation)))+
  geom_bar(width=1)+
  theme(legend.position="none")+
  ggtitle(title)+
  coord_polar("y")+
  scale_fill_brewer(palette = "Set2")
a

# 2.1)e)1) What is the frequency of OGs with at least 1 DEG from both of the Hypothesis 1 expanded species in "Conserved" OGs?
HOGs_DEG_both_tol <- HOG_level_list$hypothesis_1_cons[HOG_level_list$hypothesis_1_cons$Arabidopsis_lyrata_sigDE>=1 & !is.na(HOG_level_list$hypothesis_1_cons$Arabidopsis_lyrata_sigDE)
                                                      &
                                                        HOG_level_list$hypothesis_1_cons$Eutrema_salsugineum_sigDE>=1 & !is.na(HOG_level_list$hypothesis_1_cons$Eutrema_salsugineum_sigDE)
                                                      ,]$HOG

### change the dataset to plot here ###
dataset=HOGs_DEG_both_tol
title="HOGs_DEG_both_tol"
b <- ggplot(Not_min_2_expr_Conserved_with_expression_at_least_2_expressed_and_at_least_2_DEG_and_category_unique_HOGs[Not_min_2_expr_Conserved_with_expression_at_least_2_expressed_and_at_least_2_DEG_and_category_unique_HOGs$HOG %in% dataset,], aes(x="", fill=as.factor(category_of_regulation)))+
  geom_bar(width=1)+
  theme(legend.position="none")+
  ggtitle(title)+
  coord_polar("y")+
  scale_fill_brewer(palette = "Set2")
b

# 2.1)c)2) Within "Conserved": Number of OGs with "Hypothesis 1 expanded==yes"
HOGs_all_tol_exp <- HOG_level_list$hypothesis_1_cons[HOG_level_list$hypothesis_1_cons$expansion == "yes",]$HOG

### change the dataset to plot here ###
dataset=HOGs_all_tol_exp
title="HOGs_all_tol_exp"
c <- ggplot(Not_min_2_expr_Conserved_with_expression_at_least_2_expressed_and_at_least_2_DEG_and_category_unique_HOGs[Not_min_2_expr_Conserved_with_expression_at_least_2_expressed_and_at_least_2_DEG_and_category_unique_HOGs$HOG %in% dataset,], aes(x="", fill=as.factor(category_of_regulation)))+
  geom_bar(width=1)+
  theme(legend.position="none")+
  ggtitle(title)+
  coord_polar("y")+
  scale_fill_brewer(palette = "Set2")
c

# 2c) Within "Conserved": What is the frequency of OGs with at least 1 DEG in "Hypothesis 1 expanded==yes" OGs?
HOGs_all_tol_exp_DEG_any <- HOG_level_list$hypothesis_1_cons[HOG_level_list$hypothesis_1_cons$expansion == "yes" & !is.na(HOG_level_list$hypothesis_1_cons$total_sigDE) & HOG_level_list$hypothesis_1_cons$total_sigDE>=1,]$HOG

### change the dataset to plot here ###
dataset=HOGs_all_tol_exp_DEG_any
title="HOGs_all_tol_exp_DEG_any"
d <- ggplot(Not_min_2_expr_Conserved_with_expression_at_least_2_expressed_and_at_least_2_DEG_and_category_unique_HOGs[Not_min_2_expr_Conserved_with_expression_at_least_2_expressed_and_at_least_2_DEG_and_category_unique_HOGs$HOG %in% dataset,], aes(x="", fill=as.factor(category_of_regulation)))+
  geom_bar(width=1)+
  theme(legend.position="none")+
  ggtitle(title)+
  coord_polar("y")+
  scale_fill_brewer(palette = "Set2")
d

# 2d) What is the frequency of OGs with at least 1 DEG from any of the expanded species in "Hypothesis 1 expanded==yes" OGs?
HOGs_all_tol_exp_DEG_any_tol <- HOG_level_list$hypothesis_1_cons[HOG_level_list$hypothesis_1_cons$expansion == "yes" & HOG_level_list$hypothesis_1_cons$Eutrema_salsugineum_sigDE>=1 & !is.na(HOG_level_list$hypothesis_1_cons$Eutrema_salsugineum_sigDE) 
                                                                 |
                                                                   HOG_level_list$hypothesis_1_cons$expansion == "yes" & HOG_level_list$hypothesis_1_cons$Arabidopsis_lyrata_sigDE>=1 & !is.na(HOG_level_list$hypothesis_1_cons$Arabidopsis_lyrata_sigDE)
                                                                 ,]$HOG

### change the dataset to plot here ###
dataset=HOGs_all_tol_exp_DEG_any_tol
title="HOGs_all_tol_exp_DEG_any_tol"
e <- ggplot(Not_min_2_expr_Conserved_with_expression_at_least_2_expressed_and_at_least_2_DEG_and_category_unique_HOGs[Not_min_2_expr_Conserved_with_expression_at_least_2_expressed_and_at_least_2_DEG_and_category_unique_HOGs$HOG %in% dataset,], aes(x="", fill=as.factor(category_of_regulation)))+
  geom_bar(width=1)+
  theme(legend.position="none")+
  ggtitle(title)+
  coord_polar("y")+
  scale_fill_brewer(palette = "Set2")
e

# 2e) What is the frequency of OGs with at least 1 DEG from both of the expanded species in "Hypothesis 1 expanded==yes" OGs?
HOGs_all_tol_exp_DEG_both_tol <- HOG_level_list$hypothesis_1_cons[HOG_level_list$hypothesis_1_cons$expansion == "yes" 
                                                                  & HOG_level_list$hypothesis_1_cons$Eutrema_salsugineum_sigDE>=1 & !is.na(HOG_level_list$hypothesis_1_cons$Eutrema_salsugineum_sigDE) 
                                                                  & HOG_level_list$hypothesis_1_cons$Arabidopsis_lyrata_sigDE>=1 & !is.na(HOG_level_list$hypothesis_1_cons$Arabidopsis_lyrata_sigDE)
                                                                  ,]$HOG
### change the dataset and title here ###
dataset=HOGs_all_tol_exp_DEG_both_tol
title="HOGs_all_tol_exp_DEG_both_tol"
pdf(paste0(OUTDIR, "/Regulation_category_ONLY_at_least_2_DEG_exp_and_DE_both_exp_Hypothesis_1_set_only_padj_less_0.1.pdf"), 12, 8)
f <- ggplot(Not_min_2_expr_Conserved_with_expression_at_least_2_expressed_and_at_least_2_DEG_and_category_unique_HOGs[Not_min_2_expr_Conserved_with_expression_at_least_2_expressed_and_at_least_2_DEG_and_category_unique_HOGs$HOG %in% dataset,], aes(x="", fill=as.factor(category_of_regulation)))+
  geom_bar(width=1)+
  ggtitle(title)+
  coord_polar("y")+
  scale_fill_manual(values = palette_OkabeIto_black[1:3])
f
dev.off()

pdf(paste0(OUTDIR, "/Regulation_category_ONLY_at_least_2_DEG_with_and_without_DEGs_in_Hypothesis_1_set_only_padj_less_0.1.pdf"), 12, 8)
ggarrange(a, b, c, d, e, f, ncol=3, nrow=2, common.legend = TRUE, legend="bottom")
dev.off()

###############################
####### Publication Plot: the same subset but as a stacked pie with the regulation categories in the Conserved Set as background ######
################################
dataset = HOGs_all_tol_exp_DEG_both_tol
ex=length(Not_min_2_expr_Conserved_with_expression_at_least_2_expressed_and_at_least_2_DEG_and_category_unique_HOGs[Not_min_2_expr_Conserved_with_expression_at_least_2_expressed_and_at_least_2_DEG_and_category_unique_HOGs$HOG %in% dataset,]$HOG)
total=length(Not_min_2_expr_Conserved_with_expression_at_least_2_expressed_and_at_least_2_DEG_and_category_unique_HOGs$HOG)
geneset1 <- names(summary(as.factor(Not_min_2_expr_Conserved_with_expression_at_least_2_expressed_and_at_least_2_DEG_and_category_unique_HOGs$category_of_regulation)))
geneset2 <- names(summary(as.factor(Not_min_2_expr_Conserved_with_expression_at_least_2_expressed_and_at_least_2_DEG_and_category_unique_HOGs[Not_min_2_expr_Conserved_with_expression_at_least_2_expressed_and_at_least_2_DEG_and_category_unique_HOGs$HOG %in% dataset,]$category_of_regulation)))
group1 <- c(total, total, total)
group2 <- c(ex, ex, ex)
At1 <- as.integer(c(summary(as.factor(Not_min_2_expr_Conserved_with_expression_at_least_2_expressed_and_at_least_2_DEG_and_category_unique_HOGs$category_of_regulation))))
At2 <- as.integer(c(summary(as.factor(Not_min_2_expr_Conserved_with_expression_at_least_2_expressed_and_at_least_2_DEG_and_category_unique_HOGs[Not_min_2_expr_Conserved_with_expression_at_least_2_expressed_and_at_least_2_DEG_and_category_unique_HOGs$HOG %in% dataset,]$category_of_regulation))))
at <- data.frame(geneset1, geneset2, group1, group2, At1, At2)
#make 2 stacked bar charts and then make the y-axis with polar coordinates with log10 x-axis
pdf(paste0(OUTDIR, "/Regulation_category_ONLY_at_least_2_DEG_exp_and_DE_both_exp_Hypothesis_1_in_Conserved_set_only_padj_less_0.1.pdf"), 12, 8)
a <- ggplot(at)+
  geom_bar(aes(x=log10(total)/2, y=At1, fill=geneset1), width = log10(total), position="fill", stat="identity", color="white", alpha=0.4)+
  geom_bar(aes(x=log10(ex)/2, y=At2, fill=geneset2), width = log10(ex), position="fill", stat="identity", color="white", alpha=1)+
  coord_polar(theta = "y", start=0)+
  scale_y_continuous(name ="Subset = Both tolerant exp. and DE, Background = Conserved and DE", limits= NULL, breaks=NULL, expand = expansion(0))+
  scale_x_continuous(name ="Proportion of HOGs per regulation category", limits= c(0,log10(total)), breaks = NULL, expand = expansion(0))+
  labs(fill="Regulation category")+
  theme(
    legend.position = "right",
    panel.grid.major = element_line(linetype = "dotted", colour="white"),
    panel.background = element_rect(fill="transparent", colour="black"))+
  geom_label(aes(x=log10(ex)-log10(ex)/2, y=0,label=paste(ex)), color="black", size=7)+
  geom_label(aes(x=log10(total)-log10(total)/6, y=0,label=paste(total)), color="black", size=7)+
  scale_fill_manual(values = palette_OkabeIto_black[1:3])
a
dev.off()
###################


# how many are only up?
length(Not_min_2_expr_Conserved_with_expression_at_least_2_expressed_and_at_least_2_DEG_and_category_unique_HOGs[Not_min_2_expr_Conserved_with_expression_at_least_2_expressed_and_at_least_2_DEG_and_category_unique_HOGs$HOG %in% dataset & Not_min_2_expr_Conserved_with_expression_at_least_2_expressed_and_at_least_2_DEG_and_category_unique_HOGs$category_of_regulation=="up",]$HOG)
# 24

# from how many categorized HOGs in total?
length(unique(Not_min_2_expr_Conserved_with_expression_at_least_2_expressed_and_at_least_2_DEG_and_category_unique_HOGs[Not_min_2_expr_Conserved_with_expression_at_least_2_expressed_and_at_least_2_DEG_and_category_unique_HOGs$HOG %in% dataset,]$HOG))
# 47

## --> No real difference for Hypothesis 1

### test if the exp and DE in both tolerant are significantly enriched for "only up"
# 2768 / 8800 = 0.315

q <- 24
m <- 2768
n <- (8800 - m)
k <- 47

p <- phyper((q-1), m, n, k, lower.tail=F)
p # 0.0039

### now do the same but the BG is the Set "Conserved with DE at least both tolerant"

dataset = HOGs_all_tol_exp_DEG_both_tol
ex=length(Not_min_2_expr_Conserved_with_expression_at_least_2_expressed_and_at_least_2_DEG_and_category_unique_HOGs[Not_min_2_expr_Conserved_with_expression_at_least_2_expressed_and_at_least_2_DEG_and_category_unique_HOGs$HOG %in% dataset,]$HOG)
total=length(Not_min_2_expr_Conserved_with_expression_at_least_2_expressed_and_at_least_2_DEG_and_category_unique_HOGs[Not_min_2_expr_Conserved_with_expression_at_least_2_expressed_and_at_least_2_DEG_and_category_unique_HOGs$HOG %in% HOGs_DEG_both_tol,]$HOG)
geneset1 <- names(summary(as.factor(Not_min_2_expr_Conserved_with_expression_at_least_2_expressed_and_at_least_2_DEG_and_category_unique_HOGs$category_of_regulation)))
geneset2 <- names(summary(as.factor(Not_min_2_expr_Conserved_with_expression_at_least_2_expressed_and_at_least_2_DEG_and_category_unique_HOGs[Not_min_2_expr_Conserved_with_expression_at_least_2_expressed_and_at_least_2_DEG_and_category_unique_HOGs$HOG %in% dataset,]$category_of_regulation)))
group1 <- c(total, total, total)
group2 <- c(ex, ex, ex)
At1 <- as.integer(c(summary(as.factor(Not_min_2_expr_Conserved_with_expression_at_least_2_expressed_and_at_least_2_DEG_and_category_unique_HOGs[
  Not_min_2_expr_Conserved_with_expression_at_least_2_expressed_and_at_least_2_DEG_and_category_unique_HOGs$HOG %in% HOGs_DEG_both_tol,]$category_of_regulation))))
At2 <- as.integer(c(summary(as.factor(Not_min_2_expr_Conserved_with_expression_at_least_2_expressed_and_at_least_2_DEG_and_category_unique_HOGs[
  Not_min_2_expr_Conserved_with_expression_at_least_2_expressed_and_at_least_2_DEG_and_category_unique_HOGs$HOG %in% dataset,]$category_of_regulation))))
at <- data.frame(geneset1, geneset2, group1, group2, At1, At2)
#make 2 stacked bar charts and then make the y-axis with polar coordinates with log10 x-axis
pdf(paste0(OUTDIR, "/Regulation_category_ONLY_at_least_2_DEG_exp_and_DE_both_exp_Hypothesis_1_in_HOGs_DEG_both_tol_only_padj_less_0.1.pdf"), 12, 8)
a <- ggplot(at)+
  geom_bar(aes(x=log10(total)/2, y=At1, fill=geneset1), width = log10(total), position="fill", stat="identity", color="white", alpha=0.4)+
  geom_bar(aes(x=log10(ex)/2, y=At2, fill=geneset2), width = log10(ex), position="fill", stat="identity", color="white", alpha=1)+
  coord_polar(theta = "y", start=0)+
  scale_y_continuous(name ="Subset = Both tolerant exp. and DE, Background = HOGs_DEG_both_tol", limits= NULL, breaks=NULL, expand = expansion(0))+
  scale_x_continuous(name ="Proportion of HOGs per regulation category", limits= c(0,log10(total)), breaks = NULL, expand = expansion(0))+
  labs(fill="Regulation category")+
  theme(
    legend.position = "right",
    panel.grid.major = element_line(linetype = "dotted", colour="white"),
    panel.background = element_rect(fill="transparent", colour="black"))+
  geom_label(aes(x=log10(ex)-log10(ex)/2, y=0,label=paste(ex)), color="black", size=7)+
  geom_label(aes(x=log10(total)-log10(total)/6, y=0,label=paste(total)), color="black", size=7)+
  scale_fill_manual(values = palette_OkabeIto_black[1:3])
a
dev.off()

## test for enrichment with only UP

# how many are only up?
length(Not_min_2_expr_Conserved_with_expression_at_least_2_expressed_and_at_least_2_DEG_and_category_unique_HOGs[Not_min_2_expr_Conserved_with_expression_at_least_2_expressed_and_at_least_2_DEG_and_category_unique_HOGs$HOG %in% dataset & Not_min_2_expr_Conserved_with_expression_at_least_2_expressed_and_at_least_2_DEG_and_category_unique_HOGs$category_of_regulation=="up",]$HOG)
# 24

# from how many categorized HOGs in total?
length(unique(Not_min_2_expr_Conserved_with_expression_at_least_2_expressed_and_at_least_2_DEG_and_category_unique_HOGs[Not_min_2_expr_Conserved_with_expression_at_least_2_expressed_and_at_least_2_DEG_and_category_unique_HOGs$HOG %in% dataset,]$HOG))
# 47

# In BG: how many are only up?
length(Not_min_2_expr_Conserved_with_expression_at_least_2_expressed_and_at_least_2_DEG_and_category_unique_HOGs[Not_min_2_expr_Conserved_with_expression_at_least_2_expressed_and_at_least_2_DEG_and_category_unique_HOGs$HOG %in% HOGs_DEG_both_tol & Not_min_2_expr_Conserved_with_expression_at_least_2_expressed_and_at_least_2_DEG_and_category_unique_HOGs$category_of_regulation=="up",]$HOG)
# 1462

# In BG: from how many HOGs in total?
length(unique(Not_min_2_expr_Conserved_with_expression_at_least_2_expressed_and_at_least_2_DEG_and_category_unique_HOGs[Not_min_2_expr_Conserved_with_expression_at_least_2_expressed_and_at_least_2_DEG_and_category_unique_HOGs$HOG %in% HOGs_DEG_both_tol,]$HOG))
# 4638

### test if the exp and DE in both tolerant are significantly enriched for "only up"
# 24 / 47 = 0.5106383
# bg 1462 / 4638 = 0.315

q <- 24
m <- 1462
n <- (4638 - m)
k <- 47

p <- phyper((q-1), m, n, k, lower.tail=F)
p # 0.00396

## --> makes no difference to using the whole Conserved Set with at least 2 DEGs = which means that probably all HOGs with min. 2 DEGs
# have min. 1 DEG from each of the tolerant, which is the largest group of DEGs


### Hypothesis 2
# 2.1)j)1) What is the frequency of OGs with at least 1 DEG from the Hypothesis 2 expanded species in "Conserved" OGs?
HOGs_DEG_Eutrema <- HOG_level_list$hypothesis_2_cons[HOG_level_list$hypothesis_2_cons$Eutrema_salsugineum_sigDE>=1 & !is.na(HOG_level_list$hypothesis_2_cons$Eutrema_salsugineum_sigDE)
                                                     ,]$HOG
### change the dataset and title here ###
dataset=HOGs_DEG_Eutrema
title="HOGs_DEG_Eutrema"
a <- ggplot(Not_min_2_expr_Conserved_with_expression_at_least_2_expressed_and_at_least_2_DEG_and_category_unique_HOGs[Not_min_2_expr_Conserved_with_expression_at_least_2_expressed_and_at_least_2_DEG_and_category_unique_HOGs$HOG %in% dataset,], aes(x="", fill=as.factor(category_of_regulation)))+
  geom_bar(width=1)+
  ggtitle(title)+
  coord_polar("y")

# 2.1)i)2) Within "Conserved": Number of OGs with "Hypothesis 2 expanded==yes"
HOGs_Eutrema_exp <- HOG_level_list$hypothesis_2_cons[HOG_level_list$hypothesis_2_cons$expansion == "yes",]$HOG

### change the dataset and title here ###
dataset=HOGs_Eutrema_exp
title="HOGs_Eutrema_exp"
b <- ggplot(Not_min_2_expr_Conserved_with_expression_at_least_2_expressed_and_at_least_2_DEG_and_category_unique_HOGs[Not_min_2_expr_Conserved_with_expression_at_least_2_expressed_and_at_least_2_DEG_and_category_unique_HOGs$HOG %in% dataset,], aes(x="", fill=as.factor(category_of_regulation)))+
  geom_bar(width=1)+
  ggtitle(title)+
  coord_polar("y")

# 2i) Within "Conserved": What is the frequency of OGs with at least 1 DEG in "Hypothesis 2 expanded==yes" OGs?
cons_1_and_2_merged_2 <-  merge(HOG_level_list$hypothesis_1_cons[,c("HOG", "total_sigDE")], HOG_level_list$hypothesis_2_cons, by="HOG")
# select for total_sigDE.x because we are interested in total_sigDE from all 4 species (inferred from data frame HOG_level_list$hypothesis_1_cons, which contains all 4 species)
HOGs_Eutrema_exp_DEG_any <- cons_1_and_2_merged_2[cons_1_and_2_merged_2$expansion == "yes" & !is.na(cons_1_and_2_merged_2$total_sigDE.x) & cons_1_and_2_merged_2$total_sigDE.x>=1,]$HOG

### change the dataset and title here ###
dataset=HOGs_Eutrema_exp_DEG_any
title="HOGs_Eutrema_exp_DEG_any"
c <- ggplot(Not_min_2_expr_Conserved_with_expression_at_least_2_expressed_and_at_least_2_DEG_and_category_unique_HOGs[Not_min_2_expr_Conserved_with_expression_at_least_2_expressed_and_at_least_2_DEG_and_category_unique_HOGs$HOG %in% dataset,], aes(x="", fill=as.factor(category_of_regulation)))+
  geom_bar(width=1)+
  ggtitle(title)+
  coord_polar("y")

# 2j) What is the frequency of OGs with at least 1 DEG from any of the expanded species in "Hypothesis 2 expanded==yes" OGs?
HOGs_Eutrema_exp_DEG_Eutrema <- HOG_level_list$hypothesis_2_cons[HOG_level_list$hypothesis_2_cons$expansion == "yes" & HOG_level_list$hypothesis_2_cons$Eutrema_salsugineum_sigDE>=1 & !is.na(HOG_level_list$hypothesis_2_cons$Eutrema_salsugineum_sigDE) 
                                                                 ,]$HOG

### change the dataset and title here ###
dataset=HOGs_Eutrema_exp_DEG_Eutrema
title="HOGs_Eutrema_exp_DEG_Eutrema"
d <- ggplot(Not_min_2_expr_Conserved_with_expression_at_least_2_expressed_and_at_least_2_DEG_and_category_unique_HOGs[Not_min_2_expr_Conserved_with_expression_at_least_2_expressed_and_at_least_2_DEG_and_category_unique_HOGs$HOG %in% dataset,], aes(x="", fill=as.factor(category_of_regulation)))+
  geom_bar(width=1)+
  ggtitle(title)+
  coord_polar("y")

pdf(paste0(OUTDIR, "/Regulation_category_ONLY_at_least_2_DEG_with_and_without_DEGs_in_Hypothesis_2_set_only_padj_less_0.1.pdf"), 12, 8)
ggarrange(a, b, c, d, ncol=2, nrow=2, common.legend = TRUE, legend="bottom")
dev.off()

## --> No real difference for Hypothesis 2

## Plot the same subset but as a stacked pie with the regulation categories in the Conserved Set as background

dataset = HOGs_Eutrema_exp_DEG_Eutrema
ex=length(Not_min_2_expr_Conserved_with_expression_at_least_2_expressed_and_at_least_2_DEG_and_category_unique_HOGs[Not_min_2_expr_Conserved_with_expression_at_least_2_expressed_and_at_least_2_DEG_and_category_unique_HOGs$HOG %in% dataset,]$HOG)
total=length(Not_min_2_expr_Conserved_with_expression_at_least_2_expressed_and_at_least_2_DEG_and_category_unique_HOGs$HOG)
geneset1 <-names(summary(as.factor(Not_min_2_expr_Conserved_with_expression_at_least_2_expressed_and_at_least_2_DEG_and_category_unique_HOGs$category_of_regulation)))
geneset2 <- names(summary(as.factor(Not_min_2_expr_Conserved_with_expression_at_least_2_expressed_and_at_least_2_DEG_and_category_unique_HOGs[Not_min_2_expr_Conserved_with_expression_at_least_2_expressed_and_at_least_2_DEG_and_category_unique_HOGs$HOG %in% dataset,]$category_of_regulation)))
group1 <- c(total, total, total)
group2 <- c(ex, ex, ex)
At1 <- as.integer(c(summary(as.factor(Not_min_2_expr_Conserved_with_expression_at_least_2_expressed_and_at_least_2_DEG_and_category_unique_HOGs$category_of_regulation))))
At2 <- as.integer(c(summary(as.factor(Not_min_2_expr_Conserved_with_expression_at_least_2_expressed_and_at_least_2_DEG_and_category_unique_HOGs[Not_min_2_expr_Conserved_with_expression_at_least_2_expressed_and_at_least_2_DEG_and_category_unique_HOGs$HOG %in% dataset,]$category_of_regulation))))
at <- data.frame(geneset1, geneset2, group1, group2, At1, At2)
#make 2 stacked bar charts and then make the y-axis with polar coordinates with log10 x-axis
pdf(paste0(OUTDIR, "/Regulation_category_ONLY_at_least_2_DEG_exp_and_DE_exp_Hypothesis_2_in_Conserved_set_only_padj_less_0.1.pdf"), 12, 8)
b <- ggplot(at)+
  geom_bar(aes(x=log10(total)/2, y=At1, fill=geneset1), width = log10(total), position="fill", stat="identity", color="white", alpha=0.4)+
  geom_bar(aes(x=log10(ex)/2, y=At2, fill=geneset2), width = log10(ex), position="fill", stat="identity", color="white", alpha=1)+
  coord_polar(theta = "y", start=0)+
  scale_y_continuous(name ="Subset = Eutrema exp. and DE, Background = Conserved and DE", limits= NULL, breaks=NULL, expand = expansion(0))+
  scale_x_continuous(name ="Proportion of HOGs per regulation category", limits= c(0,log10(total)), breaks = NULL, expand = expansion(0))+
  labs(fill="Regulation category")+
  theme(
    legend.position = "right",
    panel.grid.major = element_line(linetype = "dotted", colour="white"),
    panel.background = element_rect(fill="transparent", colour="black"))+
  geom_label(aes(x=log10(ex)-log10(ex)/2, y=0,label=paste(ex)), color="black", size=7)+
  geom_label(aes(x=log10(total)-log10(total)/6, y=0,label=paste(total)), color="black", size=7)+
  scale_fill_manual(values = palette_OkabeIto_black[1:3])
b
dev.off()

### now do the same but the BG is the Set "Conserved with DE at least Eutrema"

dataset = HOGs_Eutrema_exp_DEG_Eutrema
ex=length(Not_min_2_expr_Conserved_with_expression_at_least_2_expressed_and_at_least_2_DEG_and_category_unique_HOGs[Not_min_2_expr_Conserved_with_expression_at_least_2_expressed_and_at_least_2_DEG_and_category_unique_HOGs$HOG %in% dataset,]$HOG)
total=length(Not_min_2_expr_Conserved_with_expression_at_least_2_expressed_and_at_least_2_DEG_and_category_unique_HOGs[Not_min_2_expr_Conserved_with_expression_at_least_2_expressed_and_at_least_2_DEG_and_category_unique_HOGs$HOG %in% HOGs_DEG_Eutrema,]$HOG)
geneset1 <-names(summary(as.factor(Not_min_2_expr_Conserved_with_expression_at_least_2_expressed_and_at_least_2_DEG_and_category_unique_HOGs$category_of_regulation)))
geneset2 <- names(summary(as.factor(Not_min_2_expr_Conserved_with_expression_at_least_2_expressed_and_at_least_2_DEG_and_category_unique_HOGs[Not_min_2_expr_Conserved_with_expression_at_least_2_expressed_and_at_least_2_DEG_and_category_unique_HOGs$HOG %in% dataset,]$category_of_regulation)))
group1 <- c(total, total, total)
group2 <- c(ex, ex, ex)
At1 <- as.integer(c(summary(as.factor(Not_min_2_expr_Conserved_with_expression_at_least_2_expressed_and_at_least_2_DEG_and_category_unique_HOGs[
  Not_min_2_expr_Conserved_with_expression_at_least_2_expressed_and_at_least_2_DEG_and_category_unique_HOGs$HOG %in% HOGs_DEG_Eutrema,]$category_of_regulation))))
At2 <- as.integer(c(summary(as.factor(Not_min_2_expr_Conserved_with_expression_at_least_2_expressed_and_at_least_2_DEG_and_category_unique_HOGs[
  Not_min_2_expr_Conserved_with_expression_at_least_2_expressed_and_at_least_2_DEG_and_category_unique_HOGs$HOG %in% dataset,]$category_of_regulation))))
at <- data.frame(geneset1, geneset2, group1, group2, At1, At2)
#make 2 stacked bar charts and then make the y-axis with polar coordinates with log10 x-axis
pdf(paste0(OUTDIR, "/Regulation_category_ONLY_at_least_2_DEG_exp_and_DE_exp_Hypothesis_2_in_Eutrema_DE_set_only_padj_less_0.1.pdf"), 12, 8)
b <- ggplot(at)+
  geom_bar(aes(x=log10(total)/2, y=At1, fill=geneset1), width = log10(total), position="fill", stat="identity", color="white", alpha=0.4)+
  geom_bar(aes(x=log10(ex)/2, y=At2, fill=geneset2), width = log10(ex), position="fill", stat="identity", color="white", alpha=1)+
  coord_polar(theta = "y", start=0)+
  scale_y_continuous(name ="Subset = Eutrema exp. and DE, Background = Conserved and Eutrema DE", limits= NULL, breaks=NULL, expand = expansion(0))+
  scale_x_continuous(name ="Proportion of HOGs per regulation category", limits= c(0,log10(total)), breaks = NULL, expand = expansion(0))+
  labs(fill="Regulation category")+
  theme(
    legend.position = "right",
    panel.grid.major = element_line(linetype = "dotted", colour="white"),
    panel.background = element_rect(fill="transparent", colour="black"))+
  geom_label(aes(x=log10(ex)-log10(ex)/2, y=0,label=paste(ex)), color="black", size=7)+
  geom_label(aes(x=log10(total)-log10(total)/6, y=0,label=paste(total)), color="black", size=7)+
  scale_fill_manual(values = palette_OkabeIto_black[1:3])
b
dev.off()

## test for enrichment with only UP
# how many are only up?
length(Not_min_2_expr_Conserved_with_expression_at_least_2_expressed_and_at_least_2_DEG_and_category_unique_HOGs[Not_min_2_expr_Conserved_with_expression_at_least_2_expressed_and_at_least_2_DEG_and_category_unique_HOGs$HOG %in% dataset & Not_min_2_expr_Conserved_with_expression_at_least_2_expressed_and_at_least_2_DEG_and_category_unique_HOGs$category_of_regulation=="up",]$HOG)
# 112

# from how many categorized HOGs in total?
length(unique(Not_min_2_expr_Conserved_with_expression_at_least_2_expressed_and_at_least_2_DEG_and_category_unique_HOGs[Not_min_2_expr_Conserved_with_expression_at_least_2_expressed_and_at_least_2_DEG_and_category_unique_HOGs$HOG %in% dataset,]$HOG))
# 294

# In BG: how many are only up?
length(Not_min_2_expr_Conserved_with_expression_at_least_2_expressed_and_at_least_2_DEG_and_category_unique_HOGs[Not_min_2_expr_Conserved_with_expression_at_least_2_expressed_and_at_least_2_DEG_and_category_unique_HOGs$HOG %in% HOGs_DEG_Eutrema & Not_min_2_expr_Conserved_with_expression_at_least_2_expressed_and_at_least_2_DEG_and_category_unique_HOGs$category_of_regulation=="up",]$HOG)
# 1769

# In BG: from how many HOGs in total?
length(unique(Not_min_2_expr_Conserved_with_expression_at_least_2_expressed_and_at_least_2_DEG_and_category_unique_HOGs[Not_min_2_expr_Conserved_with_expression_at_least_2_expressed_and_at_least_2_DEG_and_category_unique_HOGs$HOG %in% HOGs_DEG_Eutrema,]$HOG))
# 6050

### test if the exp and DE in Eutrema are significantly enriched for "only up"
# 112 / 294 = 0.38
# bg 1769/6050 = 0.2923967
q <- 112
m <- 1769
n <- (6050 - m)
k <- 294

p <- phyper((q-1), m, n, k, lower.tail=F)
p # 0.0005


### Hypothesis 3
# 2.1)l)1) What is the frequency of OGs with at least 1 DEG from the Hypothesis 3 expanded species in "Conserved" OGs?
HOGs_DEG_Alyrata <- HOG_level_list$hypothesis_3_cons[HOG_level_list$hypothesis_3_cons$Arabidopsis_lyrata_sigDE>=1 & !is.na(HOG_level_list$hypothesis_3_cons$Arabidopsis_lyrata_sigDE)
                                                     ,]$HOG

### change the dataset and title here ###
dataset=HOGs_DEG_Alyrata
title="HOGs_DEG_Alyrata"
a <- ggplot(Not_min_2_expr_Conserved_with_expression_at_least_2_expressed_and_at_least_2_DEG_and_category_unique_HOGs[Not_min_2_expr_Conserved_with_expression_at_least_2_expressed_and_at_least_2_DEG_and_category_unique_HOGs$HOG %in% dataset,], aes(x="", fill=as.factor(category_of_regulation)))+
  geom_bar(width=1)+
  ggtitle(title)+
  coord_polar("y")

# 2.1)k)2) Within "Conserved": Number of OGs with "Hypothesis 3 expanded==yes"
HOGs_Alyrata_exp <- HOG_level_list$hypothesis_3_cons[HOG_level_list$hypothesis_3_cons$expansion == "yes",]$HOG

### change the dataset and title here ###
dataset=HOGs_Alyrata_exp
title="HOGs_Alyrata_exp"
b <- ggplot(Not_min_2_expr_Conserved_with_expression_at_least_2_expressed_and_at_least_2_DEG_and_category_unique_HOGs[Not_min_2_expr_Conserved_with_expression_at_least_2_expressed_and_at_least_2_DEG_and_category_unique_HOGs$HOG %in% dataset,], aes(x="", fill=as.factor(category_of_regulation)))+
  geom_bar(width=1)+
  ggtitle(title)+
  coord_polar("y")

# 2k) Within "Conserved": What is the frequency of OGs with at least 1 DEG in "Hypothesis 3 expanded==yes" OGs?
cons_1_and_3_merged_2 <-  merge(HOG_level_list$hypothesis_1_cons[,c("HOG", "total_sigDE")], HOG_level_list$hypothesis_3_cons, by="HOG")
# select for total_sigDE.x because we are interested in total_sigDE from all 4 species (inferred from data frame HOG_level_list$hypothesis_1_cons, which contains all 4 species)
HOGs_Alyrata_exp_DEG_any <- cons_1_and_3_merged_2[cons_1_and_3_merged_2$expansion == "yes" & !is.na(cons_1_and_3_merged_2$total_sigDE.x) & cons_1_and_3_merged_2$total_sigDE.x>=1,]$HOG

### change the dataset and title here ###
dataset=HOGs_Alyrata_exp_DEG_any
title="HOGs_Alyrata_exp_DEG_any"
c <- ggplot(Not_min_2_expr_Conserved_with_expression_at_least_2_expressed_and_at_least_2_DEG_and_category_unique_HOGs[Not_min_2_expr_Conserved_with_expression_at_least_2_expressed_and_at_least_2_DEG_and_category_unique_HOGs$HOG %in% dataset,], aes(x="", fill=as.factor(category_of_regulation)))+
  geom_bar(width=1)+
  ggtitle(title)+
  coord_polar("y")

# 2l) What is the frequency of OGs with at least 1 DEG from any of the expanded species in "Hypothesis 3 expanded==yes" OGs?
HOGs_Alyrata_exp_DEG_Alyrata <- HOG_level_list$hypothesis_3_cons[HOG_level_list$hypothesis_3_cons$expansion == "yes" & HOG_level_list$hypothesis_3_cons$Arabidopsis_lyrata_sigDE>=1 & !is.na(HOG_level_list$hypothesis_3_cons$Arabidopsis_lyrata_sigDE) 
                                                                 ,]$HOG

### change the dataset and title here ###
dataset=HOGs_Alyrata_exp_DEG_Alyrata
title="HOGs_Alyrata_exp_DEG_Alyrata"
d <- ggplot(Not_min_2_expr_Conserved_with_expression_at_least_2_expressed_and_at_least_2_DEG_and_category_unique_HOGs[Not_min_2_expr_Conserved_with_expression_at_least_2_expressed_and_at_least_2_DEG_and_category_unique_HOGs$HOG %in% dataset,], aes(x="", fill=as.factor(category_of_regulation)))+
  geom_bar(width=1)+
  ggtitle(title)+
  coord_polar("y")

pdf(paste0(OUTDIR, "/Regulation_category_ONLY_at_least_2_DEG_with_and_without_DEGs_in_Hypothesis_3_set_only_padj_less_0.1.pdf"), 12, 8)
ggarrange(a, b, c, d, ncol=2, nrow=2, common.legend = TRUE, legend="bottom")
dev.off()

## --> No real difference for Hypothesis 3

## Plot the same subset but as a stacked pie with the regulation categories in the Conserved Set as background

dataset=HOGs_Alyrata_exp_DEG_Alyrata
ex=length(Not_min_2_expr_Conserved_with_expression_at_least_2_expressed_and_at_least_2_DEG_and_category_unique_HOGs[Not_min_2_expr_Conserved_with_expression_at_least_2_expressed_and_at_least_2_DEG_and_category_unique_HOGs$HOG %in% dataset,]$HOG)
total=length(Not_min_2_expr_Conserved_with_expression_at_least_2_expressed_and_at_least_2_DEG_and_category_unique_HOGs$HOG)
geneset1 <-names(summary(as.factor(Not_min_2_expr_Conserved_with_expression_at_least_2_expressed_and_at_least_2_DEG_and_category_unique_HOGs$category_of_regulation)))
geneset2 <- names(summary(as.factor(Not_min_2_expr_Conserved_with_expression_at_least_2_expressed_and_at_least_2_DEG_and_category_unique_HOGs[Not_min_2_expr_Conserved_with_expression_at_least_2_expressed_and_at_least_2_DEG_and_category_unique_HOGs$HOG %in% dataset,]$category_of_regulation)))
group1 <- c(total, total, total)
group2 <- c(ex, ex, ex)
At1 <- as.integer(c(summary(as.factor(Not_min_2_expr_Conserved_with_expression_at_least_2_expressed_and_at_least_2_DEG_and_category_unique_HOGs$category_of_regulation))))
At2 <- as.integer(c(summary(as.factor(Not_min_2_expr_Conserved_with_expression_at_least_2_expressed_and_at_least_2_DEG_and_category_unique_HOGs[Not_min_2_expr_Conserved_with_expression_at_least_2_expressed_and_at_least_2_DEG_and_category_unique_HOGs$HOG %in% dataset,]$category_of_regulation))))
at <- data.frame(geneset1, geneset2, group1, group2, At1, At2)
#make 2 stacked bar charts and then make the y-axis with polar coordinates with log10 x-axis
pdf(paste0(OUTDIR, "/Regulation_category_ONLY_at_least_2_DEG_exp_and_DE_exp_Hypothesis_3_in_Conserved_set_only_padj_less_0.1.pdf"), 12, 8)
c <- ggplot(at)+
  geom_bar(aes(x=log10(total)/2, y=At1, fill=geneset1), width = log10(total), position="fill", stat="identity", color="white", alpha=0.4)+
  geom_bar(aes(x=log10(ex)/2, y=At2, fill=geneset2), width = log10(ex), position="fill", stat="identity", color="white", alpha=1)+
  coord_polar(theta = "y", start=0)+
  scale_y_continuous(name ="Subset = Alyrata exp. and DE, Background = Conserved with DE", limits= NULL, breaks=NULL, expand = expansion(0))+
  scale_x_continuous(name ="Proportion of HOGs per regulation category", limits= c(0,log10(total)), breaks = NULL, expand = expansion(0))+
  labs(fill="Regulation category")+
  theme(
    legend.position = "right",
    panel.grid.major = element_line(linetype = "dotted", colour="white"),
    panel.background = element_rect(fill="transparent", colour="black"))+
  geom_label(aes(x=log10(ex)-log10(ex)/2, y=0,label=paste(ex)), color="black", size=7)+
  geom_label(aes(x=log10(total)-log10(total)/6, y=0,label=paste(total)), color="black", size=7)+
  scale_fill_manual(values = palette_OkabeIto_black[1:3])
c
dev.off()


## Plot the same subset but as a stacked pie with the regulation categories in the HOGs with DEG from Alyrata within Conserved Set as background
dataset=HOGs_Alyrata_exp_DEG_Alyrata
ex=length(Not_min_2_expr_Conserved_with_expression_at_least_2_expressed_and_at_least_2_DEG_and_category_unique_HOGs[Not_min_2_expr_Conserved_with_expression_at_least_2_expressed_and_at_least_2_DEG_and_category_unique_HOGs$HOG %in% dataset,]$HOG)
total=length(Not_min_2_expr_Conserved_with_expression_at_least_2_expressed_and_at_least_2_DEG_and_category_unique_HOGs[Not_min_2_expr_Conserved_with_expression_at_least_2_expressed_and_at_least_2_DEG_and_category_unique_HOGs$HOG %in% HOGs_DEG_Alyrata,]$HOG)
geneset1 <-names(summary(as.factor(Not_min_2_expr_Conserved_with_expression_at_least_2_expressed_and_at_least_2_DEG_and_category_unique_HOGs$category_of_regulation)))
geneset2 <- names(summary(as.factor(Not_min_2_expr_Conserved_with_expression_at_least_2_expressed_and_at_least_2_DEG_and_category_unique_HOGs[Not_min_2_expr_Conserved_with_expression_at_least_2_expressed_and_at_least_2_DEG_and_category_unique_HOGs$HOG %in% dataset,]$category_of_regulation)))
group1 <- c(total, total, total)
group2 <- c(ex, ex, ex)
At1 <- as.integer(c(summary(as.factor(Not_min_2_expr_Conserved_with_expression_at_least_2_expressed_and_at_least_2_DEG_and_category_unique_HOGs[
  Not_min_2_expr_Conserved_with_expression_at_least_2_expressed_and_at_least_2_DEG_and_category_unique_HOGs$HOG %in% HOGs_DEG_Alyrata,]$category_of_regulation))))
At2 <- as.integer(c(summary(as.factor(Not_min_2_expr_Conserved_with_expression_at_least_2_expressed_and_at_least_2_DEG_and_category_unique_HOGs[
  Not_min_2_expr_Conserved_with_expression_at_least_2_expressed_and_at_least_2_DEG_and_category_unique_HOGs$HOG %in% dataset,]$category_of_regulation))))
at <- data.frame(geneset1, geneset2, group1, group2, At1, At2)
#make 2 stacked bar charts and then make the y-axis with polar coordinates with log10 x-axis
pdf(paste0(OUTDIR, "/Regulation_category_ONLY_at_least_2_DEG_exp_and_DE_exp_Hypothesis_3_in_HOGs_DE_Alyrata_only_padj_less_0.1.pdf"), 12, 8)
c <- ggplot(at)+
  geom_bar(aes(x=log10(total)/2, y=At1, fill=geneset1), width = log10(total), position="fill", stat="identity", color="white", alpha=0.4)+
  geom_bar(aes(x=log10(ex)/2, y=At2, fill=geneset2), width = log10(ex), position="fill", stat="identity", color="white", alpha=1)+
  coord_polar(theta = "y", start=0)+
  scale_y_continuous(name ="Subset = Alyrata exp. and DE, Background = Alyrata DE", limits= NULL, breaks=NULL, expand = expansion(0))+
  scale_x_continuous(name ="Proportion of HOGs per regulation category", limits= c(0,log10(total)), breaks = NULL, expand = expansion(0))+
  labs(fill="Regulation category")+
  theme(
    legend.position = "right",
    panel.grid.major = element_line(linetype = "dotted", colour="white"),
    panel.background = element_rect(fill="transparent", colour="black"))+
  geom_label(aes(x=log10(ex)-log10(ex)/2, y=0,label=paste(ex)), color="black", size=7)+
  geom_label(aes(x=log10(total)-log10(total)/6, y=0,label=paste(total)), color="black", size=7)+
  scale_fill_manual(values = palette_OkabeIto_black[1:3])
c
dev.off()


## test for enrichment with only UP in Alyrata

# how many are only up?
length(Not_min_2_expr_Conserved_with_expression_at_least_2_expressed_and_at_least_2_DEG_and_category_unique_HOGs[Not_min_2_expr_Conserved_with_expression_at_least_2_expressed_and_at_least_2_DEG_and_category_unique_HOGs$HOG %in% dataset & Not_min_2_expr_Conserved_with_expression_at_least_2_expressed_and_at_least_2_DEG_and_category_unique_HOGs$category_of_regulation=="up",]$HOG)
# 178

# from how many categorized HOGs in total?
length(unique(Not_min_2_expr_Conserved_with_expression_at_least_2_expressed_and_at_least_2_DEG_and_category_unique_HOGs[Not_min_2_expr_Conserved_with_expression_at_least_2_expressed_and_at_least_2_DEG_and_category_unique_HOGs$HOG %in% dataset,]$HOG))
# 452

# In BG: how many are only up?
length(Not_min_2_expr_Conserved_with_expression_at_least_2_expressed_and_at_least_2_DEG_and_category_unique_HOGs[Not_min_2_expr_Conserved_with_expression_at_least_2_expressed_and_at_least_2_DEG_and_category_unique_HOGs$HOG %in% HOGs_DEG_Alyrata & Not_min_2_expr_Conserved_with_expression_at_least_2_expressed_and_at_least_2_DEG_and_category_unique_HOGs$category_of_regulation=="up",]$HOG)
# 2193

# In BG: from how many HOGs in total?
length(unique(Not_min_2_expr_Conserved_with_expression_at_least_2_expressed_and_at_least_2_DEG_and_category_unique_HOGs[Not_min_2_expr_Conserved_with_expression_at_least_2_expressed_and_at_least_2_DEG_and_category_unique_HOGs$HOG %in% HOGs_DEG_Alyrata,]$HOG))
# 6840

### test if the exp and DE in Alyrata are significantly enriched for "only up"
# 2193 / 6840 = 0.32
# 178/452 = 0.3938
q <- 178
m <- 2193
n <- (6840 - m)
k <- 452

p <- phyper((q-1), m, n, k, lower.tail=F)
p # 0.0004057926


### Hypothesis 4
# 2.1)g)1) What is the frequency of OGs with at least 1 DEG from any of the Hypothesis 4 expanded species in "Conserved" OGs?
HOGs_DEG_any_sens <- HOG_level_list$hypothesis_4_cons[HOG_level_list$hypothesis_4_cons$Arabidopsis_thaliana_sigDE>=1 & !is.na(HOG_level_list$hypothesis_4_cons$Arabidopsis_thaliana_sigDE)
                                                      |
                                                        HOG_level_list$hypothesis_4_cons$Brassica_napus_sigDE>=1 & !is.na(HOG_level_list$hypothesis_4_cons$Brassica_napus_sigDE)
                                                      ,]$HOG

### change the dataset to plot here ###
dataset=HOGs_DEG_any_sens
title="HOGs_DEG_any_sens"
a <- ggplot(Not_min_2_expr_Conserved_with_expression_at_least_2_expressed_and_at_least_2_DEG_and_category_unique_HOGs[Not_min_2_expr_Conserved_with_expression_at_least_2_expressed_and_at_least_2_DEG_and_category_unique_HOGs$HOG %in% dataset,], aes(x="", fill=as.factor(category_of_regulation)))+
  geom_bar(width=1)+
  ggtitle(title)+
  coord_polar("y")

# 2.1)h)1) What is the frequency of OGs with at least 1 DEG from both of the Hypothesis 4 expanded species in "Conserved" OGs?
HOGs_DEG_both_sens <- HOG_level_list$hypothesis_4_cons[HOG_level_list$hypothesis_4_cons$Arabidopsis_thaliana_sigDE>=1 & !is.na(HOG_level_list$hypothesis_4_cons$Arabidopsis_thaliana_sigDE)
                                                       &
                                                         HOG_level_list$hypothesis_4_cons$Brassica_napus_sigDE>=1 & !is.na(HOG_level_list$hypothesis_4_cons$Brassica_napus_sigDE)
                                                       ,]$HOG
### change the dataset to plot here ###
dataset=HOGs_DEG_both_sens
title="HOGs_DEG_both_sens"
b <- ggplot(Not_min_2_expr_Conserved_with_expression_at_least_2_expressed_and_at_least_2_DEG_and_category_unique_HOGs[Not_min_2_expr_Conserved_with_expression_at_least_2_expressed_and_at_least_2_DEG_and_category_unique_HOGs$HOG %in% dataset,], aes(x="", fill=as.factor(category_of_regulation)))+
  geom_bar(width=1)+
  ggtitle(title)+
  coord_polar("y")

# 2.1)f)2) Within "Conserved": Number of OGs with "Hypothesis 4 expanded==yes"
HOGs_all_sens_exp <- HOG_level_list$hypothesis_4_cons[HOG_level_list$hypothesis_4_cons$expansion == "yes",]$HOG

### change the dataset to plot here ###
dataset=HOGs_all_sens_exp
title="HOGs_all_sens_exp"
c <- ggplot(Not_min_2_expr_Conserved_with_expression_at_least_2_expressed_and_at_least_2_DEG_and_category_unique_HOGs[Not_min_2_expr_Conserved_with_expression_at_least_2_expressed_and_at_least_2_DEG_and_category_unique_HOGs$HOG %in% dataset,], aes(x="", fill=as.factor(category_of_regulation)))+
  geom_bar(width=1)+
  ggtitle(title)+
  coord_polar("y")

# 2f) Within "Conserved": What is the frequency of OGs with at least 1 DEG in "Hypothesis 4 expanded==yes" OGs?
HOGs_all_sens_exp_DEG_any <- HOG_level_list$hypothesis_4_cons[HOG_level_list$hypothesis_4_cons$expansion == "yes" & !is.na(HOG_level_list$hypothesis_4_cons$total_sigDE) & HOG_level_list$hypothesis_4_cons$total_sigDE>=1,]$HOG

### change the dataset to plot here ###
dataset=HOGs_all_sens_exp_DEG_any
title="HOGs_all_sens_exp_DEG_any"
d <- ggplot(Not_min_2_expr_Conserved_with_expression_at_least_2_expressed_and_at_least_2_DEG_and_category_unique_HOGs[Not_min_2_expr_Conserved_with_expression_at_least_2_expressed_and_at_least_2_DEG_and_category_unique_HOGs$HOG %in% dataset,], aes(x="", fill=as.factor(category_of_regulation)))+
  geom_bar(width=1)+
  ggtitle(title)+
  coord_polar("y")

# 2g) What is the frequency of OGs with at least 1 DEG from any of the expanded species in "Hypothesis 4 expanded==yes" OGs?
HOGs_all_sens_exp_DEG_any_sens <- HOG_level_list$hypothesis_4_cons[HOG_level_list$hypothesis_4_cons$expansion == "yes" & HOG_level_list$hypothesis_4_cons$Arabidopsis_thaliana_sigDE>=1 & !is.na(HOG_level_list$hypothesis_4_cons$Arabidopsis_thaliana_sigDE) 
                                                                   |
                                                                     HOG_level_list$hypothesis_4_cons$expansion == "yes" & HOG_level_list$hypothesis_4_cons$Brassica_napus_sigDE>=1 & !is.na(HOG_level_list$hypothesis_4_cons$Brassica_napus_sigDE)
                                                                   ,]$HOG
### change the dataset to plot here ###
dataset=HOGs_all_sens_exp_DEG_any_sens
title="HOGs_all_sens_exp_DEG_any_sens"
e <- ggplot(Not_min_2_expr_Conserved_with_expression_at_least_2_expressed_and_at_least_2_DEG_and_category_unique_HOGs[Not_min_2_expr_Conserved_with_expression_at_least_2_expressed_and_at_least_2_DEG_and_category_unique_HOGs$HOG %in% dataset,], aes(x="", fill=as.factor(category_of_regulation)))+
  geom_bar(width=1)+
  ggtitle(title)+
  coord_polar("y")


# 2h) What is the frequency of OGs with at least 1 DEG from both of the expanded species in "Hypothesis 4 expanded==yes" OGs?
HOGs_all_sens_exp_DEG_both_sens <- HOG_level_list$hypothesis_4_cons[HOG_level_list$hypothesis_4_cons$expansion == "yes" 
                                                                    & HOG_level_list$hypothesis_4_cons$Arabidopsis_thaliana_sigDE>=1 & !is.na(HOG_level_list$hypothesis_4_cons$Arabidopsis_thaliana_sigDE) 
                                                                    & HOG_level_list$hypothesis_4_cons$Brassica_napus_sigDE>=1 & !is.na(HOG_level_list$hypothesis_4_cons$Brassica_napus_sigDE)
                                                                    ,]$HOG

### change the dataset and title here ###
dataset=HOGs_all_sens_exp_DEG_both_sens
title="HOGs_all_sens_exp_DEG_both_sens"
f <- ggplot(Not_min_2_expr_Conserved_with_expression_at_least_2_expressed_and_at_least_2_DEG_and_category_unique_HOGs[Not_min_2_expr_Conserved_with_expression_at_least_2_expressed_and_at_least_2_DEG_and_category_unique_HOGs$HOG %in% dataset,], aes(x="", fill=as.factor(category_of_regulation)))+
  geom_bar(width=1)+
  ggtitle(title)+
  coord_polar("y")

pdf(paste0(OUTDIR, "/Regulation_category_ONLY_at_least_2_DEG_with_and_without_DEGs_in_Hypothesis_4_set_only_padj_less_0.1.pdf"), 12, 8)
ggarrange(a, b, c, d, e, f, ncol=3, nrow=2, common.legend = TRUE, legend="bottom")
dev.off()

## Plot the same subset but as a stacked pie with the regulation categories in the Conserved Set as background

dataset=HOGs_all_sens_exp_DEG_both_sens
ex=length(Not_min_2_expr_Conserved_with_expression_at_least_2_expressed_and_at_least_2_DEG_and_category_unique_HOGs[Not_min_2_expr_Conserved_with_expression_at_least_2_expressed_and_at_least_2_DEG_and_category_unique_HOGs$HOG %in% dataset,]$HOG)
total=length(Not_min_2_expr_Conserved_with_expression_at_least_2_expressed_and_at_least_2_DEG_and_category_unique_HOGs$HOG)
geneset1 <-names(summary(as.factor(Not_min_2_expr_Conserved_with_expression_at_least_2_expressed_and_at_least_2_DEG_and_category_unique_HOGs$category_of_regulation)))
geneset2 <- names(summary(as.factor(Not_min_2_expr_Conserved_with_expression_at_least_2_expressed_and_at_least_2_DEG_and_category_unique_HOGs[Not_min_2_expr_Conserved_with_expression_at_least_2_expressed_and_at_least_2_DEG_and_category_unique_HOGs$HOG %in% dataset,]$category_of_regulation)))
group1 <- c(total, total, total)
group2 <- c(ex, ex, ex)
At1 <- as.integer(c(summary(as.factor(Not_min_2_expr_Conserved_with_expression_at_least_2_expressed_and_at_least_2_DEG_and_category_unique_HOGs$category_of_regulation))))
At2 <- as.integer(c(summary(as.factor(Not_min_2_expr_Conserved_with_expression_at_least_2_expressed_and_at_least_2_DEG_and_category_unique_HOGs[Not_min_2_expr_Conserved_with_expression_at_least_2_expressed_and_at_least_2_DEG_and_category_unique_HOGs$HOG %in% dataset,]$category_of_regulation))))
at <- data.frame(geneset1, geneset2, group1, group2, At1, At2)
#make 2 stacked bar charts and then make the y-axis with polar coordinates with log10 x-axis
pdf(paste0(OUTDIR, "/Regulation_category_ONLY_at_least_2_DEG_exp_and_DE_both_exp_Hypothesis_4_in_Conserved_set_only_padj_less_0.1.pdf"), 12, 8)
d <- ggplot(at)+
  geom_bar(aes(x=log10(total)/2, y=At1, fill=geneset1), width = log10(total), position="fill", stat="identity", color="white", alpha=0.4)+
  geom_bar(aes(x=log10(ex)/2, y=At2, fill=geneset2), width = log10(ex), position="fill", stat="identity", color="white", alpha=1)+
  coord_polar(theta = "y", start=0)+
  scale_y_continuous(name ="Subset = Both sensitive exp. and DE, Background = Conserved and DE", limits= NULL, breaks=NULL, expand = expansion(0))+
  scale_x_continuous(name ="Proportion of HOGs per regulation category", limits= c(0,log10(total)), breaks = NULL, expand = expansion(0))+
  labs(fill="Regulation category")+
  theme(
    legend.position = "right",
    panel.grid.major = element_line(linetype = "dotted", colour="white"),
    panel.background = element_rect(fill="transparent", colour="black"))+
  geom_label(aes(x=log10(ex)-log10(ex)/2, y=0,label=paste(ex)), color="black", size=7)+
  geom_label(aes(x=log10(total)-log10(total)/6, y=0,label=paste(total)), color="black", size=7)+
  scale_fill_manual(values = palette_OkabeIto_black[1:3])
d
dev.off()

# plot the same but the background is "HOGs_DEG_both_sens"
dataset=HOGs_all_sens_exp_DEG_both_sens
ex=length(Not_min_2_expr_Conserved_with_expression_at_least_2_expressed_and_at_least_2_DEG_and_category_unique_HOGs[Not_min_2_expr_Conserved_with_expression_at_least_2_expressed_and_at_least_2_DEG_and_category_unique_HOGs$HOG %in% dataset,]$HOG)
total=length(Not_min_2_expr_Conserved_with_expression_at_least_2_expressed_and_at_least_2_DEG_and_category_unique_HOGs[Not_min_2_expr_Conserved_with_expression_at_least_2_expressed_and_at_least_2_DEG_and_category_unique_HOGs$HOG %in% HOGs_DEG_both_sens,]$HOG)
geneset1 <-names(summary(as.factor(Not_min_2_expr_Conserved_with_expression_at_least_2_expressed_and_at_least_2_DEG_and_category_unique_HOGs$category_of_regulation)))
geneset2 <- names(summary(as.factor(Not_min_2_expr_Conserved_with_expression_at_least_2_expressed_and_at_least_2_DEG_and_category_unique_HOGs[Not_min_2_expr_Conserved_with_expression_at_least_2_expressed_and_at_least_2_DEG_and_category_unique_HOGs$HOG %in% dataset,]$category_of_regulation)))
group1 <- c(total, total, total)
group2 <- c(ex, ex, ex)
At1 <- as.integer(c(summary(as.factor(Not_min_2_expr_Conserved_with_expression_at_least_2_expressed_and_at_least_2_DEG_and_category_unique_HOGs[
  Not_min_2_expr_Conserved_with_expression_at_least_2_expressed_and_at_least_2_DEG_and_category_unique_HOGs$HOG %in% HOGs_DEG_both_sens,]$category_of_regulation))))
At2 <- as.integer(c(summary(as.factor(Not_min_2_expr_Conserved_with_expression_at_least_2_expressed_and_at_least_2_DEG_and_category_unique_HOGs[
  Not_min_2_expr_Conserved_with_expression_at_least_2_expressed_and_at_least_2_DEG_and_category_unique_HOGs$HOG %in% dataset,]$category_of_regulation))))
at <- data.frame(geneset1, geneset2, group1, group2, At1, At2)
#make 2 stacked bar charts and then make the y-axis with polar coordinates with log10 x-axis
pdf(paste0(OUTDIR, "/Regulation_category_ONLY_at_least_2_DEG_exp_and_DE_both_exp_Hypothesis_4_in_DE_both_sens_only_padj_less_0.1.pdf"), 12, 8)
d <- ggplot(at)+
  geom_bar(aes(x=log10(total)/2, y=At1, fill=geneset1), width = log10(total), position="fill", stat="identity", color="white", alpha=0.4)+
  geom_bar(aes(x=log10(ex)/2, y=At2, fill=geneset2), width = log10(ex), position="fill", stat="identity", color="white", alpha=1)+
  coord_polar(theta = "y", start=0)+
  scale_y_continuous(name ="Subset = Both sensitive exp. and DE, Background = both sens DE", limits= NULL, breaks=NULL, expand = expansion(0))+
  scale_x_continuous(name ="Proportion of HOGs per regulation category", limits= c(0,log10(total)), breaks = NULL, expand = expansion(0))+
  labs(fill="Regulation category")+
  theme(
    legend.position = "right",
    panel.grid.major = element_line(linetype = "dotted", colour="white"),
    panel.background = element_rect(fill="transparent", colour="black"))+
  geom_label(aes(x=log10(ex)-log10(ex)/2, y=0,label=paste(ex)), color="black", size=7)+
  geom_label(aes(x=log10(total)-log10(total)/6, y=0,label=paste(total)), color="black", size=7)+
  scale_fill_manual(values = palette_OkabeIto_black[1:3])
d
dev.off()

## test for enrichment with UP_and_DOWN

# how many are up_and_down?
length(Not_min_2_expr_Conserved_with_expression_at_least_2_expressed_and_at_least_2_DEG_and_category_unique_HOGs[Not_min_2_expr_Conserved_with_expression_at_least_2_expressed_and_at_least_2_DEG_and_category_unique_HOGs$HOG %in% dataset & Not_min_2_expr_Conserved_with_expression_at_least_2_expressed_and_at_least_2_DEG_and_category_unique_HOGs$category_of_regulation=="down and up",]$HOG)
# 4

# from how many categorized HOGs in total?
length(unique(Not_min_2_expr_Conserved_with_expression_at_least_2_expressed_and_at_least_2_DEG_and_category_unique_HOGs[Not_min_2_expr_Conserved_with_expression_at_least_2_expressed_and_at_least_2_DEG_and_category_unique_HOGs$HOG %in% dataset,]$HOG))
# 6

# In BG: how many are up and down?
length(Not_min_2_expr_Conserved_with_expression_at_least_2_expressed_and_at_least_2_DEG_and_category_unique_HOGs[Not_min_2_expr_Conserved_with_expression_at_least_2_expressed_and_at_least_2_DEG_and_category_unique_HOGs$HOG %in% HOGs_DEG_both_sens & Not_min_2_expr_Conserved_with_expression_at_least_2_expressed_and_at_least_2_DEG_and_category_unique_HOGs$category_of_regulation=="down and up",]$HOG)
# 591

# In BG: from how many HOGs in total?
length(unique(Not_min_2_expr_Conserved_with_expression_at_least_2_expressed_and_at_least_2_DEG_and_category_unique_HOGs[Not_min_2_expr_Conserved_with_expression_at_least_2_expressed_and_at_least_2_DEG_and_category_unique_HOGs$HOG %in% HOGs_DEG_both_sens,]$HOG))
# 1436

### test if the exp and DE in both tolerant are significantly enriched for "only up"
# 591 / 1436 = 0.412
# 4/6=

q <- 4
m <- 591
n <- (1436 - m)
k <- 6

p <- phyper((q-1), m, n, k, lower.tail=F)
p # 0.195

# --> the enrichment is not sign. (prob. due to small sample size)


### Hypothesis 5
# 2.1)n)1) What is the frequency of OGs with at least 1 DEG from the Hypothesis 5 expanded species in "Conserved" OGs?
HOGs_DEG_Athaliana <- HOG_level_list$hypothesis_5_cons[HOG_level_list$hypothesis_5_cons$Arabidopsis_thaliana_sigDE>=1 & !is.na(HOG_level_list$hypothesis_5_cons$Arabidopsis_thaliana_sigDE)
                                                       ,]$HOG
### change the dataset and title here ###
dataset=HOGs_DEG_Athaliana
title="HOGs_DEG_Athaliana"
a <- ggplot(Not_min_2_expr_Conserved_with_expression_at_least_2_expressed_and_at_least_2_DEG_and_category_unique_HOGs[Not_min_2_expr_Conserved_with_expression_at_least_2_expressed_and_at_least_2_DEG_and_category_unique_HOGs$HOG %in% dataset,], aes(x="", fill=as.factor(category_of_regulation)))+
  geom_bar(width=1)+
  ggtitle(title)+
  coord_polar("y")

# 2.1)m)2) Within "Conserved": Number of OGs with "Hypothesis 5 expanded==yes"
HOGs_Athaliana_exp <- HOG_level_list$hypothesis_5_cons[HOG_level_list$hypothesis_5_cons$expansion == "yes",]$HOG

### change the dataset and title here ###
dataset=HOGs_Athaliana_exp
title="HOGs_Athaliana_exp"
b <- ggplot(Not_min_2_expr_Conserved_with_expression_at_least_2_expressed_and_at_least_2_DEG_and_category_unique_HOGs[Not_min_2_expr_Conserved_with_expression_at_least_2_expressed_and_at_least_2_DEG_and_category_unique_HOGs$HOG %in% dataset,], aes(x="", fill=as.factor(category_of_regulation)))+
  geom_bar(width=1)+
  ggtitle(title)+
  coord_polar("y")

# 2m) Within "Conserved": What is the frequency of OGs with at least 1 DEG in "Hypothesis 5 expanded==yes" OGs?
cons_1_and_5_merged_2 <-  merge(HOG_level_list$hypothesis_1_cons[,c("HOG", "total_sigDE")], HOG_level_list$hypothesis_5_cons, by="HOG")
# select for total_sigDE.x because we are interested in total_sigDE from all 4 species (inferred from data frame HOG_level_list$hypothesis_1_cons, which contains all 4 species)
HOGs_Athaliana_exp_DEG_any <- cons_1_and_5_merged_2[cons_1_and_5_merged_2$expansion == "yes" & !is.na(cons_1_and_5_merged_2$total_sigDE.x) & cons_1_and_5_merged_2$total_sigDE.x>=1,]$HOG

### change the dataset and title here ###
dataset=HOGs_Athaliana_exp_DEG_any
title="HOGs_Athaliana_exp_DEG_any"
c <- ggplot(Not_min_2_expr_Conserved_with_expression_at_least_2_expressed_and_at_least_2_DEG_and_category_unique_HOGs[Not_min_2_expr_Conserved_with_expression_at_least_2_expressed_and_at_least_2_DEG_and_category_unique_HOGs$HOG %in% dataset,], aes(x="", fill=as.factor(category_of_regulation)))+
  geom_bar(width=1)+
  ggtitle(title)+
  coord_polar("y")

# 2n) What is the frequency of OGs with at least 1 DEG from any of the expanded species in "Hypothesis 5 expanded==yes" OGs?
HOGs_Athaliana_exp_DEG_Athaliana <- HOG_level_list$hypothesis_5_cons[HOG_level_list$hypothesis_5_cons$expansion == "yes" & HOG_level_list$hypothesis_5_cons$Arabidopsis_thaliana_sigDE>=1 & !is.na(HOG_level_list$hypothesis_5_cons$Arabidopsis_thaliana_sigDE)
                                                                     ,]$HOG

### change the dataset and title here ###
dataset=HOGs_Athaliana_exp_DEG_Athaliana
title="HOGs_Athaliana_exp_DEG_Athaliana"
d <- ggplot(Not_min_2_expr_Conserved_with_expression_at_least_2_expressed_and_at_least_2_DEG_and_category_unique_HOGs[Not_min_2_expr_Conserved_with_expression_at_least_2_expressed_and_at_least_2_DEG_and_category_unique_HOGs$HOG %in% dataset,], aes(x="", fill=as.factor(category_of_regulation)))+
  geom_bar(width=1)+
  ggtitle(title)+
  coord_polar("y")

pdf(paste0(OUTDIR, "/Regulation_category_ONLY_at_least_2_DEG_with_and_without_DEGs_in_Hypothesis_5_set_only_padj_less_0.1.pdf"), 12, 8)
ggarrange(a, b, c, d, ncol=2, nrow=2, common.legend = TRUE, legend="bottom")
dev.off()


## --> No real difference for Hypothesis 5

## Plot the same subset but as a stacked pie with the regulation categories in the Conserved Set as background

dataset=HOGs_Athaliana_exp_DEG_Athaliana
ex=length(Not_min_2_expr_Conserved_with_expression_at_least_2_expressed_and_at_least_2_DEG_and_category_unique_HOGs[Not_min_2_expr_Conserved_with_expression_at_least_2_expressed_and_at_least_2_DEG_and_category_unique_HOGs$HOG %in% dataset,]$HOG)
total=length(Not_min_2_expr_Conserved_with_expression_at_least_2_expressed_and_at_least_2_DEG_and_category_unique_HOGs$HOG)
geneset1 <-names(summary(as.factor(Not_min_2_expr_Conserved_with_expression_at_least_2_expressed_and_at_least_2_DEG_and_category_unique_HOGs$category_of_regulation)))
geneset2 <- names(summary(as.factor(Not_min_2_expr_Conserved_with_expression_at_least_2_expressed_and_at_least_2_DEG_and_category_unique_HOGs[Not_min_2_expr_Conserved_with_expression_at_least_2_expressed_and_at_least_2_DEG_and_category_unique_HOGs$HOG %in% dataset,]$category_of_regulation)))
group1 <- c(total, total, total)
group2 <- c(ex, ex, ex)
At1 <- as.integer(c(summary(as.factor(Not_min_2_expr_Conserved_with_expression_at_least_2_expressed_and_at_least_2_DEG_and_category_unique_HOGs$category_of_regulation))))
At2 <- as.integer(c(summary(as.factor(Not_min_2_expr_Conserved_with_expression_at_least_2_expressed_and_at_least_2_DEG_and_category_unique_HOGs[Not_min_2_expr_Conserved_with_expression_at_least_2_expressed_and_at_least_2_DEG_and_category_unique_HOGs$HOG %in% dataset,]$category_of_regulation))))
at <- data.frame(geneset1, geneset2, group1, group2, At1, At2)
#make 2 stacked bar charts and then make the y-axis with polar coordinates with log10 x-axis
pdf(paste0(OUTDIR, "/Regulation_category_ONLY_at_least_2_DEG_exp_and_DE_exp_Hypothesis_5_in_Conserved_set_only_padj_less_0.1.pdf"), 12, 8)
e <- ggplot(at)+
  geom_bar(aes(x=log10(total)/2, y=At1, fill=geneset1), width = log10(total), position="fill", stat="identity", color="white", alpha=0.4)+
  geom_bar(aes(x=log10(ex)/2, y=At2, fill=geneset2), width = log10(ex), position="fill", stat="identity", color="white", alpha=1)+
  coord_polar(theta = "y", start=0)+
  scale_y_continuous(name ="Subset = Athaliana exp. and DE, Background = Conserved and DE", limits= NULL, breaks=NULL, expand = expansion(0))+
  scale_x_continuous(name ="Proportion of HOGs per regulation category", limits= c(0,log10(total)), breaks = NULL, expand = expansion(0))+
  labs(fill="Regulation category")+
  theme(
    legend.position = "right",
    panel.grid.major = element_line(linetype = "dotted", colour="white"),
    panel.background = element_rect(fill="transparent", colour="black"))+
  geom_label(aes(x=log10(ex)-log10(ex)/2, y=0,label=paste(ex)), color="black", size=7)+
  geom_label(aes(x=log10(total)-log10(total)/6, y=0,label=paste(total)), color="black", size=7)+
  scale_fill_manual(values = palette_OkabeIto_black[1:3])
e
dev.off()

dataset=HOGs_Athaliana_exp_DEG_Athaliana
ex=length(Not_min_2_expr_Conserved_with_expression_at_least_2_expressed_and_at_least_2_DEG_and_category_unique_HOGs[Not_min_2_expr_Conserved_with_expression_at_least_2_expressed_and_at_least_2_DEG_and_category_unique_HOGs$HOG %in% dataset,]$HOG)
total=length(Not_min_2_expr_Conserved_with_expression_at_least_2_expressed_and_at_least_2_DEG_and_category_unique_HOGs[Not_min_2_expr_Conserved_with_expression_at_least_2_expressed_and_at_least_2_DEG_and_category_unique_HOGs$HOG %in% HOGs_DEG_Athaliana,]$HOG)
geneset1 <-names(summary(as.factor(Not_min_2_expr_Conserved_with_expression_at_least_2_expressed_and_at_least_2_DEG_and_category_unique_HOGs$category_of_regulation)))
geneset2 <- names(summary(as.factor(Not_min_2_expr_Conserved_with_expression_at_least_2_expressed_and_at_least_2_DEG_and_category_unique_HOGs[Not_min_2_expr_Conserved_with_expression_at_least_2_expressed_and_at_least_2_DEG_and_category_unique_HOGs$HOG %in% dataset,]$category_of_regulation)))
group1 <- c(total, total, total)
group2 <- c(ex, ex, ex)
At1 <- as.integer(c(summary(as.factor(Not_min_2_expr_Conserved_with_expression_at_least_2_expressed_and_at_least_2_DEG_and_category_unique_HOGs[
  Not_min_2_expr_Conserved_with_expression_at_least_2_expressed_and_at_least_2_DEG_and_category_unique_HOGs$HOG %in% HOGs_DEG_Athaliana,]$category_of_regulation))))
At2 <- as.integer(c(summary(as.factor(Not_min_2_expr_Conserved_with_expression_at_least_2_expressed_and_at_least_2_DEG_and_category_unique_HOGs[
  Not_min_2_expr_Conserved_with_expression_at_least_2_expressed_and_at_least_2_DEG_and_category_unique_HOGs$HOG %in% dataset,]$category_of_regulation))))
at <- data.frame(geneset1, geneset2, group1, group2, At1, At2)
#make 2 stacked bar charts and then make the y-axis with polar coordinates with log10 x-axis
pdf(paste0(OUTDIR, "/Regulation_category_ONLY_at_least_2_DEG_exp_and_DE_exp_Hypothesis_5_in_DEG_Athaliana_set_only_padj_less_0.1.pdf"), 12, 8)
e <- ggplot(at)+
  geom_bar(aes(x=log10(total)/2, y=At1, fill=geneset1), width = log10(total), position="fill", stat="identity", color="white", alpha=0.4)+
  geom_bar(aes(x=log10(ex)/2, y=At2, fill=geneset2), width = log10(ex), position="fill", stat="identity", color="white", alpha=1)+
  coord_polar(theta = "y", start=0)+
  scale_y_continuous(name ="Subset = Athaliana exp. and DE, Background = Athaliana DE", limits= NULL, breaks=NULL, expand = expansion(0))+
  scale_x_continuous(name ="Proportion of HOGs per regulation category", limits= c(0,log10(total)), breaks = NULL, expand = expansion(0))+
  labs(fill="Regulation category")+
  theme(
    legend.position = "right",
    panel.grid.major = element_line(linetype = "dotted", colour="white"),
    panel.background = element_rect(fill="transparent", colour="black"))+
  geom_label(aes(x=log10(ex)-log10(ex)/2, y=0,label=paste(ex)), color="black", size=7)+
  geom_label(aes(x=log10(total)-log10(total)/6, y=0,label=paste(total)), color="black", size=7)+
  scale_fill_manual(values = palette_OkabeIto_black[1:3])
e
dev.off()

## test for enrichment with UP_and_DOWN

# how many are up_and_down?
length(Not_min_2_expr_Conserved_with_expression_at_least_2_expressed_and_at_least_2_DEG_and_category_unique_HOGs[Not_min_2_expr_Conserved_with_expression_at_least_2_expressed_and_at_least_2_DEG_and_category_unique_HOGs$HOG %in% dataset & Not_min_2_expr_Conserved_with_expression_at_least_2_expressed_and_at_least_2_DEG_and_category_unique_HOGs$category_of_regulation=="down and up",]$HOG)
# 31

# from how many categorized HOGs in total?
length(unique(Not_min_2_expr_Conserved_with_expression_at_least_2_expressed_and_at_least_2_DEG_and_category_unique_HOGs[Not_min_2_expr_Conserved_with_expression_at_least_2_expressed_and_at_least_2_DEG_and_category_unique_HOGs$HOG %in% dataset,]$HOG))
# 79

# In BG: how many are up and down?
length(Not_min_2_expr_Conserved_with_expression_at_least_2_expressed_and_at_least_2_DEG_and_category_unique_HOGs[Not_min_2_expr_Conserved_with_expression_at_least_2_expressed_and_at_least_2_DEG_and_category_unique_HOGs$HOG %in% HOGs_DEG_Athaliana & Not_min_2_expr_Conserved_with_expression_at_least_2_expressed_and_at_least_2_DEG_and_category_unique_HOGs$category_of_regulation=="down and up",]$HOG)
# 716

# In BG: from how many HOGs in total?
length(unique(Not_min_2_expr_Conserved_with_expression_at_least_2_expressed_and_at_least_2_DEG_and_category_unique_HOGs[Not_min_2_expr_Conserved_with_expression_at_least_2_expressed_and_at_least_2_DEG_and_category_unique_HOGs$HOG %in% HOGs_DEG_Athaliana,]$HOG))
# 2591

### test if the exp and DE in both tolerant are significantly enriched for "only up"
# 716 / 2591 = 0.276
# 31/79= 0.3924

q <- 31
m <- 716
n <- (2591 - m)
k <- 79

p <- phyper((q-1), m, n, k, lower.tail=F)
p # 0.015

# --> the enrichment with up_and_down is sign.

## test for enrichment with only UP

# how many are only up?
length(Not_min_2_expr_Conserved_with_expression_at_least_2_expressed_and_at_least_2_DEG_and_category_unique_HOGs[Not_min_2_expr_Conserved_with_expression_at_least_2_expressed_and_at_least_2_DEG_and_category_unique_HOGs$HOG %in% dataset & Not_min_2_expr_Conserved_with_expression_at_least_2_expressed_and_at_least_2_DEG_and_category_unique_HOGs$category_of_regulation=="up",]$HOG)
# 25

# from how many categorized HOGs in total?
length(unique(Not_min_2_expr_Conserved_with_expression_at_least_2_expressed_and_at_least_2_DEG_and_category_unique_HOGs[Not_min_2_expr_Conserved_with_expression_at_least_2_expressed_and_at_least_2_DEG_and_category_unique_HOGs$HOG %in% dataset,]$HOG))
# 79

# In BG: how many are only up?
length(Not_min_2_expr_Conserved_with_expression_at_least_2_expressed_and_at_least_2_DEG_and_category_unique_HOGs[Not_min_2_expr_Conserved_with_expression_at_least_2_expressed_and_at_least_2_DEG_and_category_unique_HOGs$HOG %in% HOGs_DEG_Athaliana & Not_min_2_expr_Conserved_with_expression_at_least_2_expressed_and_at_least_2_DEG_and_category_unique_HOGs$category_of_regulation=="up",]$HOG)
# 695

# In BG: from how many HOGs in total?
length(unique(Not_min_2_expr_Conserved_with_expression_at_least_2_expressed_and_at_least_2_DEG_and_category_unique_HOGs[Not_min_2_expr_Conserved_with_expression_at_least_2_expressed_and_at_least_2_DEG_and_category_unique_HOGs$HOG %in% HOGs_DEG_Athaliana,]$HOG))
# 2591

### test if the exp and DE in both tolerant are significantly enriched for "only up"
# 695 / 2591 = 0.268
# 25/ 79 = 0.3164

q <- 25
m <- 695
n <- (2591 - m)
k <- 79

p <- phyper((q-1), m, n, k, lower.tail=F)
p # 0.195

# --> the increase in only UP is not. Sign.

### Hypothesis 6
# 2.1)p)1) What is the frequency of OGs with at least 1 DEG from the Hypothesis 6 expanded species in "Conserved" OGs?
HOGs_DEG_Bnapus <- HOG_level_list$hypothesis_6_cons[HOG_level_list$hypothesis_6_cons$Brassica_napus_sigDE>=1 & !is.na(HOG_level_list$hypothesis_6_cons$Brassica_napus_sigDE)
                                                    ,]$HOG

### change the dataset and title here ###
dataset=HOGs_DEG_Bnapus
title="HOGs_DEG_Bnapus"
a <- ggplot(Not_min_2_expr_Conserved_with_expression_at_least_2_expressed_and_at_least_2_DEG_and_category_unique_HOGs[Not_min_2_expr_Conserved_with_expression_at_least_2_expressed_and_at_least_2_DEG_and_category_unique_HOGs$HOG %in% dataset,], aes(x="", fill=as.factor(category_of_regulation)))+
  geom_bar(width=1)+
  ggtitle(title)+
  coord_polar("y")

# 2.1)o)2) Within "Conserved": Number of OGs with "Hypothesis 6 expanded==yes"
HOGs_Bnapus_exp <- HOG_level_list$hypothesis_6_cons[HOG_level_list$hypothesis_6_cons$expansion == "yes",]$HOG

### change the dataset and title here ###
dataset=HOGs_Bnapus_exp
title="HOGs_Bnapus_exp"
b <- ggplot(Not_min_2_expr_Conserved_with_expression_at_least_2_expressed_and_at_least_2_DEG_and_category_unique_HOGs[Not_min_2_expr_Conserved_with_expression_at_least_2_expressed_and_at_least_2_DEG_and_category_unique_HOGs$HOG %in% dataset,], aes(x="", fill=as.factor(category_of_regulation)))+
  geom_bar(width=1)+
  ggtitle(title)+
  coord_polar("y")

# 2o) Within "Conserved": What is the frequency of OGs with at least 1 DEG in "Hypothesis 6 expanded==yes" OGs?
cons_1_and_6_merged_2 <-  merge(HOG_level_list$hypothesis_1_cons[,c("HOG", "total_sigDE")], HOG_level_list$hypothesis_6_cons, by="HOG")
# select for total_sigDE.x because we are interested in total_sigDE from all 4 species (inferred from data frame HOG_level_list$hypothesis_1_cons, which contains all 4 species)
HOGs_Bnapus_exp_DEG_any <- cons_1_and_6_merged_2[cons_1_and_6_merged_2$expansion == "yes" & !is.na(cons_1_and_6_merged_2$total_sigDE.x) & cons_1_and_6_merged_2$total_sigDE.x>=1,]$HOG

### change the dataset and title here ###
dataset=HOGs_Bnapus_exp_DEG_any
title="HOGs_Bnapus_exp_DEG_any"
c <- ggplot(Not_min_2_expr_Conserved_with_expression_at_least_2_expressed_and_at_least_2_DEG_and_category_unique_HOGs[Not_min_2_expr_Conserved_with_expression_at_least_2_expressed_and_at_least_2_DEG_and_category_unique_HOGs$HOG %in% dataset,], aes(x="", fill=as.factor(category_of_regulation)))+
  geom_bar(width=1)+
  ggtitle(title)+
  coord_polar("y")

# 2p) What is the frequency of OGs with at least 1 DEG from any of the expanded species in "Hypothesis 6 expanded==yes" OGs?
HOGs_Bnapus_exp_DEG_Bnapus <- HOG_level_list$hypothesis_6_cons[HOG_level_list$hypothesis_6_cons$expansion == "yes" & HOG_level_list$hypothesis_6_cons$Brassica_napus_sigDE>=1 & !is.na(HOG_level_list$hypothesis_6_cons$Brassica_napus_sigDE)
                                                               ,]$HOG
### change the dataset and title here ###
dataset=HOGs_Bnapus_exp_DEG_Bnapus
title="HOGs_Bnapus_exp_DEG_Bnapus"
d <- ggplot(Not_min_2_expr_Conserved_with_expression_at_least_2_expressed_and_at_least_2_DEG_and_category_unique_HOGs[Not_min_2_expr_Conserved_with_expression_at_least_2_expressed_and_at_least_2_DEG_and_category_unique_HOGs$HOG %in% dataset,], aes(x="", fill=as.factor(category_of_regulation)))+
  geom_bar(width=1)+
  ggtitle(title)+
  coord_polar("y")

pdf(paste0(OUTDIR, "/Regulation_category_ONLY_at_least_2_DEG_with_and_without_DEGs_in_Hypothesis_6_set_only_padj_less_0.1.pdf"), 12, 8)
ggarrange(a, b, c, d, ncol=2, nrow=2, common.legend = TRUE, legend="bottom")
dev.off()

## --> No real difference for Hypothesis 6

## Plot the same subset but as a stacked pie with the regulation categories in the Conserved Set as background

dataset=HOGs_Bnapus_exp_DEG_Bnapus
ex=length(Not_min_2_expr_Conserved_with_expression_at_least_2_expressed_and_at_least_2_DEG_and_category_unique_HOGs[Not_min_2_expr_Conserved_with_expression_at_least_2_expressed_and_at_least_2_DEG_and_category_unique_HOGs$HOG %in% dataset,]$HOG)
total=length(Not_min_2_expr_Conserved_with_expression_at_least_2_expressed_and_at_least_2_DEG_and_category_unique_HOGs$HOG)
geneset1 <-names(summary(as.factor(Not_min_2_expr_Conserved_with_expression_at_least_2_expressed_and_at_least_2_DEG_and_category_unique_HOGs$category_of_regulation)))
geneset2 <- names(summary(as.factor(Not_min_2_expr_Conserved_with_expression_at_least_2_expressed_and_at_least_2_DEG_and_category_unique_HOGs[Not_min_2_expr_Conserved_with_expression_at_least_2_expressed_and_at_least_2_DEG_and_category_unique_HOGs$HOG %in% dataset,]$category_of_regulation)))
group1 <- c(total, total, total)
group2 <- c(ex, ex, ex)
At1 <- as.integer(c(summary(as.factor(Not_min_2_expr_Conserved_with_expression_at_least_2_expressed_and_at_least_2_DEG_and_category_unique_HOGs$category_of_regulation))))
At2 <- as.integer(c(summary(as.factor(Not_min_2_expr_Conserved_with_expression_at_least_2_expressed_and_at_least_2_DEG_and_category_unique_HOGs[Not_min_2_expr_Conserved_with_expression_at_least_2_expressed_and_at_least_2_DEG_and_category_unique_HOGs$HOG %in% dataset,]$category_of_regulation))))
at <- data.frame(geneset1, geneset2, group1, group2, At1, At2)
#make 2 stacked bar charts and then make the y-axis with polar coordinates with log10 x-axis
pdf(paste0(OUTDIR, "/Regulation_category_ONLY_at_least_2_DEG_exp_and_DE_exp_Hypothesis_6_in_Conserved_set_only_padj_less_0.1.pdf"), 12, 8)
f <- ggplot(at)+
  geom_bar(aes(x=log10(total)/2, y=At1, fill=geneset1), width = log10(total), position="fill", stat="identity", color="white", alpha=0.4)+
  geom_bar(aes(x=log10(ex)/2, y=At2, fill=geneset2), width = log10(ex), position="fill", stat="identity", color="white", alpha=1)+
  coord_polar(theta = "y", start=0)+
  scale_y_continuous(name ="Subset = Bnapus exp. and DE, Background = Conserved and DE", limits= NULL, breaks=NULL, expand = expansion(0))+
  scale_x_continuous(name ="Proportion of HOGs per regulation category", limits= c(0,log10(total)), breaks = NULL, expand = expansion(0))+
  labs(fill="Regulation category")+
  theme(
    legend.position = "right",
    panel.grid.major = element_line(linetype = "dotted", colour="white"),
    panel.background = element_rect(fill="transparent", colour="black"))+
  geom_label(aes(x=log10(ex)-log10(ex)/2, y=0,label=paste(ex)), color="black", size=7)+
  geom_label(aes(x=log10(total)-log10(total)/6, y=0,label=paste(total)), color="black", size=7)+
  scale_fill_manual(values = palette_OkabeIto_black[1:3])
f
dev.off()

# with background DE in Bnapus
dataset=HOGs_Bnapus_exp_DEG_Bnapus
ex=length(Not_min_2_expr_Conserved_with_expression_at_least_2_expressed_and_at_least_2_DEG_and_category_unique_HOGs[Not_min_2_expr_Conserved_with_expression_at_least_2_expressed_and_at_least_2_DEG_and_category_unique_HOGs$HOG %in% dataset,]$HOG)
total=length(Not_min_2_expr_Conserved_with_expression_at_least_2_expressed_and_at_least_2_DEG_and_category_unique_HOGs[Not_min_2_expr_Conserved_with_expression_at_least_2_expressed_and_at_least_2_DEG_and_category_unique_HOGs$HOG %in% HOGs_DEG_Bnapus,]$HOG)
geneset1 <-names(summary(as.factor(Not_min_2_expr_Conserved_with_expression_at_least_2_expressed_and_at_least_2_DEG_and_category_unique_HOGs$category_of_regulation)))
geneset2 <- names(summary(as.factor(Not_min_2_expr_Conserved_with_expression_at_least_2_expressed_and_at_least_2_DEG_and_category_unique_HOGs[Not_min_2_expr_Conserved_with_expression_at_least_2_expressed_and_at_least_2_DEG_and_category_unique_HOGs$HOG %in% dataset,]$category_of_regulation)))
group1 <- c(total, total, total)
group2 <- c(ex, ex, ex)
At1 <- as.integer(c(summary(as.factor(Not_min_2_expr_Conserved_with_expression_at_least_2_expressed_and_at_least_2_DEG_and_category_unique_HOGs[Not_min_2_expr_Conserved_with_expression_at_least_2_expressed_and_at_least_2_DEG_and_category_unique_HOGs$HOG %in% HOGs_DEG_Bnapus
                                                                                      ,]$category_of_regulation))))
At2 <- as.integer(c(summary(as.factor(Not_min_2_expr_Conserved_with_expression_at_least_2_expressed_and_at_least_2_DEG_and_category_unique_HOGs[Not_min_2_expr_Conserved_with_expression_at_least_2_expressed_and_at_least_2_DEG_and_category_unique_HOGs$HOG %in% dataset
                                                                                      ,]$category_of_regulation))))
at <- data.frame(geneset1, geneset2, group1, group2, At1, At2)
#make 2 stacked bar charts and then make the y-axis with polar coordinates with log10 x-axis
pdf(paste0(OUTDIR, "/Regulation_category_ONLY_at_least_2_DEG_exp_and_DE_exp_Hypothesis_6_in_Bnapus_DE_only_padj_less_0.1.pdf"), 12, 8)
f <- ggplot(at)+
  geom_bar(aes(x=log10(total)/2, y=At1, fill=geneset1), width = log10(total), position="fill", stat="identity", color="white", alpha=0.4)+
  geom_bar(aes(x=log10(ex)/2, y=At2, fill=geneset2), width = log10(ex), position="fill", stat="identity", color="white", alpha=1)+
  coord_polar(theta = "y", start=0)+
  scale_y_continuous(name ="Subset = Bnapus exp. and DE, Background = Bnapus DE", limits= NULL, breaks=NULL, expand = expansion(0))+
  scale_x_continuous(name ="Proportion of HOGs per regulation category", limits= c(0,log10(total)), breaks = NULL, expand = expansion(0))+
  labs(fill="Regulation category")+
  theme(
    legend.position = "right",
    panel.grid.major = element_line(linetype = "dotted", colour="white"),
    panel.background = element_rect(fill="transparent", colour="black"))+
  geom_label(aes(x=log10(ex)-log10(ex)/2, y=0,label=paste(ex)), color="black", size=7)+
  geom_label(aes(x=log10(total)-log10(total)/6, y=0,label=paste(total)), color="black", size=7)+
  scale_fill_manual(values = palette_OkabeIto_black[1:3])
f
dev.off()


## test for enrichment with UP_and_DOWN
# 398/851 = 0.4676851
# bg 2679/5909 = 0.4533762

# how many are up_and_down?
length(Not_min_2_expr_Conserved_with_expression_at_least_2_expressed_and_at_least_2_DEG_and_category_unique_HOGs[Not_min_2_expr_Conserved_with_expression_at_least_2_expressed_and_at_least_2_DEG_and_category_unique_HOGs$HOG %in% dataset & Not_min_2_expr_Conserved_with_expression_at_least_2_expressed_and_at_least_2_DEG_and_category_unique_HOGs$category_of_regulation=="down and up",]$HOG)
# 398

# from how many categorized HOGs in total?
length(unique(Not_min_2_expr_Conserved_with_expression_at_least_2_expressed_and_at_least_2_DEG_and_category_unique_HOGs[Not_min_2_expr_Conserved_with_expression_at_least_2_expressed_and_at_least_2_DEG_and_category_unique_HOGs$HOG %in% dataset,]$HOG))
# 851

# In BG: how many are up and down?
length(Not_min_2_expr_Conserved_with_expression_at_least_2_expressed_and_at_least_2_DEG_and_category_unique_HOGs[Not_min_2_expr_Conserved_with_expression_at_least_2_expressed_and_at_least_2_DEG_and_category_unique_HOGs$HOG %in% HOGs_DEG_Bnapus & Not_min_2_expr_Conserved_with_expression_at_least_2_expressed_and_at_least_2_DEG_and_category_unique_HOGs$category_of_regulation=="down and up",]$HOG)
# 2679

# In BG: from how many HOGs in total?
length(unique(Not_min_2_expr_Conserved_with_expression_at_least_2_expressed_and_at_least_2_DEG_and_category_unique_HOGs[Not_min_2_expr_Conserved_with_expression_at_least_2_expressed_and_at_least_2_DEG_and_category_unique_HOGs$HOG %in% HOGs_DEG_Bnapus,]$HOG))
# 5909

### test if the exp and DE in Bna are significantly enriched for "up and down"
q <- 398
m <- 2679
n <- (5909 - m)
k <- 851

p <- phyper((q-1), m, n, k, lower.tail=F)
p # 0.192

# --> the enrichment with up_and_down is NOT sign.


#### 20240206 ####### --> makes no real difference in any subset, if we  apply the filter for expressed = basemean <= 10 or if we do not apply it !!!


###### 20230918: PER SPECIES: UpSetR plots = selection criterion: plot the distribution of OGs and their regulation pattern specific per species in an UpSetR plot
# For each HOG, the value of the new column "category_of_regulation" is -1 where all values of "log2FoldChange" are <0 for this HOG
# and select only those gene families which contain at least 2 DEGs
### without selecting basemean >=10 (=Not_min_2_expr_...) (20240821)

Not_min_2_expr_Conserved_with_expression_at_least_2_DEG_and_species_specific_category <-
  Conserved_with_expression_at_least_2_expressed_and_at_least_2_DEG[Conserved_with_expression_at_least_2_expressed_and_at_least_2_DEG$two_DEG=="yes" & !is.na(Conserved_with_expression_at_least_2_expressed$log2FoldChange) & !is.na(Conserved_with_expression_at_least_2_expressed$padj) & (Conserved_with_expression_at_least_2_expressed$padj<=0.1),] %>%
  group_by(HOG, species) %>%
  mutate(category_of_regulation_per_species=case_when(all(log2FoldChange < 0) ~ "down" ,
                                                      all(log2FoldChange > 0) ~ "up",
                                                      any(log2FoldChange < 0) && any(log2FoldChange > 0) ~ "down_and_up"))

Not_min_2_expr_Conserved_with_expression_at_least_2_DEG_and_species_specific_category$category_of_regulation_per_species_with_name <-
  paste(Not_min_2_expr_Conserved_with_expression_at_least_2_DEG_and_species_specific_category$species, Not_min_2_expr_Conserved_with_expression_at_least_2_DEG_and_species_specific_category$category_of_regulation_per_species, sep=" ")

### length(unique(Not_min_2_expr_Conserved_with_expression_at_least_2_DEG_and_species_specific_category$HOG))
# 8800

## Plot the regulation categories per species in an UpSetR plot:
library("UpSetR")
listInput <- list(Eutrema_down = Not_min_2_expr_Conserved_with_expression_at_least_2_DEG_and_species_specific_category[Not_min_2_expr_Conserved_with_expression_at_least_2_DEG_and_species_specific_category$category_of_regulation_per_species_with_name=="Eutrema_salsugineum down",]$HOG, 
                  Alyrata_down = Not_min_2_expr_Conserved_with_expression_at_least_2_DEG_and_species_specific_category[Not_min_2_expr_Conserved_with_expression_at_least_2_DEG_and_species_specific_category$category_of_regulation_per_species_with_name=="Arabidopsis_lyrata down",]$HOG,
                  Athaliana_down = Not_min_2_expr_Conserved_with_expression_at_least_2_DEG_and_species_specific_category[Not_min_2_expr_Conserved_with_expression_at_least_2_DEG_and_species_specific_category$category_of_regulation_per_species_with_name=="Arabidopsis_thaliana down",]$HOG,
                  Bnapus_down = Not_min_2_expr_Conserved_with_expression_at_least_2_DEG_and_species_specific_category[Not_min_2_expr_Conserved_with_expression_at_least_2_DEG_and_species_specific_category$category_of_regulation_per_species_with_name=="Brassica_napus down",]$HOG,
                  Eutrema_up = Not_min_2_expr_Conserved_with_expression_at_least_2_DEG_and_species_specific_category[Not_min_2_expr_Conserved_with_expression_at_least_2_DEG_and_species_specific_category$category_of_regulation_per_species_with_name=="Eutrema_salsugineum up",]$HOG, 
                  Alyrata_up = Not_min_2_expr_Conserved_with_expression_at_least_2_DEG_and_species_specific_category[Not_min_2_expr_Conserved_with_expression_at_least_2_DEG_and_species_specific_category$category_of_regulation_per_species_with_name=="Arabidopsis_lyrata up",]$HOG,
                  Athaliana_up = Not_min_2_expr_Conserved_with_expression_at_least_2_DEG_and_species_specific_category[Not_min_2_expr_Conserved_with_expression_at_least_2_DEG_and_species_specific_category$category_of_regulation_per_species_with_name=="Arabidopsis_thaliana up",]$HOG,
                  Bnapus_up = Not_min_2_expr_Conserved_with_expression_at_least_2_DEG_and_species_specific_category[Not_min_2_expr_Conserved_with_expression_at_least_2_DEG_and_species_specific_category$category_of_regulation_per_species_with_name=="Brassica_napus up",]$HOG,
                  Eutrema_down_and_up = Not_min_2_expr_Conserved_with_expression_at_least_2_DEG_and_species_specific_category[Not_min_2_expr_Conserved_with_expression_at_least_2_DEG_and_species_specific_category$category_of_regulation_per_species_with_name=="Eutrema_salsugineum down_and_up",]$HOG, 
                  Alyrata_down_and_up = Not_min_2_expr_Conserved_with_expression_at_least_2_DEG_and_species_specific_category[Not_min_2_expr_Conserved_with_expression_at_least_2_DEG_and_species_specific_category$category_of_regulation_per_species_with_name=="Arabidopsis_lyrata down_and_up",]$HOG,
                  Athaliana_down_and_up = Not_min_2_expr_Conserved_with_expression_at_least_2_DEG_and_species_specific_category[Not_min_2_expr_Conserved_with_expression_at_least_2_DEG_and_species_specific_category$category_of_regulation_per_species_with_name=="Arabidopsis_thaliana down_and_up",]$HOG,
                  Bnapus_down_and_up = Not_min_2_expr_Conserved_with_expression_at_least_2_DEG_and_species_specific_category[Not_min_2_expr_Conserved_with_expression_at_least_2_DEG_and_species_specific_category$category_of_regulation_per_species_with_name=="Brassica_napus down_and_up",]$HOG
)
names(listInput) <- c("Eutrema salsugineum down",
                      "Arabidopsis lyrata down",
                      "Arabidopsis thaliana down",
                      "Brassica napus down",
                      "Eutrema salsugineum up",
                      "Arabidopsis lyrata up",
                      "Arabidopsis thaliana up",
                      "Brassica napus up",
                      "Eutrema salsugineum down and up",
                      "Arabidopsis lyrata down and up",
                      "Arabidopsis thaliana down and up",
                      "Brassica napus down and up")

### color the rows per group (up, down or up_and_down)
metadata <-
  data.frame(names(listInput), c(rep("down",4), rep("up", 4), rep("down_and_up", 4)))
names(metadata) = c("sets", "category_of_regulation")

pdf(paste0(OUTDIR, "/Intersection_of_OGs_with_regulation_categories_per_species_ONLY_at_least_2_sig_Conserved.pdf"), 16, 8, onefile = FALSE)
a <- upset(fromList(listInput), nsets=12, nintersects = 40, order.by = "freq", 
      sets.x.label = "Number of HOGs per \nspecies-specific regulation category \n- cut this set size plot",
      mainbar.y.label = "Size of Intersection",
      set.metadata = list(data=metadata, plots=list(list(type="matrix_rows", column="category_of_regulation",
      colors=c(down = palette_OkabeIto_black[1:3][1], down_and_up = palette_OkabeIto_black[1:3][2], up = palette_OkabeIto_black[1:3][3]),alpha=0.6))),
      # and color the intersects per total regulation category
      queries =
        list(
          list(query = intersects, params=list("Arabidopsis lyrata up", "Brassica napus down"), color= palette_OkabeIto_black[1:3][2], active = T),
          list(query = intersects, params=list("Arabidopsis lyrata up","Eutrema salsugineum up", "Brassica napus down"), color= palette_OkabeIto_black[1:3][2], active = T),
          list(query = intersects, params=list("Arabidopsis lyrata up","Eutrema salsugineum down", "Brassica napus up"), color= palette_OkabeIto_black[1:3][2], active = T),
          list(query = intersects, params=list("Arabidopsis lyrata down", "Brassica napus up"), color= palette_OkabeIto_black[1:3][2], active = T),
          list(query = intersects, params=list("Brassica napus up","Eutrema salsugineum down"), color= palette_OkabeIto_black[1:3][2], active = T),
          list(query = intersects, params=list("Brassica napus up","Eutrema salsugineum down", "Arabidopsis lyrata down"), color= palette_OkabeIto_black[1:3][2], active = T),
          list(query = intersects, params=list("Arabidopsis lyrata down", "Arabidopsis thaliana down", "Eutrema salsugineum down", "Brassica napus up"), color= palette_OkabeIto_black[1:3][2], active = T),
          list(query = intersects, params=list("Arabidopsis lyrata down","Eutrema salsugineum up"), color= palette_OkabeIto_black[1:3][2], active = T),
          list(query = intersects, params=list("Arabidopsis lyrata up","Eutrema salsugineum down"), color= palette_OkabeIto_black[1:3][2], active = T),
          list(query = intersects, params=list("Arabidopsis lyrata up","Eutrema salsugineum down", "Brassica napus up"), color= palette_OkabeIto_black[1:3][2], active = T),
          list(query = intersects, params=list("Arabidopsis lyrata up","Eutrema salsugineum down", "Brassica napus down"), color= palette_OkabeIto_black[1:3][2], active = T),
          list(query = intersects, params=list("Arabidopsis lyrata up","Eutrema salsugineum up", "Brassica napus down", "Arabidopsis thaliana up"), color= palette_OkabeIto_black[1:3][2], active = T),
          list(query = intersects, params=list("Brassica napus down","Arabidopsis lyrata down","Eutrema salsugineum up"), color= palette_OkabeIto_black[1:3][2], active = T),
          list(query = intersects, params=list("Arabidopsis lyrata down", "Arabidopsis thaliana down", "Eutrema salsugineum down", "Brassica napus up"), color= palette_OkabeIto_black[1:3][2], active = T),
          list(query = intersects, params=list("Arabidopsis lyrata down","Eutrema salsugineum up"), color= palette_OkabeIto_black[1:3][2], active = T),
          list(query = intersects, params=list("Arabidopsis lyrata up","Eutrema salsugineum down"), color= palette_OkabeIto_black[1:3][2], active = T),
          list(query = intersects, params=list("Arabidopsis lyrata up","Eutrema salsugineum down"), color= palette_OkabeIto_black[1:3][2], active = T),
          list(query = intersects, params=list("Arabidopsis lyrata up","Eutrema salsugineum down", "Brassica napus up"), color= palette_OkabeIto_black[1:3][2], active = T),
          list(query = intersects, params=list("Arabidopsis lyrata up","Eutrema salsugineum down", "Brassica napus down"), color= palette_OkabeIto_black[1:3][2], active = T),
          list(query = intersects, params=list("Arabidopsis lyrata up", "Arabidopsis thaliana up", "Eutrema salsugineum up", "Brassica napus down"), color= palette_OkabeIto_black[1:3][2], active = T),
          list(query = intersects, params=list("Arabidopsis lyrata down","Eutrema salsugineum up", "Brassica napus down"), color= palette_OkabeIto_black[1:3][2], active = T),
          list(query = intersects, params=list("Arabidopsis lyrata down","Arabidopsis thaliana down", "Brassica napus up"), color= palette_OkabeIto_black[1:3][2], active = T),
          list(query = intersects, params=list("Eutrema salsugineum down", "Arabidopsis thaliana down", "Brassica napus up"), color= palette_OkabeIto_black[1:3][2], active = T),
          list(query = intersects, params=list("Eutrema salsugineum down", "Arabidopsis lyrata down", "Brassica napus down and up"), color= palette_OkabeIto_black[1:3][2], active = T),
          list(query = intersects, params=list("Arabidopsis lyrata up","Arabidopsis thaliana up", "Brassica napus down"), color= palette_OkabeIto_black[1:3][2], active = T),
          list(query = intersects, params=list("Eutrema salsugineum up", "Brassica napus down"), color= palette_OkabeIto_black[1:3][2], active = T)
                                      ),
      matrix.color = "white",
      matrix.dot.alpha=0.3, 
      shade.alpha = 0,
      set_size.show = TRUE,
      set_size.scale_max = 4100,
      number.angles = 0,
      point.size = 2, line.size = 1,
      text.scale = c(2,1.5,1.6,1.5,1.5,1.5)
)
a
dev.off()

### Plot the same for the expanded sets ###

# Save expanded HOGs in a vector
### Hypothesis 1
Hypothesis_1_exp_OGs <- unique(data.frame(read.table(paste0(OUTDIR, "Hypothesis_1_Exp_in_Conserved_dataset_matrix.tsv"), header=T))$HOG)
### Hypothesis 2
Hypothesis_2_exp_OGs <- unique(data.frame(read.table(paste0(OUTDIR, "Hypothesis_2_Exp_in_Conserved_dataset_matrix.tsv"), header=T))$HOG)
### Hypothesis 3
Hypothesis_3_exp_OGs <- unique(data.frame(read.table(paste0(OUTDIR, "Hypothesis_3_Exp_in_Conserved_dataset_matrix.tsv"), header=T))$HOG)
### Hypothesis 4
Hypothesis_4_exp_OGs <- unique(data.frame(read.table(paste0(OUTDIR, "Hypothesis_4_Exp_in_Conserved_dataset_matrix.tsv"), header=T))$HOG)
### Hypothesis 5
Hypothesis_5_exp_OGs <- unique(data.frame(read.table(paste0(OUTDIR, "Hypothesis_5_Exp_in_Conserved_dataset_matrix.tsv"), header=T))$HOG)
### Hypothesis 6
Hypothesis_6_exp_OGs <- unique(data.frame(read.table(paste0(OUTDIR, "Hypothesis_6_Exp_in_Conserved_dataset_matrix.tsv"), header=T))$HOG)

# Select the expanded HOGs from the A2TEA_dataframe:
### Hypothesis 1
Hypothesis_1_Not_min_2_expr_Conserved_with_expression_at_least_2_DEG_and_species_specific_category <- Not_min_2_expr_Conserved_with_expression_at_least_2_DEG_and_species_specific_category[Not_min_2_expr_Conserved_with_expression_at_least_2_DEG_and_species_specific_category$HOG %in% Hypothesis_1_exp_OGs,]
# length(unique(Hypothesis_1_Not_min_2_expr_Conserved_with_expression_at_least_2_DEG_and_species_specific_category$HOG))
# not 73
# but 75

### Hypothesis 2
Hypothesis_2_Not_min_2_expr_Conserved_with_expression_at_least_2_DEG_and_species_specific_category <- Not_min_2_expr_Conserved_with_expression_at_least_2_DEG_and_species_specific_category[Not_min_2_expr_Conserved_with_expression_at_least_2_DEG_and_species_specific_category$HOG %in% Hypothesis_2_exp_OGs,]
# length(unique(Hypothesis_2_Not_min_2_expr_Conserved_with_expression_at_least_2_DEG_and_species_specific_category$HOG))
# not 361
# but 366

### Hypothesis 3
Hypothesis_3_Not_min_2_expr_Conserved_with_expression_at_least_2_DEG_and_species_specific_category <- Not_min_2_expr_Conserved_with_expression_at_least_2_DEG_and_species_specific_category[Not_min_2_expr_Conserved_with_expression_at_least_2_DEG_and_species_specific_category$HOG %in% Hypothesis_3_exp_OGs,]
# length(unique(Hypothesis_3_Not_min_2_expr_Conserved_with_expression_at_least_2_DEG_and_species_specific_category$HOG))
# not 531
# but 537

### Hypothesis 4
Hypothesis_4_Not_min_2_expr_Conserved_with_expression_at_least_2_DEG_and_species_specific_category <- Not_min_2_expr_Conserved_with_expression_at_least_2_DEG_and_species_specific_category[Not_min_2_expr_Conserved_with_expression_at_least_2_DEG_and_species_specific_category$HOG %in% Hypothesis_4_exp_OGs,]
# length(unique(Hypothesis_4_Not_min_2_expr_Conserved_with_expression_at_least_2_DEG_and_species_specific_category$HOG))
# stays 32

### Hypothesis 5
Hypothesis_5_Not_min_2_expr_Conserved_with_expression_at_least_2_DEG_and_species_specific_category <- Not_min_2_expr_Conserved_with_expression_at_least_2_DEG_and_species_specific_category[Not_min_2_expr_Conserved_with_expression_at_least_2_DEG_and_species_specific_category$HOG %in% Hypothesis_5_exp_OGs,]
# length(unique(Hypothesis_5_Not_min_2_expr_Conserved_with_expression_at_least_2_DEG_and_species_specific_category$HOG))
# not 189
# but 191

### Hypothesis 6
Hypothesis_6_Not_min_2_expr_Conserved_with_expression_at_least_2_DEG_and_species_specific_category <- Not_min_2_expr_Conserved_with_expression_at_least_2_DEG_and_species_specific_category[Not_min_2_expr_Conserved_with_expression_at_least_2_DEG_and_species_specific_category$HOG %in% Hypothesis_6_exp_OGs,]
# length(unique(Hypothesis_6_Not_min_2_expr_Conserved_with_expression_at_least_2_DEG_and_species_specific_category$HOG))
# not 1045
# but 1059


#######################
##### Publication Plot #######

## Plot the regulation categories per species in a UpSetR plot for the hypotheses:

### Hypothesis 1
listInput <- list(Eutrema_down = Hypothesis_1_Not_min_2_expr_Conserved_with_expression_at_least_2_DEG_and_species_specific_category[Hypothesis_1_Not_min_2_expr_Conserved_with_expression_at_least_2_DEG_and_species_specific_category$category_of_regulation_per_species_with_name=="Eutrema_salsugineum down",]$HOG, 
                  Alyrata_down = Hypothesis_1_Not_min_2_expr_Conserved_with_expression_at_least_2_DEG_and_species_specific_category[Hypothesis_1_Not_min_2_expr_Conserved_with_expression_at_least_2_DEG_and_species_specific_category$category_of_regulation_per_species_with_name=="Arabidopsis_lyrata down",]$HOG,
                  Athaliana_down = Hypothesis_1_Not_min_2_expr_Conserved_with_expression_at_least_2_DEG_and_species_specific_category[Hypothesis_1_Not_min_2_expr_Conserved_with_expression_at_least_2_DEG_and_species_specific_category$category_of_regulation_per_species_with_name=="Arabidopsis_thaliana down",]$HOG,
                  Bnapus_down = Hypothesis_1_Not_min_2_expr_Conserved_with_expression_at_least_2_DEG_and_species_specific_category[Hypothesis_1_Not_min_2_expr_Conserved_with_expression_at_least_2_DEG_and_species_specific_category$category_of_regulation_per_species_with_name=="Brassica_napus down",]$HOG,
                  Eutrema_up = Hypothesis_1_Not_min_2_expr_Conserved_with_expression_at_least_2_DEG_and_species_specific_category[Hypothesis_1_Not_min_2_expr_Conserved_with_expression_at_least_2_DEG_and_species_specific_category$category_of_regulation_per_species_with_name=="Eutrema_salsugineum up",]$HOG, 
                  Alyrata_up = Hypothesis_1_Not_min_2_expr_Conserved_with_expression_at_least_2_DEG_and_species_specific_category[Hypothesis_1_Not_min_2_expr_Conserved_with_expression_at_least_2_DEG_and_species_specific_category$category_of_regulation_per_species_with_name=="Arabidopsis_lyrata up",]$HOG,
                  Athaliana_up = Hypothesis_1_Not_min_2_expr_Conserved_with_expression_at_least_2_DEG_and_species_specific_category[Hypothesis_1_Not_min_2_expr_Conserved_with_expression_at_least_2_DEG_and_species_specific_category$category_of_regulation_per_species_with_name=="Arabidopsis_thaliana up",]$HOG,
                  Bnapus_up = Hypothesis_1_Not_min_2_expr_Conserved_with_expression_at_least_2_DEG_and_species_specific_category[Hypothesis_1_Not_min_2_expr_Conserved_with_expression_at_least_2_DEG_and_species_specific_category$category_of_regulation_per_species_with_name=="Brassica_napus up",]$HOG,
                  Eutrema_down_and_up = Hypothesis_1_Not_min_2_expr_Conserved_with_expression_at_least_2_DEG_and_species_specific_category[Hypothesis_1_Not_min_2_expr_Conserved_with_expression_at_least_2_DEG_and_species_specific_category$category_of_regulation_per_species_with_name=="Eutrema_salsugineum down_and_up",]$HOG, 
                  Alyrata_down_and_up = Hypothesis_1_Not_min_2_expr_Conserved_with_expression_at_least_2_DEG_and_species_specific_category[Hypothesis_1_Not_min_2_expr_Conserved_with_expression_at_least_2_DEG_and_species_specific_category$category_of_regulation_per_species_with_name=="Arabidopsis_lyrata down_and_up",]$HOG,
                  Athaliana_down_and_up = Hypothesis_1_Not_min_2_expr_Conserved_with_expression_at_least_2_DEG_and_species_specific_category[Hypothesis_1_Not_min_2_expr_Conserved_with_expression_at_least_2_DEG_and_species_specific_category$category_of_regulation_per_species_with_name=="Arabidopsis_thaliana down_and_up",]$HOG,
                  Bnapus_down_and_up = Hypothesis_1_Not_min_2_expr_Conserved_with_expression_at_least_2_DEG_and_species_specific_category[Hypothesis_1_Not_min_2_expr_Conserved_with_expression_at_least_2_DEG_and_species_specific_category$category_of_regulation_per_species_with_name=="Brassica_napus down_and_up",]$HOG
)

names(listInput) <- c("Eutrema salsugineum down",
                      "Arabidopsis lyrata down",
                      "Arabidopsis thaliana down",
                      "Brassica napus down",
                      "Eutrema salsugineum up",
                      "Arabidopsis lyrata up",
                      "Arabidopsis thaliana up",
                      "Brassica napus up",
                      "Eutrema salsugineum down and up",
                      "Arabidopsis lyrata down and up",
                      "Arabidopsis thaliana down and up",
                      "Brassica napus down and up")

### color the rows per group (up, down or up_and_down)
metadata <-
  data.frame(names(listInput), c(rep("down",4), rep("up", 4), rep("down_and_up", 4)))
names(metadata) = c("sets", "category_of_regulation")


pdf(paste0(OUTDIR, "/Intersection_of_OGs_with_regulation_categories_per_species_ONLY_at_least_2_sig_Hypothesis_1.pdf"), 16, 8, onefile = FALSE)
a <- upset(fromList(listInput), nsets=12, nintersects = 40, order.by = "freq", 
           sets.x.label = "Number of HOGs",
           mainbar.y.label = "Size of Intersection",
           set.metadata = list(data=metadata, plots=list(list(type="matrix_rows", column="category_of_regulation",
                                                              colors=c(down = palette_OkabeIto_black[1:3][1], down_and_up = palette_OkabeIto_black[1:3][2], up = palette_OkabeIto_black[1:3][3]),alpha=0.5))),
                      # and color the intersects per total regulation category
           queries =
             list(
               list(query = intersects, params=list("Arabidopsis lyrata up","Eutrema salsugineum up"), color= palette_OkabeIto_black[1:3][3], active = T),
               list(query = intersects, params=list("Arabidopsis lyrata up","Eutrema salsugineum up", "Arabidopsis thaliana up"), color= palette_OkabeIto_black[1:3][3], active = T),
               list(query = intersects, params=list("Arabidopsis lyrata up","Eutrema salsugineum up", "Arabidopsis thaliana up", "Brassica napus up"), color= palette_OkabeIto_black[1:3][3], active = T),
               list(query = intersects, params=list("Arabidopsis lyrata up","Eutrema salsugineum up", "Brassica napus up"), color= palette_OkabeIto_black[1:3][3], active = T),
               list(query = intersects, params=list("Arabidopsis lyrata down","Eutrema salsugineum down"), color= palette_OkabeIto_black[1:3][1], active = T),
               list(query = intersects, params=list("Arabidopsis lyrata down","Eutrema salsugineum down", "Arabidopsis thaliana down"), color= palette_OkabeIto_black[1:3][1], active = T),
               # not present in data set          list(query = intersects, params=list("Arabidopsis lyrata down","Eutrema salsugineum down", "Arabidopsis thaliana down", "Brassica napus down"), color= "red", active = T)
               list(query = intersects, params=list("Arabidopsis lyrata down","Eutrema salsugineum down", "Brassica napus down"), color= palette_OkabeIto_black[1:3][1], active = T)
#               list(query = intersects, params=list("Arabidopsis lyrata down","Eutrema salsugineum up"), color= "green", active = T),
#               list(query = intersects, params=list("Arabidopsis lyrata up","Eutrema salsugineum down"), color= "green", active = T),
#               list(query = intersects, params=list("Arabidopsis lyrata up","Eutrema salsugineum up", "Brassica napus down"), color= "green", active = T),
#               list(query = intersects, params=list("Arabidopsis lyrata down","Eutrema salsugineum up", "Brassica napus down"), color= "green", active = T),
#               list(query = intersects, params=list("Arabidopsis lyrata up","Eutrema salsugineum down", "Brassica napus up", "Arabidopsis thaliana down"), color= "green", active = T),
#               list(query = intersects, params=list("Arabidopsis lyrata down","Eutrema salsugineum down and up"), color= "green", active = T)
             ),
           matrix.color = "white",
           matrix.dot.alpha=0.7, 
           shade.alpha = 0,
           set_size.show = TRUE,
           set_size.scale_max = 41,
           number.angles = 0,
           point.size = 2, line.size = 1,
           text.scale = c(2,1.5,1.6,1.5,1.5,1.5)
)
a
dev.off()

#######################


### Hypothesis 2
listInput <- list(Eutrema_down = Hypothesis_2_Not_min_2_expr_Conserved_with_expression_at_least_2_DEG_and_species_specific_category[Hypothesis_2_Not_min_2_expr_Conserved_with_expression_at_least_2_DEG_and_species_specific_category$category_of_regulation_per_species_with_name=="Eutrema_salsugineum down",]$HOG, 
                  Alyrata_down = Hypothesis_2_Not_min_2_expr_Conserved_with_expression_at_least_2_DEG_and_species_specific_category[Hypothesis_2_Not_min_2_expr_Conserved_with_expression_at_least_2_DEG_and_species_specific_category$category_of_regulation_per_species_with_name=="Arabidopsis_lyrata down",]$HOG,
                  Athaliana_down = Hypothesis_2_Not_min_2_expr_Conserved_with_expression_at_least_2_DEG_and_species_specific_category[Hypothesis_2_Not_min_2_expr_Conserved_with_expression_at_least_2_DEG_and_species_specific_category$category_of_regulation_per_species_with_name=="Arabidopsis_thaliana down",]$HOG,
                  Bnapus_down = Hypothesis_2_Not_min_2_expr_Conserved_with_expression_at_least_2_DEG_and_species_specific_category[Hypothesis_2_Not_min_2_expr_Conserved_with_expression_at_least_2_DEG_and_species_specific_category$category_of_regulation_per_species_with_name=="Brassica_napus down",]$HOG,
                  Eutrema_up = Hypothesis_2_Not_min_2_expr_Conserved_with_expression_at_least_2_DEG_and_species_specific_category[Hypothesis_2_Not_min_2_expr_Conserved_with_expression_at_least_2_DEG_and_species_specific_category$category_of_regulation_per_species_with_name=="Eutrema_salsugineum up",]$HOG, 
                  Alyrata_up = Hypothesis_2_Not_min_2_expr_Conserved_with_expression_at_least_2_DEG_and_species_specific_category[Hypothesis_2_Not_min_2_expr_Conserved_with_expression_at_least_2_DEG_and_species_specific_category$category_of_regulation_per_species_with_name=="Arabidopsis_lyrata up",]$HOG,
                  Athaliana_up = Hypothesis_2_Not_min_2_expr_Conserved_with_expression_at_least_2_DEG_and_species_specific_category[Hypothesis_2_Not_min_2_expr_Conserved_with_expression_at_least_2_DEG_and_species_specific_category$category_of_regulation_per_species_with_name=="Arabidopsis_thaliana up",]$HOG,
                  Bnapus_up = Hypothesis_2_Not_min_2_expr_Conserved_with_expression_at_least_2_DEG_and_species_specific_category[Hypothesis_2_Not_min_2_expr_Conserved_with_expression_at_least_2_DEG_and_species_specific_category$category_of_regulation_per_species_with_name=="Brassica_napus up",]$HOG,
                  Eutrema_down_and_up = Hypothesis_2_Not_min_2_expr_Conserved_with_expression_at_least_2_DEG_and_species_specific_category[Hypothesis_2_Not_min_2_expr_Conserved_with_expression_at_least_2_DEG_and_species_specific_category$category_of_regulation_per_species_with_name=="Eutrema_salsugineum down_and_up",]$HOG, 
                  Alyrata_down_and_up = Hypothesis_2_Not_min_2_expr_Conserved_with_expression_at_least_2_DEG_and_species_specific_category[Hypothesis_2_Not_min_2_expr_Conserved_with_expression_at_least_2_DEG_and_species_specific_category$category_of_regulation_per_species_with_name=="Arabidopsis_lyrata down_and_up",]$HOG,
                  Athaliana_down_and_up = Hypothesis_2_Not_min_2_expr_Conserved_with_expression_at_least_2_DEG_and_species_specific_category[Hypothesis_2_Not_min_2_expr_Conserved_with_expression_at_least_2_DEG_and_species_specific_category$category_of_regulation_per_species_with_name=="Arabidopsis_thaliana down_and_up",]$HOG,
                  Bnapus_down_and_up = Hypothesis_2_Not_min_2_expr_Conserved_with_expression_at_least_2_DEG_and_species_specific_category[Hypothesis_2_Not_min_2_expr_Conserved_with_expression_at_least_2_DEG_and_species_specific_category$category_of_regulation_per_species_with_name=="Brassica_napus down_and_up",]$HOG
)

names(listInput) <- c("Eutrema salsugineum down",
                      "Arabidopsis lyrata down",
                      "Arabidopsis thaliana down",
                      "Brassica napus down",
                      "Eutrema salsugineum up",
                      "Arabidopsis lyrata up",
                      "Arabidopsis thaliana up",
                      "Brassica napus up",
                      "Eutrema salsugineum down and up",
                      "Arabidopsis lyrata down and up",
                      "Arabidopsis thaliana down and up",
                      "Brassica napus down and up")

### color the rows per group (up, down or up_and_down)
metadata <-
  data.frame(names(listInput), c(rep("down",4), rep("up", 4), rep("down_and_up", 4)))
names(metadata) = c("sets", "category_of_regulation")

pdf(paste0(OUTDIR, "/Intersection_of_OGs_with_regulation_categories_per_species_ONLY_at_least_2_sig_Hypothesis_2.pdf"), 16, 8, onefile = FALSE)
a <- upset(fromList(listInput), nsets=12, nintersects = 40, order.by = "freq", 
           sets.x.label = "Number of HOGs per \nspecies-specific regulation category",
           mainbar.y.label = "Size of Intersection",
           set.metadata = list(data=metadata, plots=list(list(type="matrix_rows", column="category_of_regulation",
                                                              colors=c(down = palette_OkabeIto_black[1:3][1], down_and_up = palette_OkabeIto_black[1:3][2], up = palette_OkabeIto_black[1:3][3]),alpha=0.5))),
           matrix.color = "white",
           matrix.dot.alpha=0.7, 
           shade.alpha = 0,
           set_size.show = TRUE,
           set_size.scale_max = 170,
           number.angles = 0,
           
           point.size = 2, line.size = 1,
           text.scale = c(2,1.5,1.6,1.5,1.5,1.5)
)
a
dev.off()


### Hypothesis 3
listInput <- list(Eutrema_down = Hypothesis_3_Not_min_2_expr_Conserved_with_expression_at_least_2_DEG_and_species_specific_category[Hypothesis_3_Not_min_2_expr_Conserved_with_expression_at_least_2_DEG_and_species_specific_category$category_of_regulation_per_species_with_name=="Eutrema_salsugineum down",]$HOG, 
                  Alyrata_down = Hypothesis_3_Not_min_2_expr_Conserved_with_expression_at_least_2_DEG_and_species_specific_category[Hypothesis_3_Not_min_2_expr_Conserved_with_expression_at_least_2_DEG_and_species_specific_category$category_of_regulation_per_species_with_name=="Arabidopsis_lyrata down",]$HOG,
                  Athaliana_down = Hypothesis_3_Not_min_2_expr_Conserved_with_expression_at_least_2_DEG_and_species_specific_category[Hypothesis_3_Not_min_2_expr_Conserved_with_expression_at_least_2_DEG_and_species_specific_category$category_of_regulation_per_species_with_name=="Arabidopsis_thaliana down",]$HOG,
                  Bnapus_down = Hypothesis_3_Not_min_2_expr_Conserved_with_expression_at_least_2_DEG_and_species_specific_category[Hypothesis_3_Not_min_2_expr_Conserved_with_expression_at_least_2_DEG_and_species_specific_category$category_of_regulation_per_species_with_name=="Brassica_napus down",]$HOG,
                  Eutrema_up = Hypothesis_3_Not_min_2_expr_Conserved_with_expression_at_least_2_DEG_and_species_specific_category[Hypothesis_3_Not_min_2_expr_Conserved_with_expression_at_least_2_DEG_and_species_specific_category$category_of_regulation_per_species_with_name=="Eutrema_salsugineum up",]$HOG, 
                  Alyrata_up = Hypothesis_3_Not_min_2_expr_Conserved_with_expression_at_least_2_DEG_and_species_specific_category[Hypothesis_3_Not_min_2_expr_Conserved_with_expression_at_least_2_DEG_and_species_specific_category$category_of_regulation_per_species_with_name=="Arabidopsis_lyrata up",]$HOG,
                  Athaliana_up = Hypothesis_3_Not_min_2_expr_Conserved_with_expression_at_least_2_DEG_and_species_specific_category[Hypothesis_3_Not_min_2_expr_Conserved_with_expression_at_least_2_DEG_and_species_specific_category$category_of_regulation_per_species_with_name=="Arabidopsis_thaliana up",]$HOG,
                  Bnapus_up = Hypothesis_3_Not_min_2_expr_Conserved_with_expression_at_least_2_DEG_and_species_specific_category[Hypothesis_3_Not_min_2_expr_Conserved_with_expression_at_least_2_DEG_and_species_specific_category$category_of_regulation_per_species_with_name=="Brassica_napus up",]$HOG,
                  Eutrema_down_and_up = Hypothesis_3_Not_min_2_expr_Conserved_with_expression_at_least_2_DEG_and_species_specific_category[Hypothesis_3_Not_min_2_expr_Conserved_with_expression_at_least_2_DEG_and_species_specific_category$category_of_regulation_per_species_with_name=="Eutrema_salsugineum down_and_up",]$HOG, 
                  Alyrata_down_and_up = Hypothesis_3_Not_min_2_expr_Conserved_with_expression_at_least_2_DEG_and_species_specific_category[Hypothesis_3_Not_min_2_expr_Conserved_with_expression_at_least_2_DEG_and_species_specific_category$category_of_regulation_per_species_with_name=="Arabidopsis_lyrata down_and_up",]$HOG,
                  Athaliana_down_and_up = Hypothesis_3_Not_min_2_expr_Conserved_with_expression_at_least_2_DEG_and_species_specific_category[Hypothesis_3_Not_min_2_expr_Conserved_with_expression_at_least_2_DEG_and_species_specific_category$category_of_regulation_per_species_with_name=="Arabidopsis_thaliana down_and_up",]$HOG,
                  Bnapus_down_and_up = Hypothesis_3_Not_min_2_expr_Conserved_with_expression_at_least_2_DEG_and_species_specific_category[Hypothesis_3_Not_min_2_expr_Conserved_with_expression_at_least_2_DEG_and_species_specific_category$category_of_regulation_per_species_with_name=="Brassica_napus down_and_up",]$HOG
)

names(listInput) <- c("Eutrema salsugineum down",
                      "Arabidopsis lyrata down",
                      "Arabidopsis thaliana down",
                      "Brassica napus down",
                      "Eutrema salsugineum up",
                      "Arabidopsis lyrata up",
                      "Arabidopsis thaliana up",
                      "Brassica napus up",
                      "Eutrema salsugineum down and up",
                      "Arabidopsis lyrata down and up",
                      "Arabidopsis thaliana down and up",
                      "Brassica napus down and up")

### color the rows per group (up, down or up_and_down)
metadata <-
  data.frame(names(listInput), c(rep("down",4), rep("up", 4), rep("down_and_up", 4)))
names(metadata) = c("sets", "category_of_regulation")

pdf(paste0(OUTDIR, "/Intersection_of_OGs_with_regulation_categories_per_species_ONLY_at_least_2_sig_Hypothesis_3.pdf"), 16, 8, onefile = FALSE)
a <- upset(fromList(listInput), nsets=12, nintersects = 40, order.by = "freq", 
           sets.x.label = "Number of HOGs per \nspecies-specific regulation category",
           mainbar.y.label = "Size of Intersection",
           set.metadata = list(data=metadata, plots=list(list(type="matrix_rows", column="category_of_regulation",
                                                              colors=c(down = palette_OkabeIto_black[1:3][1], down_and_up = palette_OkabeIto_black[1:3][2], up = palette_OkabeIto_black[1:3][3]),alpha=0.5))),
           matrix.color = "white",
           matrix.dot.alpha=0.7, 
           shade.alpha = 0,
           set_size.show = TRUE,
           set_size.scale_max = 270,
           number.angles = 0,
           
           point.size = 2, line.size = 1,
           text.scale = c(2,1.5,1.6,1.5,1.5,1.5)
)
a
dev.off()


### Hypothesis 4
listInput <- list(Eutrema_down = Hypothesis_4_Not_min_2_expr_Conserved_with_expression_at_least_2_DEG_and_species_specific_category[Hypothesis_4_Not_min_2_expr_Conserved_with_expression_at_least_2_DEG_and_species_specific_category$category_of_regulation_per_species_with_name=="Eutrema_salsugineum down",]$HOG, 
                  Alyrata_down = Hypothesis_4_Not_min_2_expr_Conserved_with_expression_at_least_2_DEG_and_species_specific_category[Hypothesis_4_Not_min_2_expr_Conserved_with_expression_at_least_2_DEG_and_species_specific_category$category_of_regulation_per_species_with_name=="Arabidopsis_lyrata down",]$HOG,
                  Athaliana_down = Hypothesis_4_Not_min_2_expr_Conserved_with_expression_at_least_2_DEG_and_species_specific_category[Hypothesis_4_Not_min_2_expr_Conserved_with_expression_at_least_2_DEG_and_species_specific_category$category_of_regulation_per_species_with_name=="Arabidopsis_thaliana down",]$HOG,
                  Bnapus_down = Hypothesis_4_Not_min_2_expr_Conserved_with_expression_at_least_2_DEG_and_species_specific_category[Hypothesis_4_Not_min_2_expr_Conserved_with_expression_at_least_2_DEG_and_species_specific_category$category_of_regulation_per_species_with_name=="Brassica_napus down",]$HOG,
                  Eutrema_up = Hypothesis_4_Not_min_2_expr_Conserved_with_expression_at_least_2_DEG_and_species_specific_category[Hypothesis_4_Not_min_2_expr_Conserved_with_expression_at_least_2_DEG_and_species_specific_category$category_of_regulation_per_species_with_name=="Eutrema_salsugineum up",]$HOG, 
                  Alyrata_up = Hypothesis_4_Not_min_2_expr_Conserved_with_expression_at_least_2_DEG_and_species_specific_category[Hypothesis_4_Not_min_2_expr_Conserved_with_expression_at_least_2_DEG_and_species_specific_category$category_of_regulation_per_species_with_name=="Arabidopsis_lyrata up",]$HOG,
                  Athaliana_up = Hypothesis_4_Not_min_2_expr_Conserved_with_expression_at_least_2_DEG_and_species_specific_category[Hypothesis_4_Not_min_2_expr_Conserved_with_expression_at_least_2_DEG_and_species_specific_category$category_of_regulation_per_species_with_name=="Arabidopsis_thaliana up",]$HOG,
                  Bnapus_up = Hypothesis_4_Not_min_2_expr_Conserved_with_expression_at_least_2_DEG_and_species_specific_category[Hypothesis_4_Not_min_2_expr_Conserved_with_expression_at_least_2_DEG_and_species_specific_category$category_of_regulation_per_species_with_name=="Brassica_napus up",]$HOG,
                  Eutrema_down_and_up = Hypothesis_4_Not_min_2_expr_Conserved_with_expression_at_least_2_DEG_and_species_specific_category[Hypothesis_4_Not_min_2_expr_Conserved_with_expression_at_least_2_DEG_and_species_specific_category$category_of_regulation_per_species_with_name=="Eutrema_salsugineum down_and_up",]$HOG, 
                  Alyrata_down_and_up = Hypothesis_4_Not_min_2_expr_Conserved_with_expression_at_least_2_DEG_and_species_specific_category[Hypothesis_4_Not_min_2_expr_Conserved_with_expression_at_least_2_DEG_and_species_specific_category$category_of_regulation_per_species_with_name=="Arabidopsis_lyrata down_and_up",]$HOG,
                  Athaliana_down_and_up = Hypothesis_4_Not_min_2_expr_Conserved_with_expression_at_least_2_DEG_and_species_specific_category[Hypothesis_4_Not_min_2_expr_Conserved_with_expression_at_least_2_DEG_and_species_specific_category$category_of_regulation_per_species_with_name=="Arabidopsis_thaliana down_and_up",]$HOG,
                  Bnapus_down_and_up = Hypothesis_4_Not_min_2_expr_Conserved_with_expression_at_least_2_DEG_and_species_specific_category[Hypothesis_4_Not_min_2_expr_Conserved_with_expression_at_least_2_DEG_and_species_specific_category$category_of_regulation_per_species_with_name=="Brassica_napus down_and_up",]$HOG
)

names(listInput) <- c("Eutrema salsugineum down",
                      "Arabidopsis lyrata down",
                      "Arabidopsis thaliana down",
                      "Brassica napus down",
                      "Eutrema salsugineum up",
                      "Arabidopsis lyrata up",
                      "Arabidopsis thaliana up",
                      "Brassica napus up",
                      "Eutrema salsugineum down and up",
                      "Arabidopsis lyrata down and up",
                      "Arabidopsis thaliana down and up",
                      "Brassica napus down and up")

### color the rows per group (up, down or up_and_down)
metadata <-
  data.frame(names(listInput), c(rep("down",4), rep("up", 4), rep("down_and_up", 4)))
names(metadata) = c("sets", "category_of_regulation")

pdf(paste0(OUTDIR, "/Intersection_of_OGs_with_regulation_categories_per_species_ONLY_at_least_2_sig_Hypothesis_4.pdf"), 16, 8, onefile = FALSE)
a <- upset(fromList(listInput), nsets=12, nintersects = 40, order.by = "freq", 
           sets.x.label = "Number of HOGs per \nspecies-specific regulation category",
           mainbar.y.label = "Size of Intersection",
           set.metadata = list(data=metadata, plots=list(list(type="matrix_rows", column="category_of_regulation",
                                                              colors=c(down = palette_OkabeIto_black[1:3][1], down_and_up = palette_OkabeIto_black[1:3][2], up = palette_OkabeIto_black[1:3][3]),alpha=0.5))),
           matrix.color = "white",
           matrix.dot.alpha=0.7, 
           shade.alpha = 0,
           set_size.show = TRUE,
           set_size.scale_max = 13,
           number.angles = 0,
           
           point.size = 2, line.size = 1,
           text.scale = c(2,1.5,1.6,1.5,1.5,1.5)
)
a
dev.off()


### Hypothesis 5
listInput <- list(Eutrema_down = Hypothesis_5_Not_min_2_expr_Conserved_with_expression_at_least_2_DEG_and_species_specific_category[Hypothesis_5_Not_min_2_expr_Conserved_with_expression_at_least_2_DEG_and_species_specific_category$category_of_regulation_per_species_with_name=="Eutrema_salsugineum down",]$HOG, 
                  Alyrata_down = Hypothesis_5_Not_min_2_expr_Conserved_with_expression_at_least_2_DEG_and_species_specific_category[Hypothesis_5_Not_min_2_expr_Conserved_with_expression_at_least_2_DEG_and_species_specific_category$category_of_regulation_per_species_with_name=="Arabidopsis_lyrata down",]$HOG,
                  Athaliana_down = Hypothesis_5_Not_min_2_expr_Conserved_with_expression_at_least_2_DEG_and_species_specific_category[Hypothesis_5_Not_min_2_expr_Conserved_with_expression_at_least_2_DEG_and_species_specific_category$category_of_regulation_per_species_with_name=="Arabidopsis_thaliana down",]$HOG,
                  Bnapus_down = Hypothesis_5_Not_min_2_expr_Conserved_with_expression_at_least_2_DEG_and_species_specific_category[Hypothesis_5_Not_min_2_expr_Conserved_with_expression_at_least_2_DEG_and_species_specific_category$category_of_regulation_per_species_with_name=="Brassica_napus down",]$HOG,
                  Eutrema_up = Hypothesis_5_Not_min_2_expr_Conserved_with_expression_at_least_2_DEG_and_species_specific_category[Hypothesis_5_Not_min_2_expr_Conserved_with_expression_at_least_2_DEG_and_species_specific_category$category_of_regulation_per_species_with_name=="Eutrema_salsugineum up",]$HOG, 
                  Alyrata_up = Hypothesis_5_Not_min_2_expr_Conserved_with_expression_at_least_2_DEG_and_species_specific_category[Hypothesis_5_Not_min_2_expr_Conserved_with_expression_at_least_2_DEG_and_species_specific_category$category_of_regulation_per_species_with_name=="Arabidopsis_lyrata up",]$HOG,
                  Athaliana_up = Hypothesis_5_Not_min_2_expr_Conserved_with_expression_at_least_2_DEG_and_species_specific_category[Hypothesis_5_Not_min_2_expr_Conserved_with_expression_at_least_2_DEG_and_species_specific_category$category_of_regulation_per_species_with_name=="Arabidopsis_thaliana up",]$HOG,
                  Bnapus_up = Hypothesis_5_Not_min_2_expr_Conserved_with_expression_at_least_2_DEG_and_species_specific_category[Hypothesis_5_Not_min_2_expr_Conserved_with_expression_at_least_2_DEG_and_species_specific_category$category_of_regulation_per_species_with_name=="Brassica_napus up",]$HOG,
                  Eutrema_down_and_up = Hypothesis_5_Not_min_2_expr_Conserved_with_expression_at_least_2_DEG_and_species_specific_category[Hypothesis_5_Not_min_2_expr_Conserved_with_expression_at_least_2_DEG_and_species_specific_category$category_of_regulation_per_species_with_name=="Eutrema_salsugineum down_and_up",]$HOG, 
                  Alyrata_down_and_up = Hypothesis_5_Not_min_2_expr_Conserved_with_expression_at_least_2_DEG_and_species_specific_category[Hypothesis_5_Not_min_2_expr_Conserved_with_expression_at_least_2_DEG_and_species_specific_category$category_of_regulation_per_species_with_name=="Arabidopsis_lyrata down_and_up",]$HOG,
                  Athaliana_down_and_up = Hypothesis_5_Not_min_2_expr_Conserved_with_expression_at_least_2_DEG_and_species_specific_category[Hypothesis_5_Not_min_2_expr_Conserved_with_expression_at_least_2_DEG_and_species_specific_category$category_of_regulation_per_species_with_name=="Arabidopsis_thaliana down_and_up",]$HOG,
                  Bnapus_down_and_up = Hypothesis_5_Not_min_2_expr_Conserved_with_expression_at_least_2_DEG_and_species_specific_category[Hypothesis_5_Not_min_2_expr_Conserved_with_expression_at_least_2_DEG_and_species_specific_category$category_of_regulation_per_species_with_name=="Brassica_napus down_and_up",]$HOG
)

names(listInput) <- c("Eutrema salsugineum down",
                      "Arabidopsis lyrata down",
                      "Arabidopsis thaliana down",
                      "Brassica napus down",
                      "Eutrema salsugineum up",
                      "Arabidopsis lyrata up",
                      "Arabidopsis thaliana up",
                      "Brassica napus up",
                      "Eutrema salsugineum down and up",
                      "Arabidopsis lyrata down and up",
                      "Arabidopsis thaliana down and up",
                      "Brassica napus down and up")

### color the rows per group (up, down or up_and_down)
metadata <-
  data.frame(names(listInput), c(rep("down",4), rep("up", 4), rep("down_and_up", 4)))
names(metadata) = c("sets", "category_of_regulation")

pdf(paste0(OUTDIR, "/Intersection_of_OGs_with_regulation_categories_per_species_ONLY_at_least_2_sig_Hypothesis_5.pdf"), 16, 8, onefile = FALSE)
a <- upset(fromList(listInput), nsets=12, nintersects = 40, order.by = "freq", 
           sets.x.label = "Number of HOGs per \nspecies-specific regulation category",
           mainbar.y.label = "Size of Intersection",
           set.metadata = list(data=metadata, plots=list(list(type="matrix_rows", column="category_of_regulation",
                                                              colors=c(down = palette_OkabeIto_black[1:3][1], down_and_up = palette_OkabeIto_black[1:3][2], up = palette_OkabeIto_black[1:3][3]),alpha=0.5))),
           matrix.color = "white",
           matrix.dot.alpha=0.7, 
           shade.alpha = 0,
           set_size.show = TRUE,
           set_size.scale_max = 77,
           number.angles = 0,
           
           point.size = 2, line.size = 1,
           text.scale = c(2,1.5,1.6,1.5,1.5,1.5)
)
a
dev.off()

### Hypothesis 6
listInput <- list(Eutrema_down = Hypothesis_6_Not_min_2_expr_Conserved_with_expression_at_least_2_DEG_and_species_specific_category[Hypothesis_6_Not_min_2_expr_Conserved_with_expression_at_least_2_DEG_and_species_specific_category$category_of_regulation_per_species_with_name=="Eutrema_salsugineum down",]$HOG, 
                  Alyrata_down = Hypothesis_6_Not_min_2_expr_Conserved_with_expression_at_least_2_DEG_and_species_specific_category[Hypothesis_6_Not_min_2_expr_Conserved_with_expression_at_least_2_DEG_and_species_specific_category$category_of_regulation_per_species_with_name=="Arabidopsis_lyrata down",]$HOG,
                  Athaliana_down = Hypothesis_6_Not_min_2_expr_Conserved_with_expression_at_least_2_DEG_and_species_specific_category[Hypothesis_6_Not_min_2_expr_Conserved_with_expression_at_least_2_DEG_and_species_specific_category$category_of_regulation_per_species_with_name=="Arabidopsis_thaliana down",]$HOG,
                  Bnapus_down = Hypothesis_6_Not_min_2_expr_Conserved_with_expression_at_least_2_DEG_and_species_specific_category[Hypothesis_6_Not_min_2_expr_Conserved_with_expression_at_least_2_DEG_and_species_specific_category$category_of_regulation_per_species_with_name=="Brassica_napus down",]$HOG,
                  Eutrema_up = Hypothesis_6_Not_min_2_expr_Conserved_with_expression_at_least_2_DEG_and_species_specific_category[Hypothesis_6_Not_min_2_expr_Conserved_with_expression_at_least_2_DEG_and_species_specific_category$category_of_regulation_per_species_with_name=="Eutrema_salsugineum up",]$HOG, 
                  Alyrata_up = Hypothesis_6_Not_min_2_expr_Conserved_with_expression_at_least_2_DEG_and_species_specific_category[Hypothesis_6_Not_min_2_expr_Conserved_with_expression_at_least_2_DEG_and_species_specific_category$category_of_regulation_per_species_with_name=="Arabidopsis_lyrata up",]$HOG,
                  Athaliana_up = Hypothesis_6_Not_min_2_expr_Conserved_with_expression_at_least_2_DEG_and_species_specific_category[Hypothesis_6_Not_min_2_expr_Conserved_with_expression_at_least_2_DEG_and_species_specific_category$category_of_regulation_per_species_with_name=="Arabidopsis_thaliana up",]$HOG,
                  Bnapus_up = Hypothesis_6_Not_min_2_expr_Conserved_with_expression_at_least_2_DEG_and_species_specific_category[Hypothesis_6_Not_min_2_expr_Conserved_with_expression_at_least_2_DEG_and_species_specific_category$category_of_regulation_per_species_with_name=="Brassica_napus up",]$HOG,
                  Eutrema_down_and_up = Hypothesis_6_Not_min_2_expr_Conserved_with_expression_at_least_2_DEG_and_species_specific_category[Hypothesis_6_Not_min_2_expr_Conserved_with_expression_at_least_2_DEG_and_species_specific_category$category_of_regulation_per_species_with_name=="Eutrema_salsugineum down_and_up",]$HOG, 
                  Alyrata_down_and_up = Hypothesis_6_Not_min_2_expr_Conserved_with_expression_at_least_2_DEG_and_species_specific_category[Hypothesis_6_Not_min_2_expr_Conserved_with_expression_at_least_2_DEG_and_species_specific_category$category_of_regulation_per_species_with_name=="Arabidopsis_lyrata down_and_up",]$HOG,
                  Athaliana_down_and_up = Hypothesis_6_Not_min_2_expr_Conserved_with_expression_at_least_2_DEG_and_species_specific_category[Hypothesis_6_Not_min_2_expr_Conserved_with_expression_at_least_2_DEG_and_species_specific_category$category_of_regulation_per_species_with_name=="Arabidopsis_thaliana down_and_up",]$HOG,
                  Bnapus_down_and_up = Hypothesis_6_Not_min_2_expr_Conserved_with_expression_at_least_2_DEG_and_species_specific_category[Hypothesis_6_Not_min_2_expr_Conserved_with_expression_at_least_2_DEG_and_species_specific_category$category_of_regulation_per_species_with_name=="Brassica_napus down_and_up",]$HOG
)

names(listInput) <- c("Eutrema salsugineum down",
                      "Arabidopsis lyrata down",
                      "Arabidopsis thaliana down",
                      "Brassica napus down",
                      "Eutrema salsugineum up",
                      "Arabidopsis lyrata up",
                      "Arabidopsis thaliana up",
                      "Brassica napus up",
                      "Eutrema salsugineum down and up",
                      "Arabidopsis lyrata down and up",
                      "Arabidopsis thaliana down and up",
                      "Brassica napus down and up")

### color the rows per group (up, down or up_and_down)
metadata <-
  data.frame(names(listInput), c(rep("down",4), rep("up", 4), rep("down_and_up", 4)))
names(metadata) = c("sets", "category_of_regulation")

pdf(paste0(OUTDIR, "/Intersection_of_OGs_with_regulation_categories_per_species_ONLY_at_least_2_sig_Hypothesis_6.pdf"), 16, 8, onefile = FALSE)
a <- upset(fromList(listInput), nsets=12, nintersects = 40, order.by = "freq", 
           sets.x.label = "Number of HOGs per \nspecies-specific regulation category",
           mainbar.y.label = "Size of Intersection",
           set.metadata = list(data=metadata, plots=list(list(type="matrix_rows", column="category_of_regulation",
                                                              colors=c(down = palette_OkabeIto_black[1:3][1], down_and_up = palette_OkabeIto_black[1:3][2], up = palette_OkabeIto_black[1:3][3]),alpha=0.5))),
           matrix.color = "white",
           matrix.dot.alpha=0.7, 
           shade.alpha = 0,
           set_size.show = TRUE,
           set_size.scale_max = 448,
           number.angles = 0,
           
           point.size = 2, line.size = 1,
           text.scale = c(2,1.5,1.6,1.5,1.5,1.5)
)
a
dev.off()
```
