## Additional File 3 for "Integration of differential expression under drought with gene family expansion unique to drought tolerant species predicts candidate genes for drought adaptation in Brassicaceae species": filter_min100aa.sh

```
#!/bin/bash

#filter_min100aa.sh

WDIR=/data/home/uebermuth/fdi_genofam_mount/uebermuth/brassicaceae-drought/Genomic_Data_BR-dr-02
OUTDIR=/data/home/uebermuth/fdi_genofam_mount/uebermuth/brassicaceae-drought/Genomic_Data_BR-dr-03/min100aa
cd $WDIR
normal=$(ls *.fa)

for f in $normal; do
        var=${f%.fa}
#linearize fasta:
	cat $f |\
	awk '/^>/ {if(N>0) printf("\n"); printf("%s\t",$0);N++;next;} {printf("%s",$0);} END {if(N>0) printf("\n");}' |\
#extract lenght:
	awk -F '\t'  '{printf("%s\t%d\n",$0,length($2));}' |\
#keep only if length>99:
	awk -F '\t' '{if($3>99) printf("%s\t%s\n",$1, $2);}' |\
#back to fasta:
#	tr "\t" "\n" |\
#	fold -w 100 > ${OUTDIR}/${var}_min100aa.fa
	tr "\t" "\n" > ${OUTDIR}/${var}_min100aa.fa
done;
```
