## Additional File 3 for "Integration of differential expression under drought with gene family expansion unique to drought tolerant species predicts candidate genes for drought adaptation in Brassicaceae species": merge_all_info_of_candidate_gene_families.R

```
# for publication purposes, I merged all info I have gathered while performing all analyses which I have saved in different tables
# I merge the data per gene, clustered per HOG and save them into two main files: 
# one for the HOGs which are expanded in Eutrema salsginuem and one for the HOGs which are expanded in Arabidopsis lyrata
# And I saved two subsets of these: "expanded and DE in both tolerant" and "expanded in both and DE in any of the tolerant"

setwd("/home/cropbio/sciebo/Documents/PhD/Brassicaceae-drought/BR-dr-03/Analysis/")
load("/home/cropbio/sciebo/Documents/PhD/Brassicaceae-drought/BR-dr-03/A2TEA_finished.RData")
Conserved_Selection_results_genes_pvalues <- read.table("/home/cropbio/sciebo/Documents/PhD/Brassicaceae-drought/BR-dr-03/Selection/Analysis/results_20221027_Conserved_Dataset/OGs_pos_selection_pvalues.tsv", sep = "\t", header=F)
names(Conserved_Selection_results_genes_pvalues) <- c("HOG", "gene", "uncorrected_pvalue", "selection_pvalue")

Alyrata <- read.csv("HOGs_Alyrata_exp_DEG_Alyrata_with_HRD_and_expression.tsv", sep = "\t", header=T)
Eutrema <-read.csv("HOGs_Eutrema_exp_DEG_Eutrema_with_HRD_and_expression.tsv", sep = "\t", header=T)


merged_1 <- merge(Alyrata, Eutrema, all=T)
merged_2 <- merged_1[order(merged_1$HOG),]
# make a column "gene_separator_edited" which contains the gene IDs from Alyrata which contain "." replaced by the gene IDs from Alyrata which contain "_"
merged_2$Protein.Accession_separator_edited <- gsub(".", "_", merged_2$Protein.Accession, fixed = T)
length(unique(merged_2$HOG.x))
# 928

# info tables with hand made notes on the up regulated genes from Alyrata and Eutrema of the 17 HOGs in the focus set (48 HOGs)
Both_tolerant_exp_and_only_up_notes <- read.csv("/home/cropbio/sciebo/Documents/PhD/Brassicaceae-drought/BR-dr-03/Candidate_Gene_OGs/Exp_and_upregulated_only_tolerant/Conserved_with_expression_with_species_specific_category_only_both_tol_up_and_exp_both_tol_NOTES.csv", header=T)

merged_3 <- merge(merged_2, Both_tolerant_exp_and_only_up_notes[,c("gene", "Notes", "Priority..1.3.", "Function_Notes")], 
                  by.x="Protein.Accession", by.y="gene", all=T)
merged_4 <- merged_3[order(merged_3$HOG.x),]
length(unique(merged_4$HOG.x))
# 928
Eutrema_DEG_and_Pos_sel_notes <- read.csv("/home/cropbio/sciebo/Documents/PhD/Brassicaceae-drought/BR-dr-03/Selection/Analysis/Selection_and_pos_sel_Eutrema_OGs/Conserved_Selection_results_genes_pvalues_HOG_DE_Es_pos_and_DE_and_exp_notes.csv", header=T)
# contains info on 41 genes

merged_5 <- merge(merged_4, Eutrema_DEG_and_Pos_sel_notes[,c("gene", "Priority..0.3.", "Notes", "Function_Notes", "alignment.length",
                  "Subfunctionalization")], by.x="Protein.Accession_separator_edited", 
                  by.y="gene", all=T)
merged_6 <- merged_5[order(merged_5$HOG.x),]
names(merged_6) <- c(names(merged_6[c(1,2)]), "HOG", names(merged_6[4:10]), "Notes_focus_set", "Prio_focus_set", 
                     "Notes_on_function_focus_set", "Prio_Selection_Eutrema", "Notes_Selection_Eutrema", 
                     "Notes_on_function_Selection_Eutrema", "alignment_length_Selection_Eutrema", "Subfunctionalization_Selection_Eutrema")
length(unique(merged_6$HOG))
# 928

# note that we have the ID with "_" instead of "." in some gene ID from Alyrata in the following table. 
# Merge via Protein.Accession_separator_edited in merged_2
Alyrata_DEG_and_Pos_sel_notes <- read.csv("/home/cropbio/sciebo/Documents/PhD/Brassicaceae-drought/BR-dr-03/Selection/Analysis/Selection_and_pos_sel_Alyrata_OGs/20240430_Conserved_Selection_results_genes_pvalues_HOG_DE_Al_exp_and_DEG_is_pos_Notes.csv", header=T)
# contains info on 47 genes
merged_7 <- base::merge(merged_6, Alyrata_DEG_and_Pos_sel_notes[,c("gene", "Priority..0.3.", "Notes", 
                        "Function_Notes", "Subfunctionalization")], by.x="Protein.Accession_separator_edited", 
                        by.y="gene", all=T)
merged_8 <- merged_7[order(merged_7$HOG),]
names(merged_8) <- c(names(merged_8[1:18]),"Prio_Selection_Alyrata", "Notes_Selection_Alyrata", 
                     "Notes_on_function_Selection_Alyrata", "Subfunctionalization_Selection_Alyrata")
length(unique(merged_8$HOG))
# 928

# merge the Notes_Selection per species to one Notes_Selection column
# merge the Prio_Selection per tolerant species to one Prio_Selection column
library(tidyr)
library(stringr)
library(dplyr)
# convert missing value to "" in order to later merge the columns "Prio_Selection_Eutrema" and "Prio_Selection_Alyrata","Notes_Selection_Eutrema" and "Notes_Selection_Alyrata",
# "Notes_on_function_Selection_Eutrema" and "Notes_on_function_Selection_Alyrata", "Subfunctionalization_Selection_Eutrema" and "Subfunctionalization_Selection_Alyrata"

merged_9 <-
merged_8 %>%
  mutate(
    Prio_Selection_Eutrema_2 = str_replace_na(Prio_Selection_Eutrema, replacement=""),
    Prio_Selection_Alyrata_2 = str_replace_na(Prio_Selection_Alyrata, replacement=""),
    Notes_Selection_Eutrema_2 = str_replace_na(Notes_Selection_Eutrema, replacement=""),
    Notes_Selection_Alyrata_2 = str_replace_na(Notes_Selection_Alyrata, replacement=""),
    Subfunctionalization_Selection_Eutrema_2 = str_replace_na(Subfunctionalization_Selection_Eutrema, replacement=""),
    Subfunctionalization_Selection_Alyrata_2 = str_replace_na(Subfunctionalization_Selection_Alyrata,replacement=""),
    Notes_on_function_Selection_Eutrema_2 = str_replace_na(Notes_on_function_Selection_Eutrema, replacement=""),
    Notes_on_function_Selection_Alyrata_2 = str_replace_na(Notes_on_function_Selection_Alyrata, replacement="")
  ) %>%
  mutate(
  Prio_Selection = str_c(Prio_Selection_Eutrema_2, "", Prio_Selection_Alyrata_2),
  Notes_Selection = str_c(Notes_Selection_Eutrema_2, "", Notes_Selection_Alyrata_2),
  Subfunctionalization = str_c(Subfunctionalization_Selection_Eutrema_2, "", Subfunctionalization_Selection_Alyrata_2),
  Notes_on_function_Selection = str_c(Notes_on_function_Selection_Eutrema_2, "", Notes_on_function_Selection_Alyrata_2)
  )

length(unique(merged_9$HOG))
# 928

# extract the selection padj values from "Conserved_Selection_results"
merged_10 <-
  merge(merged_9, Conserved_Selection_results_genes_pvalues[,c("gene", "selection_pvalue")], by.x="Protein.Accession_separator_edited",
        by.y="gene", all.x=T, all.y=F) # all.x and all.y because we do not want all HOGs which were evaluated for positive selection
# but only the ones which are also in the sets of expanded and DE HOGs either in Aly or in Esa
length(unique(merged_10$HOG))
# 928

# what HOG is extra when adding all=T instead of all.x=T and all.y=F?
#merged_10_test <-
#  merge(merged_9, Conserved_Selection_results_genes_pvalues[,c("gene", "selection_pvalue")], by.x="Protein.Accession_separator_edited",
#        by.y="gene", all=T)
#setdiff(merged_10_test$HOG, merged_10$HOG)
# NAs! probably genestested for pos selection which are not in either of Aly exp and DE or Esa exp and DE


# I forgot to put the parameter all=T to not loose rows which do not find a match (the ones which were not tested for pos selection?)

# keep only the relevant columns, rename them if necessary and order by HOG, species, log2FC (decreasing)
merged_11 <- merged_10[with(merged_10, order(HOG, species, -log2FoldChange)), c(1:10, 35, 33, 31, 12, 11, 32, 13, 34)]
# replace NA values by "" also in "Prio_focus_set", "Notes_focus_set" and "Notes_on_function_focus_set" to be consistent in all character string variables
merged_11$Prio_focus_set <- str_replace_na(merged_11$Prio_focus_set, replacement = "")
merged_11$Notes_focus_set <- str_replace_na(merged_11$Notes_focus_set, replacement = "")
merged_11$Notes_on_function_focus_set <- str_replace_na(merged_11$Notes_on_function_focus_set, replacement = "")

length(unique(merged_11$HOG))
# 928

# add a variable which marks the HOGs in the Candidate Gene Set
candidate_gene_set <- read.table("HOG_level_list_all_tol_exp_DEG_any_tol.tsv", header=T)
merged_12 <-
  merged_11 %>%
  mutate(Candidate_Gene_Set = case_when(HOG %in% candidate_gene_set$HOG ~ "candidate_gene_set",
                                      TRUE ~ "no"))
# and add a variable which marks the HOGs in the focus set of Candidate gene families which are up regulated only in both tolerant species
HOG_level_list_all_tol_exp_DEG_both_tol <- read.table("HOG_level_list_all_tol_exp_DEG_both_tol.tsv", header=T)
merged_13 <-
  merged_12 %>%
mutate(Only_both_tol_up = case_when(HOG %in% HOG_level_list_all_tol_exp_DEG_both_tol$HOG ~ "only_both_tol_up",
                                      TRUE ~ "no"))

# export the table with all genes within the sets "expanded and DE in Eutrema" and "expanded and DE in Alyrata"
# and the relevant notes and info on the subsets of these sets which contain either DEGs from Eutrema or Alyrata 
# reagrding diversifying selection or whether they are in the Candidate gene set or in its subset of gene families uniquely up regulated in Eutrema AND Alyrata
write.csv(merged_13[,c(1:11,19:20,12:18)], "Candidate_gene_families_and_tolerant_specific_expansions.csv", row.names = F)
```
