## Additional File 3 for "Integration of differential expression under drought with gene family expansion unique to drought tolerant species predicts candidate genes for drought adaptation in Brassicaceae species": normalize.sh

```
#!/bin/bash

#normalize.sh

WDIR=/data/home/uebermuth/fdi_genofam_mount/uebermuth/brassicaceae-drought/Genomic_Data_BR-dr-03/min100aa/
OUTDIR=${WDIR}/normalized
cd $WDIR
#mkdir $OUTDIR
raw=$(ls *.fa)

for f in $raw; do
        var=${f%.fa}
#	echo "${var}"
        gatk NormalizeFasta -I $f -O ${OUTDIR}/${var}_norm.fa
done;
```
