## Additional File 3 for "Integration of differential expression under drought with gene family expansion unique to drought tolerant species predicts candidate genes for drought adaptation in Brassicaceae species": Selection_overrepresentation_v04_with_corrected_pvalues.pdf

### Gene family expansion and drought DE in Brassicaceae - tests for over representation with genes under positive Selection

#### Combine the results from the A2TEA Analysis with the results from the Selection Analysis

```
setwd("/home/cropbio/sciebo/Documents/PhD/Brassicaceae-drought/BR-dr-03/Selection/Analysis/")
OUTDIR <- "/home/cropbio/sciebo/Documents/PhD/Brassicaceae-drought/BR-dr-03/Selection/Analysis/"
# script to explore a2tea output BR-dr-03 together with results from Selection Analysis and perform over
    representation analyses
suppressMessages(library(Biostrings))
suppressMessages(library(ggplot2))
suppressMessages(library(reshape2))
suppressMessages(library(tidyr))
suppressMessages(library(dplyr))

load("/home/cropbio/sciebo/Documents/PhD/Brassicaceae-drought/BR-dr-03/A2TEA_finished.RData")
```

Count the number of genes which were in the input data set per species (the ones which are in the Conserved OGs file and were considered in the DE Analysis)

```
Conserved_Selection_input_dataset <-  read.table("./results_20221027_Conserved_Dataset/Conserved_dataset_
    matrix.tsv", sep = "\t", header=T)
Conserved_genes_HOG_DE <- merge(Conserved_Selection_input_dataset, HOG_DE.a2tea, by=c("gene", "HOG"))

length(Conserved_genes_HOG_DE[Conserved_genes_HOG_DE$species=="Eutrema_salsugineum",]$gene)
```

```
## [1] 19990
```

```
length(Conserved_genes_HOG_DE[Conserved_genes_HOG_DE$species=="Arabidopsis_lyrata",]$gene)
```

```
## [1] 21173
```

```
length(Conserved_genes_HOG_DE[Conserved_genes_HOG_DE$species=="Arabidopsis_thaliana",]$gene)
```

```
## [1] 19919
```

```
length(Conserved_genes_HOG_DE[Conserved_genes_HOG_DE$species=="Brassica_napus",]$gene)
```

```
## [1] 38861
```

-> the input is more ore less even (Bnapus has double, it is tetraploid)

And from that the OGs that were evaluated in the Selection Analysis load the results from the Selection Analysis

```
Conserved_Selection_results_numbers <- read.table("./results_20221027_Conserved_Dataset/OGs_number_of_
     branches_under_pos_selection.tsv", sep = "\t", header=F)
names(Conserved_Selection_results_numbers) <- c("HOG", "number_of_pos_sel")
Conserved_Selection_results_genes_pvalues <- read.table("./results_20221027_Conserved_Dataset/OGs_pos_
     selection_pvalues.tsv", sep = "\t", header=F)
names(Conserved_Selection_results_genes_pvalues) <- c("HOG", "gene", "uncorrected_pvalue", "selection_
     pvalue")

HOGs_in_results_selection_analysis <- Conserved_Selection_results_numbers$HOG
```

For all calculations on gene level, merge Conserved_Selection_results_genes_pvalues with HOG_DE.a2tea (also to obtain the species per gene) Problem: in Selection results, some IDs of Alyrata are with “_1” in the end or with “pg.” or with “kg.” in the middle. In DE results, the IDs of Alyrata are with “.1” in the end or “pg.” or “kg.”

there are 699 genes with “pg_” in Conserved_Selection_results_genes_pvalues and 8188 genes with “kg_” in Conserved_Selection_results_genes_pvalues

there are 2154 genes with “pg.” in HOG_DE.a2tea$genethereare12428geneswith”kg.”inHOG_DE.a2tea$gene

–> change the “.” used as separator in some Alyrata IDs to “_” in the HOG_DE.a2tea dataset

```
HOG_DE.a2tea_edited_01<- HOG_DE.a2tea
HOG_DE.a2tea_edited_01$gene <- gsub(".", "_", HOG_DE.a2tea_edited_01$gene, fixed = T)


Conserved_Selection_results_genes_pvalues_HOG_DE <- merge(Conserved_Selection_results_genes_pvalues, HOG_
     DE.a2tea_edited_01, by=c("gene", "HOG"))
```

Count the number of genes which were evaluated per species (the ones which are in the results files and as such have a p-value)

```
length(Conserved_Selection_results_genes_pvalues_HOG_DE[Conserved_Selection_results_genes_pvalues_HOG_DE$
     species=="Eutrema_salsugineum",]$gene)
```

```
## [1] 14412
```

```
length(Conserved_Selection_results_genes_pvalues_HOG_DE[Conserved_Selection_results_genes_pvalues_HOG_DE$
     species=="Arabidopsis_lyrata",]$gene)
```

```
## [1] 14974
```

```
length(Conserved_Selection_results_genes_pvalues_HOG_DE[Conserved_Selection_results_genes_pvalues_HOG_DE$
     species=="Arabidopsis_thaliana",]$gene)
```

```
## [1] 14408
```

```
length(Conserved_Selection_results_genes_pvalues_HOG_DE[Conserved_Selection_results_genes_pvalues_HOG_DE$
     species=="Brassica_napus",]$gene)
```

```
## [1] 27962
```

#### Select the genes and HOGs per species which are 1) pos. Sel, 2) DE and 3) exp. and save them in tables per species

### then use these values for over representation tests

load the list of expanded HOGs

```
Eutrema_exp <- read.table("/home/cropbio/sciebo/Documents/PhD/Brassicaceae-drought/BR-dr-03/Analysis/
     Hypothesis_2_Exp_in_Conserved_dataset_matrix.tsv", sep="\t", header=T)
Alyrata_exp <- read.table("/home/cropbio/sciebo/Documents/PhD/Brassicaceae-drought/BR-dr-03/Analysis/
     Hypothesis_3_Exp_in_Conserved_dataset_matrix.tsv", sep="\t", header=T)
Athaliana_exp <- read.table("/home/cropbio/sciebo/Documents/PhD/Brassicaceae-drought/BR-dr-03/Analysis/
     Hypothesis_5_Exp_in_Conserved_dataset_matrix.tsv", sep="\t", header=T)
Bnapus_exp <- read.table("/home/cropbio/sciebo/Documents/PhD/Brassicaceae-drought/BR-dr-03/Analysis/
     Hypothesis_6_Exp_in_Conserved_dataset_matrix.tsv", sep="\t", header=T)
# Phylogenetic expansions
Both_Arabidopsis_exp <- read.table("/home/cropbio/sciebo/Documents/PhD/Brassicaceae-drought/BR-dr-03/
     Analysis/HOGs_exp_2ArabidopsisvsBnapusandEutrema.tsv")
Both_Arabidopsis_and_Eutrema_exp <- read.table("/home/cropbio/sciebo/Documents/PhD/Brassicaceae-drought/
     BR-dr-03/Analysis/HOGs_exp_3vsBnapus.tsv")
Bnapus_vs_all_3_exp <- read.table("/home/cropbio/sciebo/Documents/PhD/Brassicaceae-drought/BR-dr-03/
     Analysis/HOGs_Bnapusexp_vsall3.tsv")
Bnapus_and_Eutrema_vs_Arabidpsis_exp <- read.table("/home/cropbio/sciebo/Documents/PhD/Brassicaceae-
     drought/BR-dr-03/Analysis/HOGS_Esa_Bnaexp_vs_2_Arabidopsis.tsv")
```

create a df that links the abbreviations and the drought phenotype to the species

```
df <- data.frame(abbrev = c("Al", "At", "Es", "Bn"), species = unique(Conserved_Selection_results_genes_
     pvalues_HOG_DE$species), tolerance = c(rep(c("tolerant", "sensitive"), 2)))
#define the expanded HOGs
Al_exp <- unique(Alyrata_exp$HOG)
At_exp <- unique(Athaliana_exp$HOG)
Es_exp <- unique(Eutrema_exp$HOG)
Bn_exp <- unique(Bnapus_exp$HOG)
tolerant_exp <- unique(intersect(Al_exp, Es_exp))
sensitive_exp <- unique(intersect(At_exp, Bn_exp))
```

Loop through the species and count the set sizes. Save this in an output table per species

```
for (x in seq_len(nrow(df))) {
  a <- df[x,1]
  b <- df[x,2]
  c <- df[x,3]
  print(paste0("Conserved_Selection_results_genes_pvalues_HOG_DE_", df[x,1], "_pos", " species is ", df[x
     ,2], "which is", c))
# print(head(get(paste0(a, "_exp"))))
# 1. Select the dataframes with genes from $species which are 1) pos. Sel
  mame <- paste0("Conserved_Selection_results_genes_pvalues_HOG_DE_", a, "_pos")
  data <-
    Conserved_Selection_results_genes_pvalues_HOG_DE[Conserved_Selection_results_genes_pvalues_HOG_DE$
     species==b
                                                     & Conserved_Selection_results_genes_pvalues_HOG_DE$
     selection_pvalue<=0.05,]
#print(head(assign(mame, data)))
# 1a. Select the dataframes with genes from$species which are 2) DE
mame2 <- paste0("Conserved_Selection_results_genes_pvalues_HOG_DE_", a,"_DE")
data2 <-
  Conserved_Selection_results_genes_pvalues_HOG_DE[Conserved_Selection_results_genes_pvalues_HOG_DE$
     species==b
                                                   & (!is.na(Conserved_Selection_results_genes_pvalues_
     HOG_DE$padj))
                                                   & Conserved_Selection_results_genes_pvalues_HOG_DE$
     padj<=0.1,]
#print(head(assign(mame2, data2)))

# 1b. Select the dataframes with all genes from $species from all OGs which have at least one
# 1) pos. Sel and at least one 2) DE from $species
mame2a <- paste0("Conserved_Selection_results_genes_pvalues_HOG_DE_", a,"_pos_and_DE")
data2a <-
  Conserved_Selection_results_genes_pvalues_HOG_DE[Conserved_Selection_results_genes_pvalues_HOG_DE$HOG %
     in% data2$HOG
```

```
                                                    & Conserved_Selection_results_genes_pvalues_HOG_DE$
    species==b
                                                    & Conserved_Selection_results_genes_pvalues_HOG_DE$
    selection_pvalue<=0.05,]

# More detailed:
# Select the dataframes with genes from $species which are 1) pos. Sel and 2) DE
mame3 <- paste0("Conserved_Selection_results_genes_pvalues_HOG_DE_", a, "_DEG_is_pos")
data3 <-
  data2a[(!is.na(data2a$padj)) & data2a$padj<=0.1,]

write.table(data3,paste0(mame3,".tsv"), sep="\t", row.names = F)

# Select the dataframes with genes from $species where the pos. Sel. is NON-DE
mame3a <- paste0("Conserved_Selection_results_genes_pvalues_HOG_DE_", a, "_DE_and_NON_DEG_is_pos")
data3a <-
  data2a[((is.na(data2a$padj)) | data2a$padj>0.1),]

# Select the dataframes with genes from $species which are 1) pos. Sel and 2) exp
mame4 <- paste0("Conserved_Selection_results_genes_pvalues_HOG_DE_", a, "_exp_and_pos")
data4 <-
  data[data$HOG %in% get(paste0(a, "_exp")),]

## in species pos and in both species with same tolerance exp
mame44 <- paste0("Conserved_Selection_results_genes_pvalues_HOG_DE_", c, "_exp_and_", a, "_pos")
data44 <-
  data[data$HOG %in% get(paste0(c, "_exp")),]


# Select the dataframes with genes from $species which are 1) pos. Sel and 2) exp and 3) have a DE from $
    species
# (it is not necessarily the one under pos. Sel.)
mame4a <- paste0("Conserved_Selection_results_genes_pvalues_HOG_DE_", a, "_exp_and_DE_and_pos")
data4a <- data4[data4$HOG %in% data2$HOG,]
## in species pos and DE and in both species with same tolerance exp
mame44a <- paste0("Conserved_Selection_results_genes_pvalues_HOG_DE_", c, "_exp_and_", a, "_DE_and_pos")
data44a <- data4[data4$HOG %in% data2$HOG,]

# More detailed:
# 3. Select the dataframes with genes from $species which are 1) pos. Sel and 2) the same gene is DE and
    3) exp. in $species
mame5 <- paste0("Conserved_Selection_results_genes_pvalues_HOG_DE_", a, "_exp_and_DEG_is_pos")
data5 <-
  data3[data3$HOG %in% get(paste0(a, "_exp")),]

write.table(data5,paste0(mame5,".tsv"), sep="\t", row.names = F)

# 1) pos. Sel and 2) the same gene is DE and 3) exp. in both species with same tolerance
mame55 <- paste0("Conserved_Selection_results_genes_pvalues_HOG_DE_",c , "_exp_and_", a, "_DEG_is_pos")
data55 <-
  data3[data3$HOG %in% get(paste0(c, "_exp")),]


# 3x. Select the dataframes with genes from $species which are in a DE in $species HOG,
# and 1) pos. Sel and 2) a NON-DE gene from $species is pos. sel. and 3) exp. in $species
mame5a <- paste0("Conserved_Selection_results_genes_pvalues_HOG_DE_", a, "_exp_and_DE_and_NON-DEG_is_pos"
    )
data5a <-
  data3a[data3a$HOG %in% get(paste0(a, "_exp")),]
#1) pos. Sel and 2) a NON-DE gene from $species is pos. sel. and 3) exp. in both species with same
    toleranve
mame55a <- paste0("Conserved_Selection_results_genes_pvalues_HOG_DE_", c, "_exp_and_", a, "_DE_and_NON-
    DEG_is_pos")
data55a <-
  data3a[data3a$HOG %in% get(paste0(c, "_exp")),]

# i missed the simple case:
mame5b <- paste0("Conserved_Selection_results_genes_pvalues_HOG_DE_", a, "_exp_and_DE")
data5b <- Conserved_Selection_results_genes_pvalues_HOG_DE[Conserved_Selection_results_genes_pvalues_HOG_
    DE$HOG %in% get(paste0(a, "_exp"))
               & Conserved_Selection_results_genes_pvalues_HOG_DE$HOG %in% data2$HOG,]
# i missed the simple case also with both species with same tolerance exp
mame55b <- paste0("Conserved_Selection_results_genes_pvalues_HOG_DE_", c,"_exp_and_", a, "_DE")
data55b <- Conserved_Selection_results_genes_pvalues_HOG_DE[Conserved_Selection_results_genes_pvalues_HOG
    _DE$HOG %in% get(paste0(c, "_exp"))
               & Conserved_Selection_results_genes_pvalues_HOG_DE$HOG %in% data2$HOG,]



# 4. count number of HOGs (or genes) in the subsets 1) pos. Sel and 2) DE and 3) exp. per $species
```

```
#1)
mame6 <- paste0("number_of_OGs_", a, "_pos")
data6 <- length(unique(data$HOG))
# gene level
mame7 <- paste0("genes_", a, "_pos")
data7 <- length(unique(data[data$species==b,]$gene))

#2a)
mame8 <- paste0("number_of_OGs_", a, "_DEG_is_pos")
data8 <- length(unique(data3$HOG))
# gene level
mame9 <- paste0("genes_", a, "_DEG_is_pos")
data9 <- length(unique(data3[data3$species==b,]$gene))

# calculate the number of HOGs within the HOGs DE in species, where the pos. sel. gene from species is
    NOT DE
mame10 <- paste0("number_of_OGs_", a, "_DE_and_NON-DEG_is_pos")
data10 <- length(unique(data3a$HOG))
# gene level
mame11 <- paste0("genes_", a, "_DE_and_NON-DEG_is_pos")
data11 <- length(unique(data3a[data3a$species==b,]$gene))


# more general background:
mame12 <- paste0("number_of_OGs_", a, "_DE")
data12 <- length(unique(data2$HOG))
# gene level:
mame13 <- paste0("genes_", a, "_DE")
data13 <-length(unique(Conserved_Selection_results_genes_pvalues_HOG_DE[Conserved_Selection_results_genes
    _pvalues_HOG_DE$HOG %in% data2$HOG & Conserved_Selection_results_genes_pvalues_HOG_DE$species==b,]$
    gene))
# DEGs:
mame14 <- paste0("DEGs_", a, "_DE")
data14 <- length(unique(data2[data2$species==b,]$gene))


#more general with pos sel.:
mame15 <- paste0("number_of_OGs_", a, "_pos_and_DE")
data15 <- length(unique(data2a$HOG))
# gene level:
mame16 <- paste0("genes_", a, "_in_pos_and_DE")
data16 <- length(unique(data2a[data2a$species==b,]$gene))

#2b)
mame17 <- paste0("number_of_OGs_", a, "_pos_and_exp")
data17 <- length(unique(data4$HOG))
# gene level:
mame18 <- paste0("genes_", a, "_pos_and_exp")
data18 <- length(unique(data4[data4$species==b,]$gene))

# same but exp in both species with same tolerance
mame33 <- paste0("number_of_OGs_", a, "_pos_and_", c, "_exp")
data33 <- length(unique(data44$HOG))
# gene level:
mame34 <- paste0("genes_", a, "_pos_and_", c, "_exp")
data34 <- length(unique(data44[data44$species==b,]$gene))

# exp and DE and pos
mame19 <- paste0("number_of_OGs_", a, "_exp_and_DE_and_pos")
data19 <- length(unique(data4a$HOG))
# gene level:
mame20 <- paste0("genes_", a, "_in_exp_and_DE_and_pos")
data20 <- length(unique(data4a[data4a$species==b,]$gene))

# same but exp in both species with same tolerance
# exp and DE and pos
mame35 <- paste0("number_of_OGs_", c, "_exp_and_", a, "_DE_and_pos")
data35 <- length(unique(data44a$HOG))
# gene level:
mame36 <- paste0("genes_", a, "_in_", c, "_exp_and_", a, "_DE_and_pos")
data36 <- length(unique(data4a[data4a$species==b,]$gene))

#3) exp and DE is pos
mame21 <- paste0("number_of_OGs_", a, "_exp_and_DE_is_pos")
data21 <- length(unique(data5$HOG))
# gene level
mame22 <- paste0("genes_", a, "_exp_and_DE_is_pos")
data22 <- length(unique(data5$gene))

# same but exp in both species with same tolerance
#3) exp and DE is pos
```

```
mame37 <- paste0("number_of_OGs_", c, "_exp_and_", a, "_DE_is_pos")
data37 <- length(unique(data55$HOG))
# gene level
mame38 <- paste0("genes_", c, "_exp_and_", a, "_DE_is_pos")
data38 <- length(unique(data55$gene))


# the contrast: exp and DE and the NON-DE is pos
mame23 <- paste0("number_of_OGs_", a, "_exp_and_DE_and_NON-DE_is_pos")
data23 <- length(unique(data5a$HOG))
# gene level
mame24 <- paste0("genes_", a, "_exp_and_DE_and_NON-DE_is_pos")
data24 <- length(unique(data5a$gene))

# same but exp in both species with same tolerance
# the contrast: exp and DE and the NON-DE is pos
mame39 <- paste0("number_of_OGs_", c, "_exp_and_", a, "_DE_and_NON-DE_is_pos")
data39 <- length(unique(data55a$HOG))
# gene level
mame40 <- paste0("genes_", c, "_exp_and_", a, "_DE_and_NON-DE_is_pos")
data40 <- length(unique(data55a$gene))


# background set (all conserved HOGs which were evaluated for pos. sel.)
mame26 <- paste0("genes_", a, "_in_Conserved")
data26 <- length(unique(Conserved_Selection_results_genes_pvalues_HOG_DE[
  Conserved_Selection_results_genes_pvalues_HOG_DE$species==b,]$gene))
# same on HOG level
mame25 <- paste0("number_of_OGs_", a, "_in_Conserved")
data25 <- length(unique(Conserved_Selection_results_genes_pvalues_HOG_DE[
  Conserved_Selection_results_genes_pvalues_HOG_DE$species==b,]$HOG))

# background exp
mame27 <- paste0("number_of_OGs_", a, "_exp")
data27 <- length(unique(Conserved_Selection_results_genes_pvalues_HOG_DE[
  Conserved_Selection_results_genes_pvalues_HOG_DE$HOG %in% get(paste0(a, "_exp")),]$HOG))
# gene level
mame28 <- paste0("genes_", a, "_exp")
data28 <- length(unique(Conserved_Selection_results_genes_pvalues_HOG_DE[
  Conserved_Selection_results_genes_pvalues_HOG_DE$HOG %in% get(paste0(a, "_exp"))
  & Conserved_Selection_results_genes_pvalues_HOG_DE$species==b,]$gene))

# same but exp in both species with same tolerance
# background exp
mame41 <- paste0("number_of_OGs_", c, "_exp")
data41 <- length(unique(Conserved_Selection_results_genes_pvalues_HOG_DE[
  Conserved_Selection_results_genes_pvalues_HOG_DE$HOG %in% get(paste0(c, "_exp")),]$HOG))
# gene level
mame42 <- paste0("genes_", a, "_in_", c, "_exp")
data42 <- length(unique(Conserved_Selection_results_genes_pvalues_HOG_DE[
  Conserved_Selection_results_genes_pvalues_HOG_DE$HOG %in% get(paste0(c, "_exp"))
  & Conserved_Selection_results_genes_pvalues_HOG_DE$species==b,]$gene))


mame29 <- paste0("number_of_OGs_", a, "_exp_and_DE")
data29 <- length(unique(data5b$HOG))
# gene level:
mame30 <- paste0("genes_", a, "_exp_and_DE")
data30 <- length(unique(data5b[data5b$species==b,]$gene))
# only the DEGs:
mame31 <- paste0("DEGs_", a, "_exp_and_DE")
data31 <- length(unique(data5b[data5b$species==b
                              & (!is.na(data5b$padj))
                              & data5b$padj<=0.1,]$gene))
# only the NON-DEGs:
mame32 <- paste0("NON-DEGs_", a, "_exp_and_DE")
data32 <- length(unique(data5b[data5b$species==b
                              & (is.na(data5b$padj)
                              | data5b$padj>0.1),]$gene))

# same but exp in both species with same tolerance
mame43 <- paste0("number_of_OGs_", c, "_exp_and_", a, "_DE")
data43 <- length(unique(data55b$HOG))
# gene level:
mame44 <- paste0("genes_", a, "_in_", c, "_exp_and_", a, "_DE")
data44 <- length(unique(data55b[data55b$species==b,]$gene))
# only the DEGs:
mame45 <- paste0("DEGs_", a, "_in_", c, "_exp_and_", a, "_DE")
data45 <- length(unique(data55b[data55b$species==b
                              & (!is.na(data55b$padj))
                              & data55b$padj<=0.1,]$gene))
```

```r
# only the NON-DEGs:
mame46 <- paste0("NON-DEGs_", a, "_in_", c, "_exp_and_", a, "_DE")
data46 <- length(unique(data55b[data55b$species==b
                               & (is.na(data55b$padj)
                               | data55b$padj>0.1),]$gene))


# put the names of the sets into a character vector
set_names <- c(paste0("mame", seq(6,46)))
#print(set_names)
print(as.character(sapply(set_names, get)))

# put the counts of the sets into a numeric vector
set_values <- c(paste0("data", seq(6,46)))
#print(set_values)
print(as.numeric(sapply(set_values, get)))

# combine the names and the counts of the sets to a data.frame
df_name <- paste0("counts_pos_sel_", a)
output_df <- data.frame(set_names=as.character(sapply(set_names, get)), counts=as.numeric(sapply(set_
    values, get)))
# save the output_df in a variable
assign(df_name, output_df)
# export the data.frame
write.table(output_df, paste0(OUTDIR, df_name, ".tsv"), sep="\t", row.names = F, col.names = T)
}
```

```
## [1] "Conserved_Selection_results_genes_pvalues_HOG_DE_Al_pos species is Arabidopsis_lyratawhich
    istolerant"
##  [1] "number_of_OGs_Al_pos"
##  [2] "genes_Al_pos"
##  [3] "number_of_OGs_Al_DEG_is_pos"
##  [4] "genes_Al_DEG_is_pos"
##  [5] "number_of_OGs_Al_DE_and_NON-DEG_is_pos"
##  [6] "genes_Al_DE_and_NON-DEG_is_pos"
##  [7] "number_of_OGs_Al_DE"
##  [8] "genes_Al_DE"
##  [9] "DEGs_Al_DE"
## [10] "number_of_OGs_Al_pos_and_DE"
## [11] "genes_Al_in_pos_and_DE"
## [12] "number_of_OGs_Al_pos_and_exp"
## [13] "genes_Al_pos_and_exp"
## [14] "number_of_OGs_Al_exp_and_DE_and_pos"
## [15] "genes_Al_in_exp_and_DE_and_pos"
## [16] "number_of_OGs_Al_exp_and_DE_is_pos"
## [17] "genes_Al_exp_and_DE_is_pos"
## [18] "number_of_OGs_Al_exp_and_DE_and_NON-DE_is_pos"
## [19] "genes_Al_exp_and_DE_and_NON-DE_is_pos"
## [20] "number_of_OGs_Al_in_Conserved"
## [21] "genes_Al_in_Conserved"
## [22] "number_of_OGs_Al_exp"
## [23] "genes_Al_exp"
## [24] "number_of_OGs_Al_exp_and_DE"
## [25] "genes_Al_exp_and_DE"
## [26] "DEGs_Al_exp_and_DE"
## [27] "NON-DEGs_Al_exp_and_DE"
## [28] "number_of_OGs_Al_pos_and_tolerant_exp"
## [29] "genes_Al_pos_and_tolerant_exp"
## [30] "number_of_OGs_tolerant_exp_and_Al_DE_and_pos"
## [31] "genes_Al_in_tolerant_exp_and_Al_DE_and_pos"
## [32] "number_of_OGs_tolerant_exp_and_Al_DE_is_pos"
## [33] "genes_tolerant_exp_and_Al_DE_is_pos"
## [34] "number_of_OGs_tolerant_exp_and_Al_DE_and_NON-DE_is_pos"
## [35] "genes_tolerant_exp_and_Al_DE_and_NON-DE_is_pos"
## [36] "number_of_OGs_tolerant_exp"
## [37] "genes_Al_in_tolerant_exp"
## [38] "number_of_OGs_tolerant_exp_and_Al_DE"
## [39] "genes_Al_in_tolerant_exp_and_Al_DE"
## [40] "DEGs_Al_in_tolerant_exp_and_Al_DE"
## [41] "NON-DEGs_Al_in_tolerant_exp_and_Al_DE"
##  [1]  1335  1423   536   544   187   204  6852  7919  7209   696   748   272
## [13]   326   164   197    64    70   115   127 13257 14974   583  1330   332
## [25]   747   423   324    33    41   164   197     5     5     7     7    69
## [37]   174    32    68    44    24
## [1] "Conserved_Selection_results_genes_pvalues_HOG_DE_At_pos species is Arabidopsis_thalianawhich
    issensitive"
##  [1] "number_of_OGs_At_pos"
##  [2] "genes_At_pos"
```

```
## [3] "number_of_OGs_At_DEG_is_pos"
## [4] "genes_At_DEG_is_pos"
## [5] "number_of_OGs_At_DE_and_NON-DEG_is_pos"
## [6] "genes_At_DE_and_NON-DEG_is_pos"
## [7] "number_of_OGs_At_DE"
## [8] "genes_At_DE"
## [9] "DEGs_At_DE"
## [10] "number_of_OGs_At_pos_and_DE"
## [11] "genes_At_in_pos_and_DE"
## [12] "number_of_OGs_At_pos_and_exp"
## [13] "genes_At_pos_and_exp"
## [14] "number_of_OGs_At_exp_and_DE_and_pos"
## [15] "genes_At_in_exp_and_DE_and_pos"
## [16] "number_of_OGs_At_exp_and_DE_is_pos"
## [17] "genes_At_exp_and_DE_is_pos"
## [18] "number_of_OGs_At_exp_and_DE_and_NON-DE_is_pos"
## [19] "genes_At_exp_and_DE_and_NON-DE_is_pos"
## [20] "number_of_OGs_At_in_Conserved"
## [21] "genes_At_in_Conserved"
## [22] "number_of_OGs_At_exp"
## [23] "genes_At_exp"
## [24] "number_of_OGs_At_exp_and_DE"
## [25] "genes_At_exp_and_DE"
## [26] "DEGs_At_exp_and_DE"
## [27] "NON-DEGs_At_exp_and_DE"
## [28] "number_of_OGs_At_pos_and_sensitive_exp"
## [29] "genes_At_pos_and_sensitive_exp"
## [30] "number_of_OGs_sensitive_exp_and_At_DE_and_pos"
## [31] "genes_At_in_sensitive_exp_and_At_DE_and_pos"
## [32] "number_of_OGs_sensitive_exp_and_At_DE_is_pos"
## [33] "genes_sensitive_exp_and_At_DE_is_pos"
## [34] "number_of_OGs_sensitive_exp_and_At_DE_and_NON-DE_is_pos"
## [35] "genes_sensitive_exp_and_At_DE_and_NON-DE_is_pos"
## [36] "number_of_OGs_sensitive_exp"
## [37] "genes_At_in_sensitive_exp"
## [38] "number_of_OGs_sensitive_exp_and_At_DE"
## [39] "genes_At_in_sensitive_exp_and_At_DE"
## [40] "DEGs_At_in_sensitive_exp_and_At_DE"
## [41] "NON-DEGs_At_in_sensitive_exp_and_At_DE"
## [1]  1127  1165   133   133    31    33  2178  2524  2253   160   166    83
## [13]    97    16    20     6     6    12    14 13255 14408   223   488    50
## [25]   118    62    56    15    17    16    20     1     1     2     2    32
## [37]    68     9    19    13     6
## [1] "Conserved_Selection_results_genes_pvalues_HOG_DE_Es_pos species is Eutrema_salsugineumwhich
    istolerant"
## [1] "number_of_OGs_Es_pos"
## [2] "genes_Es_pos"
## [3] "number_of_OGs_Es_DEG_is_pos"
## [4] "genes_Es_DEG_is_pos"
## [5] "number_of_OGs_Es_DE_and_NON-DEG_is_pos"
## [6] "genes_Es_DE_and_NON-DEG_is_pos"
## [7] "number_of_OGs_Es_DE"
## [8] "genes_Es_DE"
## [9] "DEGs_Es_DE"
## [10] "number_of_OGs_Es_pos_and_DE"
## [11] "genes_Es_in_pos_and_DE"
## [12] "number_of_OGs_Es_pos_and_exp"
## [13] "genes_Es_pos_and_exp"
## [14] "number_of_OGs_Es_exp_and_DE_and_pos"
## [15] "genes_Es_in_exp_and_DE_and_pos"
## [16] "number_of_OGs_Es_exp_and_DE_is_pos"
## [17] "genes_Es_exp_and_DE_is_pos"
## [18] "number_of_OGs_Es_exp_and_DE_and_NON-DE_is_pos"
## [19] "genes_Es_exp_and_DE_and_NON-DE_is_pos"
## [20] "number_of_OGs_Es_in_Conserved"
## [21] "genes_Es_in_Conserved"
## [22] "number_of_OGs_Es_exp"
## [23] "genes_Es_exp"
## [24] "number_of_OGs_Es_exp_and_DE"
## [25] "genes_Es_exp_and_DE"
## [26] "DEGs_Es_exp_and_DE"
## [27] "NON-DEGs_Es_exp_and_DE"
## [28] "number_of_OGs_Es_pos_and_tolerant_exp"
## [29] "genes_Es_pos_and_tolerant_exp"
## [30] "number_of_OGs_tolerant_exp_and_Es_DE_and_pos"
## [31] "genes_Es_in_tolerant_exp_and_Es_DE_and_pos"
## [32] "number_of_OGs_tolerant_exp_and_Es_DE_is_pos"
## [33] "genes_tolerant_exp_and_Es_DE_is_pos"
## [34] "number_of_OGs_tolerant_exp_and_Es_DE_and_NON-DE_is_pos"
## [35] "genes_tolerant_exp_and_Es_DE_and_NON-DE_is_pos"
## [36] "number_of_OGs_tolerant_exp"
```

```
## [37] "genes_Es_in_tolerant_exp"
## [38] "number_of_OGs_tolerant_exp_and_Es_DE"
## [39] "genes_Es_in_tolerant_exp_and_Es_DE"
## [40] "DEGs_Es_in_tolerant_exp_and_Es_DE"
## [41] "NON-DEGs_Es_in_tolerant_exp_and_Es_DE"
##  [1]  1318  1371   486   489    82    86  5750  6397  5971   551   575   179
## [13]   209    90   104    39    41    60    63 13256 14412   438   987   230
## [25]   512   296   216    26    33    90   104     6     7     6     6    69
## [37]   167    34    75    48    27
## [1] "Conserved_Selection_results_genes_pvalues_HOG_DE_Bn_pos species is Brassica_napuswhich
    issensitive"
##  [1] "number_of_OGs_Bn_pos"
##  [2] "genes_Bn_pos"
##  [3] "number_of_OGs_Bn_DEG_is_pos"
##  [4] "genes_Bn_DEG_is_pos"
##  [5] "number_of_OGs_Bn_DE_and_NON-DEG_is_pos"
##  [6] "genes_Bn_DE_and_NON-DEG_is_pos"
##  [7] "number_of_OGs_Bn_DE"
##  [8] "genes_Bn_DE"
##  [9] "DEGs_Bn_DE"
## [10] "number_of_OGs_Bn_pos_and_DE"
## [11] "genes_Bn_in_pos_and_DE"
## [12] "number_of_OGs_Bn_pos_and_exp"
## [13] "genes_Bn_pos_and_exp"
## [14] "number_of_OGs_Bn_exp_and_DE_and_pos"
## [15] "genes_Bn_in_exp_and_DE_and_pos"
## [16] "number_of_OGs_Bn_exp_and_DE_is_pos"
## [17] "genes_Bn_exp_and_DE_is_pos"
## [18] "number_of_OGs_Bn_exp_and_DE_and_NON-DE_is_pos"
## [19] "genes_Bn_exp_and_DE_and_NON-DE_is_pos"
## [20] "number_of_OGs_Bn_in_Conserved"
## [21] "genes_Bn_in_Conserved"
## [22] "number_of_OGs_Bn_exp"
## [23] "genes_Bn_exp"
## [24] "number_of_OGs_Bn_exp_and_DE"
## [25] "genes_Bn_exp_and_DE"
## [26] "DEGs_Bn_exp_and_DE"
## [27] "NON-DEGs_Bn_exp_and_DE"
## [28] "number_of_OGs_Bn_pos_and_sensitive_exp"
## [29] "genes_Bn_pos_and_sensitive_exp"
## [30] "number_of_OGs_sensitive_exp_and_Bn_DE_and_pos"
## [31] "genes_Bn_in_sensitive_exp_and_Bn_DE_and_pos"
## [32] "number_of_OGs_sensitive_exp_and_Bn_DE_is_pos"
## [33] "genes_sensitive_exp_and_Bn_DE_is_pos"
## [34] "number_of_OGs_sensitive_exp_and_Bn_DE_and_NON-DE_is_pos"
## [35] "genes_sensitive_exp_and_Bn_DE_and_NON-DE_is_pos"
## [36] "number_of_OGs_sensitive_exp"
## [37] "genes_Bn_in_sensitive_exp"
## [38] "number_of_OGs_sensitive_exp_and_Bn_DE"
## [39] "genes_Bn_in_sensitive_exp_and_Bn_DE"
## [40] "DEGs_Bn_in_sensitive_exp_and_Bn_DE"
## [41] "NON-DEGs_Bn_in_sensitive_exp_and_Bn_DE"
##  [1]  2320  2732   558   596   472   543  5432 12290  7773   957  1139   405
## [13]   565   226   313    95   109   157   204 13257 27962  1228  5429   675
## [25]  3018  1351  1667    14    21   226   313     2     3     5     6    32
## [37]   144    15    64    36    28
```

**Over representation analyses**

### Hypergeometric test for over representation of HOGs with pos sel in HOG in "Conserved" Data Set Hypergeometric test (comparable to Fisher's exact test) for enrichment of exp. SPECIES OGs with pos selection over exp.SPECIES OGs in Conserved

q = number of white balls (success) drawn without replacement from an urn which contains both black and white balls m = the number of white balls in the urn. (success in BG) n = the number of black balls in the urn. (BG - m) k = the number of balls drawn from the urn (number of balls in sample)

**first try only Eutrema expanded HOGs**

#### Pos Sel $species in $species expanded HOGs compared to Pos Sel $species in Conserved HOGs

Hypothesis 2: Eutrema expanded vs. Conserved

```
q <- counts_pos_sel_Es[counts_pos_sel_Es$set_names=="number_of_OGs_Es_pos_and_exp",]$counts
m <- counts_pos_sel_Es[counts_pos_sel_Es$set_names=="number_of_OGs_Es_pos",]$counts
n <- (counts_pos_sel_Es[counts_pos_sel_Es$set_names=="number_of_OGs_Es_in_Conserved",]$counts - m)
k <- counts_pos_sel_Es[counts_pos_sel_Es$set_names=="number_of_OGs_Es_exp",]$counts

p <- phyper((q-1), m, n, k, lower.tail=F)
p # 3.387887e-68
```

```
## [1] 3.387887e-68
```

**Generalize the test per species:**

#### Pos Sel $species in $species exp HOGs compared to Pos Sel $species in Conserved HOGs

```
df <- data.frame(abbrev = c("Al", "At", "Es", "Bn"), species = unique(Conserved_Selection_results_genes_
     pvalues_HOG_DE$species))


for (x in seq_len(nrow(df))) {
# define the name of the test
    c <- "exp_in_Conserved"
  a <- df[x,1]
  b <- df[x,2]

# select the proper sets to compare their sizes
  q <- get(paste0("counts_pos_sel_", a))[get(paste0("counts_pos_sel_", a))$set_names==(paste0("number_of_
     OGs_", a, "_pos_and_exp")),]$counts
  q_name <- paste0(c, "_success_in_sample_", a)
  assign(q_name, q)

  m <- get(paste0("counts_pos_sel_", a))[get(paste0("counts_pos_sel_", a))$set_names==(paste0("number_of_
     OGs_", a, "_pos")),]$counts
   m_name <- paste0(c, "_success_in_BG_", a)
  assign(m_name, m)

  n <- (get(paste0("counts_pos_sel_", a))[get(paste0("counts_pos_sel_", a))$set_names==(paste0("number_of
     _OGs_", a, "_in_Conserved")),]$counts - m)
   n_name <- paste0(c, "_non_success_in_BG_", a)
  assign(n_name, n)

  k <- get(paste0("counts_pos_sel_", a))[get(paste0("counts_pos_sel_", a))$set_names==(paste0("number_of_
     OGs_", a, "_exp")),]$counts
 k_name <- paste0(c, "_sample_", a)
  assign(k_name, k)

  p_name <- paste0(c, "_phyper_", a)
  p <- phyper((q-1), m, n, k, lower.tail=F)
  assign(p_name, p)

  print(paste(b, "phyper = ", p, sep= ))
  #print(p)
  my_results <-c(round(q,2), round(k,2), round((q/k),2), round(m,2), round(n,2), round((m/(n+m)),2))
  print(format(my_results, scientific = F))
}
```

```
## [1] "Arabidopsis_lyrata phyper =  9.19942087739887e-122"
## [1] "  272.00" "  583.00" "    0.47" " 1335.00" "11922.00" "    0.10"
## [1] "Arabidopsis_thaliana phyper =  4.08411089394162e-33"
## [1] "   83.00" "  223.00" "    0.37" " 1127.00" "12128.00" "    0.09"
## [1] "Eutrema_salsugineum phyper =  3.38788742282084e-68"
## [1] "  179.00" "  438.00" "    0.41" " 1318.00" "11938.00" "    0.10"
## [1] "Brassica_napus phyper =  6.5380864091395e-44"
## [1] "  405.00" " 1228.00" "    0.33" " 2320.00" "10937.00" "    0.18"
```

–> All HOGs expanded in species contain more often a gene under pos sel from species.

2 species expanded: Pos Sel species in both species with same tolerance expanded HOGs compared to Pos Sel $species in Conserved HOGs

```
df <- data.frame(abbrev = c("Al", "At", "Es", "Bn"), species = unique(Conserved_Selection_results_genes_
    pvalues_HOG_DE$species), tolerance=c(rep(c("tolerant", "sensitive"), 2)))


for (x in seq_len(nrow(df))) {
# define the name of the test
    c <- "same_tolerance_both_exp_in_Conserved"
  a <- df[x,1]
  b <- df[x,2]
  c <- df[x,3]

# select the proper sets to compare their sizes
  q <- get(paste0("counts_pos_sel_", a))[get(paste0("counts_pos_sel_", a))$set_names==(paste0("number_of_
    OGs_", a, "_pos_and_", c, "_exp")),]$counts
  q_name <- paste0(c, "_success_in_sample_", a)
  assign(q_name, q)

  m <- get(paste0("counts_pos_sel_", a))[get(paste0("counts_pos_sel_", a))$set_names==(paste0("number_of_
    OGs_", a, "_pos")),]$counts
   m_name <- paste0(c, "_success_in_BG_", a)
  assign(m_name, m)

  n <- (get(paste0("counts_pos_sel_", a))[get(paste0("counts_pos_sel_", a))$set_names==(paste0("number_of
    _OGs_", a, "_in_Conserved")),]$counts - m)
   n_name <- paste0(c, "_non_success_in_BG_", a)
  assign(n_name, n)

  k <- get(paste0("counts_pos_sel_", a))[get(paste0("counts_pos_sel_", a))$set_names==(paste0("number_of_
    OGs_", c, "_exp")),]$counts
 k_name <- paste0(c, "_sample_", a)
  assign(k_name, k)

  p_name <- paste0(c, "_phyper_", a)
  p <- phyper((q-1), m, n, k, lower.tail=F)
  assign(p_name, p)

  print(paste(b, c, "phyper = ", p, sep= ))
  #print(p)
  my_results <-c(round(q,2), round(k,2), round((q/k),2), round(m,2), round(n,2), round((m/(n+m)),2))
  print(format(my_results, scientific = F))
}
```

```
## [1] "Arabidopsis_lyrata tolerant phyper =  1.25537813236025e-15"
## [1] "   33.00" "   69.00" "    0.48" " 1335.00" "11922.00" "    0.10"
## [1] "Arabidopsis_thaliana sensitive phyper =  1.13369292844997e-08"
## [1] "   15.00" "   32.00" "    0.47" " 1127.00" "12128.00" "    0.09"
## [1] "Eutrema_salsugineum tolerant phyper =  6.9768112199441e-10"
## [1] "   26.00" "   69.00" "    0.38" " 1318.00" "11938.00" "    0.10"
## [1] "Brassica_napus sensitive phyper =  0.000486551639644687"
## [1] "   14.00" "   32.00" "    0.44" " 2320.00" "10937.00" "    0.18"
```

–> All HOGs in expanded in both species with same tolerance contain more genes from species under pos sel.

### On gene level: Pos Sel gene from species in species exp HOGs compared to Pos Sel gene from species in Conserved HOGs

```
df <- data.frame(abbrev = c("Al", "At", "Es", "Bn"), species = unique(Conserved_Selection_results_genes_
    pvalues_HOG_DE$species))


for (x in seq_len(nrow(df))) {
# define the name of the test
    c <- "genes_exp_in_Conserved"
  a <- df[x,1]
  b <- df[x,2]

# select the proper sets to compare their sizes
  q <- get(paste0("counts_pos_sel_", a))[get(paste0("counts_pos_sel_", a))$set_names==(paste0("genes_", a
    , "_pos_and_exp")),]$counts
  q_name <- paste0(c, "_success_in_sample_", a)
  assign(q_name, q)

  m <- get(paste0("counts_pos_sel_", a))[get(paste0("counts_pos_sel_", a))$set_names==(paste0("genes_", a
    , "_pos")),]$counts
   m_name <- paste0(c, "_success_in_BG_", a)
  assign(m_name, m)

  n <- (get(paste0("counts_pos_sel_", a))[get(paste0("counts_pos_sel_", a))$set_names==(paste0("genes_",
    a, "_in_Conserved")),]$counts - m)
   n_name <- paste0(c, "_non_success_in_BG_", a)
  assign(n_name, n)

  k <- get(paste0("counts_pos_sel_", a))[get(paste0("counts_pos_sel_", a))$set_names==(paste0("genes_", a
    , "_exp")),]$counts
 k_name <- paste0(c, "_sample_", a)
  assign(k_name, k)

  p_name <- paste0(c, "_phyper_", a)
  p <- phyper((q-1), m, n, k, lower.tail=F)
  assign(p_name, p)

  print(paste(b, "phyper = ", p, sep= ))
  #print(p)
  my_results <-c(round(q,2), round(k,2), round((q/k),2), round(m,2), round(n,2), round((m/(n+m)),2))
  print(format(my_results, scientific = F))
}
```

```
## [1] "Arabidopsis_lyrata phyper =  5.89317757026031e-65"
## [1] "  326.00" " 1330.00" "    0.25" " 1423.00" "13551.00" "    0.10"
## [1] "Arabidopsis_thaliana phyper =  3.65744500446229e-17"
## [1] "   97.00" "  488.00" "    0.20" " 1165.00" "13243.00" "    0.08"
## [1] "Eutrema_salsugineum phyper =  1.39126399424335e-30"
## [1] "  209.00" "  987.00" "    0.21" " 1371.00" "13041.00" "    0.10"
## [1] "Brassica_napus phyper =  0.0421312092965297"
## [1] "  565.0" " 5429.0" "    0.1" " 2732.0" "25230.0" "    0.1"
```

–> Enriched in Al, At and Es but only slightly significant in Bn. Genes in expanded HOGs are more often under pos. Selection, but only slightly the ones in HOGs expanded in Bn. –> this could be due to the more recent WGD.

2 species expanded: Pos Sel species in both species with same tolerance expanded HOGs compared to Pos Sel $species in Conserved HOGs

```
df <- data.frame(abbrev = c("Al", "At", "Es", "Bn"), species = unique(Conserved_Selection_results_genes_
    pvalues_HOG_DE$species), tolerance=c(rep(c("tolerant", "sensitive"), 2)))


for (x in seq_len(nrow(df))) {
# define the name of the test
    c <- "genes_same_tolerance_both_exp_in_Conserved"
  a <- df[x,1]
  b <- df[x,2]
  c <- df[x,3]
```

```
# select the proper sets to compare their sizes
  q <- get(paste0("counts_pos_sel_", a))[get(paste0("counts_pos_sel_", a))$set_names==(paste0("genes_", a
     , "_pos_and_", c, "_exp")),]$counts
  q_name <- paste0(c, "_success_in_sample_", a)
  assign(q_name, q)

  m <- get(paste0("counts_pos_sel_", a))[get(paste0("counts_pos_sel_", a))$set_names==(paste0("genes_", a
     , "_pos")),]$counts
   m_name <- paste0(c, "_success_in_BG_", a)
  assign(m_name, m)

  n <- (get(paste0("counts_pos_sel_", a))[get(paste0("counts_pos_sel_", a))$set_names==(paste0("genes_",
     a, "_in_Conserved")),]$counts - m)
   n_name <- paste0(c, "_non_success_in_BG_", a)
  assign(n_name, n)

  k <- get(paste0("counts_pos_sel_", a))[get(paste0("counts_pos_sel_", a))$set_names==(paste0("genes_", a
     , "_in_", c, "_exp")),]$counts
 k_name <- paste0(c, "_sample_", a)
  assign(k_name, k)

  p_name <- paste0(c, "_phyper_", a)
  p <- phyper((q-1), m, n, k, lower.tail=F)
  assign(p_name, p)

  print(paste(b, c, "phyper = ", p, sep= ))
  #print(p)
  my_results <-c(round(q,2), round(k,2), round((q/k),2), round(m,2), round(n,2), round((m/(n+m)),2))
  print(format(my_results, scientific = F))
}
```

```
## [1] "Arabidopsis_lyrata tolerant phyper =  3.2183245850215e-08"
## [1] "    41.00" "  174.00" "     0.24" " 1423.00" "13551.00" "     0.10"
## [1] "Arabidopsis_thaliana sensitive phyper =  2.05293420662187e-05"
## [1] "    17.00" "   68.00" "     0.25" " 1165.00" "13243.00" "     0.08"
## [1] "Eutrema_salsugineum tolerant phyper =  3.80311352336892e-05"
## [1] "    33.0" "  167.0" "     0.2" " 1371.0" "13041.0" "     0.1"
## [1] "Brassica_napus sensitive phyper =  0.0408287238531039"
## [1] "    21.00" "  144.00" "     0.15" " 2732.00" "25230.00" "     0.10"
```

–> Enriched in all subsets, but only slightly in pos sel genes from Bnapus in all sens exp –> I interpret that in the HOGs, which are expanded in both sensitive, there are more adaptive genes from Bnapus (the older gene duplicates!) than in the HOGs which are expanded only in Bnapus (newer duplications)

#### Pos Sel from species in both species with same tolerance expanded HOGs compared to Pos Sel from species in in species expanded HOGs

```
df <- data.frame(abbrev = c("Al", "At", "Es", "Bn"), species = unique(Conserved_Selection_results_genes_
     pvalues_HOG_DE$species), tolerance=c(rep(c("tolerant", "sensitive"), 2)))


for (x in seq_len(nrow(df))) {
# define the name of the test
    c <- "same_tolerance_both_exp_in_species_exp"
  a <- df[x,1]
  b <- df[x,2]
  c <- df[x,3]

# select the proper sets to compare their sizes
  q <- get(paste0("counts_pos_sel_", a))[get(paste0("counts_pos_sel_", a))$set_names==(paste0("number_of_
     OGs_", a, "_pos_and_", c, "_exp")),]$counts
  q_name <- paste0(c, "_success_in_sample_", a)
  assign(q_name, q)

  m <- get(paste0("counts_pos_sel_", a))[get(paste0("counts_pos_sel_", a))$set_names==(paste0("number_of_
     OGs_", a, "_pos_and_exp")),]$counts
   m_name <- paste0(c, "_success_in_BG_", a)
  assign(m_name, m)

  n <- (get(paste0("counts_pos_sel_", a))[get(paste0("counts_pos_sel_", a))$set_names==(paste0("number_of
     _OGs_", a, "_exp")),]$counts - m)
```

```
  n_name <- paste0(c, "_non_success_in_BG_", a)
  assign(n_name, n)

  k <- get(paste0("counts_pos_sel_", a))[get(paste0("counts_pos_sel_", a))$set_names==(paste0("number_of_
    OGs_", c, "_exp")),]$counts
 k_name <- paste0(c, "_sample_", a)
  assign(k_name, k)

  p_name <- paste0(c, "_phyper_", a)
  p <- phyper((q-1), m, n, k, lower.tail=F)
  assign(p_name, p)

  print(paste(b, c, "phyper = ", p, sep= ))
  #print(p)
  my_results <-c(round(q,2), round(k,2), round((q/k),2), round(m,2), round(n,2), round((m/(n+m)),2))
  print(format(my_results, scientific = F))
}
```

```
## [1] "Arabidopsis_lyrata tolerant phyper =  0.467625484322633"
## [1] " 33.00" " 69.00" "  0.48" "272.00" "311.00" "  0.47"
## [1] "Arabidopsis_thaliana sensitive phyper =  0.153135479129356"
## [1] " 15.00" " 32.00" "  0.47" " 83.00" "140.00" "  0.37"
## [1] "Eutrema_salsugineum tolerant phyper =  0.763303190204148"
## [1] " 26.00" " 69.00" "  0.38" "179.00" "259.00" "  0.41"
## [1] "Brassica_napus sensitive phyper =  0.131615419008599"
## [1] " 14.00" " 32.00" "  0.44" "405.00" "823.00" "  0.33"
```

–> None is significantly enriched.

### On gene level:

```
df <- data.frame(abbrev = c("Al", "At", "Es", "Bn"), species = unique(Conserved_Selection_results_genes_
    pvalues_HOG_DE$species), tolerance=c(rep(c("tolerant", "sensitive"), 2)))


for (x in seq_len(nrow(df))) {
# define the name of the test
    c <- "same_tolerance_both_exp_in_species_exp_gene_level"
  a <- df[x,1]
  b <- df[x,2]
  c <- df[x,3]

# select the proper sets to compare their sizes
  q <- get(paste0("counts_pos_sel_", a))[get(paste0("counts_pos_sel_", a))$set_names==(paste0("genes_", a
    , "_pos_and_", c, "_exp")),]$counts
  q_name <- paste0(c, "_success_in_sample_", a)
  assign(q_name, q)

  m <- get(paste0("counts_pos_sel_", a))[get(paste0("counts_pos_sel_", a))$set_names==(paste0("genes_", a
    , "_pos_and_exp")),]$counts
   m_name <- paste0(c, "_success_in_BG_", a)
  assign(m_name, m)

  n <- (get(paste0("counts_pos_sel_", a))[get(paste0("counts_pos_sel_", a))$set_names==(paste0("genes_",
    a, "_exp")),]$counts - m)
   n_name <- paste0(c, "_non_success_in_BG_", a)
  assign(n_name, n)

  k <- get(paste0("counts_pos_sel_", a))[get(paste0("counts_pos_sel_", a))$set_names==(paste0("genes_", a
    , "_in_", c, "_exp")),]$counts
 k_name <- paste0(c, "_sample_", a)
  assign(k_name, k)

  p_name <- paste0(c, "_phyper_", a)
  p <- phyper((q-1), m, n, k, lower.tail=F)
  assign(p_name, p)

  print(paste(b, c, "phyper = ", p, sep= ))
  #print(p)
  my_results <-c(round(q,2), round(k,2), round((q/k),2), round(m,2), round(n,2), round((m/(n+m)),2))
  print(format(my_results, scientific = F))
```

```
}
```

```
## [1] "Arabidopsis_lyrata tolerant phyper =  0.654028186693694"
## [1] "  41.00" " 174.00" "   0.24" " 326.00" "1004.00" "   0.25"
## [1] "Arabidopsis_thaliana sensitive phyper =  0.16376115253321"
## [1] " 17.00" " 68.00" "  0.25" " 97.00" "391.00" "  0.20"
## [1] "Eutrema_salsugineum tolerant phyper =  0.721080329126301"
## [1] " 33.00" "167.00" "  0.20" "209.00" "778.00" "  0.21"
## [1] "Brassica_napus sensitive phyper =  0.0684018799387704"
## [1] "  21.00" " 144.00" "   0.15" " 565.00" "4864.00" "   0.10"
```

–> The sensitive species are slightly enriched, Bnapus nearly significantly.The tolerant species are rather depleted (tendency, not tested).

##### DE HOGs

#### Pos Sel $species in $species DE HOGs compared to Pos Sel $species in Conserved HOGs

```
df <- data.frame(abbrev = c("Al", "At", "Es", "Bn"), species = unique(Conserved_Selection_results_genes_
    pvalues_HOG_DE$species))


for (x in seq_len(nrow(df))) {
# define the name of the test
    c <- "DE_in_Conserved"
  a <- df[x,1]
  b <- df[x,2]

# select the proper sets to compare their sizes
  q <- get(paste0("counts_pos_sel_", a))[get(paste0("counts_pos_sel_", a))$set_names==(paste0("number_of_
    OGs_", a, "_pos_and_DE")),]$counts
  q_name <- paste0(c, "_success_in_sample_", a)
  assign(q_name, q)

  m <- get(paste0("counts_pos_sel_", a))[get(paste0("counts_pos_sel_", a))$set_names==(paste0("number_of_
    OGs_", a, "_pos")),]$counts
   m_name <- paste0(c, "_success_in_BG_", a)
  assign(m_name, m)

  n <- (get(paste0("counts_pos_sel_", a))[get(paste0("counts_pos_sel_", a))$set_names==(paste0("number_of
    _OGs_", a, "_in_Conserved")),]$counts - m)
   n_name <- paste0(c, "_non_success_in_BG_", a)
  assign(n_name, n)

  k <- get(paste0("counts_pos_sel_", a))[get(paste0("counts_pos_sel_", a))$set_names==(paste0("number_of_
    OGs_", a, "_DE")),]$counts
 k_name <- paste0(c, "_sample_", a)
  assign(k_name, k)

  p_name <- paste0(c, "_phyper_", a)
  p <- phyper((q-1), m, n, k, lower.tail=F)
  assign(p_name, p)

  print(paste(b, "phyper = ", p, sep= ))
  #print(p)
  my_results <-c(round(q,2), round(k,2), round((q/k),2), round(m,2), round(n,2), round((m/(n+m)),2))
  print(format(my_results, scientific = F))
}
```

```
## [1] "Arabidopsis_lyrata phyper =  0.375618393229233"
## [1] "  696.0" " 6852.0" "    0.1" " 1335.0" "11922.0" "    0.1"
## [1] "Arabidopsis_thaliana phyper =  0.985706821525994"
## [1] "  160.00" " 2178.00" "    0.07" " 1127.00" "12128.00" "    0.09"
## [1] "Eutrema_salsugineum phyper =  0.892961097799181"
## [1] "  551.0" " 5750.0" "    0.1" " 1318.0" "11938.0" "    0.1"
## [1] "Brassica_napus phyper =  0.39181878385913"
```

```
## [1] "  957.00" " 5432.00" "    0.18" " 2320.00" "10937.00" "    0.18"
```

–> not enriched!

### On gene level: Pos Sel in DEG from $species compared to the Conserved genes from species.

```
df <- data.frame(abbrev = c("Al", "At", "Es", "Bn"), species = unique(Conserved_Selection_results_genes_
    pvalues_HOG_DE$species))


for (x in seq_len(nrow(df))) {
# define the name of the test
    c <- "DEGs_in_Conserved"
  a <- df[x,1]
  b <- df[x,2]

# select the proper sets to compare their sizes
  q <- get(paste0("counts_pos_sel_", a))[get(paste0("counts_pos_sel_", a))$set_names==(paste0("genes_", a
     , "_DEG_is_pos")),]$counts
  q_name <- paste0(c, "_success_in_sample_", a)
  assign(q_name, q)

  m <- get(paste0("counts_pos_sel_", a))[get(paste0("counts_pos_sel_", a))$set_names==(paste0("genes_", a
     , "_pos")),]$counts
   m_name <- paste0(c, "_success_in_BG_", a)
  assign(m_name, m)

  n <- (get(paste0("counts_pos_sel_", a))[get(paste0("counts_pos_sel_", a))$set_names==(paste0("genes_",
     a, "_in_Conserved")),]$counts - m)
   n_name <- paste0(c, "_non_success_in_BG_", a)
  assign(n_name, n)

  k <- get(paste0("counts_pos_sel_", a))[get(paste0("counts_pos_sel_", a))$set_names==(paste0("DEGs_", a,
      "_DE")),]$counts
 k_name <- paste0(c, "_sample_", a)
  assign(k_name, k)

  p_name <- paste0(c, "_phyper_", a)
  p <- phyper((q-1), m, n, k, lower.tail=F)
  assign(p_name, p)

  print(paste(b, "phyper = ", p, sep= ))
  #print(p)
  my_results <-c(round(q,2), round(k,2), round((q/k),2), round(m,2), round(n,2), round((m/(n+m)),2))
  print(format(my_results, scientific = F))
}
```

```
## [1] "Arabidopsis_lyrata phyper =  0.999999999999999"
## [1] "  544.00" " 7209.00" "    0.08" " 1423.00" "13551.00" "    0.10"
## [1] "Arabidopsis_thaliana phyper =  0.999992638343701"
## [1] "  133.00" " 2253.00" "    0.06" " 1165.00" "13243.00" "    0.08"
## [1] "Eutrema_salsugineum phyper =  0.999998033593514"
## [1] "  489.00" " 5971.00" "    0.08" " 1371.00" "13041.00" "    0.10"
## [1] "Brassica_napus phyper =  0.999999999999975"
## [1] "  596.00" " 7773.00" "    0.08" " 2732.00" "25230.00" "    0.10"
```

–> None is enriched.

### And are they sig. depleted? Only change the formula for calculating the p

```r
df <- data.frame(abbrev = c("Al", "At", "Es", "Bn"), species = unique(Conserved_Selection_results_genes_
     pvalues_HOG_DE$species))


for (x in seq_len(nrow(df))) {
# define the name of the test
    c <- "DEGs_in_Conserved"
  a <- df[x,1]
  b <- df[x,2]

# select the proper sets to compare their sizes
  q <- get(paste0("counts_pos_sel_", a))[get(paste0("counts_pos_sel_", a))$set_names==(paste0("genes_", a
     , "_DEG_is_pos")),]$counts
  q_name <- paste0(c, "_success_in_sample_", a)
  assign(q_name, q)

  m <- get(paste0("counts_pos_sel_", a))[get(paste0("counts_pos_sel_", a))$set_names==(paste0("genes_", a
     , "_pos")),]$counts
   m_name <- paste0(c, "_success_in_BG_", a)
  assign(m_name, m)

  n <- (get(paste0("counts_pos_sel_", a))[get(paste0("counts_pos_sel_", a))$set_names==(paste0("genes_",
     a, "_in_Conserved")),]$counts - m)
   n_name <- paste0(c, "_non_success_in_BG_", a)
  assign(n_name, n)

  k <- get(paste0("counts_pos_sel_", a))[get(paste0("counts_pos_sel_", a))$set_names==(paste0("DEGs_", a,
      "_DE")),]$counts
 k_name <- paste0(c, "_sample_", a)
  assign(k_name, k)

  p_name <- paste0(c, "_phyper_", a)
  p <- phyper((q), m, n, k, lower.tail=T)
  assign(p_name, p)

  print(paste(b, "phyper = ", p, sep= ))
  #print(p)
  my_results <-c(round(q,2), round(k,2), round((q/k),2), round(m,2), round(n,2), round((m/(n+m)),2))
  print(format(my_results, scientific = F))
}
```

```
## [1] "Arabidopsis_lyrata phyper =  1.62554279158999e-15"
## [1] "  544.00" " 7209.00" "    0.08" " 1423.00" "13551.00" "    0.10"
## [1] "Arabidopsis_thaliana phyper =  1.10907928867381e-05"
## [1] "  133.00" " 2253.00" "    0.06" " 1165.00" "13243.00" "    0.08"
## [1] "Eutrema_salsugineum phyper =  2.60239254073304e-06"
## [1] "  489.00" " 5971.00" "    0.08" " 1371.00" "13041.00" "    0.10"
## [1] "Brassica_napus phyper =  3.52495939065106e-14"
## [1] "  596.00" " 7773.00" "    0.08" " 2732.00" "25230.00" "    0.10"
```

–> yes, they are sig. depleted

Pos Sel in NON-DEG from $species in DE HOGs compared to the Conserved genes from species in the conserved Set.

```r
df <- data.frame(abbrev = c("Al", "At", "Es", "Bn"), species = unique(Conserved_Selection_results_genes_
     pvalues_HOG_DE$species))


for (x in seq_len(nrow(df))) {
# define the name of the test
    c <- "NON-DEGs_in_Conserved"
  a <- df[x,1]
  b <- df[x,2]

# select the proper sets to compare their sizes
  q <- get(paste0("counts_pos_sel_", a))[get(paste0("counts_pos_sel_", a))$set_names==(paste0("genes_", a
     , "_DE_and_NON-DEG_is_pos")),]$counts
  q_name <- paste0(c, "_success_in_sample_", a)
  assign(q_name, q)

  m <- get(paste0("counts_pos_sel_", a))[get(paste0("counts_pos_sel_", a))$set_names==(paste0("genes_", a
     , "_pos")),]$counts
   m_name <- paste0(c, "_success_in_BG_", a)
  assign(m_name, m)
```

```
  n <- (get(paste0("counts_pos_sel_", a))[get(paste0("counts_pos_sel_", a))$set_names==(paste0("genes_",
     a, "_in_Conserved")),]$counts - m)
   n_name <- paste0(c, "_non_success_in_BG_", a)
  assign(n_name, n)

  k <- get(paste0("counts_pos_sel_", a))[get(paste0("counts_pos_sel_", a))$set_names==(paste0("genes_", a
     , "_DE")),]$counts - get(paste0("counts_pos_sel_", a))[get(paste0("counts_pos_sel_", a))$set_names
     ==(paste0("DEGs_", a, "_DE")),]$counts
 k_name <- paste0(c, "_sample_", a)
  assign(k_name, k)

  p_name <- paste0(c, "_phyper_", a)
  p <- phyper((q-1), m, n, k, lower.tail=F)
  assign(p_name, p)

  print(paste(b, "phyper = ", p, sep= ))
  #print(p)
  my_results <-c(round(q,2), round(k,2), round((q/k),2), round(m,2), round(n,2), round((m/(n+m)),2))
  print(format(my_results, scientific = F))
}
```

```
## [1] "Arabidopsis_lyrata phyper =  5.37545990640208e-51"
## [1] "  204.00" "  710.00" "    0.29" " 1423.00" "13551.00" "    0.10"
## [1] "Arabidopsis_thaliana phyper =  0.0117190006673827"
## [1] "   33.00" "  271.00" "    0.12" " 1165.00" "13243.00" "    0.08"
## [1] "Eutrema_salsugineum phyper =  9.28135798758627e-12"
## [1] "   86.0" "  426.0" "    0.2" " 1371.0" "13041.0" "    0.1"
## [1] "Brassica_napus phyper =  3.26270625072852e-08"
## [1] "  543.00" " 4517.00" "    0.12" " 2732.00" "25230.00" "    0.10"
```

–> All species are enriched for NON-DEGs which are under pos sel. in HOGs which are DE, most prominent is Alyrata.

###### Expanded and DE

##### General: Within all DE HOGs: are there more HOGs with a pos sel. from species in the expanded HOGs compared to the non-expanded HOGs?

### HOG level:

```
df <- data.frame(abbrev = c("Al", "At", "Es", "Bn"), species = unique(Conserved_Selection_results_genes_
     pvalues_HOG_DE$species))


for (x in seq_len(nrow(df))) {
# define the name of the test
    c <- "DE_and_pos_sel_in_exp_HOGs_in_contrast_to_non-exp_HOGs"
  a <- df[x,1]
  b <- df[x,2]

### HOG level!

# select the proper sets to compare their sizes
  q <- get(paste0("counts_pos_sel_", a))[get(paste0("counts_pos_sel_", a))$set_names==(paste0("number_of_
     OGs_", a, "_exp_and_DE_and_pos")),]$counts
  q_name <- paste0(c, "_success_in_sample_", a)
  assign(q_name, q)

  m <- get(paste0("counts_pos_sel_", a))[get(paste0("counts_pos_sel_", a))$set_names==(paste0("number_of_
     OGs_", a, "_pos_and_DE")),]$counts
  m_name <- paste0(c, "_success_in_BG_", a)
  assign(m_name, m)

  n <- (get(paste0("counts_pos_sel_", a))[get(paste0("counts_pos_sel_", a))$set_names==(paste0("number_of
     _OGs_", a, "_DE")),]$counts - m)
  n_name <- paste0(c, "_non_success_in_BG_", a)
  assign(n_name, n)
```

```
  k <- get(paste0("counts_pos_sel_", a))[get(paste0("counts_pos_sel_", a))$set_names==(paste0("number_of_
     OGs_", a, "_exp_and_DE")),]$counts
  k_name <- paste0(c, "_sample_", a)
  assign(k_name, k)

  p_name <- paste0(c, "_phyper_", a)
  p <- phyper((q-1), m, n, k, lower.tail=F)
  assign(p_name, p)

  print(paste(b, "phyper = ", p, sep= ))
  #print(p)
  my_results <-c(round(q,2), round(k,2), round((q/k),2), round(m,2), round(n,2), round((m/(n+m)),2))
  print(format(my_results, scientific = F))
}
```

```
## [1] "Arabidopsis_lyrata phyper =  2.34948731031184e-79"
## [1] " 164.00" " 332.00" "   0.49" " 696.00" "6156.00" "   0.10"
## [1] "Arabidopsis_thaliana phyper =  1.89493454288113e-07"
## [1] "  16.00" "  50.00" "   0.32" " 160.00" "2018.00" "   0.07"
## [1] "Eutrema_salsugineum phyper =  5.19744800261928e-35"
## [1] "  90.00" " 230.00" "   0.39" " 551.00" "5199.00" "   0.10"
## [1] "Brassica_napus phyper =  5.10968812644808e-27"
## [1] " 226.00" " 675.00" "   0.33" " 957.00" "4475.00" "   0.18"
```

–> All species are enriched! Again: this could be due to the larger number of genes from species in the exp HOGs? –> better test on gene level

#### Gene level and More detailed: Within all DE HOGs: are there more DEGs under pos sel. in the expanded HOGs compared to the non-expanded HOGs?

```
df <- data.frame(abbrev = c("Al", "At", "Es", "Bn"), species = unique(Conserved_Selection_results_genes_
     pvalues_HOG_DE$species))


for (x in seq_len(nrow(df))) {
# define the name of the test
    c <- "DEG_is_pos_sel_in_exp_HOGs_in_contrast_to_non-exp_HOGs"
  a <- df[x,1]
  b <- df[x,2]

### gene level!

# select the proper sets to compare their sizes
  q <- get(paste0("counts_pos_sel_", a))[get(paste0("counts_pos_sel_", a))$set_names==(paste0("genes_", a
     , "_exp_and_DE_is_pos")),]$counts
  q_name <- paste0(c, "_success_in_sample_", a)
  assign(q_name, q)

  m <- get(paste0("counts_pos_sel_", a))[get(paste0("counts_pos_sel_", a))$set_names==(paste0("genes_", a
     , "_DEG_is_pos")),]$counts
  m_name <- paste0(c, "_success_in_BG_", a)
  assign(m_name, m)

  n <- (get(paste0("counts_pos_sel_", a))[get(paste0("counts_pos_sel_", a))$set_names==(paste0("DEGs_", a
     , "_DE")),]$counts - m)
  n_name <- paste0(c, "_non_success_in_BG_", a)
  assign(n_name, n)

  k <- get(paste0("counts_pos_sel_", a))[get(paste0("counts_pos_sel_", a))$set_names==(paste0("DEGs_", a,
      "_exp_and_DE")),]$counts
  k_name <- paste0(c, "_sample_", a)
  assign(k_name, k)

  p_name <- paste0(c, "_phyper_", a)
  p <- phyper((q-1), m, n, k, lower.tail=F)
  assign(p_name, p)

  print(paste(b, "phyper = ", p, sep= ))
  #print(p)
  my_results <-c(round(q,2), round(k,2), round((q/k),2), round(m,2), round(n,2), round((m/(n+m)),2))
```

```
  print(format(my_results, scientific = F))
}
```

```
## [1] "Arabidopsis_lyrata phyper =  1.25292653559441e-10"
## [1] "  70.00" " 423.00" "   0.17" " 544.00" "6665.00" "   0.08"
## [1] "Arabidopsis_thaliana phyper =  0.15560443749026"
## [1] "   6.00" "  62.00" "   0.10" " 133.00" "2120.00" "   0.06"
## [1] "Eutrema_salsugineum phyper =  0.000498475535266016"
## [1] "  41.00" " 296.00" "   0.14" " 489.00" "5482.00" "   0.08"
## [1] "Brassica_napus phyper =  0.287872925643616"
## [1] " 109.00" "1351.00" "   0.08" " 596.00" "7177.00" "   0.08"
```

–> Enriched in Al and in Es but not in At and in Bn!

2 species expanded: Pos Sel DEG from species in both species with same tolerance expanded HOGs compared to Pos Sel DEG from species in Conserved HOGs

```
df <- data.frame(abbrev = c("Al", "At", "Es", "Bn"), species = unique(Conserved_Selection_results_genes_
    pvalues_HOG_DE$species), tolerance=c(rep(c("tolerant", "sensitive"), 2)))


for (x in seq_len(nrow(df))) {
# define the name of the test
    c <- "genes_same_tolerance_both_exp_and_DEG_species_in_Conserved"
  a <- df[x,1]
  b <- df[x,2]
  c <- df[x,3]

# select the proper sets to compare their sizes
  q <- get(paste0("counts_pos_sel_", a))[get(paste0("counts_pos_sel_", a))$set_names==(paste0("genes_", c
    , "_exp_and_", a, "_DE_is_pos")),]$counts
  q_name <- paste0(c, "_success_in_sample_", a)
  assign(q_name, q)

  m <- get(paste0("counts_pos_sel_", a))[get(paste0("counts_pos_sel_", a))$set_names==(paste0("genes_", a
    , "_DEG_is_pos")),]$counts
   m_name <- paste0(c, "_success_in_BG_", a)
  assign(m_name, m)

  n <- (get(paste0("counts_pos_sel_", a))[get(paste0("counts_pos_sel_", a))$set_names==(paste0("DEGs_", a
    , "_DE")),]$counts - m)
   n_name <- paste0(c, "_non_success_in_BG_", a)
  assign(n_name, n)
  k <- get(paste0("counts_pos_sel_", a))[get(paste0("counts_pos_sel_", a))$set_names==(paste0("DEGs_", a,
     "_in_", c, "_exp_and_", a, "_DE")),]$counts
 k_name <- paste0(c, "_sample_", a)
  assign(k_name, k)

  p_name <- paste0(c, "_phyper_", a)
  p <- phyper((q-1), m, n, k, lower.tail=F)
  assign(p_name, p)

  print(paste(b, c, "phyper = ", p, sep= ))
  #print(p)
  my_results <-c(round(q,2), round(k,2), round((q/k),2), round(m,2), round(n,2), round((m/(n+m)),2))
  print(format(my_results, scientific = F))
}
```

```
## [1] "Arabidopsis_lyrata tolerant phyper =  0.235597432180182"
## [1] "   5.00" "  44.00" "   0.11" " 544.00" "6665.00" "   0.08"
## [1] "Arabidopsis_thaliana sensitive phyper =  0.547598723897982"
## [1] "   1.00" "  13.00" "   0.08" " 133.00" "2120.00" "   0.06"
## [1] "Eutrema_salsugineum tolerant phyper =  0.0937707171708073"
## [1] "   7.00" "  48.00" "   0.15" " 489.00" "5482.00" "   0.08"
## [1] "Brassica_napus sensitive phyper =  0.528755487245412"
## [1] "   3.00" "  36.00" "   0.08" " 596.00" "7177.00" "   0.08"
```

–> !!! Small sample sizes !!! Only Eutrema is sig. All are slightly enriched.

Now consider DEGs from ANY of Alyrata and Eutrema in subset and bg 2 species expanded: Pos Sel DEG from any of the two species with same tolerance in both species with same tolerance expanded HOGs compared to Pos Sel DEG from any of the two species with same tolerance in Conserved HOGs

```
df <- data.frame(abbrev = c("Al", "At", "Es", "Bn"), species = unique(Conserved_Selection_results_genes_
    pvalues_HOG_DE$species), tolerance=as.factor(c(rep(c("tolerant", "sensitive"), 2))))

for (a in levels(df$tolerance)) {
b <- df[df$tolerance==a,]$species
# select the relevant HOGs:
mamex1 <- paste0("DEG_under_pos_sel_from_any_", a, "_in_DEGs_from_any_", a, "_which_are_exp_in_both_", a)
datax1 <- length(unique(
  Conserved_Selection_results_genes_pvalues_HOG_DE[Conserved_Selection_results_genes_pvalues_HOG_DE$HOG %
    in% get(paste0(a, "_exp"))
                                                    & Conserved_Selection_results_genes_pvalues_HOG_DE$
    species %in% b
                                                    & Conserved_Selection_results_genes_pvalues_HOG_DE$
    selection_pvalue<=0.05
                                                    & (!is.na(Conserved_Selection_results_genes_pvalues_
    HOG_DE$padj))
                                                    & Conserved_Selection_results_genes_pvalues_HOG_DE$
    padj<=0.1,]
  $gene))

mamex2 <- paste0("DEG_under_pos_sel_from_any_", a, "_in_DEGs_from_any_", a)
datax2 <- length(unique(
  Conserved_Selection_results_genes_pvalues_HOG_DE[Conserved_Selection_results_genes_pvalues_HOG_DE$
    species %in% b
                                                    & Conserved_Selection_results_genes_pvalues_HOG_DE$
    selection_pvalue<=0.05
                                                    & (!is.na(Conserved_Selection_results_genes_pvalues_
    HOG_DE$padj))
                                                    & Conserved_Selection_results_genes_pvalues_HOG_DE$
    padj<=0.1,]
  $gene))

mamex3 <- paste0("DEG_from_any_", a)
datax3 <- length(unique(
  Conserved_Selection_results_genes_pvalues_HOG_DE[Conserved_Selection_results_genes_pvalues_HOG_DE$
    species %in% b
                                                    & (!is.na(Conserved_Selection_results_genes_pvalues_
    HOG_DE$padj))
                                                    & Conserved_Selection_results_genes_pvalues_HOG_DE$
    padj<=0.1,]
  $gene))

mamex4 <- paste0("DEG_from_any_", a, "_in_exp_in_both_", a)
datax4 <- length(unique(
  Conserved_Selection_results_genes_pvalues_HOG_DE[Conserved_Selection_results_genes_pvalues_HOG_DE$HOG %
    in% get(paste0(a, "_exp"))
                                                    & Conserved_Selection_results_genes_pvalues_HOG_DE$
    species %in% b
                                                    & (!is.na(Conserved_Selection_results_genes_pvalues_
    HOG_DE$padj))
                                                    & Conserved_Selection_results_genes_pvalues_HOG_DE$
    padj<=0.1,]
  $gene))


# put the names of the sets into a character vector
set_names <- c(paste0("mamex", seq(1,4)))
#print(set_names)
print(as.character(sapply(set_names, get)))

# put the counts of the sets into a numeric vector
set_values <- c(paste0("datax", seq(1,4)))
#print(set_values)
print(as.numeric(sapply(set_values, get)))

# combine the names and the counts of the sets to a data.frame
df_name <- paste0("counts_pos_sel_", a)
output_df <- data.frame(set_names=as.character(sapply(set_names, get)), counts=as.numeric(sapply(set_
    values, get)))
# save the output_df in a variable
assign(df_name, output_df)
# export the data.frame
write.table(output_df, paste0(OUTDIR, df_name, ".tsv"), sep="\t", row.names = F, col.names = T)
}
```

```
## [1] "DEG_under_pos_sel_from_any_sensitive_in_DEGs_from_any_sensitive_which_are_exp_in_both_sensitive"
## [2] "DEG_under_pos_sel_from_any_sensitive_in_DEGs_from_any_sensitive"
## [3] "DEG_from_any_sensitive"
## [4] "DEG_from_any_sensitive_in_exp_in_both_sensitive"
## [1]     4   729 10026    49
## [1] "DEG_under_pos_sel_from_any_tolerant_in_DEGs_from_any_tolerant_which_are_exp_in_both_tolerant"
## [2] "DEG_under_pos_sel_from_any_tolerant_in_DEGs_from_any_tolerant"
## [3] "DEG_from_any_tolerant"
## [4] "DEG_from_any_tolerant_in_exp_in_both_tolerant"
## [1]    12  1033 13180    92
```

```
# perform the test

for (a in levels(df$tolerance)) {

# select the proper sets to compare their sizes
  q <- get(paste0("counts_pos_sel_", a))[get(paste0("counts_pos_sel_", a))$set_names==(paste0("DEG_under_
    pos_sel_from_any_", a, "_in_DEGs_from_any_", a, "_which_are_exp_in_both_", a)),]$counts
  q_name <- paste0(a, "_success_in_sample_", a)
  assign(q_name, q)

  m <- get(paste0("counts_pos_sel_", a))[get(paste0("counts_pos_sel_", a))$set_names==(paste0("DEG_under_
    pos_sel_from_any_", a, "_in_DEGs_from_any_", a)),]$counts
  m_name <- paste0(a, "_success_in_BG_", a)
  assign(m_name, m)

  n <- (get(paste0("counts_pos_sel_", a))[get(paste0("counts_pos_sel_", a))$set_names==(paste0("DEG_from_
    any_", a)),]$counts - m)
  n_name <- paste0(a, "_non_success_in_BG_", a)
  assign(n_name, n)

  k <- get(paste0("counts_pos_sel_", a))[get(paste0("counts_pos_sel_", a))$set_names==(paste0("DEG_from_
    any_", a, "_in_exp_in_both_", a)),]$counts
  k_name <- paste0(a, "_sample_", a)
  assign(k_name, k)

  p_name <- paste0(c, "_phyper_", a)
  p <- phyper((q-1), m, n, k, lower.tail=F)
  assign(p_name, p)

  print(paste(a, "phyper = ", p, sep= ))
  #print(p)
  my_results <-c(round(q,2), round(k,2), round((q/k),2), round(m,2), round(n,2), round((m/(n+m)),2))
  print(format(my_results, scientific = F))
}
```

```
## [1] "sensitive phyper =  0.481661270601279"
## [1] "   4.00" "  49.00" "   0.08" " 729.00" "9297.00" "   0.07"
## [1] "tolerant phyper =  0.0550605023119113"
## [1] "   12.00" "   92.00" "    0.13" " 1033.00" "12147.00" "    0.08"
```

2 species expanded: Pos Sel DEG from species in both species with same tolerance expanded HOGs compared to Pos Sel DEG from species in in species expanded HOGs.

```
df <- data.frame(abbrev = c("Al", "At", "Es", "Bn"), species = unique(Conserved_Selection_results_genes_
    pvalues_HOG_DE$species), tolerance=c(rep(c("tolerant", "sensitive"), 2)))


for (x in seq_len(nrow(df))) {
# define the name of the test
    c <- "DE_species_same_tolerance_both_exp_in_species_exp_and_DE_gene_level"
  a <- df[x,1]
  b <- df[x,2]
  c <- df[x,3]

# select the proper sets to compare their sizes
  q <- get(paste0("counts_pos_sel_", a))[get(paste0("counts_pos_sel_", a))$set_names==(paste0("genes_", c
    , "_exp_and_", a, "_DE_is_pos")),]$counts
  q_name <- paste0(c, "_success_in_sample_", a)
  assign(q_name, q)

  m <- get(paste0("counts_pos_sel_", a))[get(paste0("counts_pos_sel_", a))$set_names==(paste0("genes_", a
    , "_exp_and_DE_is_pos")),]$counts
```

```
  m_name <- paste0(c, "_success_in_BG_", a)
  assign(m_name, m)

    n <- (get(paste0("counts_pos_sel_", a))[get(paste0("counts_pos_sel_", a))$set_names==(paste0("DEGs_",
      a, "_exp_and_DE")),]$counts - m)
   n_name <- paste0(c, "_non_success_in_BG_", a)
  assign(n_name, n)

  k <- get(paste0("counts_pos_sel_", a))[get(paste0("counts_pos_sel_", a))$set_names==(paste0("DEGs_", a,
      "_in_", c, "_exp_and_", a, "_DE")),]$counts
 k_name <- paste0(c, "_sample_", a)
  assign(k_name, k)

  p_name <- paste0(c, "_phyper_", a)
  p <- phyper((q-1), m, n, k, lower.tail=F)
  assign(p_name, p)

  print(paste(b, c, "phyper = ", p, sep= ))
  #print(p)
  my_results <-c(round(q,2), round(k,2), round((q/k),2), round(m,2), round(n,2), round((m/(n+m)),2))
  print(format(my_results, scientific = F))
}
```

```
## [1] "Arabidopsis_lyrata tolerant phyper =  0.887583223214119"
## [1] "  5.00" " 44.00" "  0.11" " 70.00" "353.00" "  0.17"
## [1] "Arabidopsis_thaliana sensitive phyper =  0.772526630098562"
## [1] " 1.00" "13.00" " 0.08" " 6.00" "56.00" " 0.10"
## [1] "Eutrema_salsugineum tolerant phyper =  0.511861826075353"
## [1] "  7.00" " 48.00" "  0.15" " 41.00" "255.00" "  0.14"
## [1] "Brassica_napus sensitive phyper =  0.5665785449429"
## [1] "   3.00" "  36.00" "   0.08" " 109.00" "1242.00" "   0.08"
```

–> !!! Small sample sizes !!! None enriched!

And on HOGs level? Are there more HOGs with DEGs under pos sel. in the expanded HOGs compared to the non-expanded HOGs?

```
df <- data.frame(abbrev = c("Al", "At", "Es", "Bn"), species = unique(Conserved_Selection_results_genes_
     pvalues_HOG_DE$species))


for (x in seq_len(nrow(df))) {
# define the name of the test
    c <- "HOGs_where_DEG_is_pos_sel_in_exp_HOGs_in_contrast_to_non-exp_HOGs"
  a <- df[x,1]
  b <- df[x,2]

### HOG level!

# select the proper sets to compare their sizes
  q <- get(paste0("counts_pos_sel_", a))[get(paste0("counts_pos_sel_", a))$set_names==(paste0("number_of_
     OGs_", a, "_exp_and_DE_is_pos")),]$counts
  q_name <- paste0(c, "_success_in_sample_", a)
  assign(q_name, q)

  m <- get(paste0("counts_pos_sel_", a))[get(paste0("counts_pos_sel_", a))$set_names==(paste0("number_of_
     OGs_", a, "_DEG_is_pos")),]$counts
  m_name <- paste0(c, "_success_in_BG_", a)
  assign(m_name, m)

  n <- (get(paste0("counts_pos_sel_", a))[get(paste0("counts_pos_sel_", a))$set_names==(paste0("number_of
     _OGs_", a, "_DE")),]$counts - m)
  n_name <- paste0(c, "_non_success_in_BG_", a)
  assign(n_name, n)

  k <- get(paste0("counts_pos_sel_", a))[get(paste0("counts_pos_sel_", a))$set_names==(paste0("number_of_
     OGs_", a, "_exp_and_DE")),]$counts
  k_name <- paste0(c, "_sample_", a)
  assign(k_name, k)

  p_name <- paste0(c, "_phyper_", a)
  p <- phyper((q-1), m, n, k, lower.tail=F)
  assign(p_name, p)

  print(paste(b, "phyper = ", p, sep= ))
```

```
  #print(p)
  my_results <-c(round(q,2), round(k,2), round((q/k),2), round(m,2), round(n,2), round((m/(n+m)),2))
  print(format(my_results, scientific = F))
}
```

```
## [1] "Arabidopsis_lyrata phyper =  4.2528381025703e-12"
## [1] "  64.00" " 332.00" "   0.19" " 536.00" "6316.00" "   0.08"
## [1] "Arabidopsis_thaliana phyper =  0.0805676779054649"
## [1] "   6.00" "  50.00" "   0.12" " 133.00" "2045.00" "   0.06"
## [1] "Eutrema_salsugineum phyper =  1.55102188968515e-05"
## [1] "  39.00" " 230.00" "   0.17" " 486.00" "5264.00" "   0.08"
## [1] "Brassica_napus phyper =  0.00050795252231995"
## [1] "  95.00" " 675.00" "   0.14" " 558.00" "4874.00" "   0.10"
```

–> Enriched in all species, but not significant in At. –> on gene level is the better way because it normalizes for the total number of genes from species per HOG.

### And the contrast: Within all DE HOGs: are there more NON-DEGs under pos sel. in the expanded HOGs compared to the non-expanded HOGs?

```
df <- data.frame(abbrev = c("Al", "At", "Es", "Bn"), species = unique(Conserved_Selection_results_genes_
    pvalues_HOG_DE$species))


for (x in seq_len(nrow(df))) {
# define the name of the test
    c <- "NON-DEG_is_pos_sel_in_exp_HOGs_in_contrast_to_non-exp_HOGs"
  a <- df[x,1]
  b <- df[x,2]

### try on gene level!

# select the proper sets to compare their sizes
  q <- get(paste0("counts_pos_sel_", a))[get(paste0("counts_pos_sel_", a))$set_names==(paste0("genes_", a
    , "_exp_and_DE_and_NON-DE_is_pos")),]$counts
  q_name <- paste0(c, "_success_in_sample_", a)
  assign(q_name, q)

  m <- get(paste0("counts_pos_sel_", a))[get(paste0("counts_pos_sel_", a))$set_names==(paste0("genes_", a
    , "_DE_and_NON-DEG_is_pos")),]$counts
  m_name <- paste0(c, "_success_in_BG_", a)
  assign(m_name, m)

  n <- (get(paste0("counts_pos_sel_", a))[get(paste0("counts_pos_sel_", a))$set_names==(paste0("genes_",
    a, "_in_pos_and_DE")),]$counts - m)
  n_name <- paste0(c, "_non_success_in_BG_", a)
  assign(n_name, n)

  # the NON-DEGs in exp_and_DE = genes - DEGs
  k <- get(paste0("counts_pos_sel_", a))[get(paste0("counts_pos_sel_", a))$set_names==(paste0("genes_", a
    , "_exp_and_DE")),]$counts - get(paste0("counts_pos_sel_", a))[get(paste0("counts_pos_sel_", a))$set
    _names==(paste0("DEGs_", a, "_exp_and_DE")),]$counts
  k_name <- paste0(c, "_sample_", a)
  assign(k_name, k)

  p_name <- paste0(c, "_phyper_", a)
  p <- phyper((q-1), m, n, k, lower.tail=F)
  assign(p_name, p)

  print(paste(b, "phyper = ", p, sep= ))
  #print(p)
  my_results <-c(round(q,2), round(k,2), round((q/k),2), round(m,2), round(n,2), round((m/(n+m)),2))
  print(format(my_results, scientific = F))

}
```

```
## [1] "Arabidopsis_lyrata phyper =  1.39530725846974e-10"
## [1] "127.00" "324.00" "  0.39" "204.00" "544.00" "  0.27"
## [1] "Arabidopsis_thaliana phyper =  0.164855023916685"
## [1] " 14.00" " 56.00" "  0.25" " 33.00" "133.00" "  0.20"
## [1] "Eutrema_salsugineum phyper =  3.02710542656994e-13"
## [1] " 63.00" "216.00" "  0.29" " 86.00" "489.00" "  0.15"
```

```
## Warning in phyper((q - 1), m, n, k, lower.tail = F): NaNs produced
```

```
## [1] "Brassica_napus phyper =  NaN"
## [1] " 204.00" "1667.00" "   0.12" " 543.00" " 596.00" "   0.48"
```

–> Al, Es and At (not. sign.) are enriched. but Bn is not (p is 1).

#### Are there more DEGs under pos. Sel. in the exp and DE HOGs than in the Conserved and DE HOGs?

```
df <- data.frame(abbrev = c("Al", "At", "Es", "Bn"), species = unique(Conserved_Selection_results_genes_
    pvalues_HOG_DE$species))


for (x in seq_len(nrow(df))) {
# define the name of the test
    c <- "DEG_is_pos_sel_in_DE_and_exp_HOGs_compared_to_DEG_is_pos_sel_in_Conserved_an"
  a <- df[x,1]
  b <- df[x,2]

#q = pos sel (= success) in a DEG from species within the genes from species in HOGs which are exp + DE
    in species (= success in sample) (genes_Es_exp_and_DE_is_pos)
#m =  pos sel (= success) in any gene from species within the genes from species in HOGs which are exp +
    DE in species (= success in background) (genes_Es_in_exp_and_DE_and_pos)
#n = genes from species in HOGs exp and DE in species (= background) - m (genes_Es_exp_and_DE - m)
#k = DEGs from species in HOGs exp and DE in species (= sample) (DEGs_Es_exp_and_DE)

### try on gene level!

#  s in sample = genes_Es_exp_and_DE_is_pos
#  sample = DEGs_Es_exp_and_DE
#  s in bg = genes_Es_pos_and_exp
#  bg = genes_Es_exp


# select the proper sets to compare their sizes
  q <- get(paste0("counts_pos_sel_", a))[get(paste0("counts_pos_sel_", a))$set_names==(paste0("genes_", a
    , "_exp_and_DE_is_pos")),]$counts
  q_name <- paste0(c, "_success_in_sample_", a)
  assign(q_name, q)

  m <- get(paste0("counts_pos_sel_", a))[get(paste0("counts_pos_sel_", a))$set_names==(paste0("genes_", a
    , "_pos_and_exp")),]$counts
  m_name <- paste0(c, "_success_in_BG_", a)
  assign(m_name, m)

  n <- (get(paste0("counts_pos_sel_", a))[get(paste0("counts_pos_sel_", a))$set_names==(paste0("genes_",
    a, "_exp")),]$counts - m)
  n_name <- paste0(c, "_non_success_in_BG_", a)
  assign(n_name, n)

  k <- get(paste0("counts_pos_sel_", a))[get(paste0("counts_pos_sel_", a))$set_names==(paste0("DEGs_", a,
     "_exp_and_DE")),]$counts
  k_name <- paste0(c, "_sample_", a)
  assign(k_name, k)

  p_name <- paste0(c, "_phyper_", a)
  p <- phyper((q-1), m, n, k, lower.tail=F)
  assign(p_name, p)

  print(paste(b, "phyper = ", p, sep= ))
  #print(p)
```

```
  my_results <-c(round(q,2), round(k,2), round((q/k),2), round(m,2), round(n,2), round((m/(n+m)),2))
  print(format(my_results, scientific = F))
}
```

```
## [1] "Arabidopsis_lyrata phyper =  0.999999185442643"
## [1] "  70.00" " 423.00" "   0.17" " 326.00" "1004.00" "   0.25"
## [1] "Arabidopsis_thaliana phyper =  0.993534554796277"
## [1] "  6.0" " 62.0" "  0.1" " 97.0" "391.0" "  0.2"
## [1] "Eutrema_salsugineum phyper =  0.999948574278826"
## [1] " 41.00" "296.00" "  0.14" "209.00" "778.00" "  0.21"
## [1] "Brassica_napus phyper =  0.999633769165058"
## [1] " 109.00" "1351.00" "   0.08" " 565.00" "4864.00" "   0.10"
```

–> no enrichment

### and are they sig. depleted? only change the formula for the calculion of p

```
df <- data.frame(abbrev = c("Al", "At", "Es", "Bn"), species = unique(Conserved_Selection_results_genes_
    pvalues_HOG_DE$species))


for (x in seq_len(nrow(df))) {
# define the name of the test
    c <- "DEG_is_pos_sel_in_DE_and_exp_HOGs_compared_to_DEG_is_pos_sel_in_Conserved_an"
  a <- df[x,1]
  b <- df[x,2]

#q = pos sel (= success) in a DEG from species within the genes from species in HOGs which are exp + DE
    in species (= success in sample) (genes_Es_exp_and_DE_is_pos)
#m =  pos sel (= success) in any gene from species within the genes from species in HOGs which are exp +
    DE in species (= success in background) (genes_Es_in_exp_and_DE_and_pos)
#n = genes from species in HOGs exp and DE in species (= background) - m (genes_Es_exp_and_DE - m)
#k = DEGs from species in HOGs exp and DE in species (= sample) (DEGs_Es_exp_and_DE)

### try on gene level!

#  s in sample = genes_Es_exp_and_DE_is_pos
#  sample = DEGs_Es_exp_and_DE
#  s in bg = genes_Es_pos_and_exp
#  bg = genes_Es_exp


# select the proper sets to compare their sizes
  q <- get(paste0("counts_pos_sel_", a))[get(paste0("counts_pos_sel_", a))$set_names==(paste0("genes_", a
    , "_exp_and_DE_is_pos")),]$counts
  q_name <- paste0(c, "_success_in_sample_", a)
  assign(q_name, q)

  m <- get(paste0("counts_pos_sel_", a))[get(paste0("counts_pos_sel_", a))$set_names==(paste0("genes_", a
    , "_pos_and_exp")),]$counts
  m_name <- paste0(c, "_success_in_BG_", a)
  assign(m_name, m)

  n <- (get(paste0("counts_pos_sel_", a))[get(paste0("counts_pos_sel_", a))$set_names==(paste0("genes_",
    a, "_exp")),]$counts - m)
  n_name <- paste0(c, "_non_success_in_BG_", a)
  assign(n_name, n)

  k <- get(paste0("counts_pos_sel_", a))[get(paste0("counts_pos_sel_", a))$set_names==(paste0("DEGs_", a,
     "_exp_and_DE")),]$counts
  k_name <- paste0(c, "_sample_", a)
  assign(k_name, k)

  p_name <- paste0(c, "_phyper_", a)
  p <- phyper((q), m, n, k, lower.tail=T)
  assign(p_name, p)

  print(paste(b, "phyper = ", p, sep= ))
  #print(p)
```

```
  my_results <-c(round(q,2), round(k,2), round((q/k),2), round(m,2), round(n,2), round((m/(n+m)),2))
  print(format(my_results, scientific = F))
}
```

```
## [1] "Arabidopsis_lyrata phyper =  1.66025431129948e-06"
## [1] "  70.00" " 423.00" "   0.17" " 326.00" "1004.00" "   0.25"
## [1] "Arabidopsis_thaliana phyper =  0.018526944121884"
## [1] "  6.0" " 62.0" "  0.1" " 97.0" "391.0" "  0.2"
## [1] "Eutrema_salsugineum phyper =  0.00010705472137481"
## [1] " 41.00" "296.00" "  0.14" "209.00" "778.00" "  0.21"
## [1] "Brassica_napus phyper =  0.000540009463561822"
## [1] " 109.00" "1351.00" "   0.08" " 565.00" "4864.00" "   0.10"
```

–> yes, they are sig. depleted

Test the phylogenetic expansions for over representation of genes from species with diversifying selection in the expanded gene families

```
# Phylogenetic expansions
Both_Arabidopsis_exp <- read.table("/home/cropbio/sciebo/Documents/PhD/Brassicaceae-drought/BR-dr-03/
     Analysis/HOGs_exp_2ArabidopsisvsBnapusandEutrema.tsv", header=T)
Both_Arabidopsis_and_Eutrema_exp <- read.table("/home/cropbio/sciebo/Documents/PhD/Brassicaceae-drought/
     BR-dr-03/Analysis/HOGs_exp_3vsBnapus.tsv", header=T)
Bnapus_vs_all_3_exp <- read.table("/home/cropbio/sciebo/Documents/PhD/Brassicaceae-drought/BR-dr-03/
     Analysis/HOGs_Bnapusexp_vsall3.tsv", header=T)
Bnapus_and_Eutrema_vs_Arabidopsis_exp <- read.table("/home/cropbio/sciebo/Documents/PhD/Brassicaceae-
     drought/BR-dr-03/Analysis/HOGS_Esa_Bnaexp_vs_2_Arabidopsis.tsv", header=T)

### select the relevant HOGs:
a <- c("Arabidopsis_thaliana", "Arabidopsis_lyrata")
succes_in_sample <- length(unique(
  Conserved_Selection_results_genes_pvalues_HOG_DE[Conserved_Selection_results_genes_pvalues_HOG_DE$HOG %
     in% Both_Arabidopsis_exp$HOG
                                                    & Conserved_Selection_results_genes_pvalues_HOG_DE$
     species %in% a
                                                    & Conserved_Selection_results_genes_pvalues_HOG_DE$
     selection_pvalue<=0.05,]
  $gene))
success_in_bg <- length(unique(
  Conserved_Selection_results_genes_pvalues_HOG_DE[Conserved_Selection_results_genes_pvalues_HOG_DE$
     species %in% a
                                                    & Conserved_Selection_results_genes_pvalues_HOG_DE$
     selection_pvalue<=0.05,]
  $gene))
sample <- length(unique(
  Conserved_Selection_results_genes_pvalues_HOG_DE[Conserved_Selection_results_genes_pvalues_HOG_DE$HOG %
     in% Both_Arabidopsis_exp$HOG
                                                    & Conserved_Selection_results_genes_pvalues_HOG_DE$
     species %in% a,]
  $gene))
bg <- length(unique(
  Conserved_Selection_results_genes_pvalues_HOG_DE[Conserved_Selection_results_genes_pvalues_HOG_DE$
     species %in% a,]
  $gene))

## perform the test for over representation
# select the proper sets to compare their sizes
  q <- succes_in_sample
  q_name <- paste0("Both_Arabidopsis_exp", "_success_in_sample")
  assign(q_name, q)

  m <- success_in_bg
  m_name <- paste0("Both_Arabidopsis_exp", "_success_in_BG")
  assign(m_name, m)

  n <- (bg - m)
  n_name <- paste0("Both_Arabidopsis_exp", "_non_success_in_BG")
  assign(n_name, n)

  k <- sample
  k_name <- paste0("Both_Arabidopsis_exp", "_sample")
  assign(k_name, k)

  p_name <- paste0("Both_Arabidopsis_exp", "_phyper")
  p <- phyper((q-1), m, n, k, lower.tail=F)
```

```
  assign(p_name, p)

  my_results <-c(round(q,2), round(k,2), round((q/k),2), round(m,2), round(n,2), round((m/(n+m)),2))
  print(paste0("Both_Arabidopsis_exp: ", p))
```

```
## [1] "Both_Arabidopsis_exp: 6.86072958983792e-06"
```

```
  print(format(my_results, scientific = F))
```

```
## [1] "  141.00" " 1113.00" "    0.13" " 2588.00" "26794.00" "    0.09"
```

```
### select the relevant HOGs:
a <- c("Arabidopsis_thaliana", "Arabidopsis_lyrata", "Eutrema_salsugineum")
succes_in_sample <- length(unique(
  Conserved_Selection_results_genes_pvalues_HOG_DE[Conserved_Selection_results_genes_pvalues_HOG_DE$HOG %
    in% Both_Arabidopsis_and_Eutrema_exp$HOG
                                                    & Conserved_Selection_results_genes_pvalues_HOG_DE$
    species %in% a
                                                    & Conserved_Selection_results_genes_pvalues_HOG_DE$
    selection_pvalue<=0.05,]
  $gene))
success_in_bg <- length(unique(
  Conserved_Selection_results_genes_pvalues_HOG_DE[Conserved_Selection_results_genes_pvalues_HOG_DE$
    species %in% a
                                                    & Conserved_Selection_results_genes_pvalues_HOG_DE$
    selection_pvalue<=0.05,]
  $gene))
sample <- length(unique(
  Conserved_Selection_results_genes_pvalues_HOG_DE[Conserved_Selection_results_genes_pvalues_HOG_DE$HOG %
    in% Both_Arabidopsis_and_Eutrema_exp$HOG
                                                    & Conserved_Selection_results_genes_pvalues_HOG_DE$
    species %in% a,]
  $gene))
bg <- length(unique(
  Conserved_Selection_results_genes_pvalues_HOG_DE[Conserved_Selection_results_genes_pvalues_HOG_DE$
    species %in% a,]
  $gene))

## perform the test for over representation
# select the proper sets to compare their sizes
  q <- succes_in_sample
  q_name <- paste0("Both_Arabidopsis_and_Eutrema_exp", "_success_in_sample")
  assign(q_name, q)

  m <- success_in_bg
  m_name <- paste0("Both_Arabidopsis_and_Eutrema_exp", "_success_in_BG")
  assign(m_name, m)

  n <- (bg - m)
  n_name <- paste0("Both_Arabidopsis_and_Eutrema_exp", "_non_success_in_BG")
  assign(n_name, n)

  k <- sample
  k_name <- paste0("Both_Arabidopsis_and_Eutrema_exp", "_sample")
  assign(k_name, k)

  p_name <- paste0("Both_Arabidopsis_and_Eutrema_exp", "_phyper")
  p <- phyper((q-1), m, n, k, lower.tail=F)
  assign(p_name, p)

  my_results <-c(round(q,2), round(k,2), round((q/k),2), round(m,2), round(n,2), round((m/(n+m)),2))
  print(paste0("Both_Arabidopsis_and_Eutrema_exp: ", p))
```

```
## [1] "Both_Arabidopsis_and_Eutrema_exp: 5.70433522259995e-05"
```

```
print(format(my_results, scientific = F))
```

```
## [1] "  217.00" " 1860.00" "    0.12" " 3959.00" "39835.00" "    0.09"
```

```
### select the relevant HOGs:
a <- c("Brassica_napus")
succes_in_sample <- length(unique(
  Conserved_Selection_results_genes_pvalues_HOG_DE[Conserved_Selection_results_genes_pvalues_HOG_DE$HOG %
    in% Bnapus_vs_all_3_exp$HOG
                                                   & Conserved_Selection_results_genes_pvalues_HOG_DE$
    species %in% a
                                                   & Conserved_Selection_results_genes_pvalues_HOG_DE$
    selection_pvalue<=0.05,]
  $gene))
success_in_bg <- length(unique(
  Conserved_Selection_results_genes_pvalues_HOG_DE[Conserved_Selection_results_genes_pvalues_HOG_DE$
    species %in% a
                                                   & Conserved_Selection_results_genes_pvalues_HOG_DE$
    selection_pvalue<=0.05,]
  $gene))
sample <- length(unique(
  Conserved_Selection_results_genes_pvalues_HOG_DE[Conserved_Selection_results_genes_pvalues_HOG_DE$HOG %
    in% Bnapus_vs_all_3_exp$HOG
                                                   & Conserved_Selection_results_genes_pvalues_HOG_DE$
    species %in% a,]
  $gene))
bg <- length(unique(
  Conserved_Selection_results_genes_pvalues_HOG_DE[Conserved_Selection_results_genes_pvalues_HOG_DE$
    species %in% a,]
  $gene))

## perform the test for over representation
# select the proper sets to compare their sizes
  q <- succes_in_sample
  q_name <- paste0("Bnapus_vs_all_3_exp", "_success_in_sample")
  assign(q_name, q)

  m <- success_in_bg
  m_name <- paste0("Bnapus_vs_all_3_exp", "_success_in_BG")
  assign(m_name, m)

  n <- (bg - m)
  n_name <- paste0("Bnapus_vs_all_3_exp", "_non_success_in_BG")
  assign(n_name, n)

  k <- sample
  k_name <- paste0("Bnapus_vs_all_3_exp", "_sample")
  assign(k_name, k)

  p_name <- paste0("Bnapus_vs_all_3_exp", "_phyper")
  p <- phyper((q-1), m, n, k, lower.tail=F)
  assign(p_name, p)

  my_results <-c(round(q,2), round(k,2), round((q/k),2), round(m,2), round(n,2), round((m/(n+m)),2))
  print(paste0("Bnapus_vs_all_3_exp: ", p))
```

```
## [1] "Bnapus_vs_all_3_exp: 0.0745562529038334"
```

```
print(format(my_results, scientific = F))
```

```
## [1] "  544.0" " 5275.0" "    0.1" " 2732.0" "25230.0" "    0.1"
```

```
### select the relevant HOGs:
a <- c("Brassica_napus", "Eutrema_salsugineum")
succes_in_sample <- length(unique(
  Conserved_Selection_results_genes_pvalues_HOG_DE[Conserved_Selection_results_genes_pvalues_HOG_DE$HOG %
    in% Bnapus_and_Eutrema_vs_Arabidopsis_exp$HOG
                                                    & Conserved_Selection_results_genes_pvalues_HOG_DE$
    species %in% a
                                                    & Conserved_Selection_results_genes_pvalues_HOG_DE$
    selection_pvalue<=0.05,]
  $gene))
success_in_bg <- length(unique(
  Conserved_Selection_results_genes_pvalues_HOG_DE[Conserved_Selection_results_genes_pvalues_HOG_DE$
    species %in% a
                                                    & Conserved_Selection_results_genes_pvalues_HOG_DE$
    selection_pvalue<=0.05,]
  $gene))
sample <- length(unique(
  Conserved_Selection_results_genes_pvalues_HOG_DE[Conserved_Selection_results_genes_pvalues_HOG_DE$HOG %
    in% Bnapus_and_Eutrema_vs_Arabidopsis_exp$HOG
                                                    & Conserved_Selection_results_genes_pvalues_HOG_DE$
    species %in% a,]
  $gene))
bg <- length(unique(
  Conserved_Selection_results_genes_pvalues_HOG_DE[Conserved_Selection_results_genes_pvalues_HOG_DE$
    species %in% a,]
  $gene))

## perform the test for over representation
# select the proper sets to compare their sizes
  q <- succes_in_sample
  q_name <- paste0("Bnapus_and_Eutrema_vs_Arabidopsis_exp", "_success_in_sample")
  assign(q_name, q)

  m <- success_in_bg
  m_name <- paste0("Bnapus_and_Eutrema_vs_Arabidopsis_exp", "_success_in_BG")
  assign(m_name, m)

  n <- (bg - m)
  n_name <- paste0("Bnapus_and_Eutrema_vs_Arabidopsis_exp", "_non_success_in_BG")
  assign(n_name, n)

  k <- sample
  k_name <- paste0("Bnapus_and_Eutrema_vs_Arabidopsis_exp", "_sample")
  assign(k_name, k)

  p_name <- paste0("Bnapus_and_Eutrema_vs_Arabidopsis_exp", "_phyper")
  p <- phyper((q-1), m, n, k, lower.tail=F)
  assign(p_name, p)

  my_results <-c(round(q,2), round(k,2), round((q/k),2), round(m,2), round(n,2), round((m/(n+m)),2))
  print(paste0("Bnapus_and_Eutrema_vs_Arabidopsis_exp: ", p))
```

```
## [1] "Bnapus_and_Eutrema_vs_Arabidopsis_exp: 0.455140966997128"
```

```
  print(format(my_results, scientific = F))
```

```
## [1] "   34.0" "  341.0" "    0.1" " 4103.0" "38271.0" "    0.1"
```
